# FlyClockbase: Importance of Biological Model Curation for Analyzing Variability in the Circadian Clock of Drosophila melanogaster by Integrating Time Series from 25 Years of Research

**DOI:** 10.1101/099192

**Authors:** Katherine S. Scheuer, Bret Hanlon, Jerdon W. Dresel, Erik D. Nolan, John C. Davis, Laurence Loewe

## Abstract

Biological model curation provides new insights by integrating biological knowledge-fragments, assessing their uncertainty, and analyzing the reliability of potential interpretations. Here we integrate published results about circadian clocks in *Drosophila melanogaster* while exploring economies of scale in biological model curation. Clocks govern rhythms of gene-expression that impact fitness, health, cancer, memory, mental functions, and more. Human clock insights have been repeatedly pioneered in flies. Flies simplify investigating complex gene regulatory networks, which express proteins cyclically using environmentally entrained interlocking feedback loops that act as clocks. Simulations could simplify research further. We found that very few computational models test their quality directly against experimentally observed time series scattered in the literature. We designed FlyClockbase for integrating such scattered data to enable robust efficient access for biologists and modelers. To this end we have been defining data structures that simplify the construction and maintenance of Versioned Biological Information Resources (VBIRs) that prioritize simplicity, openness, and therefore maintainability. We aim to simplify the preservation of more raw data and relevant annotations from experiments in order to multiply the long-term value of wet-lab datasets for modelers interested in meta-analyses, parameter estimates, and hypothesis testing. Currently FlyClockbase contains over 400 wildtype time series of core circadian components systematically curated from 86 studies published between 1990 and 2015. Using FlyClockbase, we show that PERIOD protein amount peak time variance unexpectedly exceeds that of TIMELESS. We hypothesize that PERIOD’s exceedingly more complex phosphorylation rules are responsible. Variances of daily event times are easily confounded by errors. We improved result reliability by a human error analysis of our data handling; this revealed significance-degrading outliers, possibly violating a presumed absence of wildtype heterogeneity or lab evolution. Separate analyses revealed elevated stochasticity in PCR-based peak time variances; yet our reported core difference in peak time variances appears robust. Our study demonstrates how biological model curation enhances the understanding of circadian clocks. It also highlights diverse broader challenges that are likely to become recurrent themes if models in molecular systems biology aim to integrate ‘all relevant knowledge’. We developed a trans-disciplinary workflow, which demonstrates the importance of developing compilers for VBIRs with a more biology-friendly logic that is likely to greatly simplify biological model curation. Curation-limited grand challenges, including personalizing medicine, critically depend on such progress if they are indeed to integrate ‘all relevant knowledge’.

**General Article Summary:** Circadian clocks impact health and fitness by controlling daily rhythms of gene-expression through complex gene-regulatory networks. Deciphering how they work requires experimentally tracking changes in amounts of clock components. We designed FlyClockbase to simplify data-access for biologists and modelers, curating over 400 time series observed in wildtype fruit flies from 25 years of clock research. Substantial biological model curation was essential for identifying differences in peak time variance of the clock-proteins ‘PERIOD’ and ‘TIMELESS’, which probably stem from differences in phosphorylation-network complexity.

We repeatedly encountered systemic limitations of contemporary data analysis strategies in our work on circadian clocks. Thus, we used it as an opportunity for composing a panoramic view of the broader challenges in biological model curation, which are likely to increase as biologists aim to integrate all existing expertise in order to address diverse grand challenges. We developed and tested a trans-disciplinary research workflow, which enables biologists and compiler-architects to define biology-friendly compilers for efficiently constructing and maintaining Versioned Biological Information Resources (VBIRs). We report insights gleaned from our practical clock research that are essential for defining a VBIRs infrastructure, which improves the efficiency of biological model curation to the point where it can be democratized.

*Statement of data availability:* Stabilizing Versioned Variant of this file: **QQv1r4_2017m07d14_Lion** Before final publication *FlyClockbase* will be at **https://github.com/FlyClockbase** For review purposes *FlyClockbase* QQv1r4 will be provided as a zip-archive in the uploaded Supplemental Material; it is also available upon request from L. Loewe.

*Abbreviations:* Table 1: Molecular core clock components Table 2: Concepts for organizing FlyClockbase

*Supplemental Material:* Appendix: Supplemental Text and Tables (32 pages included in this file, QQv1v4) Supplemental Statistical Analysis (87 pages not included in this file, QQv1v4) R-Script zip file (>12K lines not included in this file, QQv1v4) FlyClockbase zip file (available upon request, QQv1v4)

## INTRODUCTION

Several grand challenges of our time, such as personalizing medicine or mapping genotypes to phenotypes, critically depend on the careful curation of biological knowledge-fragments into integrated resources of intermediate size that are easier to handle. Such resources can provide comprehensive overviews of integrated models or experimental results on a given topic. If these resources are organized well enough and are machine readable, then biological models and datasets can be explored in more automated ways and thus greatly accelerate biological discovery.

We aim to make it easier to create such intermediate resources by improving the efficiency of high-quality biological model curation. The goal of such curation work is to integrate ‘all current knowledge’ that is relevant for a research topic of reasonable complexity *while* keeping all deposited information well-organized and machine readable. This requires computational solutions that are best developed *while simultaneously* engaging deeply with the complexities of real-world biological research where results can be less than clear-cut and relevant data may be scattered across diverse sources. Curating such diffuse and scattered data can be prohibitively complicated without appropriate strategies for handling recurrent problems.

We chose the study of circadian clocks in fruit flies as our area of in-depth biological research in order to provide a real-world context for developing strategies that improve curation efficiency. While climbing onto the shoulders of giants in fly clock research, we integrated as much fly clock expertise as we could. In our opinion, a full integration is currently far beyond the scope of any single study if it is to efficiently point readers to the detailed, evidence-based evaluations of the strengths and weaknesses of a state-of-the-art fly clock model. Thus, we focused on integrating all *Drosophila melanogaster* wildtype time series observations from 25 years of research (often reported as wildtype control experiments for evaluating effects of mutants). Despite this substantial reduction of scope, our integration task is far from trivial if we aim to ensure the reproducibility, stability, and rigor of integration.

Reproducibility is pivotal for science. It also does not come easy. We aimed to increase the reproducibility of results from our research in fly clock biology while exploring strategies for simplifying reproducibility in research. Our main biological findings are differences in variances of certain time series traits that we observed between different clock components. We hypothesize that these differences hold important clues for improving our mechanistic understanding of circadian clocks. Variances are easily affected by errors that also affect reproducibility and are independent from underlying biological mechanisms. Therefore, we deem it essential to include our progress towards reproducibility in the scope of this study. We mitigate the inevitable increase of length with headings that simplify navigating its various aspects.

Our study contributes to the foundations of a system for integrating all expertise on the fly clock. We provide detailed experimental observations of time series ready for linking to statements in ‘big-picture’ fly clock models. We simultaneously explore how to integrate more efficiently the underpinning diffuse and scattered data. We find that such integration work is best accomplished by biological model curators with a deep biological interest in the research results that are being integrated. We also find that the efficiency of integrating and curating results can be greatly increased by access to strategies and tools designed to handle complex biological observations. We note that current computational tools repeatedly restrict representation possibilities to options that *almost* fit observed data – but not *entirely*. Such cases force model curators to inappropriately ignore data or subtly bias results by defining the closest representation as ‘good enough’. We ask if well-known data structures and logic formalisms originally developed outside of biology can appropriately capture real-world observations in biology. If not, is there merit in breaking the mold? Can economies of scale be leveraged in model curation? Abstraction is critical. If solutions are too abstract, they work everywhere – albeit poorly; if not abstract enough, they work perfectly for *one* problem – but force reinventing the wheel next time. To guide our development of computational abstractions, we find it essential to constantly face the challenging complexities of real-world experimental data as we work to advance biological research in circadian clocks of flies. This is where useful abstractions emerge naturally.

Accordingly, this study has three strands: (i) introduce FlyClockbase, the new resource we produced, and measure its reliability; (ii) present new biological insights on clocks in flies from analyzing data in FlyClockbase; (iii) evaluate emergent opportunities for abstraction as seen by a programming language compiler architect aiming to improve the efficiency of navigating the tension between the clear-cut logic formalisms in computers and the uncertain, incomplete and noisy biological data. We found that all three strands significantly strengthened each other. Each presents a distinct view on the integrated body of trans-disciplinary research presented here.

**Circadian clocks** are biochemical pathways characterized by cyclical protein expression. They play a critical role in a wide variety of behavioral and physiological processes, and a better understanding of their genetic and biochemical bases could advance research in many areas (Preussner and Heyd 2016; Sharma *et al*. 2016), including consciousness and sleep (Cirelli 2009), feeding and metabolism (Xu *et al*. 2008; Hurley *et al*. 2016), learning and memory (Xu *et al*. 2008; Chouhan *et al*. 2015), stress and immunity (Dumbell *et al*. 2016), inflammation (Carter *et al*. 2016), cancer (Sephton and Spiegel 2003; Masri *et al*. 2015; Salavaty 2015; Molina-rodrÍguez and Álvarez 2016), and psychological functioning (Mcclung 2013; Parekh *et al*. 2015; Coogan *et al*. 2016).

### Model organisms

Many model organisms have been used to study circadian rhythms, including *Synechococcus elongatus*, *Neurospora crassa*, *Arabidopsis thaliana, Mus musculus*, and *D. melanogaster* (Bell-Pedersen *et al*. 2005). Each model organism presents benefits and challenges. Here we focus on *D. melanogaster*, which is known for its ease of genetic manipulation (Stanewsky 2003; Özkaya and Rosato 2012) and its well-characterized genome (Dos Santos *et al*. 2015). Human circadian rhythms are certainly more complex than those of *D. melanogaster*. For example, it is not uncommon for one fly circadian clock component (e.g., *period* or *cryptochrome*) to correspond to multiple mammalian circadian clock components (e.g., *period1* and *period2*, or *cryptochrome1* and *cryptochrome2*) as reviewed elsewhere (Young and Kay 2001). Despite these differences in complexity, the *D. melanogaster* clock is similar in many aspects to the mammalian clock (Young and Kay 2001; Rosato *et al*. 2006). Insights from the fly clock have substantially contributed to understanding aspects of the mammalian clock in general and in particular related disorders such as familial advanced sleep phase syndrome (FASPS) (Rosato *et al*. 2006), pancreatic cancer (Pogue-Geile *et al*. 2006), and bipolar disorder (Ko *et al*. 2010; Mcclung 2013). Increased knowledge of the *D. melanogaster* circadian clock could continue to provide important information for future work in a variety of areas pertaining to the mammalian clock, including sleep disorders (Wager-Smith and Kay 2000), Alzheimer’s disease (Long *et al*. 2014), and psychiatric disorders (Mcclung 2013; Zordan and Sandrelli 2015).

### Math models

Computer simulations of mathematical models are powerful tools for studying the dynamics of complex non-linear systems such as circadian clocks. They have been used for decades in many disciplines (Kurtz 1972; Crosby 1973; Tarantola and Valette 1982; Ascher and Petzold 1998; Law and Kelton 2000; Zeigler *et al*. 2000; Tarantola 2005; Anderson 2007; Gillespie 2007; Gillespie 2008; Anderson *et al*. 2011; Karr *et al*. 2012; Mavelli 2012; Wilkinson 2012; Zeigler 2012; Distefano 2013; Gillespie *et al*. 2013; Sanghvi *et al*. 2013; Karr *et al*. 2014; Chylek *et al*. 2015; Karr *et al*. 2015a). To be useful for the study of circadian clocks, mathematical models need to mirror relevant aspects of real-world clocks, which may include key mechanisms, reaction rates, and/or other parameters or traits. Models integrate the specified details to enable simulations of time series of amounts of circadian clock components that are based on the assumptions of the *in silico* model. The simulated distributions of amounts of different clock components at specified times is expected to match observable real-world time series if a model’s assumptions are correct. Such simulations are facilitated by rigorous simulation algorithms that have a rich history in modeling biochemical reaction networks (Kurtz 1972; Gillespie 1977; Anderson 2007; Gillespie 2007; Gillespie 2008; Anderson *et al*. 2011; Karr *et al*. 2012; Mavelli 2012; Wilkinson 2012; Distefano 2013; Gillespie *et al*. 2013; Sanghvi *et al*. 2013; Karr *et al*. 2014; Chylek *et al*. 2015; Karr *et al*. 2015a).

### Estimating unknown rates from observed time series

If the structure of a model is essentially correct but its parameter values are not, then time series observed in the real world can, in principle, be used to narrow the margins of uncertainty around poorly-known rate parameters (Tarantola and Valette 1982; Tarantola 2005). Thus, access to a high-quality collection of observed time series could substantially contribute to improving the quality of biological insights gained from computational models. Mechanistic models with firm mathematical underpinnings can be used to explore hypotheses that are impractical to investigate in the laboratory for reasons that may include the effort required to produce mutants or the difficulty of measuring particular clock features experimentally (Loewe and Hillston 2008; Loewe 2009; Loewe 2016). By exploring potential hypotheses of interest *in silico*, models ideally inform future wet-lab experiments.

#### Biological example

The clock component *clockwork orange* (*cwo*) was thought to indirectly inhibit transcription of a number of key clock genes, including *period* (*per*), *PER-aryl-domain protein 1* (*pdp1*), and *vrille* (*vri*) (Kadener *et al*. 2007; Lim *et al*. 2007), so flies with decreased *cwo* expression were expected to show increased levels of *per*, *pdp1*, and *vri*. Experimental results, however, indicated that *cwo* mutants exhibited decreased expression of these clock components (Matsumoto *et al*. 2007; Richier *et al*. 2008). In an attempt to explain these results, Fathallah-shayk et al. (2009) created a clock model that included *cwo*. This model was able to predict the experimental results previously shown and was used to develop a novel hypothesis which described a more complex interaction between *cwo* and the rest of the clock. Rather than simply repressing a transcriptional activator, the authors of this model postulated that the interaction between weak repression by *cwo* and strong activation by the transcriptional activator led to indirect activation of a different part of the clock. This explained the experimental results from *cwo* mutants and suggested that *cwo* plays a role in reducing minor variations in the clock known as “jitters” ((Fathallah-Shaykh 2010); more details (Scribner and Fathallah-shaykh 2011)). This is just one example of the powerful ways in which modeling can act as a thinking tool, helping us to understand biology better.

#### Models and reality

Computational models simulations models are fundamentally attempts to represent a simplified version of reality, and their utility hinges on their ability to faithfully capture the most important aspects of reality (Tarantola and Valette 1982; Tarantola 2005). Complex processes with well-defined inputs and outputs are easily simplified by assuming that the timing of these processes remains essentially unchanged; then complicated sub-models of such processes can be substituted by simple transformations that merely reproduce the correct timing. If a model is to help understand, describe, or predict a biological system, such simplifications must be grounded in observations and a biological understanding of the phenomena to be modeled (Wooley and Lin 2005; Brodland 2015). For example, a model for predicting credible functions for *cwo* in the clock (Fathallah-Shaykh *et al*. 2009) needed to be able to replicate previous experimental results first, before it could make useful predictions. Such work is facilitated by the simplifying assumption that transcription, translation, and degradation in clock models can be replaced by simple reactions with rates appropriately chosen to match experimental data. Thus, access to a broad array of curated, high-quality experimental observations is critical for efficiently constructing and refining computational models.

### Reproducibility of research

The advanced mechanistic simulations and complex statistical inference methods above are necessary for arriving at a rigorous understanding of circadian clocks. They require a complex software stack and substantial efforts to implement dedicated code, workflows, and data organization schemes. It is not easy to develop scientific computing solutions of such complexity without loss of usability or sacrificing the reproducibility of earlier results. Yet the importance of reproducibility for science is undisputed and has recently received some attention (Ioannidis 2005b; Jasny *et al*. 2011; Huang and Gottardo 2013; Mcnutt 2014; Aarts *et al*. 2015; Freedman *et al*. 2015a; Allison *et al*. 2016; Baker 2016; Barba 2016; Lewis *et al*. 2016; Stodden *et al*. 2016). Reproducibility is an extremely broad topic that frequently requires input from many experimental, statistical, computational, theoretical and applied disciplines to arrive at rigorous solutions (Huang and Gottardo 2013). Conducting research reproducibly requires more effort than commonly realized (Donoho 2009; Stodden *et al*. 2014; James *et al*. 2015; Loewe 2016; Mesnard and Barba 2016). Yet, the steep upfront costs of entry seem to pay off: research teams with a reproducible research workflow report substantial benefits (Donoho 2009; Mesnard and Barba 2016). Evaluations of computational tools Such reports inspired us to work towards improving reproducibility in our efforts. While useful recommendations and tools exist e.g. see https://www.xsede.org/web/reproducibility and (Ince *et al*. 2012; Stodden *et al*. 2014; Poldrack and Poline 2015; Lewis *et al*. 2016; Stodden *et al*. 2016), there is no silver bullet and standards are still evolving. We aimed to keep computational requirements to a minimum.

#### Firm foundations

Here we cannot investigate the reproducibility of complex mechanistic circadian clock simulations or their underpinning parameter estimates. However, we can prepare a firm foundation for later studies. Rigorous reproducible reports of new parameter estimates need to provide many of the details reported here. This includes details on (i) literature database search strategies, (ii) initial screening processes and criteria, (iii) filtering of candidate studies and other special selection methods, (iv) reasons for combining some datasets but not others, (v) justifications for approaches that handle noise, outliers, or other complications, (vi) bibliographic references, (vii) a version of all raw data before preprocessing, and (viii) a well-integrated final version of the modified data used for inference after preprocessing. Ideally, such a study would include additional analyses like (ix) a human error analysis providing estimates of some low-level error rates quantifying the quality of internal raw data handling, (x) an error analysis and justifiable correction strategies for errors inherent to the raw data as received by this study, (xi) some high-level summary statistics description of the observed data, (xii) justifications and results from conducting various reasonable consistency checks, (xiii) reviews of reasonable biological interpretations of typical observations, outliers, or other patterns of interest, (xiv) critical assessments summarizing sufficient biological and other context to help readers evaluate thoroughly, skeptically, and efficiently how much trust is justified by the quality of the best and most complete dataset in this study, and (xv) any other potential limitations. Since circadian clocks in flies have been an active area of research for some time, a substantial number of studies report time series of potential interest. Time series of clock components are foundational for understanding circadian clocks. Appropriately integrating them raises various subtle issues that require decisions in order to build a strong foundation for further studies. We initially underestimated the complexity of dealing with their combined impact on the reproducibility of a steep data processing pyramid that aims to eventually integrate parameter estimation and biologically reasonable fly clock model ensembles. Since later steps such as parameter estimation cannot correct quality problems at earlier steps, we decided to dedicate this study to ensuring the availability of a durable high-quality set of time series observations ready to serve beyond this study. Such efforts rival the complexity of wet-lab experiments, except that they occur in a dry lab. The substantial investments in manual curation of high quality datasets are thus justified by the well-known GIGO principle that applies to simulations and experiments alike (Garbage In, Garbage Out). We next present some background on questions of basic reproducibility and data quality that arise for integration efforts at the scale of our study.

#### Problems with label reproducibility

Irreproducibility can be caused by seemingly trivial errors while executing deceptively simple work, such as pipetting errors (Broman *et al*. 2015) or (mis)labeling a line of descent in the lab (Lorsch *et al*. 2014; Freedman *et al*. 2015b). Assigning sequence annotations in GeneOntology databases is neither trivial nor always correct (Jones *et al*. 2007); incorrect assignments can replicate via uninformed users and can also be generated easily by using spreadsheets with inappropriate auto-conversion (Zeeberg *et al*. 2004; Ziemann *et al*. 2016). Activities like labeling or pipetting in array shaped micro-titer plates appear simple, but their simplicity is deceptive because they involve naming – an often-underestimated problem of extremely varying complexity (Loewe 2016). The stakes are high and have led to calls for systematically improving research at the bench and beyond (Collins and Tabak 2014; Lorsch *et al*. 2014; Allison *et al*. 2016). The impact of these problems on our study is immediate. We have little choice but to start with the assumption that *all errors had been corrected* by the time of publication, implying correctness of all name-related operations of all researchers involved in the production of the over 400 time series we report below. Time of publication matters, as authors and readers might struggle to get errors corrected if found later (Allison *et al*. 2016). The list of implications is long: no accidental swaps anywhere, neither in any fly strain used throughout its relevant history of descent, nor in the vials of final experiments, nor in the raw data, nor in averaging repeated observations, nor in labeling the final plots for publication. We appreciate that every single team of authors did their best to ensure that all errors were corrected in the final publication. Also, label-errors in published time series of clock components are probably less frequent than extrapolations of single-person initial error rates might suggest (assuming scrutiny from co-authors and peer review). However, human error analyses performed over decades in very diverse disciplines and for tasks of varying complexity have quantified in numerous experiments that “*to err is human*” (Panko 2016). Measured error rates observed in one type of experiment do not easily transfer to other contexts, but the existence of labeling errors in labs is well documented (Lorsch *et al*. 2014; Broman *et al*. 2015). Thus, it would be surprising if not a single error existed in the published time series data we integrated. Equally, it would be surprising if such a complex set of diffuse and scattered data could be integrated without adding a single error from data handling. These observations highlight the importance of assessing error rates and providing a defined protocol for reducing data handling errors. Thus, high-quality data curation requires (in reverse order) mature strategies for efficiently

- monitoring and handling all relevant error types,
- defining data structures that enable true data integration (and efficient querying)
- collecting all relevant scattered data in one place (and pre-sort for integration).

All this requires substantial efforts and biological model curators could probably learn from the substantial methodologies for human error analysis that have been developed elsewhere (NASA *et al*. 2006-07; NASA *et al*. 2011). Some of these approaches are too complex for the application to individual studies in biology. However, meta-analyses aiming to draw conclusions from noisy biological data need to find a way of handling the errors that occur during data handling. They also have to address reproducibility in the domain of statistics.

#### Statistical reproducibility

A substantial fraction of recent problems with reproducibility is caused by a lack of statistical reproducibility (Aarts *et al*. 2015; Halsey *et al*. 2015; Stodden 2015). These problems easily arise while designing experiments or analyzing data without the necessary statistical background (Salsburg 1985; Vaux 2012). Here, interpretations of ‘necessary’ are the subject of much discussion (Sterne 2003; Cumming 2013; Sharpe 2013; Leek and Peng 2015) as guidelines on statistical best practices are being updated (Altman *et al*. 1983; Millis 2003; Plowman 2008; Mazumdar *et al*. 2010; Cumming 2013; Drummond and Vowler 2013; Johnson 2013; Cumming 2014; Huang *et al*. 2015; Savalei and Dunn 2015; Trafimow and Marks 2015; Woolston 2015; Taroni *et al*. 2016; Wasserstein and Lazar 2016), but not necessarily followed everywhere (Lew 2012; Tressoldi *et al*. 2013). For example, a short-sighted over-reliance on *P*-values easily generates irreproducible or misleading results (Lew 2012; Cumming 2013; Nuzzo 2014; Halsey *et al*. 2015; Leek and Peng 2015), a criticism with history (Loftus 1993; Sterne 2003). Briefly, *P-*values are the probability that an observation can be explained by a given null-hypothesis, which usually represents the ‘most boring explanation’. Thus, *P*-values are often seen as ‘null-hypothesis significance tests’, but they do not make any statements about alternative hypotheses, of which there could be many. Yet researchers often use *P*-values to draw unsafe conclusions of deceptive simplicity about their respective favorite alternative hypotheses (Lew 2012; Tressoldi *et al*. 2013). They do so with such regularity that this error’s pervasiveness might one day motivate a fascinating human error analysis. *P*-values may offer substantial attractions as they combine the apparent reassurance of a precise number, the obvious simplicity of a single dimension, and the clear choice between a boring and a seemingly interesting option. In comparison, careful time-consuming analyses might be less appealing as they often reveal complex ensembles of less-than-clear-cut alternatives in a world of multi-dimensional trade-offs, requiring qualitative reasoning to decide which quantitative methods to use for producing precise numbers. Such analyses offer more nuance, albeit at greater cost and require more expertise in advanced statistics (Wilcox 2012), and aspects of type systems (Pierce 2002), semantics, and naming (Loewe 2016). These complex analyses underscore a conclusion that is intuitively well understood: biology does not present itself in a black and white picture of only interesting or boring parts. Instead it offers not only shades (allowing for gradients in addition to cutoffs at significance thresholds), but also colors (additional dimensions that otherwise might be inappropriately collapsed into a single dimension). The recent interest in statistical reproducibility has produced guidelines that recommend a closer look at some of these additional dimensions by estimating confidence intervals and other measures instead of testing arbitrary significance thresholds (Killeen 2005; Nakagawa and Cuthill 2007; Curran-Everett 2009; Cumming 2013; Cumming 2014; Demidenko 2016). This does not mean that *P-*values have no merit (Murtaugh 2014; Stanton-Geddes *et al*. 2014) and hence a pragmatic approach might be most appropriate (Boos and Stefanski 2011), if the high variability of *P*-values is accounted for (Halsey *et al*. 2015). In either case, close attention to the robustness of statistical methods is warranted (Wilcox 2012), and any statistical conclusions should be supported by some analysis of their statistical reproducibility (Halsey *et al*. 2015). Finally, showing more raw data is preferable (Loftus 1993; Drummond and Vowler 2011).

#### Statistical error iceberg

Recent interest in statistical reproducibility has drawn attention to other aspects of statistical analysis workflows. In this context, *P*-values have been described as the tip of the iceberg (Leek and Peng 2015). To arrive at fully rigorous conclusions requires investigating numerous detailed decisions about which data to include, which outliers to remove, which tests to use, and which simplifying assumptions to employ. Such analyses are more complex to produce and read, but they are currently essential for exploring the most efficient approaches for arriving at reliable statistical results. The fundamental nature of time series data for understanding clocks in flies motivated us to invest in corresponding statistical reliability. Therefore, we explore below several alternative ways to construct the statistical analysis pipeline for this study.

### Reproducibility in genetics

The recent surge of interest in reproducibility has resulted in a number of studies of additional relevance to FlyClockbase. For example, the reproducibility of genotype-phenotype associations has been investigated (Nci-Nhgri Working Group on Replication in Association Studies *et al*. 2007; Ioannidis *et al*. 2009b; Janssens *et al*. 2009; Kraft *et al*. 2009). Analysis of gene expression are an important tool of genetic analysis and have therefore seen substantial standardizing efforts (Bammler *et al*. 2005). Analysis of standardized and non-standardized measurements have found improved reproducibility when standardized experimental protocols were used (Bammler *et al*. 2005). For independent studies collected from the literature, repeatability of microarray gene expression analyses has met limited success (Ioannidis *et al*. 2009a). Reasons for failure included the unavailability of data, incomplete annotations, and missing documentation on data processing (Ioannidis *et al*. 2009a). Other relevant observations that can hamper reproducibility include pipetting errors (Broman *et al*. 2015) and pedigree errors (Broman 1999).

### Versioned Biological Information Resources (VBIRs)

The importance of biological information resources is undisputed and has motivated the construction of hundreds of heterogeneous resources as reviewed elsewhere (Brooksbank *et al*. 2005; Ng *et al*. 2006; Laibe and Le Novere 2007; Wierling *et al*. 2007; van Gend and Snoep 2008; Sullivan *et al*. 2010; Drager and Palsson 2014; Najafi *et al*. 2014). Sizes vary, as do scope and topics ranging from general (e.g. https://datascience.nih.gov/commons; https://kbase.us), to organism specific (e.g. http://flybase.org organized around *Drosophila* genomes (Gramates *et al*. 2016; Marygold *et al*. 2016)), modeling specific (Le Novere *et al*. 2006; Chelliah *et al*. 2015), approach specific (Cusick *et al*. 2009), and down to pathway or molecule specific resources (e.g. ClotBase (Sonawani *et al*. 2010) or SwissLipids (Aimo *et al*. 2015)). Their heterogeneity remains a challenge and motivated development of the FAIR Principles (Wilkinson *et al*. 2016). The FAIR Principles were designed for evaluating credible solutions for the problem of exchanging data in biology and emphasize important principles that make data sharing FAIR and data Findable, Accessible, Interoperable, and Reusable (Wilkinson *et al*. 2016). The FAIR Principles do not aim to provide any standards or implementations, but leave the actual development of solutions to others, such as the proposed standard for Minimal Information Requested In the Annotation of biochemical Models (MIRIAM) (Laibe and Le Novere 2007), Brief, Explicit, Summarizing, Technical (BEST) Names (Loewe 2016), and the many proposals reviewed elsewhere (Drager and Palsson 2014). Overall solutions will need to solve the extraordinarily difficult challenge of data integration (Doan *et al*. 2012). Accordingly, efforts to exchange detailed data more efficiently in these complex contexts have become top priorities in biological research contexts (NIH *et al*. 2012; Drager and Palsson 2014; NIH 2015; NIH 2016; Wilkinson *et al*. 2016).

#### Importance of versioned data integrators

Versioned interoperable information resources of intermediate size are likely to play a permanent role as hubs of integration for the biological expertise in an area. The versioning is important to enable users to access a stable state of information, without the unrealistic demand that these resources have reached their final stage of development. Any *VBIR* worth developing will likely be updated and improved for an extended period of time. While rates of such change are likely to vary substantially, none of these changes should imperil the reproducibility of some result that is based on the earlier state of the resourced as accessed by the authors of that result. How to achieve interoperable long-term stable versioning that is flexible enough to accommodate the broad range of needs of resources as heterogeneous as *VBIR*s is an open question for research in the semantics of naming. Currently, innumerable, incompatible, and inconsistent versioning systems are actively used by numerous projects. Integrating them without reflection would create a system that is almost incomprehensible and inflict on users intolerable amounts of inessential complexity. Experience has shown that such a complex system would be very brittle and would jeopardize reproducibility by its complexity. However, the value of consistent easily reproducible integration is in the quality to which usable resources offer expert curated relevant data that is continuously updated. Updates could be triggered by detecting errors or integrating future experiments, possibly expanding scope or precision through improved data models (but always increasing some versioning number). Managing such updating processes works best for *VBIR*s of some intermediate size. Thus, *VBIR*s size is defined at the lower end by exceeding the limited scope of single publications, reviews, or meta-analyses that are all frozen in time once completed. At their typical size, *VBIR*s enable the functional, ongoing integration of information evaluating multiple studies and reviews from the perspective of a well-defined scope. At the upper end, *VBIR*s generally remain at a much lower complexity than grand challenges, and thereby avoid many additional complications caused by their excessive complexity. This intermediate size and their stability enables *VBIR* to act as reliable building blocks for accumulating biological expertise and address existing grand challenges more efficiently.

#### Flexibility of VBIRs

These minimal constraints allow *VBIR*s to take on a great diversity of organizational forms and the size of their scope may vary widely. However, not all biological information resources currently perceived as useful do satisfy these requirements. For example, not all repositories and databases in biology that aim to continually integrate information provide reliable access to clearly defined states in the past that are easy to access and to cite. Traditionally, the biological information resources that are easiest to cite are journal articles, but these do not usually provide information in a form that is structured enough for further processing and they do not usually update information (Allison *et al*. 2016). Some authors complement their articles with online databases or more static resources that can contain valuable material. However, the lack of standards and tools that are easy to use means that such efforts usually require substantial programming and data science expertise when they are set up and when they are to be maintained. Such barriers of entry make it very difficult for non-programming biologists with interesting datasets to set up and publish a VBIR in a form that facilitates further data integration.

*Database integration* is a special case of data integration. Both are enormous general challenges, whenever non-trivial datasets are to be used together (Doan *et al*. 2012). For example, a substantial research collaboration worked towards integrating data scattered across 81 geospatial temporal ecology datasets from 7 provider types in an effort to build LAGOS, the LAke multi-scaled GeOSpatial & temporal database (Soranno *et al*. 2015). The substantial supporting online material of the initial LAGOS description (Soranno *et al*. 2015) provides an impression of the numerous data-handling and type system synchronization challenges the LAGOS team had to face in order to obtain some state of data integration (see http://csilimno.cse.msu.edu/lagos_status.php for updates).

*Cochrane reviews* provide a completely different approach to data integration. To reduce arbitrary bias that easily arises in more limited non-systematic reviews, http://www.cochranelibrary.com aims to stimulate the use of systematic methods for finding and integrating all peer-reviewed information about a given topic. The resulting retrospective and prospective meta-analyses have substantially advanced the integration of biomedical observations. Ongoing development of methodologies for systematic bias reduction has greatly increased awareness and approaches available for reducing the influence of important biasing factors (Tharyan 1998; Ades *et al*. 2008; Mckenzie *et al*. 2013; Doi 2014; Onitilo 2014; Debray *et al*. 2015; Efthimiou *et al*. 2016). This success does not imply that further improvements are impossible; there are biases that are notoriously difficult to address, such as certain types of ascertainment bias (Amos *et al*. 2003; Clark *et al*. 2005; Lachance and Tishkoff 2013; Minikel *et al*. 2014) and biases against the publication of negative results (Johnson and Dickersin 2007). In fact, it can be argued that biases are almost everywhere (e.g.: Ioannidis 2005a; Patil *et al*. 2015). Thus, quantifying biases and analyzing their impact appropriately may be more important than demanding their absence. Cochrane reviews are not VBIRs, because traditional publications cannot be updated regularly. If VBIRs could be published easily, without the currently required database programming overheads, then Cochrane reviews with a reasonably well-defined data model could in principle become VBIRs.

### Genome projects as a model for VBIRs development on a broader scale

*VBIR*s increase the speed of hypothesis testing and greatly add to the long-term value of properly annotated wet-lab data. They offer the raw material for diverse meta-analyses, opening up entirely new research perspectives. In this respect, *VBIR*s mirror similar efficiencies known from genome projects. The similarities do not end here. *VBIR*s development also shows similar strong dependencies on software and data organization. Genome projects demonstrated efficiencies of scale by separating data collection and various stages of data interpretation (Lander *et al*. 2001; Venter *et al*. 2001). The efficiency of post-genomic sequencing workflows critically depends on the development of appropriate data structures, processing tools and exchange protocols (Wilkinson *et al*. 2016). We expect similar boosts to efficiency from developing *VBIR*-tools. For example, they could support more biology-friendly data structures to increase the efficiency and precision of integrating inherently imprecise biological observations. They could also greatly accelerate the adoption of sophisticated statistical analysis by the biological community, simply by implementing the appropriate statistical methods into the corresponding automated workflows. This would reduce problems of confusing SD and SEM (Salsburg 1985), and opens up a new avenue for communicating recommended best practice for statistical analyses, that could be provided right next to a user-friendly implementation (Mazumdar *et al*. 2010; Sharpe 2013).

#### Genome projects have revolutionized biology

Here we want to explore whether efficiencies of scale in biological model curation organized in a VBIRs project might hold a similar potential for accelerating the pace of biological discovery. The success of genome projects and other targeted efforts has been built on efficiently organizing and exchanging new data and interpretations (Wilkinson *et al*. 2016). For VBIRs exchanging data is more complicated, because their data is more structured and more diverse than typical genome data. Accordingly, efforts to exchange details data more efficiently in these complex contexts have become top priorities (NIH *et al*. 2012; Drager and Palsson 2014; NIH 2015; NIH 2016; Wilkinson *et al*. 2016). Such work is essential for progress towards meeting increasingly complex grand challenges like personalizing medicine, constructing genotype-phenotype-maps, or predicting how cancer cell populations evolve using mechanistic models in evolutionary systems biology (Loewe 2016). By definition, these grand challenges all exceed the problem-solving skills of any single research unit, and therefore critically depend on the efficient communication of the latest progress. This progress could be captured in high-quality *VBIR*s that probably will bring the same efficiency benefits as genome projects, once corresponding tools become available for dealing with their more diverse data types.

### Importance of biological model curation

None of the benefits above can be realized without substantial human input in the form of biological model curation. Despite many advances in machine learning, the gold standard for data curation is still the eye of a domain expert. While machines are extraordinary in exploiting regular patterns, it has been difficult to teach machines how to correctly handle the many exceptions that are readily recognized by human experts as deviations from ‘common sense’ (Burkhardt *et al*. 2006; Salimi and Vita 2006). Biological model curation by itself is not new. In fact, one could argue that very essence of research is the construction and curation of biological expertise that could also be described as a model. Thus, the curation of biological information is at least as old as Linnaean taxonomy (LinnÉ 1758). None of the recent resources that systems biology depends on could have been put together without substantial model curation efforts (see e.g. Drager and Palsson 2014). The work of biological model curators, or biocurators, has only recently come into focus as an increasingly important avenue of biological research (Bourne and Mcentyre 2006; Burkhardt *et al*. 2006; Salimi and Vita 2006; Howe *et al*. 2008; St Pierre and Mcquilton 2009; Bateman 2010; Burge *et al*. 2012; Hirschman *et al*. 2012; Zhang *et al*. 2014b; Mitchell *et al*. 2015; Orchard and Hermjakob 2015; Rodriguez-Esteban 2015; Gibson *et al*. 2016; Kim *et al*. 2016; Reiser *et al*. 2016; Singhal *et al*. 2016). By now biocurators have an international society (http://biocuration.org) and an official journal (http://database.oxfordjournals.org). Funding for digital depositories in biology has historically been complicated because few have realized the essential contributions of biological model curators to the overall scientific enterprise. This problem has been recognized and efforts are underway to address this discrepancy (Ember *et al*. 2013). One potential contribution to these efforts could be to find a way that substantially reduces the cost of initiating, growing, and maintaining *VBIR*s. We explore a potential approach for simplifying model curation by exploiting advances in computer science that have greatly simplified the design and construction of compliers for new programming languages.

### How a compiler could help in biological model curation

Programing language compliers are extraordinary efficient tools for guaranteeing that a given collection of texts (source code) conform to a well-defined standard (the complier’s language) and are transformed into output that is guaranteed to conform to strict rules. The construction of compliers requires an advanced understanding of computer science, but decades of research have produced a substantial body of data structures, algorithms, and tools that greatly simplify the construction of compilers today (e.g. Cooper 2012; Grune 2012). Thus, the question today is not if a compiler can be constructed for a given language, but rather what should a language look like for which it is worth constructing a compiler. After the construction of uncounted programing languages each with their own strengths and weaknesses, rather compelling reasons are necessary for creating a new one and any such efforts should learn from the diverse shortcomings of their many predecessors (Mandrioli and Pradella 2015). Traditional approaches to designing new programming languages have not included the very large amounts of feedback from research biologists that are necessary for creating a language design that could efficiently support biologists in their work (Loewe 2016).

#### Enforce best practices

For example, such a compiler could greatly reduce the confusion between SD and SEM that has plagued the reporting of biological results for some time (Salsburg 1985). A compiler that implements the latest statistical testing methods could greatly improve the adoption of statistical best practices in the biological research community (Mazumdar *et al*. 2010; Sharpe 2013). Such a compiler could also advance standards for facilitating interoperability in systems biology (Drager and Palsson 2014) and thereby contribute towards solving the extraordinarily difficult challenge of data integration (Doan *et al*. 2012), improve the semantic reproducibility of biological data (Loewe 2016), facilitate the sharing of meaningful data based on the FAIR Principles (Wilkinson *et al*. 2016), and encourage biologists to provide the Minimal Information Requested In the Annotation of biochemical Models (MIRIAM) (Laibe and Le Novere 2007). VBIRs construction does not require the existence of such a compiler, as every task can also be performed manually. Manual work is slower, but also more flexible, and can therefore better attend to the needs of high-quality biological model curation of a given data set. If such curation work is combined with the perspective of a compiler architect, then it provides extraordinary opportunities for designing efficient abstractions for data structures and tasks that can later be supported by a fully automated compiler.

#### Efficiencies of scale

Many VBIRs are likely to have similar needs that can be served by the same compiler if they share a standard for storing data. Thus, the costs of compiler development can then benefit several *VBIR*s where they reduce the cost of *VBIR* development and maintenance, which have been difficult to fund (Ember *et al*. 2013). Compiler development is also an excellent opportunity for detecting problems in logic formalisms; such errors have the potential for causing exorbitant costs (e.g. Hoare 2009; Kamp 2011; Loewe 2016). Therefore, constructing such a compiler could already be a cost-effective decision for the longer-term development and maintenance of a single VBIR alone. We aim to observe during our biological research where commonly used logic formalisms made it more complicated to accurately represent biological observations with their usual uncertainty. Representing uncertain biological data in computational structures is a sufficiently frequent problem to cause substantial frustration in efforts towards curating biological models at a reasonably high quality.

### Opportunity

Our systematic study of circadian clock gene expression patterns offers intriguing opportunities for engaging with the timely questions of reliable data handling, control experiment repeatability, human error analysis, reproducible computing, statistical reproducibility, and the semantic reproducibility of source code in research computing. These questions might be easy to dismiss at first sight, but as discussed above, in the broader context of growing data sets in VBIRs, low rates of diverse and individually rare errors can combine into a pervasive fog of confusion that can render a valuable collection of scientific results unusable. Usually these problems cannot be investigated at the level of a single experimental study, but this does not imply that errors in *VBIR*s are rare, or without consequence (e.g. see (Zeeberg *et al*. 2004; Jones *et al*. 2007; Schnoes *et al*. 2009)). The resulting irreproducibility is not cheap. For example, non-clinical biomedical studies with an estimated cost of about $7Bn/yr throughout the US come with difficulties in data analysis and reporting that hamper their reproducibility (Freedman *et al*. 2015a). Pervasive biases in biological datasets (Ioannidis 2005a) and statistical difficulties that can lead to substantially wrong conclusions (Ioannidis 2005b) can interfere with scientific discovery. To address these problems, it is important to invest in efforts towards opening science (Bartling and Friesike 2014), sharing data (Packer 2016; Wilkinson *et al*. 2016), and improving reproducibility in various areas (Ioannidis 2005b; Donoho 2009; Huang and Gottardo 2013; Loewe and Keel 2014; Stodden *et al*. 2014; Freedman *et al*. 2015a; James *et al*. 2015; Stodden 2015; Barba 2016; Loewe 2016; Loewe *et al*. 2016; Loewe *et al*. 2017). Reproducibility frameworks greatly facilitate individual scientific research studies, but take much more effort to put into place than could possibly be expected of the investigators in any individual study. Therefore, it is important to find efficient ways to achieve these goals at institutional and national scales (NIH *et al*. 2012; NIH 2015; NIH 2016). Our study is different from many typical studies in that we have attempted to simultaneously conduct high quality research while working towards a framework for improving reproducibility.

### Purpose of this study

Here we interweave several perspectives integral to one body of trans-disciplinary research. We aim to improve amount and quality of experimental time series available for parameter estimation in mechanistic simulations along with our overall understanding of *D. melanogaster* circadian clock models. To this end we present FlyClockbase, a new carefully curated biological information resource designed to maximize accessibility and ease of use for experimental biologists and modelers. We show how to use it for testing hypotheses and report our own new findings about the variability of peak times in the clock. Finally, we make trans-disciplinary observations at the interface of experimental biology, data curation, reproducibility, and the applicability of logic formalisms in biology. We present our process for working towards constructing a compiler that would substantially reduce the effort required for developing and maintaining data resources like FlyClockbase. These perspectives are next explained in a more detailed overview.

#### Formal organization of Versioned Biological Information Resources (VBIRs)

FlyClockbase is a *VBIR*. *VBIR*s store biological information using controlled immutable versioning numbers for marking each publicly released variant of the resource to ensure that previously released data remains accessible under that number. As we developed FlyClockbase, we aimed to separate the special from the general *VBIR* aspects to help make our design more applicable to the development of future *VBIR*s. Below we discuss why more *VBIR*s are needed. For ease of use, quality control, and future maintainability, we designed FlyClockbase as a file-based data resource that follows a well-defined scheme for collecting tables in text files grouped in folders. This organization as a set of tables is conceptually similar to the structure of a relational database, albeit without the speed, rule enforcement, and other amenities provided by modern database systems. As a result, our approach maximizes flexibility and openness, while minimizing certain types of administrational and long-term maintenance costs (to increase chances of long-term survival; see Discussion). The resulting system is even more general than *VBIR*s and we nicknamed it ‘TabFS’. The name TabFS highlights the central role of tabs (for delimiting), table-files (for storing) and the file-system (for organizing data). To simplify the implementation of other *VBIR*s, we have been separating the specific details of implementing FlyClockbase from general abstract features. In TabFS we aim to capture the abstractions and rules required for implementing *VBIR*s with a long-term view to developing a reliable *VBIR* standard. Our goal is to provide the simplest and most efficient organization possible without sacrificing the flexibility curators need for defining new data types that represent diverse types of complex and uncertain biological observations. Efficiently integrating new datasets in FlyClockbase requires this flexibility. It facilitates focusing on clock biology and minimizes distractions from defining or decoding data types. This approach enables FlyClockbase to integrate a diverse array of wildtype and wildtype-like time series along with the attributes necessary for documenting a broad range of experimental details. Each time series records the relative amount of a clock component as observed at various points in time.

#### Data integrated by curation

In FlyClockbase we provide a curated overview of 25 years of published observations of *D. melanogaster* clock components, which we use for retrospective meta-analyses. The types of molecules reported in FlyClockbase are based on the biological *D. melanogaster* clock model we abstracted from the relevant literature (see Table 1 for brief descriptions of core clock components). FlyClockbase contains more than 400 time series curated from the wildtype control experiments of 86 circadian clock studies. They can be compared in many ways within or between clock components for testing diverse hypotheses of potential interest for fly clock research.

#### Hypothesis tested: biological variability

We use FlyClockbase for comparing the variance of times at which the circadian mRNAs and proteins *period* (*per*) and *timeless* (*tim*) reach their relative daily peak and valley. We find significant differences in variance that are not easily explained as a statistical fluke and survived several rounds of in-depth error checking (which led to interesting conclusions in their own right). Thus, we hypothesize that the larger variance of peak times for the protein PER in comparison to TIM might have mechanistic reasons that could help illuminate interesting aspects of the clock if recovered in mechanistic models.

#### Hypothesis tested: observation method

The confluence of many diverse independently observed time series in FlyClockbase provides a unique resource for understanding such variability of fly clocks in a broad range of settings, as documented in the attributes of FlyClockbase time series. This variability can also be used for comparing the reproducibility of different approaches to measuring time series. We compared time series measured by PCR based methods (qPCR, RT-PCR) with methods that do not include self-replication (Northern Blot, RNAse Protection Assay). The variability of PCR-based time series in FlyClockbase exceeds that of non-PCR based methods; though originally surprising, this is consistent with both the exponential nature of amplification in PCR and previous reports on the reproducibility of quantitative measurements from PCR-based methods.

#### Human error analysis

We measured human error rates for a given set of tasks in FlyClockbase. Our results are broadly comparable to previous observations. The findings suggest that VBIRs would benefit from developing methods for ensuring that scientific conclusions are not affected by human errors that inevitably occur when handling or analyzing data and corrupt content or type. Designing a formal type system capturing relevant expert insights for FlyClockbase could facilitate and ultimately automate searches for logical inconsistencies.

#### Compiler logic design

We developed FlyClockbase while simultaneously exploring design options for programming language compilers that could help construct and maintain *VBIR*s. We have identified numerous pivotal features for supporting the long-term stability of FlyClockbase that are most efficiently implemented by a correspondingly designed compiler. We discuss how these and other practical aspects of working with *VBIR*s can improve the usefulness and chances of longer-term survival for *VBIR*s. We use an analogy to well-known results from population genetics to illustrate what the future might hold for a newly-born *VBIR*, such as FlyClockbase. These considerations show that cumulative practical impacts from many small complications or innovations can be unexpectedly large. We illustrate using FlyClockbase how it can be difficult to represent uncertain biological data in the certainty-demanding logic formalisms of the data types commonly used in computational tools. Most of the data types we need in FlyClockbase fall into two categories. Some are very general data types that are very common and thus ideally designed for interoperability (e.g. bibliographic references, tables, etc., see TabFS). Other data types are specific to FlyClockbase and therefore are not reusable. These types need to be defined by those expert biologists who curate FlyClockbase, and best understand the relevant biology. To facilitate these discussions, we developed a trans-disciplinary collaboration model to help with the necessary communication of biological curators with a compiler architect, who needs to be capable of bridging biology and aspects of designing compiler logic. If such a communication approach is used by a compiler architect for informing important decisions about the relevant logic formalisms to be implemented, then our observations suggest that the efficiency of biological model curation could greatly increase once a user-friendly *VBIR* compiler becomes available. Such a compiler will empower biological model curators to define their own data types that can be shaped to more appropriately representing the uncertainty of the biological observations they curate – without violating biological or computer science logic. Such compilers will also allow biologists to define their own consistency checks, which can then be automatically maintained by a correspondingly designed compiler. If this compiler also implements sufficiently reviewed standards for interoperability and data exchange, biological research will benefit from an unprecedented ability to combine models and analyses from different VBIRs.

#### Importance of efficient biological model curation

Our work highlights why topic-specific *VBIR*s like FlyClockbase have an essential, irreplaceable role to play in biological research – once curated to high quality by expert biologists. As we illustrate in FlyClockbase, *VBIR*s increase the speed of hypothesis testing and greatly add to the long-term value of properly annotated wet-lab data. They offer the raw material for diverse meta-analyses, opening up entirely new research perspectives. In this respect, *VBIR*s mirror similar efficiencies previously observed in genome projects. Our trans-disciplinary analysis suggests that the most efficient route for integrating biological information requires more work on type systems and logic formalisms in order to better capture the many uncertainties regularly found in biological data. To sample the problem space well enough, more studies like ours are needed that report how in-depth biological research challenges the expressivity of logic formalisms that have become candidates for implementation in the discussed compiler. Combining such observations with a rigorous in-depth usability and expert review process as defined elsewhere (Loewe 2016) will greatly accelerate the definition of more appropriate logic formalisms, VBIRs compiler design, biological model curation, and thus progress towards meeting the grand challenges of our time. This new efficiency would indeed allow us to stand on the shoulders of giants and no longer have to start crawling upwards from the elbow whenever a new question arises.

#### Overview of Sections

In the next Section, we review biological and computation clock models, as well as the data model of FlyClockbase from a biological perspective. We then describe how we selected the data in FlyClockbase, how we processed time series and which statistical methods we used. Our Results Section first quantifies the historic use of direct experimental time series observations in modeling studies. It then reviews the number of time series observed for each core clock component in *Drosophila melanogaster*. It provides an overview of the variability in all components before presenting a human error analysis investigating potential impacts of data handling errors in FlyClockbase on the variances of peak and valley timings of the *period* and *timeless* gene products, which are compared after defining a basic null-hypothesis for data in FlyClockbase. Our last result compares methods for observing mRNA. We start our Discussion Section by explaining, how FlyClockbase facilitates hypothesis-driven research. We then discuss the two hypotheses tested in this study and suggest mechanistic models for further testing. Next, we broaden our view to discuss the importance of model curation for molecular systems biology data. We then highlight observations that illustrate, how a tool with the capabilities of a specially crafted programming language compiler could advance work in FlyClockbase and beyond. We will pay attention to aspects like efficiency, error detection, and formal logic. Given the high likelihood of long-term loss of biological information resources, we finally discuss population genetics modeling results with some applicability to the fate of FlyClockbase. We do so in order to prioritize and motivate the next steps. We conclude with a list of the various disciplinary areas engaged by this study and how biological model curation will facilitate critical progress towards various grand challenges of our time. Our online material includes additional text with a more computational perspective and a supplementary statistical analysis to which we frequently refer in our results (including R source and data).

## MODELS

### Biological model of fly circadian clocks

The *D. melanogaster* clock is a gene regulatory network that receives environmental inputs (such as light and temperature) and produces its hallmark cyclical behavior through various interlocking positive and negative feedback loops (Kuczenski *et al*. 2007; Wang and Zhou 2010). Table 1 lists the most important clock components, and Figure 1 represents their key interactions in the Systems Biology Graphical Notation (Moodie *et al*. 2011). The timing of various clock sub-processes is essential for any clock. Circadian clocks critically depend on generic cellular processes of importance in the information processing associated with proteins, such as transcription, translation, and degradation. Thus, mutations disrupting critical functions in these generic processes are also likely to affect the clock. However, they are also likely to have many other harmful consequences; hence, we do not consider them as *core* clock components (which are the exclusive focus of our study here).

**TABLE 1.** Circadian clock components referenced in this study as introduced by their Brief, Explicit, Summarizing and Technical Name equivalents defined by using the Evolvix BEST Names concept.

| **Brief Name** | **Explicit Name** | **Summarizing Name (description)** | **Technical Name FlyBase ID** |
| --- | --- | --- | --- |
| *clk* | clock | Promotes transcription of *per*, *tim*, *pdp1*, *vri*, *cwo* | FBgn0023076 |
| *per* | period | Binds TIM, inhibits CLK action | FBgn0003068 |
| *tim* | timeless | Binds PER, inhibits CLK action, degraded in response to light | FBgn0014396 |
| *cry* | cryptochrome | Activated by light, promotes TIM degradation | FBgn0025680 |
| *cwo* | clockwork orange | Stabilizes "jitters" (minor variations) in the clock | FBgn0259938 |
| *pdp1* | PAR-domain protein 1 | Promotes transcription of *clk* | FBgn0016694 |
| *vri* | vrille | Inhibits action of PDP1 | FBgn0016076 |
| *pdf* | pigment-dispersing factor | Synchronizes different clock neurons | FBgn0023178 |
| *sgg* | shaggy | Phosphorylates TIM, promotes nuclear localization | FBgn0003371 |
| *jet* | jetlag | F-box protein, contributes to TIM degradation | FBgn0031652 |
| *ck2a* | casein kinase 2 alpha | Phosphorylates PER, promotes nuclear localization | FBgn0264492 |
| *dbt* | doubletime | Phosphorylates PER, promotes degradation | FBgn0002413 |
| *nmo* | nemo | Phosphorylates PER, promotes further phosphorylation | FBgn0011817 |
| *slmb* | slimb | F-box protein, contributes to PER degradation | FBgn0283468 |
| *cyc* | cycle | Binds to CLK | FBgn0023094 |
| *cu, noc* | curled, nocturnin | Implicated in light response, metabolism, mRNA degradation | FBgn0261808 |

Brief Names and Explicit Names represent abbreviations and full common names, respectively. Summarizing Names provide a brief description, and Technical Names correspond to hyperlinked FlyBase Identifiers. In the text, we follow the convention of denoting the proteins in CAPITAL letters and mRNAs or genes in *italics* (hence the use of italics for the brief names here). More about the way synonymous dialects (defined by the Evolvix BEST Names concept) can help navigate biological naming complexity can be found elsewhere (LOEWE et al. (2014; 2016) and LOEWE et al. (2017) for a more in-depth discussion of naming).

**FIGURE 1.**
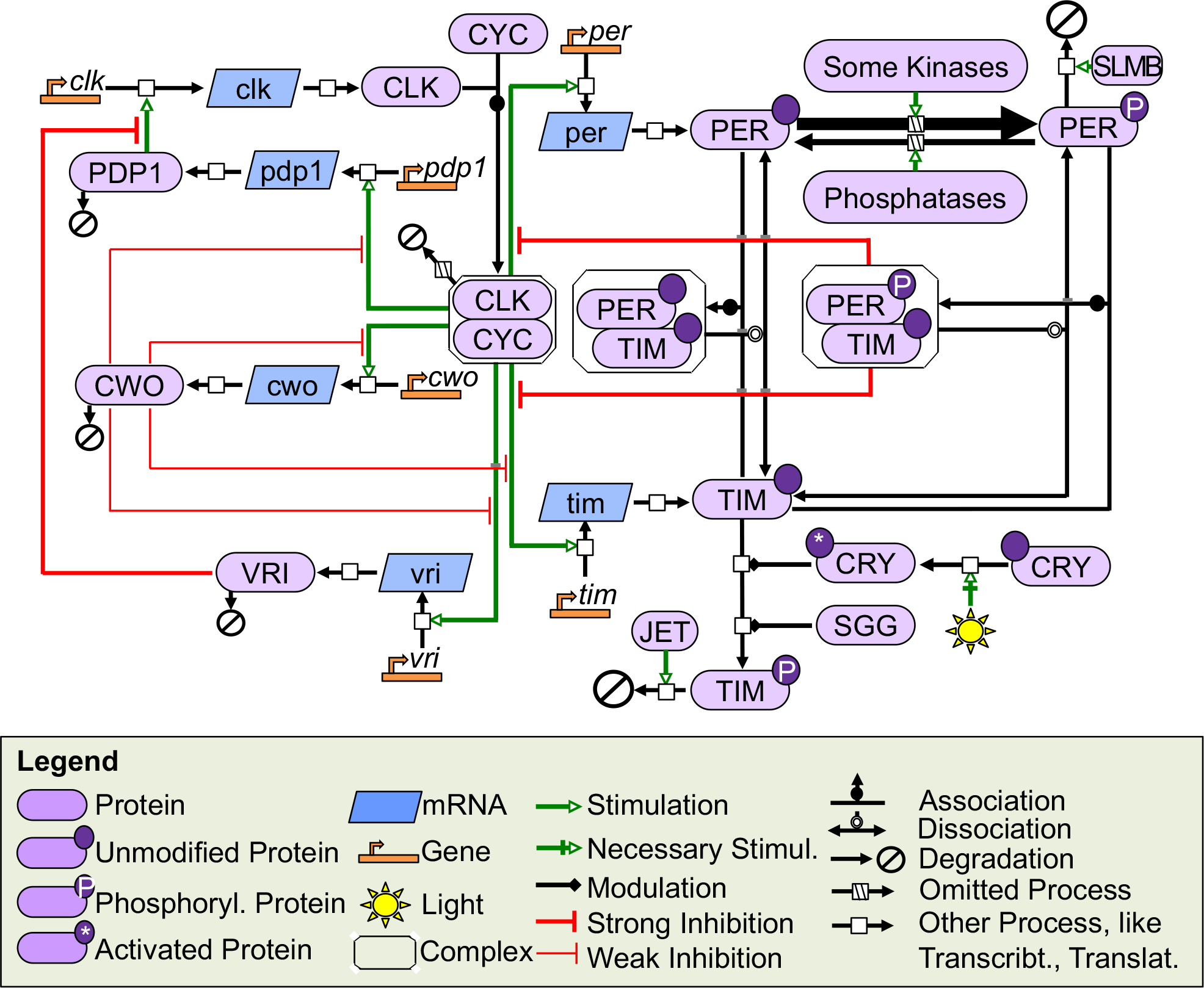
Systems biology overview of the circadian clock in *D. melanogaster*. This simplified diagram shows the basic components of the *D. melanogaster* circadian clock. In one feedback loop, CLK binds CYC and promotes the transcription of *per* and *tim*. The proteins PER and TIM then form a complex and repress this effect of CLK. Light activates CRY by inducing a conformational change (denoted by *), and CRY and SGG work with JET to promote the degradation of TIM. Without TIM, PER is phosphorylated, interacts with SLMB, and is degraded. In a second feedback loop, the CLK/CYC complex promotes transcription of *pdp1, vri, and cwo*. PDP1 promotes transcription of *clk*, and VRI inhibits the action of PDP1. CWO weakly inhibits CLK-promoted transcription of *per*, *tim*, *vri*, *pdp1*. and itself. The notation in this diagram uses Systems Biology Graphical Notation Process Description level 1 version 1.3 (Moodie *et al*. 2011), with minor modifications.

#### Main loop

Briefly, in the core (negative) feedback loop, the proteins CLOCK (CLK) and CYCLE (CYC) form a heterodimer and promote the transcription of *period* and *timeless* (Darlington *et al*. 1998; Rutila *et al*. 1998). PER protein is increasingly phosphorylated by DOUBLETIME (DBT) and several other kinases. Fully phosphorylated PER interacts with the F-box protein SLIMB (SLMB) to be marked for degradation unless TIM protein is present to form a PER/TIM complex. This complex represses the effects of the transcriptional activator CLK to form a negative feedback loop (Gekakis *et al*. 1995; Kloss *et al*. 1998; Lee *et al*. 1998; Price *et al*. 1998; Lee *et al*. 1999; Kloss *et al*. 2001; Chiu *et al*. 2008). When light is present, the protein CRYPTOCHROME (CRY) undergoes a conformational change that renders it active. As a final step before TIM degradation, activated CRY and the kinase SHAGGY (SGG) cause TIM in its phosphorylated form to interact with the F-box protein JETLAG (JET) (Martinek *et al*. 2001; Koh *et al*. 2006). If TIM is degraded thought its phosphorylated form, this will limit the formation of the PER/TIM complex. If this complex cannot form, then PER will be left in an isolated form in which it can be further phosphorylated (and thus be moved closer to its degradation). If PER pairs with TIM to form this complex, then PER cannot be phosphorylated, and it will temporarily stop its progress towards degradation. Thus degrading TIM facilitates the degradation of PER by allowing PER to become fully phosphorylated (Emery *et al*. 1998; Naidoo *et al*. 1999; Busza *et al*. 2004; Ozturk *et al*. 2011). Consequently, PER/TIM complexes no longer repress CLK transcriptional activity, and CLK proceeds to start a new cycle by again promoting again the transcription of *per* and *tim*.

#### Other loops

A second feedback loop primarily concerns *clk* transcription. CLK promotes transcription of both *PAR-domain protein 1 (pdp1)* and *vrille (vri)* (Blau 1999; Mcdonald and Rosbash 2001; Cyran *et al*. 2003). PDP1 protein then promotes the transcription of *clk*, while VRI represses the activity of PDP1 and inhibits the transcription of *clk*, creating positive and negative feedbacks, respectively (Cyran *et al*. 2003; Glossop *et al*. 2003). The relatively recently discovered *clockwork orange* (*cwo*) modulates both feedback loops by weakly repressing CLK-mediated transcription of *per*, *tim*, *pdp1*, *vri*, and *cwo* itself (Kadener *et al*. 2007; Lim *et al*. 2007; Matsumoto *et al*. 2007; Richier *et al*. 2008). The interplay between strong transcriptional activation promoted by CLK and weak repression from CWO protein counteracts “jitters,” or small variations in period (Fathallah-Shaykh *et al*. 2009; Fathallah-Shaykh 2010; Scribner and FATHALLAH-SHAYKH 2011). Other notable circadian products involved with post-translational modification, synchronization of clock neurons, and other processes include *casein kinase 2 alpha (ck2a*), *protein phosphatase 2a (pp2a*), *pigment-dispersing factor (pdf)*, *nemo (nmo)*. and others (Grima *et al*. 2002; Ko *et al*. 2002; Lin *et al*. 2002a; Sathyanarayanan *et al*. 2004). A more detailed review of the clock can be found elsewhere (Hardin 2011; Özkaya and Rosato 2012).

### *In silico* models integrating fly clock observations

Mathematical models of circadian clocks have been contributing to our understanding of clock biology for decades.

#### Biological results overview

The origins of many fly clock models can be traced back over 50 years to work by Pittendrigh & Victor (Pittendrigh and Victor 1957; Goodwin 1964; Goodwin 1965) and Goodwin (Pittendrigh and Victor 1957; Goodwin 1964; Goodwin 1965). Then Goldbeter (1995) developed a model with five ordinary differential equations that use *per* mRNA and PER protein (in various phosphorylation states) to describe a negative feedback loop created when PER represses *per* mRNA transcription. Leloup (1998a) expanded this model to include *tim* mRNA and TIM protein. Later models such as those published by Ueda (2001) and Smolen (2001) again expanded the feedback loops by adding CLK, and more recent models added a feedback loop based on *vri*, *pdp1* (Smolen *et al*. 2004; Xie and Kulasiri 2007; Kulasiri and Xie 2008) and CWO (Fathallah-Shaykh *et al*. 2009). Two other common points of interest for clock models were the importance of a positive feedback loop (Tyson *et al*. 1999; Smolen *et al*. 2001; Kuczenski *et al*. 2007; Wang and Zhou 2010) and the influence of post-translational modifications such as phosphorylation (Leise and Moin 2007; Risau-Gusman AND Gleiser 2012). Because the clock is able to adjust to temperature variations, a number of models further investigated the role of temperature on the clock (Kidd *et al*. 2015). Some temperature models, such as those by HONG (Hong and Tyson 1997) and LELOUP (Leloup and Goldbeter 1997), are based on Goldbeter’s 1995 model, while others (Ruoff and Rensing 1996; Ruoff *et al*. 1997; Ruoff *et al*. 1999) use the more general Goodwin (1965) oscillator as a foundation. While models have become sophisticated enough to explain observable biological phenomena, methods have improved for choosing more realistic values for model parameters (Kurata *et al*. 2007; Xie *et al*. 2010; Lebiedz *et al*. 2012). A number of review articles compare the various clock models and present more details on their history (Smolen *et al*. 2000b; Smolen *et al*. 2000a; Goldbeter 2002; Kurosawa *et al*. 2002; Ogawa *et al*. 2008; Gerard *et al*. 2009; Leloup 2009; Gonze 2011; Scribner and Fathallah-shaykh 2011).

#### Role of stochasticity

Researchers have also been using increasingly sophisticated computational approaches for simulating the clock. Many early models were constructed using deterministic ordinary differential equations, but some of their underlying assumptions are not always applicable to the clock. In particular, deterministic models assume large enough numbers of molecules so that any random variations caused by stochastic changes in the state of individual clock components are compensated at the level of the whole clock. Such large numbers of molecules may not be realistic for clocks at the cellular level (Ruoff *et al*. 1999). More recently, stochastic models have been constructed to overcome these limitations; these models are often derived from previously published deterministic equivalents (Barkai and Leibler 2000; Zak *et al*. 2001; Gonze *et al*. 2002a; Gonze *et al*. 2002b; Ueda *et al*. 2002; Vilar *et al*. 2002; Gonze *et al*. 2003; Gonze *et al*. 2004; Miura *et al*. 2008). This allowed for a better understanding of how intracellular stochasticity generates noisy clock observations (Barkai and Leibler 2000; Gonze *et al*. 2002b; Yi *et al*. 2006; Li and Lang 2008; Lerner *et al*. 2015) and how this noise can be reduced by synchronizing clocks across groups of neurons (Katakura and Ohmori 2006; Bagheri *et al*. 2007; Bagheri *et al*. 2008b; Diambra and Malta 2012; Risau-Gusman and Gleiser 2014).

#### Shared problems

The models above examine different aspects of the clock, but they all face two common modeling challenges: estimating parameters and testing the quality of models. Both issues require the ability to access and use high-quality experimental data, yet there is a painful lack of experimentally measured rate parameters for important circadian clock processes. Researchers have used a wide range of methods to find potentially realistic parameters (Leloup and Goldbeter 1998a; Leloup and Goldbeter 2000; Smolen *et al*. 2001; Smolen *et al*. 2002; Smolen *et al*. 2004; Ruoff *et al*. 2005; Xie and Kulasiri 2007; Bagheri *et al*. 2008a; Kulasiri and Xie 2008; Wang and Zhou 2010). These include trial-and-error approaches, but even the best systematic methods cannot guarantee finding rates that reflect nature’s values. In principle, such searches aim to find combinations of input parameter values that cause models to produce time series mirroring those observed experimentally. This ideally provides ensembles of realistic parameter combinations that cannot be ruled out by experimental evidence. It is immaterial, whether such ensembles were generated by testing deterministically or stochastically proposed parameter combinations. However, in no case is it possible to “validate” any parameter combination on principal grounds due to the open nature of the models as discussed elsewhere (Oreskes *et al*. 1994; Tarantola 2006). The best parameter estimates are thus “realistic” (up to a given stringency), and conclusions based on simulations using them are reasonable (up to the usually unknown degree to which these parameter combinations represent reality). This indirect approach has been successful in a wide range of disciplines for estimating parameters in complex models (e.g. (Stainforth *et al*. 2005)). The remaining “unknown” degree can be narrowed for a given model with unknown parameters by using statistically rigorous approaches (Tarantola and Valette 1982; Jaynes and Bretthorst 2003; Tarantola 2005; Tarantola 2006; Moura Neto and Silva Neto 2013). The problem of unknown model parameters is widespread in many disciplines that use modeling approaches and has also become known as the “inverse problem”; it can be can be solved in principle by probability theory (Tarantola and Valette 1982; Jaynes and Bretthorst 2003; Tarantola 2005; Tarantola 2006). Concisely stated, the inverse problem is the challenge to use all known data about a system for restricting the ranges of unknown causal input factors for a model that produces simulation output that is equivalent to data observed in the real system itself (even though the latter ultimately remains unknown). Solving the inverse problem for increasingly realistic biological models using growing datasets of varying quality quickly exceeds current mathematical and computational capabilities and thus remains a research challenge. Additional limits for such reverse-engineering of systems biology models may come from the large variability of their kinetic rates (Erguler and Stumpf 2011).

#### Parameter estimation in complex models

Numerous algorithms can propose sequences of input parameter combinations that repeatedly reduce computed distances between simulated and observed data. However, few frameworks can rigorously estimate the statistical uncertainty associated with their point estimates. Maximum likelihood and Bayesian statistics are currently the frameworks that are most advanced (Edwards 1992; Heyde 1997; Jaynes and Bretthorst 2003; Bishop 2006). They usually require a function that directly computes the likelihood that a given system will produce a given set of observations for a given set of input parameters. However, this likelihood function is increasingly difficult to specify for non-linear stochastic models of growing complexity such as the circadian clock models of interest. Help may come from formalized frameworks for Approximate Bayesian Computation (ABC) that have been developed in various disciplines and do not require explicit likelihood functions (Toni *et al*. 2009a; Csillery *et al*. 2010a; Robert *et al*. 2011; Sunnaker *et al*. 2013; Wilkinson 2013; Lee *et al*. 2014b; Stumpf 2014; Buzbas and Rosenberg 2015). In theory, ABC can solve inverse problems for all simulation models capable of producing output that is comparable to real observations. Briefly, ABC approximates likelihoods by (i) proposing new potentially realistic input parameters, (ii) simulating the model to predict corresponding results, (iii) calculating the distance of these results to experimentally observed data, and (iv) deciding which input parameters are actually supported by experimental evidence, based on comparing these distances to predetermined acceptance criteria. Thus, ABC generates ensembles of model variants, which describe sets of biologically realistic parameter combinations that quantify the uncertainty associated with a given model in the light of available data. Recent progress on uncertainty quantification via ensemble analysis has been reviewed by BAUER *et al*. (Bauer *et al*. 2015). The accuracy of such ensembles depends on the quality of distance measures, acceptance criteria, and sampling density in relevant regions of parameter space. The statistical, numerical and computational challenges associated with ABC increase with model complexity and data diversity. In practice, sampling speed is often limiting, and distances to observed data might have to rely on summary statistics that can create complicated biases. These biases will matter when models with different structures are compared and these summary statistics do not capture all information that is relevant for fully evaluating the models (Robert *et al*. 2011). Fortunately, these problems can be solved for simulations of biochemical systems (Toni *et al*. 2009a; Robert *et al*. 2011). Generally, ABC benefits from access to raw experimental observations such as time series to maximize the information used to estimate parameters and minimize bias from incompletely processed data. Estimating parameters using ABC in fly clock models of realistic complexity is very challenging and has not yet been attempted (to the best of our knowledge; recent work in Neurospora clocks (Deng *et al*. 2016) demonstrates some of the challenges). While the full potential of ABC still waits to be realized, the clock modeling community has estimated parameters either by fitting model output to abstract time series traits or by using time series data more directly. These approaches are discussed next.

#### Using abstract time series traits

The use of higher-level abstractions of time series traits can greatly simplify assessing the realism of a given circadian clock model, at least when compared to data-intensive work with raw time series. Despite the very general nature of abstract traits (e.g. “oscillates” or “has feedback loops”), they can provide powerful filters for removing biologically uninteresting parameter combinations when analyzing circadian clocks. More specific examples of such abstract traits are:

1. period close to 24 hours or modified as observed in mutants (Goldbeter 1995; Leloup and Goldbeter 1998a; Leloup and Goldbeter 1998b; Roenneberg and Merrow 1998; Leloup and Goldbeter 2000; Ueda *et al*. 2001; Fathallah-Shaykh *et al*. 2009; Wang and Zhou 2010; Risau-Gusman and Gleiser 2012),
2. phase changes based on light exposure (Roenneberg and Merrow 1998; Smolen *et al*. 2004),
3. ability to account for responses to light, including light pulses (Roenneberg and Merrow 1998; Tyson *et al*. 1999; Leloup and Goldbeter 2000; Leloup and Goldbeter 2001; Petri and Stengl 2001; Smolen *et al*. 2001; Smolen *et al*. 2002; Smolen *et al*. 2004; Ruoff *et al*. 2005; Bagheri *et al*. 2008a; Fathallah-Shaykh *et al*. 2009)
4. ability to be properly entrained (Smolen *et al*. 2001; Smolen *et al*. 2002),
5. robustness to small parameter changes (Smolen *et al*. 2001; Smolen 2002; Smolen *et al*. 2004),
6. ability to replicate the behavior of mutants (Roenneberg and Merrow 1998; Tyson *et al*. 1999; Smolen *et al*. 2004; Ruoff *et al*. 2005; Bagheri *et al*. 2008a; Fathallah-Shaykh *et al*. 2009; Risau-Gusman and Gleiser 2012),
7. delay between the peaks of a given mRNA and its protein (Leloup and Goldbeter 1998b; Scheper *et al*. 1999a; Scheper *et al*. 1999b; Smolen *et al*. 2001; Smolen *et al*. 2002; Xie and Kulasiri 2007; Wang and Zhou 2010; Risau-Gusman and Gleiser 2012),
8. dynamics of the combined amounts of all forms of PER protein (Petri and Stengl 2001; Smolen *et al*. 2004), and
9. time at peak expression of a given clock component (Petri and Stengl 2001; Fathallah-Shaykh *et al*. 2009).

Comparing simulation results and observed values for such abstract clock traits is generally easier than comparing simulations with complex experimental data. However, it is unclear how much information about circadian clocks is preserved and how much is biased or lost when reducing all relevant circadian clock output to the abstract measures given here.

#### Using complete observed time series

To reduce these uncontrollable biases from using abstract traits, researchers might seek to incorporate all available time series data in more direct tests to compare the distance between observed and simulated time series. The core idea is to increase statistical power by including as much information as possible when estimating parameters and to use repeated observations to obtain better estimates of the underlying distributions. It is therefore desirable to integrate all observed time series of clocks in flies in a single biological information resource. More time series also facilitate recognizing genuine clock signals among the experimental noise inevitably associated with all biological observations and thus help avoid overfitting. In practice, creating models based on experimental time series is complicated by diverse challenges:

i. All challenges of inverse problems with many dimensions discussed above are (or seem) exacerbated because time series usually provide more degrees of freedom than lower-dimensional summaries of their features. Parameter estimation is particularly difficult due to the complex, non-linear relationships between clock components (Tarantola and Valette 1982; Forger *et al*. 2005). While recent progress in developing statistical frameworks like ABC is encouraging, these are not straightforward to use for models with more than a dozen unknown parameters (Toni *et al*. 2009b; Csillery *et al*. 2010b; Soubeyrand *et al*. 2013; Sunnaker *et al*. 2013; Wu *et al*. 2014). Exploring such techniques is beyond the scope of this paper; here we aim to present a real-world, research-grade dataset that provides a non-trivial versatility and complexity test for candidate methods.
ii. It can be difficult to choose optimal measures for comparing time series, regardless of whether applied to simulations or experimental data. There are many “standard measures” for comparing time series in general (e.g. the Euclidian distance, equivalent to the assumption that amounts follow a Normal distribution), and circadian clock time series in particular (e.g. period length). Selecting one or more appropriate measures is not trivial (Glynn *et al*. 2006; Refinetti *et al*. 2007; Ding *et al*. 2008; Batista *et al*. 2011; Jin 2011; Sun *et al*. 2014; Yin *et al*. 2014; Banko and Abonyi 2015; Kotsifakos *et al*. 2016; Mori *et al*. 2016). This is particularly true when comparing time series with differently calibrated, non-linear scales that may be associated with substantial measurement errors, as is often the case for experimental observations. Thus, many diverse quantitative methods can be used to calculate diverse measures of distance between time series. However, this does not solve the substantial qualitative need for arguing which quantitative approaches are appropriate, if any.
iii. It is challenging to compile all relevant time series observations into one place and organize them in a uniformly accessible manner, as pertinent time series were observed using different methods in diverse contexts and span a rich body of literature across many years. Furthermore, data processing is complicated by many obstacles associated with scattered big data, which characterizes many types of biological information. Practical challenges include the degrees to which

a. data is rarely compiled in a uniform, directly usable data format,
b. different datasets require diverse manual corrections for special cases that are individually rare but aggregately comprise a substantial part of the data and are hence not ignorable,
c. time series data is always incomplete and gaps between observed time points are irregular,
d. data is often insufficiently documented such that it becomes impossible to determine essential information about the precise types and attributes that we collect (and which document the precise meaning of the data),
e. data has ascertainment and other biases as well as error rates that are poorly documented and difficult to control (see reports (Clark *et al*. 2005; Lachance and Tishkoff 2013) of biases in initial samples of human genomes),
f. there are other practical problems that are often associated with scattered big data (see, e.g. (Gitelman 2013; Mccallum 2013)), or that require too much data wrangling before a given information resource becomes useful (Goldston 2008).

The aggregated difficulties of navigating all these challenges make it much easier to understand why only a minority of circadian clock modelers chose to estimate parameters directly from such time-series data, and why many others preferred to match abstract time series traits (see Results below).

#### Using both, complete observed time series and abstract traits

It is obvious that both types of observations presented above have advantages and disadvantages.

*Complete experimentally observed time series* increase the information available for parameter inference, but bring the costs of handling more complex, yet incomplete datasets associated with the inevitable problems of real-world measurements. Due to experimental challenges, almost all such time series report relative amounts that are comparable within a specific observed time series. It is rarely possible to obtain reasonably precise calibrations those absolute units that matter most for modeling: the counts of different types of molecules within their respective cellular compartments. Furthermore, experiments may only report aggregated amounts, averaging over cells or other biological units. Such practical details can substantially complicate the computation of the likelihood that a given model will produce a certain observation. Solving these problems does not determine the weights of different points of observation in time series. Ideally, such weights maximize the impact of key information while minimizing noise to avoid overfitting from algorithms that focus on unimportant details, especially if no absolute calibration is available (as usual). Thus, many researchers have historically avoided raw time series and used higher-level abstractions of time series traits to assess the realism of circadian clock models. The relative difficulties of implementation may have contributed to this trend reported in the Results.

*Working with abstract time series traits* provides the ease of using higher-level traits, but comes at a price of its own. Abstract traits usually require fewer dimensions to be managed and could minimize overfitting if they provide a focused view of important clock features. However, abstract clock traits are difficult to choose and can easily omit potentially pivotal information. Higher levels of abstraction can make it easier to find parameter combinations that mimic observed ones ‘reasonably well’. Since the quality of such fits to observations can be judged in many ways, abstract traits might make it too easy to produce a ‘working clock’. Such model can easily omit details that are essential for understanding a *particular* biological circadian clock. In the worst case they degenerate into descriptions of artificial circuits that oscillate, but are unlikely to help us understand the carbon-based circadian clocks studied in biology. Abstract time series traits can reject many parameter combinations as biologically irrelevant and thus pivotally contribute to the construction of useful clock models. Any given abstract trait is not likely to extract all statistical information from the data. Thus, combining many such traits is essential for successful modeling, yet there is no guarantee that any combination will be statistically sufficient such that it can extract all relevant statistical information.

##### Distances

Furthermore, the use of multiple abstract traits raises the question of how to compute distances among and between simulated models and independent wet lab observations. Many summary statistics provide distance measures that can be adequate for some questions, yet cannot extract all information from the data and are therefore not adequate for other questions. The pervasive non-linearity of circadian clock systems complicates combining multiple traits into one reliable overall summary distance statistic. This results in unknown, unpredictable and hence uncontrollable biases when estimating parameters for clock models. To reduce these uncontrollable biases caused by abstract traits with imperfect statistical properties, researchers have started to incorporate more time series data in more direct comparisons of distances between observed and simulated time series (see below).

##### Both sides offer advantages

It might eventually be possible to combine their insights for improving the accuracy and robustness of parameter estimates in circadian clocks. It often appears easier to compare abstract clock traits than the data-rich simulated time series and corresponding experimental observations that necessarily come with many gaps and complex nuances. However, potentially important aspects of circadian clock mechanisms might be impossible to uncover, except by using a more data-rich time series based approach. This extra data often adds many more dimensions, uncertainties and complexities. It can easily overburden modeling studies with irrelevant details and noise that may lead to overfitting. To counter such difficulties, abstract traits may complement full time series data by acting as powerful filters that remove unrealistic models, which might be difficult to identify in other ways. Combining both appropaches could provide a powerful set of tests for detecting realistic oscillation patterns in new circadian clock models. Increasing the statistical power of such tests will make it increasingly difficult for them to be passed by random parameter combinations. To find values that pass all filters and move beyond a given local optimum, researchers can now use a broad array of optimization techniques combined with raw computational power (Bussieck and Meeraus 2004; Bagheri *et al*. 2008a; Leugering 2012; 2016). However, finding parameter combinations that have extremely rare desired properties does not guarantee the correctness of a model (see discussion in (Loewe 2016)). The vastness of parameter spaces requires caution when claiming that useful parameter combinations for circadian clock models describe biological reality (see Oreskes *et al*. (1994)), even if it was difficult to find working parameter combination. Other input might pass the same set of tests and thus have the same claim to be in an ensemble that might be used to represent biological reality. The purpose of FlyClockbase is to improve the availability of data for testing clock models that might be part of such ensembles.

#### Studies integrating time series data

The challenges above present significant barriers to the incorporation of experimental data into models of the *D. melanogaster* circadian clock. Thus it is not surprising that only three of the many modeling studies we surveyed (see Results below) used experimentally observed fly time series to estimate clock parameters in a more direct way. Fathallah-Shaykh (2009) used published microarray data from Kadener (2007) to fit parameters related to cry mRNA oscillation. Kuczenski (2007) used a Monte Carlo random walk method to find a set of parameters most similar to time series of circadian mRNA and proteins from twelve different experimental studies (Hardin *et al*. 1992; Zeng *et al*. 1994; Sehgal *et al*. 1995; Marrus *et al*. 1996; So and Rosbash 1997; Bae *et al*. 1998; Lee *et al*. 1998; Blau 1999; Bae *et al*. 2000; Kim *et al*. 2002; Cyran *et al*. 2003; Glossop *et al*. 2003). Leise (2007) employed a coordinate search method to estimate parameters based on time series from three papers (Lee *et al*. 1998; Bae *et al*. 2000; Shafer *et al*. 2002). Both Kuczenski (2007) and Leise (2007) point to the fit between the experimental and simulation data as evidence of the quality of their models. While further discussion of the many statistical challenges of parameter estimation in real-world datasets is beyond the scope of this study, we note here that such discussion is rather hypothetical without an actual real-world compilation of “all known” time series observations that can test how many of the real-world complications can be handled by any given approach.

One purpose of our study is to provide such an integrated dataset that paves the way for more thorough analyses of statistical approaches to assessing how good a given simulation result might fit to “all known experimental observations” of wildtype and wildtype-like *D. melanogaster* clocks. We created FlyClockbase to lower the barriers that currently limit the use of real-world data for improving simulation models.

### FlyClockbase data model overview

FlyClockbase is a file-based database for collecting and organizing experimentally observed time series of *D. melanogaster*, reporting the core circadian clock components, such as mRNAs and proteins in various states. FlyClockbase is dedicated to circadian clock research in flies and can after sufficient stabilization accept observations of circadian clocks in other organisms. It is publicly available at:

> ***https://github.com/FlyClockbase***
>
> *will become active some time before final publication. For reviewing purposes, see the simultaneously submitted (not-yet-public) zip-archive; for pre-publication access, please request a copy from Laurence Loewe, who will maintain FlyClockbase for the foreseeable future*.

Despite starting with a shared interest in the same model organism, FlyClockbase is completely independent from FlyBase (Dos Santos *et al*. 2015), a portal for genomic and other information about Drosophila as a model organism.

**FIGURE 2.**
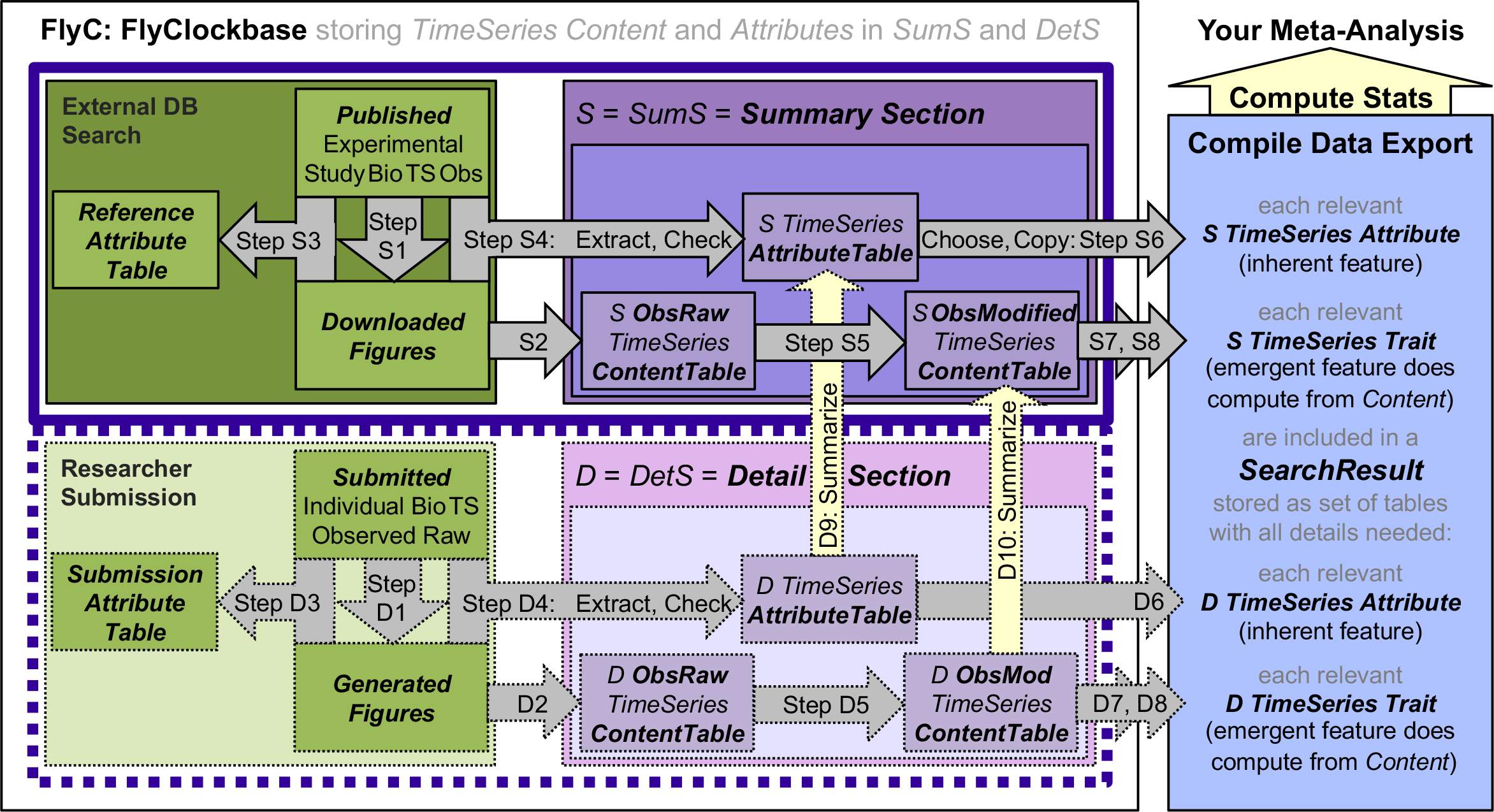
Overview of FlyClockbase organization and data model. We extracted as much raw data and as many reported experimental details as possible from plotted time series summary figures and their publications. We stored bibliographic information in the *References AttributeTable* (Step S3) and stored figures as files ready for plot digitizing (Step S1). For convenient reanalysis, we included all files of open access studies in FlyClockbase. From each figure with appropriate Attributes we extracted the FlyClockbase *TimeSeries Content* (*TS*) itself as *RawObservations* (*ObsRaw*, Step S2). *TS* report some measure of the number of molecules of a given clock component as observed at a given time and in a given volume. The volume and other scaling factors are usually unknown and may be non-linear on occasion. Here we assume that amounts of *Obs TS* indicate a measure of relative abundance among neighboring amounts of the same given *TS*. To place each *TS* in its appropriate context, we collected as many *TS Attributes* as we could reasonably extract from the experimental details given in the original study (Step S4). These *Attributes* are stored in the *SumS TS AttributeTables*. To obtain the more refined *TS ContentTables ObsModified* (*ObsMod*), we rechecked the data based on information in its associated publication (such as exact times of data collection, Step S5). Time series in *SumS ObsRaw* were not refined into *ObsMod* if they were shorter than six hours or if more than half of their values were missing or unreadable. All other *TS* were copied and modified into *ObsMod* and used with their respective *Attributes* for further analyses, ultimately aiming to produce a retrospective meta-analysis. To export the relevant data from FlyClockbase requires a *SearchResult*, which includes the specified *Content*, *Attributes*, and computed *Traits* (stored in *TS TraitTables*). Such *SearchResults* could be compiled manually or automatically and are stored in a *TableSet*, which can include *TS TraitTables*. For the retrospective meta-analysis presented here we manually compiled *TS TraitTables* reporting one peak and one valley from each *ObsMod TS* measured in *CZT* (Steps S6, S7, S8). These peak-valley *TS TraitTables* are further processed in Figure 3. *Content* and *Attributes* for >400 *TS* derived from published studies are stored in the *Summary Section* (*SumS*), which does not include raw experimental data of separately observed *TS*. These will be stored in *DetS*, the *Detail Section*, which is planned as an extension that will enable the submission and processing of more detailed and less summarized *ObsRaw* TS *Content* in order to enable researchers to compute their own independent summary statistics.

#### Overview

Constructing and maintaining a highly specialized biological information resource like FlyClockbase is only feasible for skilled biologists with a passion for flies and clocks. To improve the probability of finding capable biologist curators, we deemed it important to minimize the computational expertise required for making substantial contributions to FlyClockbase. We were aiming to minimize IT overheads of initial construction and longer-term maintenance. As discussed below and in the Supplemental Material, this goal informed important requirements for improving the efficiency of biologists curating FlyClockbase and the accuracy with which its formal type system can capture biologically relevant information. As a result, we have made a number of unconventional database design decisions. Below we summarize key differences between the FlyClockbase design presented here and other typical database designs currently used. We provide more details in the Supplemental Material, but a full technical description of FlyClockbase is beyond the scope of this study, nor can we appropriately present the many considerations that informed the current design. Instead, we focus here on how our choices help biologists, who are (i) interested in the biological question of reliably observable variability in circadian clocks of flies, or are (ii) aiming to navigate FlyClockbase for using, building on, or contributing to the quality of data in this new resource. Figure 2 provides an overview of high-level organization and Table 2 lists the Brief and Explicit Names of various FlyClockbase data structures (we use these *Italicized Proper Names* to distinguish well-specified FlyClockbase data structures from the meaning of their generic English counterparts in usual orthography). FlyClockbase is a Versioned Biological Information Resource (*VBIR*) with two *Sections*:

- *SumS*, the *Summary Section* stores statistical summaries of time series, such as arithmetic averages. We extracted these as they were presented in relevant publications (see Materials and Methods).
- *DetS*, the *Details Section* stores all individual observations available at a stage where they have not been aggregated into summary statistics. This enables independent researchers to compute the best summary statistics for investigating specific questions. Describing *DetS* is beyond the scope of this study (raw data is almost never reported among the publications in FlyClockbase; see below).

Each *Section* stores variants of time series in the form of *Raw* and *Modified Observations* (each in an *ObsRaw* or *ObsMod TimeSeries ContentTable*, respectively). The big workflow steps of importing, fully integrating data, and extracting data for analysis are marked in Figure 2 as steps D1-D10 and S1-S8. To facilitate comparisons across different datasets, we refine *ObsRaw* into *ObsMod* data in our current workflow Step S5 (for more details, see Materials and Methods Section below). Each *TimeSeries* is further characterized by some *Attributes* and may exhibit certain *Traits* (see below for details). *Attributes* denote inherent features that need to be stored and cannot be computed, such an observed genotype. *Traits* capture emergent features that need to be computed from the *Content* and *Attributes* of a time series, such as ‘peaks’. Separating *Attributes* and *Traits* helps to keep FlyClockbase organized and simplifies selecting relevant time series (see Steps S6-S8, D6-D8 in Figure 2).

**TABLE 2.**
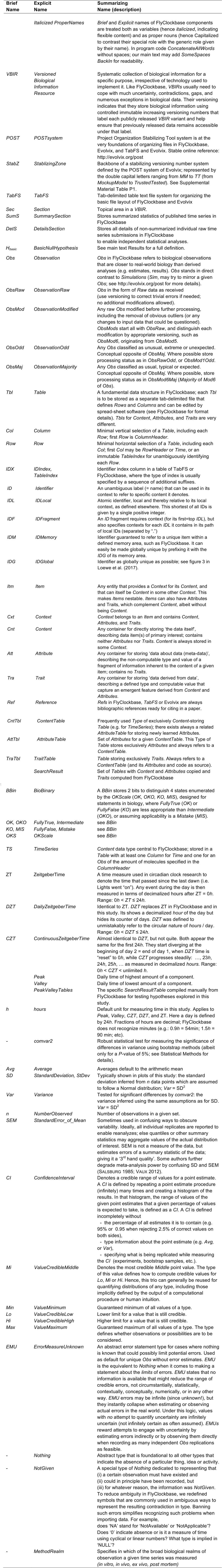
FlyClockbase and *VBIR* concepts and related keywords as introduced by their Brief, Explicit, and Summarizing Name equivalents defined by using the Evolvix BEST Names concept. Overview of concepts by naming gives the *Brief* and *Explicit* Names used in the main text for important FlyClockbase data structures (always in *italics*, usually capitalizing the beginning of significant words). Their *Summarizing* Name gives a description that could be turned into a rather long very descriptive name. The nature of these names is defined by the Evolvix BEST Names concept, see http://evolvix.org/naming and Loewe *et al*. (2017). Order of entries loosely follows conceptual relations. See main text for more details.

#### Logic in biology

Both, *Attributes* and *Traits*, may be absent in ways that may be of biological interest and differ fundamentally between time series. These challenges inspired us to investigate fundamental aspects of type systems and logic in programming languages in a search for appropriate ways of quantifying various aspects of uncertainty, unavailability, and inability to be tested shared among many scattered and diverse datasets of biological interest. In the Supplemental Material we discuss a new data type termed ‘BioBinary’ which stores one of the four alternative states termed OK, OKO, KO, MIS, which are defined by an enumeration termed ‘OKScale’. The BioBinary type is designed for handling statements in biology, where “*completely true”* or “*entirely false”*, are less appropriate than “*any transient intermediate”* or “*mistake”* (see also Discussion below, Supplemental Material, and p.16 of the online supporting material in Loewe et al. (2017); a full analysis is beyond our scope here).

**Other design decisions** of interest to biologists discussed in the Supplemental Material include:

- the intertwining and mutual stimulation of the development of FlyClockbase and the unconventional way in which the Evolvix modeling language (http://evolvix.org), is being developed (Loewe 2016) from a first prototype for recording time series in pure mass action models (Ehlert and Loewe 2014) towards adding general-purpose programming capabilities with the triple goal of maximizing expressivity, usability, and long-term backwards compatibility;
- the use of a stabilizing versioning number system for facilitating review processes in ways that improve the possibilities of working towards long-term stability without frustrating innovators by turning them away, based on the StabilizingZone of the Project Organization Stabilizing Tool (POST) system (see p.74 of online supporting material in (Loewe 2016));
- reasons beyond ease of implementation and installation for not choosing a conventional database system, but rather design-dedicated file-folder structures in a file system that can be copied easily across system boundaries;

While these points are important for questions of reproducibility and programming language development in biology and beyond, they do not directly apply to the biology of circadian clocks and are hence discussed in the Supplemental Material. We next highlight aspects of FlyClockbase that impact the ability of users to represent very diverse data in a surprisingly direct way: our particular choice of basic storage technology.

#### Simple file system storage

To increase flexibility, FlyClockbase stores data in a simple, well-defined, stable layout of files and folders in standard file systems. This design is intended to:

- maximize accessibility to biologists with very diverse levels of computational literacy and who use many different computing platforms. On all of these platforms it should be easy for any researcher to start FlyClockbase: experimental biologists, who strongly prefer to work with standard spreadsheet software as well as computational biologists, who strongly prefer direct programmatic access to raw data files to implement their own analyses,
- minimize long-term maintenance costs by delegating storage to standard file systems that maximize ease of distribution across diverse platforms,
- reduce the need for mandatory database updates that may require costly developer time or endanger the accessibility of valuable data.

These advantages come at the cost of requiring the discipline necessary to maintaining consistency, and expecting users to not irresponsibly alter data that is freely accessible to them. We expect FlyClockbase mostly to be maintained by researchers with sufficient experience and for submissions to be appropriately reviewed so it is always easy for beginners to get the last authoritative version in case they need a fresh start. Our choices of technologies and formats have numerous strategic reasons further detailed in the Supplemental Material.

#### Flexibility

In light of the enormous logic and error-reporting challenges faced by any application (see Supplemental Material), our decision to design FlyClockbase around the simpler and less restrictive technology of a filesystem has provided us with an open field of efficient experimentation to improve our way of handling the challenges of data curation. Our key insight here is the importance to empower experimental biologists with little or no computing background to efficiently *launch* important decisions about the type system and controlled lists used in FlyClockbase. Launching is not landing; developing a stable type-system requires more experience than launching the decision to consider adding a new biological special case. The importance of efficiently communicating and collaborating across very different disciplines cannot be over-emphasized: most people with enough formal experience to understand formal type systems (Pierce 2002; Pierce 2005) cannot imagine the many special biological cases that a corresponding logic would have to be able to handle. This is different for experimental biologists: if they cannot recall these from the top of their head, a few days or weeks in the lab will quickly help them to remember. However, acquiring the necessary biological expertise, usually comes at the cost of less training in the abstract art of designing consistent and stable type systems.

Practically, we developed FlyClockbase’s flexible file system folder-structure to enable the storage of content in standardized spreadsheet files easily modified by common spreadsheet programs. We found this easy to use by experimental biologists who regularly experience (and thus are best positioned to help reduce) the tension between the abstract type system (aiming to restrict chaos by setting some rules) and reality (with its own rules). Their contributions are best recorded on the spot in the most flexible form possible to ensure they are captured at all. This requires maximal flexibility and permissions and is simplest to implement by providing every local user of a local FlyClockbase installation the equivalent of full (FlyClockbase-)system administrator rights (including the ability to add, change, delete, or wreck anything and everything in their local copy of FlyClockbase). Please consult the Supplemental Material for a discussion of permissions, backups and the reliability of data storage.

These and other reasons beyond the scope of this study have motivated us to forgo the obvious speed advantages of well-known standard databases. We do this to gain the potential for a reduction in the cost of maintenance and an increase in stability, combined with the flexibility to experiment with more nuanced type systems. These type systems can better represent the complexity, diversity, uncertainty and occasional contradictions that are so pervasively found in biological data. While working on FlyClockbase, we encountered such problems regularly, as we tried to integrate all available information about the circadian clock of *D. melanogaster*. The Discussion reviews some of these experiences, but a full analysis is beyond the scope of this paper.

#### Data types for organizing content

FlyClockbase is organized around a few data types that help to structure its data (see Figure 2 for overview of components, highlighted as *ItalicizedProperNouns*). These are presented below after briefly discussing the fundamental recurring concepts of *Content*, *Attribute*. and *Trait*, which simplify navigating FlyClockbase data structures. Without loss of generality, we illustrate our definitions using observed time series as an example:

- *Content (Cnt)*: a container for directly storing ‘the data’ describing items of primary interest. For example, the *Content* of a time series observation in FlyClockbase is given by a series of pairs, each storing a time and a number - ordered so times keep increasing.
- *Attribute (Att)*: data about data. Each *Attribute* stores a type of value describing a fragment of information ‘inherent’ to a given data item, such as one of its owners, methods of observation, contexts, or types. The inherent nature of *Attributes* implies that they always have to be stored in addition to, and can never be derived from the *Content* they describe. Sometimes also called ‘metadata’, *Attributes* provide informal descriptions of the type or history of an item that can be essential for the correct interpretation of *Content*. For example, using the *Content* of time series requires *Attributes* describing which type of clock component was observed and in what context – neither of these can be derived from the *Content* itself.
- *Trait (Tra)*: data derived from data. *Traits* capture emergent features, which are externally defined properties, patterns, or conclusions derived from a given set of *Content* and its *Attributes*. For example, a time series that only records how some amount changes over time may allow the observation of one or more peaks, but neither the steps for recognizing such *Traits*, nor the annotations of peak presence or absence are part of the *Content* to which they refer.

Irrespective of how data is packaged, *Attributes* and *Traits* both characterize *Content*, but do so in different ways. In FlyClockbase the trio *Content*, *Attributes* and *Traits* form a causality chain. For example, the real-world circadian clocks of one or more flies **X** *in natura* causally affect the observed time series **Y** *in vivo*, which causally affect results of interest **Z** inferred *in silico*. Note that the flies **X** are described incompletely by *Attributes*, the time series **Y** is observed incompletely as Content and the results of interest **Z** are derived from *Content* and *Attributes* by using a set of steps that define this *Trait*. Researchers often search for some **Z** useful for investigating a **Y**, only to find their efforts undermined by loss of pivotal information on **X** (equivalent to missing type information causing many computer bugs).

FlyClockbase has been built to reduce this very problem for circadian clock research in *D. melanogaster* by providing scientists with all information about X, Y, and Z that has been made available, ideally without increasing, reducing, or biasing any existing uncertainties about X or Y. It is possible to package the same information in a myriad of different ways by nesting and re-packaging various combinations of these three. However, their diverse nature would make the use and maintenance of FlyClockbase unnecessarily complicated, as each type offers a distinct value: *Content* stores each time series that meets the required specification (a big but doable task, aiming for completion). Associated *Attributes* store as much biological and historic context information as possible (often impossibly difficult, making available Attributes very valuable). *Traits* are defined at will by active researchers investigating a given biological question or in search of new interesting *Traits* (it is always possible to define new ones, but few are interesting on the long run). Based on these, we define:

- *ContentTable (CntTbl)*: a table of frequently used data of type ‘content’ *s*uch as a specialized *TimeSeries ContentTable*; *Attributes* and *Traits* are stored separately (see below);
- *AttributeTable (AttTbl)*: a table of *Attributes* for a given *ContentTable*. Currently, the most important *AttributeTables* are those for *References*, and *Summary TimeSeries*;
- *TraitTable (TraTbl)*: a table of *Traits* as determined from the *Trait* definition and a given *ContentTable*. Currently, the most important *TraitTables* are those storing the *Peak* and *Valley* timing for the first day of each *TimeSeries* of each clock component after *ObsMod6* refinement (as given in *SearchResult*);
- *Reference (Ref)*: a specific set of *Attributes*, which combine to storing the bibliographic information about a published study that reports *Summaries* of experimentally observed *TimeSeries* or other data of interest (if not prohibited by copyright, FlyClockbase includes the corresponding files as *Content* of a *Reference*);
- *Reference AttributeTable (Ref AttTbl):* stores each *Reference* (but not the files of its study), determining once and for all its unique *Reference_IDX*, an index used throughout FlyClockbase (the next largest integer available);
- *Submission*: here a submitted set of experimental observations reporting enough *Details* to enable the independent computation of diverse *Summary* statistics of individual observations;
- *SearchResult*: a set of tables compiled automatically or manually from each *TimeSeries* in the *Details* and *Summary Section*, (i) by testing whether all *Attributes* meet the search criteria and (ii) for those that do, by testing whether the *Traits* of appropriately grouped individual or aggregated *TimeSeries* meets the *Trait* search criteria. Our results below derive from a single *SearchResult* extracted from *ObsMod6*, and analyzed in various ways as described.

The basic layout of the folder structure in FlyClockbase follows the layout specified in the Project Organization Stabilizing Tool (POST) system described elsewhere (Loewe 2016). We next discuss additional data types of biological interest before returning to the current scope and data collection strategy of FlyClockbase.

#### Identification of *TimeSeries*

This and future studies will need to refer to time series in FlyClockbase unambiguously. This requires a user-friendly system of precise and stable identification, in order to facilitate giving, using, and maintaining labels for time series with minimal effort. Defining such a system is a challenge facing various naming problems (see tables 1-2 in Loewe 2016). We aimed to avoid two extremes: (i) Using the next running number for the next time series creates efficient labels, but complicates some frequent tasks, such as determining if a pair of time series belong to the same study. (ii) Descriptive labels including author, year, figure-panel, plot-symbol, etc. can be informative, but are often too tedious, hard to automate or difficult to maintain (e.g. avoid synonyms). We therefore developed a system that combines localized integers that stand for local *Items* in different frames of reference, each of which defines a *Context* that is itself an *Item*, nested into a bigger *Context*. The resulting nestable index integers gives the outermost local item identifier (*IDLocal*, *IDL*) as the first, top, left-most integer. This top *ID* is separated by a dot (‘.’) from the next *ID* and provides the *Context* necessary for interpreting this second, next-to-top, next-to-left-most integer. This *ID* in turn is separated by a dot from the third, etc., creating as many nested *Contexts* as needed (more details are beyond the scope of this study).

*In practice*. naming *TimeSeries* unambiguously in FlyClockbase requires the following three types of local identifiers for these three levels of nesting in the *Context* provided by FlyClockbase:

1. *Reference_IDL*, points to a bibliographic reference. The *Context* for interpreting the *IDLocal* of a *Ref* is FlyClockbase itself; *Ref_IDL* is identical to *Reference_IDX* introduced above.
2. *Figure_IDL*. points to a *Figure* in the *Context* of a study, given by its *Ref_IDL*.
3. *TimeSeries_IDL* points to a *TimeSeries* in the *Context* of a figure panel. given by its *Figure_IDL*.

Any contiguous sequence of the elements above forms an *IDFragment* (*IDF*), which identifies its corresponding Items. To distinguish potentially ambiguous *IDFs* from full identifiers in a memory area, we denote the latter as ‘memory identifiers’, or *IDM*s for *IDMemory. IDM*s are unambiguous *IDFs* guaranteed to refer to a unique *Item* within a defined memory area, like unique time series IDs in FlyClockbase. Thus, we can unambiguously identify each *TimeSeries* by its *TimeSeries_IDM* in the FlyClockbase *SummarySection* by using the following form:

> `TS_IDM`
>
> `SumS.Ref_IDL.Fig_IDL.TS_IDL`
>
> `SumS.Reference_IDL.Figure_IDL.TimeSeries_IDL`

where each IDL is replaced by its respective integer. Like all other *ID*s, these *IDL*s are stored as *TimeSeriesAttribute*s in the corresponding *AttributeTable* (*SumS TS AttTbl)*, along with *Attributes* for both identifying the figure in terms used in its publication, and the time series in its figure (e.g. capturing line-type, color, plot-symbol, etc.). Thus, a *TimeSeries_IDM* such as ‘*SumS*.1.2.3’ refers to time series 3 in Figure 2 of reference 1 in the *SummarySection* of FlyClockbase. Since the numbering of time series in a figure panel (etc.), and the numbering of the latter in a study is far from clearly determined, the additional *Attributes* help identify the actual figure panel and time series denoted and would also facilitate the generation of automated reports in the future. For simplicity, we drop the leading “SumS.” from *TimeSeries_IDM*s elsewhere in this text (all TS are *Summarized*). This is appropriate until a *DetailSection* (*DetS*) is introduced for capturing non-summarized time series measurements in FlyClockbase.

*For example*, four figures from one study (*Reference_IDM* “2”) may have the *Figure_IDLs* 1, 2, 5, or 8, resulting in FlyClockbase-wide *Figure_IDMs* 2.1, 2.2, 2.5, or 2.8; currently, gaps in *Figure_IDLs* (such as 3, 6, or 7) are allowed if unavoidable and may indicate that data has been excluded when we later found that it did not fit criteria for inclusion. If the above FlyClockbase-wide *Figure_IDM* 2.5 includes the pertinent local *TimeSeries_IDL*s 3, 7, and 8, then their FlyClockbase-wide *TimeSeries_IDM*s will be 2.5.3, 2.5.7, and 2.5.8. Each such *TimeSeries_IDM* points to a unique experimental observation, unless marked in FlyClockbase as one of the rare cases where a review re-publishes an older time series along with new data. In principle, a new *IDL* can be any integer that has not yet been used in its local context. In practice, FlyClockbase will critically depend on a single naming authority for assigning integers to their corresponding items and ensuring that these assignments are never changed. Initially this naming authority will be the maintainer of FlyClockbase, until this functionality can be automated. Submissions of new studies to FlyClockbase can assign final *IDL*s for figures and time series, but only a temporary *IDL* for a reference. The final *Reference_IDM* can only be assigned once the new entry has arrived in memory area of FlyClockbase. Following proper procedures for these naming issues is essential for the integrity of FlyClockbase. Naming is complicated and the source of much concern for managing biological data in *VBIR*s (NIH *et al*. 2012; Loewe 2016). Naming time series IDs provides a microcosm of the many problems that complicate naming. Still, fully specifying a concrete dataset for further analysis requires more than particular *TimeSeries_IDMs*: it also requires specifying the type of observation, as discussed next.

#### *Raw, Mod*. and *Odd Observations*

In FlyClockbase, each *Observation* (*Obs*), is a *TimeSeries ContentTable* with time values measured in in *DZT* or *CZT* (as defined below), and an associated measure of the amount of mRNA or protein at that time. Each observation also includes measures quantifying imprecision and variability as shown in the published figure (if any). *ObsRaw* (‘raw observations’) specify time as *DZT* and contain the amount values as collected (including negative numbers or imprecision resulting from undetected human error during data collection). Each *ObsMod* (‘modified observation’) is a *TimeSeries ContentTable* measuring time in *CZT* and transforming observations to contain amounts of mRNA or protein that are easier to compare across time series (see steps S5,D5 in Figure 2). Recognizable problems with *ObsRaw* data are appropriately corrected in *ObsMod*. *ObsRaw* time series over less than 6 hours or with more than half of their *Content* marked as ‘unreadable’ were not simplified into *ObsMod*. To identify and correct errors in FlyClockbase, we found it useful to pay close attention to extreme values that might appear unusual or odd. We denote as *ObsOdd*. This does not exclude them from analyses, but motivated us to revisit the whole deduction chain that led to a given *ObsOdd*. Since any data collection will contain human errors, we thought that users of FlyClockbase might find it useful to have estimates of expected human error rates. We constructed repeated rounds of *ObsMod* (see below), which were then used for our final analyses.

#### Data types of time measurements

We follow disciplinary conventions for defining ZT (ZeitgeberTime) as *hours* since the last light period started (dawn). We initially also defined this point in time as exactly ZT = 0. However, this resulted in occasional unintended confusion of two very different meanings, simply because both are conveniently denoted by zero: (i) a valid time measurement indicating an event exactly at dawn as denoted by ‘0’ and (ii) the inappropriate use of ‘0’ for indicating that a time was *NotGiven*. While the absence of a particular expected measurement is to be indicated by the label ‘*NotGiven*’ in FlyClockbase, it proved difficult to guarantee that no unintended ‘0’ could slip in. Elsewhere, such as for elementary addition, the use of ‘0’ for indicating absence as in ‘0 apples’ is justified; it is also common enough and deeply engrained, so that every new curator would have to spend significant learning effort to avoid this ambiguity. Moreover, such errors are difficult to find, because a careful analysis is required to determine, whether a particular ‘0’ indicates ‘0*h*’ or ‘not given’. To improve the long-term quality of FlyClockbase *and* reduce curation costs, we decided to use a more robust Code2Brain interface (Loewe 2016) instead. Hence, *DZT*=0*h* has been declared a risky ambiguity to be removed from FlyClockbase as soon as possible, whenever found (process is ongoing). The old *DZT*=0*h* is replaced by the new *DZT*=24*h*, such that 0*h* < *DZT* ≤ 24*h*, while absence continues to be denoted as *NotGiven*. We think that this new approach has a robust Code2Brain interface (Loewe 2016) and provides a high-quality representation of *Null* for DZT values (see Table 2 and Discussion of Errors in Compilers below and elsewhere (White *et al*. 2013)). An important difference exists between measuring fractions of *hours* in FlyClockbase and outside. Usually, 1 hour comprises 60 minutes, but hours in FlyClockbase are decimalized; thus, fractions of *hours* are measured in decimal fractions and not minutes and the next hour is imminent at 0.99 *h*, not 59 min.

##### CZT

To simplify analyzing several days of data in sequence, we define *Continuous ZT* (*CZT*) to be an extension of ZT such that time increases without interruption at the same rate over multiple days – instead of switching back to ZT = 0 at the start of each new light period (dawn). Some studies use Circadian Time (CT), which can carry the connotation of time series recorded in unusual light schemes such as 24-hour darkness (DD) or 24-hour light (LL). *CZT* helps us to avoid any ambiguity. We use the term *Daily ZT* (or *DZT*) when we mean ZT in this study and in FlyClockbase to reduce the potential for confusion with *CZT* (also reduces ambiguity about types of time in code). Storing the respective day together with *DZT* or ZT creates a 1:1 relation to *CZT*. For example, the times of “lights on” (dawn) and “lights off” (dusk) over three days in the LD 12:12 scheme (12 hours of light (L) followed by 12 hours of darkness (D) each day) used in the experiments of the initial FlyClockbase release can be given as *DZT* (24, 12, 24, 12, 24, 12), where days are implicitly assumed to form a sequence. These times are equivalent to *CZT* (0, 12, 24, 36, 48, 60), using a different way of encoding days implicitly. *CZT* time series simplify selecting observations only from the first day in any given time series, as done in this study. We use italics for *DZT*, *CZT*, *h*, and *hours* to indicate that these types are used as defined in FlyClockbase (see Table 2). We do not italicize ZT, because we do not recommend its use in FlyClockbase.

#### Data types of amounts

None of the time series data collected reflected absolute amounts or concentrations in a cell; rather, they show the amount of mRNA or protein relative to a reference. References are different for many time series, which presents a significant challenge when attempting to compare time series. The relative amount of mRNA or protein at a given time cannot be compared across studies, so we instead turned to a trait-based comparison method. We used modified observation tables to extract two *Traits* from each day of a given time series: time of maximum expression (“peak”) and time of minimum expression (“valley”). We then combined these two *Trait* values with time series *Attributes* to produce *PeakValleyTables*, which represent *SearchResults* for further analysis. For more details, please refer to the Materials and Methods.

#### Current definition of scope

Aligned with our interest to construct the best possible circadian clock model for wildtype *D. melanogaster*, FlyClockbase currently only includes time series from wild-type or wild-type-like flies (e.g., Canton-s, yw, “control”) observed in a LD 12:12 environment from studies published between 1990 and 2015 (see search criteria below). Thus, we currently exclude on purpose any mutants that are meant to carry changes in clock genes, diversity in light-dark regimes and other species for reducing the complexity of data curation. We include as “wildtype-like” any mutants that were constructed without the intention of altering the dynamics of clock components, including reporter genes (e.g. luciferase) and modifications to body and eye color (e.g. yw). We thereby take the reported results at face value, implying that such genetic engineering actually does not affect clock dynamics. Testing this assumption is beyond the scope of this study and might become possible with the help of large numbers of replicates collected in FlyClockbase. Measurement errors associated with many observed time series are substantial and so is their variability between time series. Thus we assume in this study that “wildtype” and “wildtype-like” flies observed in the control experiments of many clock studies can be pooled. As a result, we are including time series from the 86 studies cited below in the first public release of FlyClockbase (QQv1 in the *StablizingZone* notation of the POST system (Loewe 2016), see http://evolvix.org/post). Beyond historic accident, there is no particular reason to limit FlyClockbase to this scope, as long as expansions of scope are coordinated carefully with corresponding data structures that enable the selection of desired datasets.

#### Collection of data

Unfortunately, only one study provided individual raw time series observations in addition to summaries (Shi *et al*. 2014). For the 85 other studies, we extracted observed amounts and times from the time series figures published in these papers (by plot digitizing, see details below). Future releases of the database will allow the inclusion of individual time series observations in the *Details Section*. This will enable meta-analyses to customize the statistics they report in order to choose measures of variation that may be more appropriate than the arithmetic mean and standard deviation. In our experience not all studies specify the variation measures they report with the appropriate care (e.g. failing to specify whether a figure reports standard devitations or standard errors of the mean; see Salsburg (1985) for similar experiences).

## MATERIALS AND METHODS

### Literature search

We searched the literature databases PubMed and Web of Science to collect references with time series data of the core components of the *D. melanogaster* circadian clock. Time series were broadly defined as any timed measurements of amounts of a relevant type of mRNA or protein that showed a daily peak or valley time for clock components, irrespective of the absence of scaling, calibration, and linearity. Search terms focused on variations of the terms “drosophila melanogaster” and “circadian clock” (plural, singular, or MeSH terms, or requiring any of the words in “circadian clock”). Marking phrases as specifically being MeSH terms did not influence the number of results. Using plural search terms (i.e., “circadian clocks,” “clocks”) reduced the number of results, sometimes by hundreds of articles. Requiring both words in “circadian clock” (as opposed to allowing either “circadian” or “clock”) also decreased results by up to half. To reduce the likelihood that relevant data would be excluded in the initial literature search, we chose terms that produced as many results as possible. The final literature search occurred on March 26, 2015. After we removed duplicate studies, this initial search produced 1249 results.

#### Initial eligibility assessment

We assessed the title and abstract of each study identified in the literature search based on the following three factors:

1. *Apparent content*. We excluded articles focusing on organisms other than *D. melanogaster* or centered on processes other than the core clock. We define the “core clock” to be comprised of genes integral to the functioning of the circadian clock in pacemaker cells, with a particular focus on the small ventral-lateral neurons. These genes include those shown in Figure 1 as well as *ck2a*, *sgg*, and *pdf* (see Table 1). Genes related to upstream or downstream clock processes were not included as part of the core clock; neither were genes that affect transcription, translation, or degradation rates in general. We also excluded papers if they were deemed unlikely to contain relevant time series based on the title. The articles we excluded based on this criterion focused on functional areas such as sleep, rest, arousal, locomotor rhythms, the visual system, metabolism and feeding.
2. *Type of data*. We only included papers with experimental data. As simulation data is beyond the scope of FlyClockbase, we excluded articles focusing solely on mathematical models since simulation data is beyond the scope of FlyClockbase.
3. *Format and availability*. We excluded the following reference formats because they were unlikely to contain specific experimental data: book chapters and prefaces, comments, dispatches, features, meeting reports, monitors, news, outlook articles, prediction reports, perspective articles, and reports from workshops. We also excluded one paper that was not available in English.

We then examined the full texts of the remaining 603 studies to determine which papers contained time series data. We were able to find only one article with raw time series data (Shi *et al*. 2014), so we used time series figures as a summarized proxy for raw measurement data. We found 149 studies with at least one time series figure.

#### Biological eligibility assessment

We further filtered these 149 studies with time series based on the following biological factors:

1. *External conditions*. We excluded time series with light schemes other than 12 hours each of light and darkness (12:12 LD) or with temperatures that varied over the time of collection.
2. *Observed cell specimens*. We required time series data to be based on measurements taken from biological material including at least some of the neurons closely related to the central clock in Drosophila (e.g., the small and large ventral-lateral, dorsal, dorsal-lateral, and posterior neurons, and S2 cells). For example, we included time series data taken from whole fly heads, whole flies, or the specified cell groups but excluded data from fly eyes or wings.
3. *Genotypes*. We only collected time series of wild-type or wild-type-like fly strains. We considered strains described as “wild-type”, “control”, “+/+”, or “Canton-s” to be wild-type strains. We also included other fly strains if they were natural variants such as CRY-H and CRY-s (time series IDs 9.1.1 and 9.1.2). We excluded genotypes with mutations intentionally inserted to affect levels of protein expression, phosphorylation, binding, or light response of core clock proteins. We characterized wildtype-like flies as any animals with mutations not believed to interfere with the operation of the clock. Examples include “yw” flies (have yellow bodies and white eyes) and insertions of reporter genes such as luciferase, which are co-expressed with clock genes.
4. *Amount and type of data*. We excluded time series if they were generated by mathematical extrapolation, contained fewer than three data points, or covered less than 12 hours.

The 86 remaining studies were defined as biologically eligible for inclusion in the initial release of FlyClockbase and each study, relevant figure and relevant time series were given a corresponding *Reference_IDM, Figure_IDL*, and *TimeSeries_IDL* as described above (see Figure 2, Steps S1-S4, and Section Identification of *TimeSeries*).

### *TimeSeries* data extraction

Since only a single study provided raw time series data in its online material (Shi *et al*. 2014), we extracted numbers for amounts and times from plotted time series in figures as follows. We first extracted a screenshot of each figure from an appropriately magnified downloaded PDF-formatted copy of the relevant study (using the Mac OSX program “Grab,” a simple application for taking screenshots). We then extracted the data from each individual time series from its corresponding image using the open source program “Plot Digitizer” (version 2.6.3 available at http://plotdigitizer.sourceforge.net/) and recorded the result in a *SummarySection ObsRaw TimeSeries ContentTable* (Step S2 in Figure 2). Plot Digitizer requires users to specify the plotted values and the physical locations of minima and maxima for each of the x-and y-axes in the figure. We did not detect significant curvature in the planes of plots (as might be added by careless digitizing of printed copies) and therefore assume that Plot Digitizer’s linear interpolation provides reasonably accurate numerical x and y coordinates of each point manually selected by the user. All y-axis values were directly recorded using Plot Digitizer, unless a study specifically stated the value of mRNA or protein at a given time. For x-axis, the value from Plot Digitizer was disregarded if the value was noted in the text or clearly marked on the graph.

#### Accuracy estimates of digitized *TimeSeries* data

To assess the human operator component of digitizing accuracy, we measured the variability of plot-digitizing by three authors. Each operator plot-digitized the sample time series (63.1, WTLD, green line with green squares) independently three times to produce a set of values ready for inclusion as if it was true raw data. For each operator, time point digitized, and axis, we calculated the following values: the mean of the absolute value of the relative difference between each pair of the three independently produced values. Averaging over all time points allowed us to calculate the average operator difference percentage for the time axis, *t_aod_*, representing an estimate of the relative error that one might expect for a new value added to a time series in FlyClockbase by plot-digitizing. We observed these intra-operator averages for time (with a maximal value of *t_max_* = 48):

> *t_aod_* = 0.72%, 1.79%, 1.80%, including the first point, and
>
> *t_aod_* = 0.76%, 1.07%, 0.75%, excluding the first point of the time series for operator 1, 2, and 3, respectively.

We also observed the following equivalent measures for the amount values *v_aod_* on the y-axis (with the maximal value of *v_max_* = 2), resulting in averages of

> *v_aod_* = 0.95%, 1.15%, 1.11%, including the first point, and
>
> *v_aod_* = 1.00%, 1.23%, 1.16%, excluding the first point.

Operator 3 plot-digitized all values in FlyClockbase. We averaged all values digitized by all operators and we found:

> *t_aod_* = 1.40% and *v_aod_* = 1.06%, including the first point, and
>
> *t_aod_* = 0.84% and *v_aod_* = 1.12%, excluding the first point.

Averaging coefficients of variation calculated separately for each time point by combining data from all operators gives

> *t_acv_* = 1.72% and *v_acv_* = 2.61%, including the first point, and
>
> *t_acv_* = 0.90% and *v_acv_* = 2.79%, excluding the first point.

The drop in *t_aod_* and *t_acv_* when excluding the first point of the time series stems from the uniformity of absolute errors (‘hit a point on screen’), which results in a proportionally larger impact on small values; this can help estimating precise values near zero.

Overall, these measurements indicate that relative errors introduced by plot-digitizing are small (1% or less) compared to the errors associated with the wet-lab measurements. Thus, we conclude that errors from plot-digitizing can usually be ignored. However, these are not the only potential errors in FlyClockbase. We refined each *ObsRaw TimeSeries ContentTable* (direct from plot-digitizing) into a corresponding *ObsMod TimeSeries ContentTable* (see Step S5 in Figure 2) by correcting human errors associated with data extraction and annotation, as discussed below.

#### Extracting *TimeSeries Attributes*

After extracting *ObsRaw TimeSeries Content* from published figures, we extracted associated *TimeSeries Attributes* and *Reference Attributes* from the corresponding published experimental studies (see Steps S3-S4, Figure 2). *Attributes* relevant to a study as a whole is recorded in the *Reference AttributesTable*, while *Attributes* specific to a given time series is in the *TimeSeries AttributesTable*. We collected *Attributes* for each *TimeSeries* to serve two purposes:

i. to help us to ensure *TimeSeries* fit the biological eligibility criteria previously described;
ii. *Attributes* enable later comparisons of biological, methodological, and other factors that could result in variability between time series.

We collected *Attributes* related to these three categories of questions:

1. Information about the time series

a. Which *MethodRealm* does the time series belong to (*in vitro*, *in vivo*, *ex vivo*, *post mortem*)?
b. What is the molecular type of the time series? Which protein or mRNA does it represent? Which isoform or splicing variant, or phosphorylation state of a given mRNA or protein was measured (if relevant)?
c. Does the time series reflect data based on Zeitgeber Time (*DZT* or *CZT*)?
d. Do the amounts reported in the time series come provide any information for limiting associated measurement errors? If yes, which types of errors are reported: the standard error of the mean (SEM), the standard deviation (SD) or an UnknownErrorMeasure (UKEM)?
2. Information about the method used to collect the time series

a. Which method was used to observe the time series?
b. Which machines, reagents, and software were used?
c. Which probes or antibodies and dilutions were used (if relevant)?
d. Which calibrations were applied, both mathematically (e.g., raw values were scaled by the maximum value) and biologically (e.g., values measured are relative to a specific standard mRNA or protein)?
e. How many repeats were observed, and how are those repeats defined?
f. When was the first and last data point recorded (CZT or DZT with days)?
g. How long did the overall experiment last (in hours)?
h. How long were the intervals between observed data points (if regular)?
i. At which specific times were observations recorded (if specified in the text of the study or clearly marked on the time series figure)?
3. Biological and environmental information about flies used to collect time series data

a. Were flies exposed to light:dark schemes other than 12:12 L:D?
b. To which temperature were the flies exposed?
c. How long were the flies entrained?
d. How old were the flies?
e. What were the genotypes of the flies?
f. Which sex(es) of flies were used?

In some cases, longer *Comments* directly copied from the text of a study were the most appropriate way of describing a given *Attribute* without introducing the potential for errors from paraphrasing. To avoid visual clutter and unnecessary bloating of the *TimeSeries AttributeTable* we introduced the notion of column locality, which is defined by a locality index column that is allowed to have identical values in consecutive rows and thereby define one *LocalColumn* for each such run of identical values. The purpose of this construct is to provide a formal check for the use of the equivalent of ‘*ibid*.’ in FlyClockbase. For example, adding “See Method Comments, 2.5.1” in the column “Method Comments” at time series row 2.5.2 indicates that the longer comment stored in the row above (2.5.1) also applies to the next time series. To avoid loss of context, the FlyClockbase look-up keyword “See” must be followed by an indication of the column and row that are being referenced. To maintain readability for humans and simplify the implementation of code that understands *LocalColumn*s, such pointers to previous cells of a column must not be interrupted by cells with unrelated content.

### *TimeSeries Traits* analysis of *Peaks* and *Valleys* refined by *ObsOdd* checks

We manually compiled a set of *SearchResultTables* that integrated a simplified set of *Attributes* describing the nature of a given time series. In addition, these included *Traits* that computed the respective times where amounts show a peak or a valley on the first observed day of an *ObsMod* time series in FlyClockbase. These *SearchResultTables* were termed ‘*PeakValleyTables*’ and constructed for each given mRNA and protein. Each row in a peak-valley table corresponds to one time series and records several *Attributes* (from the *SummarySection TimeSeries AttributeTable*, see Steps S6 in Figure 2) and two *Traits* (peak and valley, from the *ObsMod TimeSeries ContentTables*, see Steps S7-S8 in Figure 2 and Figure 3). Each row also stores a day index, indicating the day during which the reported peak and valley were observed, counting from the start of the experiment (*CZT* = 0h). For example, a day index of two would indicate that peak and valley of the time series described by the *Attributes* in the row were observed between 24 and 48h *CZT* after initiating the observation of this time series.

**FIGURE 3.**
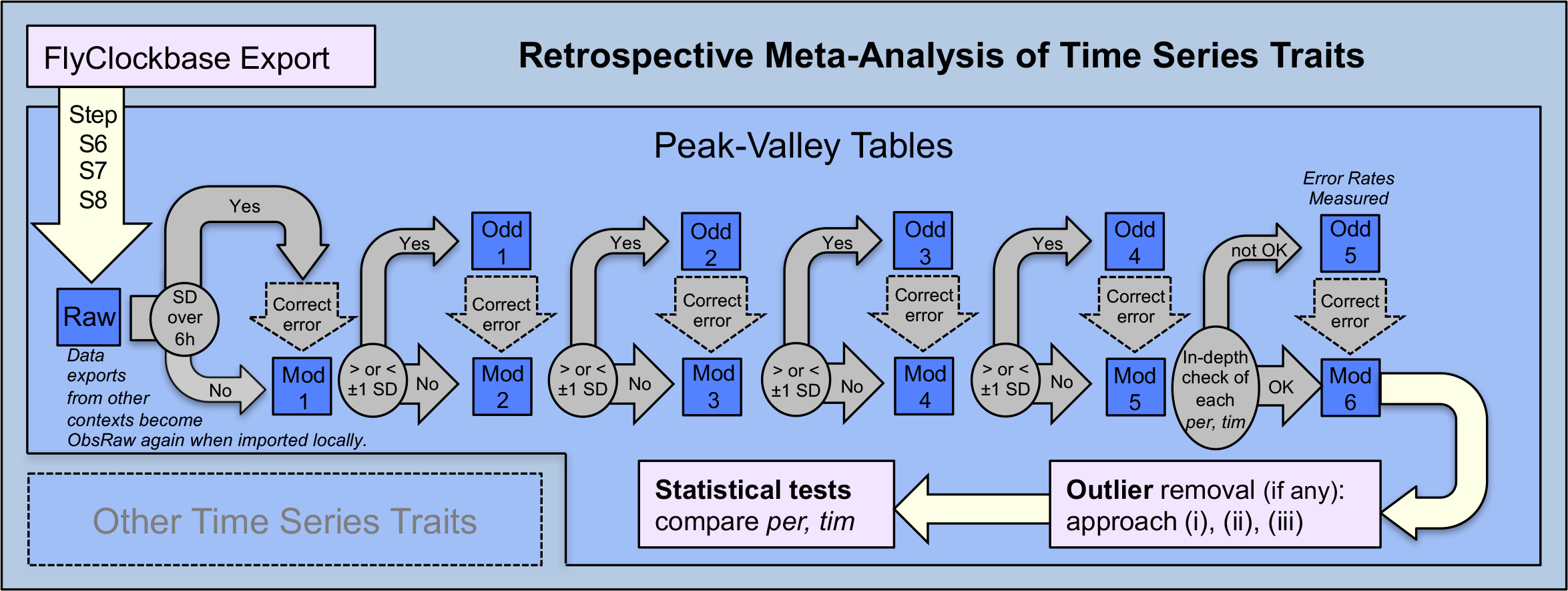
Workflow for refining the time series traits reported in peak-valley-tables. These tables represent a *SearchResult* (as in Figure 2) and contain a special set of *Traits* and *Attributes* extracted from FlyClockbase. In this study we extracted the times associated with the maximum (“*Peak*”) and minimum (“*Valley*”) amounts of the first each day of each time series. Each row of a peak-valley table contains a peak observation and a valley observation for one day of one time series, along with *Attributes* of interest. One peak-valley table was produced for each clock component of interest. *Refinement of outliers (Mod1-Mod5)*. The tables for each component formed a set that was refined together (from *Raw➔Mod1➔*…etc.). *Raw PeakValleyTables* show data for every day of the time series, while all *Modified* (“*Mod*”) *PeakValleyTables* only show data for the first day of the time series. To start identifying outliers and potential errors in these tables, we searched for clock components in which the *SD* for peak, valley, or both exceeded 6 hours. We examined these clock components more closely and corrected errors to create the first collection of modified *PeakValleyTables* (“*Mod1*”). We chose *SD* = 6 as initial cutoff value, since it approximates a uniform random distribution over all 24 hours (to us a sign of problems in correctly identifying the right values. For each modified peak-valley table in *Mod1*, we then calculated the *SD* and mean for peaks and valleys. Individual time series observations outside of the mean ±1 *SD* were noted for further investigation and recorded in a peak-valley table reserved for these “*Odd*” values (“*Odd1*”). We examined *Odd* values for potential errors that we knew could be caused by data handling in FlyClockbase and corrected them appropriately to create the next set of modified *PeakValleyTables* (“*Mod2*”). We repeated this process until we created *Odd5* and *Mod5*. Refinement up to this point was geared towards checking statistical outliers for errors in handling data or determining traits (in contrast to our error analysis below). *Human Error Analysis Context*. When the refinement process was first mapped out, the plan was to stop at *Mod5* and use this repeatedly refined dataset for calculating the final analyses of PER and TIM. Since the curators were diligent motivated researchers aware of potential errors and carefully working to avoid them, there was no expectation that substantial numbers of errors could still lurk beneath the surface. Due to the importance of our conclusions for modeling the differences in stochasticity of PER and TIM we decided to still review all FlyClockbase data before finalizing the statistical analysis of the main result of this paper. We started our review with several random tests, manually conducted on the spot. These detected enough errors to justify a more systematic approach. We decided to avoid shortcuts and focus on an exclusive complete in-depth re-review of all those time series in FlyClockbase where errors in our data handling or trait computation could have affected our conclusions about the peak or valley timing of *per* mRNA, *tim* mRNA, *PER* protein, and *TIM* protein. *Errors tested* included the following notable cases. (i) Peaks and valleys from maxima and minima at the margins of brief time series were probably not genuine, as they would have been unlikely to coincide with the start or stop of time series observations. Excluding these substantially reduced the variance we observed. (ii) The first day was non-representative in some important respect. If peak or valley times during day 1, our usually monitored interval, were noticeably different from subsequent days, we concluded that measurements probably started too early and used the peak and valley from a day that followed closely. (iii) On rare occasions fields in FlyClockbase had been swapped or misprocessed in some way. *Benefits*. While laborious, this effort payed off in multiple ways (see Results and Discussion): (i) It greatly increased confidence in our PER-TIM peak time variance analyses; see Results. (ii) We manually identified particularly extreme outliers not caused by errors on our side. This motivated us to analyze their impact by defining ‘outlier removal approach (ii)’ and choose more systematic outlier removal approaches; see Statistical Methods; see Discussion for potential implications. (iii) FlyClockbase now has an internal human error analysis, see Results. Error rates might be extrapolated to non-*per*/*tim* components or other *VBIRs* of comparable complexity. Currently not many opportunities exist for measuring real-world human error rates in advanced data organization scenarios. (iv) Valuable lessons in automation and *VBIR* compiler construction were learned as a compiler architect worked closely with expert curators during manual error identification. Such manual work is necessarily the first step towards automation of such checks in a corresponding *VBIR* compiler infrastructure as envisioned in the Discussion. *Datasets of interest*. *Mod6* is the most refined set of observations with the highest quality produced by this study and was used as the basis for all biological analyses presented here. It was produced by correcting in *Mod5* all errors in the data handling or trait calculation of *per*, *tim*, PER or TIM, while copying all data for other clock components unchanged from *Mod5*. The dataset *ObsMod7* is essentially identical to *ObsMod6*. except for replacing a few cells in the table that were not analyzed here, but caused problems when R parsed the file. *ObsMod8* is identical to *ObsMod7*, except for manual removal of the manually identified outliers (see outlier removal approach (ii) in Statistical Methods). ObsMod7 and ObsMod8 are provided with the R-source-code archive in the Supplemental Material.

#### Measuring a limit for maximal peak time variance

In an effort to limit mistakes rising from human error, we refined the initially constructed *PeakValleyTables* (‘*Raw*’) into a series of successively modified *PeakValleyTables* shown in Figure 3 (‘*Mod1*’ – ‘*Mod6*’). To create the first set of modified *PeakValleyTables* (*Mod1*), we identified *PeakValleyTables* where the standard deviation of the peak value, valley values, or both was greater than six hours. We based this threshold on two control distributions. First, we created a uniform distribution with 25 artefactual regularly placed observations covering every hour of the day effectively starting at 0*h* up to the very end at 24*h* in 1*h* steps. This distribution had an average and median of *DZT*=12*h* ± 7.36*h SD*; omitting the first or last hour reduced the standard deviation to 7.07*h*. We also randomly sampled 1000 values from a uniform distribution with a range from 0*h* to 24*h*. Repeating this exercise three times produced *DZT* medians of 11.66, 12.41, or 12.08 *h*, averages of 11.85, 12.47, 12.01 *h* and standard deviations of 6.81, 6.92, 7.02 *h*, respectively. We therefore concluded that observations of *SD* ≥ 6h effectively indicate signals that are indistinguishable from randomly distributed impulses that are not oscillating in any discernably coordinated manner. Assuming that clock researchers were probably correct when they reported oscillations, we explored the hypothesis that such high variation *SD*s might have been caused by errors in acquiring or interpreting some aspect of the data.

#### Factors contributing to increased trait variance

An important early insight was the necessity to exclude peaks or valleys that coincide with the first or last point of a time series. Although these points might appear to report the maximum and minimum expression of a clock component, the amounts following or preceding this value (not shown in the figure) could easily continue the local trend. For example, a time series figure could appear to show maximum expression at the final data point (e.g. 23 *CZT*) but reflect a system where the next theoretical data point (e.g. 25 *CZT*) corresponds to the “true” maximum expression. We excluded these time series and made additional corrections of observed human errors to construct the refined dataset *ObsMod1* from *ObsRaw*. This early success in using odd observations for detecting potential lower-level problems in datasets encouraged us to continue to investigate unusually extreme values, which we then defined as *ObsOdd* peak or valley times outside of the range defined by a given clock component’s observed *Avg* ± 1 *SD*. Time series with *ObsOdd* in *Mod1* were recorded in the peak-valley table ‘*Odd1*’. After correcting mistakes in *Odd1*, we combined the corrected values from *Odd1* with the remainder of the data from *Mod1* to create *Mod2*. We repeated this cycle of checking for mistakes, recording unusual values in *Odd PeakValleyTables*, and fixing errors until we created the final set of modified and odd *PeakValleyTables*, *Mod6* and *Odd6*. In addition to correcting more unique human errors, we also made adjustments for these potential method-based sources of errors:

1. *Local minima in peaks*: Some time series (e.g. 85.7.2, 65.1.3, 81.4.1) report a local minimum, where it is easy to intuitively suspect a peak. We only adjusted our peak estimate if (i) a peak was also expected based on other time series of the same clock component, and (ii) the data points on either side of the local minimum are the highest two values for the respective day in the time series. These local dips can result from measurements outside of linear reporting ranges for time series observation methods such as RNase protection assays (RPAs) and Northern Blots. Increases beyond their linear range no longer produce linear increases in the intensity of signals and might even decrease the signal if product inhibition phenomena occur. We therefore suggest these local minima reflect measurement inaccuracies rather than actual decreases in amounts. To correct for this error, we recorded peak time as the average time of the two surrounding near-peak amounts.
2. *Luciferase initial spike*: Time series observed using luciferase (e.g. 62.2.1) may appear to report a peak shortly after starting to record data. The timing of this first peak can be inconsistent with peaks on other days in the same time series and other time series. Often the second peak has a *DZT* timing similar to peaks on subsequent days and in other time series. Such odd initial maxima are likely to be artifacts of the bioluminescence technique used to measure such a time series. They can be caused by an initial adjustment period that is required for accurate measurements of luciferase levels. Reported extra peaks occurs shortly after the arbitrary end of such initial periods (Plautz *et al*. 1997; Stanewsky *et al*. 1997). In such cases, we ignore the initial peak and record the values associated with the second peak on that day (as opposed to the technical maximum).
3. *Minimal Duration*: Some time series (e.g. see Figure_IDs 43.1 - 43.6, 61.1 - 61.4, 75.3) had a duration of twelve or fewer hours. Although minima and maxima can be read from such figures, their use is questionable, in particular, when the actual peak or valley times are not expected in the recorded time. Thus, we mark these peak or valley times as not given.

#### Linearizing TimeSeries data

The cyclic nature of circadian rhythms must be kept in mind when statistically describing the times of peaks or valleys in circadian time series. To illustrate this point, we will use data from the first day of *clk* mRNA time series. With no alterations, mean peak time estimates suggest *DZT* = 8.79h ± 8.59h (*SD*), which is indistinguishable from randomly distributed peaks (see above). Closer inspection of the data, however, shows that these values may be misleading. Calculating the mean and standard deviation depends on finding differences between values representing time, an operation that is substantially complicated by the circular nature of hours in a day, where the value of 24*h* + 3 min results in 0.05*h*. This is in sharp contrast to any linear expectation. For example, the central peak times for the *clk* time series 35.2.1 and 35.2.2 (see Fig. S3B in Kadener *et al*. 2009) are observed at about *CZT* 23 and about *CZT* 27, respectively. Translating them into circular circadian time results in *DZT* 23 and *DZT* 3, respectively. If we then disregard the circular nature of these values, we might infer a time difference of 20h between *DZT* 23 and *DZT* 3. However, visual inspection quickly clarifies that the peak occurring at *CZT* 23 on the first day is close to the early peak of the following day, which occurs at *CZT* 27h given by the sum of (Day 1 *DZT* 24h) + (Day 2 *DZT* 3h). The difference between these peaks in time series 35.2.1 and 35.2.2 is thus more accurately calculated as *CZT* 27 - *CZT* 23 = 4h. The corresponding change is equivalent to a local linearization of time when some part of the original day crosses over into the next or last day. Peak or valley times that have been linearized in this way are hereafter called “linearized.”

We manually linearized all values in the *Mod PeakValleyTables* by moving the minority of values to an earlier or later day (i.e. adding or subtracting 24 hours). Raw *PeakValleyTables* are not linearized. Our current way of linearizing is geared towards analyses of a single 24h period. Close direct visual inspection of all relevant time series and claimed peaks makes it reasonably easy to linearize other periods; here it is beyond our scope to conduct more general analyses. We found that linearization greatly increases the overall reliability of representations of groups of time series. This is of particular importance when estimating the variability of peak and valley times, which is easily inflated artificially by omitting the linearization step.

Table S1 demonstrates the differences that can result from calculating summary statistics for *Raw* (circular time, no value linearized) or *Mod* (linearized time) peak and valley times of selected types of mRNA and protein. Results show that median, mean, and *SD* of linearized times may (but are not required to) differ dramatically from those calculated for circular raw times. For example, the linearized mean peak time for *clk* mRNA occurs 6 hours before its circular raw equivalent. Similarly, the *SD* of linearized peaks is about one-third of the *SD* of peaks measured in raw circular time.

Comparisons to summary statistics of uniform random distributions (see above) are instructive; as a rule of thumb, peak and valley times (or any daily event times) with a standard deviation greater than six hours should be treated with suspicion; they might be difficult to distinguish from randomly distributed times or could be the result of a lack of linearization. The latter affects *Raw SD* values for *clk* mRNA peak time, PER protein peak time, and *per* mRNA valley time (Table S1), which are close to *SD* values for uniform randomly drawn samples. In contrast, after linearization, *SD* for these *Traits* are much more similar to the *SD* of *Traits* of other linearized time series. Overall, this and other experiences suggest that linearization is an important step in obtaining trustworthy summary statistics from circular values such as *DZT*, even though linearization makes no difference in cases where times are already linear (e.g. TIM protein peak with peak *CZT* 18.41 ± 2.54 in Table S1).

### Statistical analyses

Our initial screening for variability of the Traits we call *Peak* time and *Valley* time (observed on day 1) surprised us by suggesting that variability might differ significantly among clock components. Given the various sources of spurious variability described above, we aimed to remove all artifacts that might randomly inflate variability estimates, including potential biases that might be introduced from analyzing more than the first day of a time series (after entrainment). Our interest in reproducible results and disappointing experiences with large untested data collections has inspired various rounds of error checking and increasingly rigorous statistical analysis.

#### Automated analysis with *R* script

The core results of our study (differences in variance between peak times of PER and TIM proteins) have been tested independently by three of us (KS, BH, LL). The most rigorous analysis is presented below and can easily be reproduced by running the script

> `FlyClockbase_PER_TIM_Methods_PeakValley_Comparisons_2016.txt`

which is provided in the Supplemental Material along with input files and the Supplemental Statistical Analysis, which is an annotated PDF, collating all pdf output from our plotting and analysis script. We executed the script on *R* version 3.2.4 (as of 2016-03-10, https://www.r-project.org). It requires the package “data.table” and the library of robust statistical testing functions implemented by this script

> http://dornsife.usc.edu/assets/sites/239/docs/Rallfun-v30.txt

All statistical analyses used data from *PeakValleyTables*, where our final results are taken from *ObsMod6* (see Figure 3 and text above). All corresponding input files are provided next to the R script and are denoted as *ObsMod7* and *ObsMod8* as described in the code. Times are given in *CZT* and have been linearized as described above.

While the script contains numerous comments, it does not attempt to be elegant code. Much of its over 12,000 lines appear at first glance to be repetitive with small variations. It is currently not clear how to simplify the documentation of this script or whether the time required for substantial code improvements would be well invested. The trade-off between readability and coding time is further discussed in the R-code and Supplemental Material under the approach to documenting code denoted as ‘DISCOVARCY’-style.

#### Outlier analysis

We addressed above those irregular values in *ObsOdd* that were demonstrably due to human errors from data processing, removing them from considerations below. Due to the substantial variability of the reported time series and the diversity of measurement methods used to collect them, we were concerned that a few substantially different outliers might obscure a robust trend exhibited by the majority of observations. Thus, we used the following three different approaches for testing the impact of outliers when analyzing Trait *X* by removing as outliers all values *X_i_*, where

i. *X_i_* is outside of the range of non-outliers given by (*q_1_* -1.5 * IQR) < *X_i_* < (*q_3_* +1.5 * IQR), where IQR = (*q_3_* - *q_1_*) is the Inter-Quartile Range, and *q_i_* are the corresponding quartiles,
ii. *X_i_* is identified an ‘extreme value’ by close visual inspection and its extreme difference to equivalent observations. This manual approach removed Protein time series 14.1.1 for TIM, and 43.2.1, 43.3.1, 43.5.1 for PER, but none for the corresponding mRNAs (see the *BestNoXtrem* input for the *R*-script above),
iii. *X_i_* is identified as an outlier by Carling’s modification to the standard boxplot approach (Carling 2000; Wilcox 2012). This technique uses sample size to adjust the range of outliers to account for the tendency to identify a greater number of outliers at smaller sample sizes. We used the implementation described by Wilcox (2012, see section 3.13.3 and 3.13.5 on p.97-98 as implemented in his R script “`Rallfun-v30.txt`” as function “`outbox`” when called with parameters “`mbox=T, gval=NA`”, so that his eq. 3.45 on p.97 is applied). This method is applied to the data analyzed by our R script described above.

#### Testing differences in variance

To test whether differences in variance are statistically significant at the level of alpha = 0.05, we ran 100,000 bootstraps of the percentile bootstrap method implemented by the function “`comvar2`” in the R script “`Rallfun-v30.txt`”, as described by Wilcox (2012) on p.175, section 5.5.2 and elsewhere (Wilcox 2002). This function provides a 0.95 confidence interval for an estimate of the difference between the variances of two groups, but was implemented in a way that only detects significance at the 5% level (without giving *P* values). We used this newer, more robust method to avoid problems associated with older methods such as Levene’s test (Nordstokke and Zumbo 2007).

#### Testing differences in mean

To test for 95% confidence in significantly different locations when comparing distributions, we used (i) the Mann−Whitney−U test as implemented in R (`wilcox.test`, 2 sided, unpaired), (ii) calculated 100,000 bootraps of the “`bootdpci`” difference as described on p.202 in section 5.9.12 of Wilcox (2012), and (iii) 100,000 bootstraps of the “`medpb2`” difference of medians as described on p.174 in section 5.4.3. of Wilcox (2012). Additional details of the function calls are easily found in the source code of our R script that performs these calculations; the results are given in the PDF output of this script, which is also available online as collated and annotated Supplemental Statistical Analysis.

#### Supplemental Statistical Analysis

The same R script produced the results plots shown in the main text below and the 81 pages of auto-generated plots shared with 6 additional pages for navigation as Supplemental Statistical Analysis in the Supplemental Material. It was generated by combining various snippets of code to test for all combinations of input data, outlier removal, observed *Traits*, molecule types and genes. This resulted in 32 distributions, combining the following features: input data (with and without the manually identified *BestNoXtrem* outliers), outlier removal (with and without applying Carling’s outlier removal), and all combinations of *Traits* (peak, valley), molecule type (mRNA, Protein) and gene (*per*, *tim*). Method comparisons employed the same set of statistical tests for comparing locations and variances of two distributions (PCR vs non-PCR).

## RESULTS

### Experimental observations used in modeling

In addition to experimental results, our extensive literature review identified 66 studies published since 1995 which focused on modeling the *D. melanogaster* core circadian clock. Figure 4 shows that about 75% (50/66) of all modeling studies reuse parameters reported in other modeling studies. Of the remaining studies, 13 relied exclusively on abstract time series traits for estimating parameters, two used exclusively direct experimental observations of time series data (Kuczenski *et al*. 2007; Fathallah-Shaykh *et al*. 2009), and one did both (Leise and Moin 2007). In addition, one study re-used parameters from one of the three studies mentioned above. Thus, only 6% (4/66) of all simulation studies were based on direct experimental observations of time series. This does not include the 13 studies that estimated parameters from abstract traits. The substantially different numbers in each category might reflect difficulties inherent in curating and incorporating experimental data into clock simulations (see above Section on Models). We experienced first-hand many such difficulties. Although models will never perfectly simulate reality and ‘validation’ is impossible on principal grounds (Oreskes *et al*. 1994; Beersma 2005), we maintain that direct experimental evidence is critical for increasing the relevance of models aiming to understand reality. Facilitating the construction of more reliable models by including more direct observations motivated us to build FlyClockbase. We found that FlyClockbase also enables interesting retrospective meta-analyses, some of which we report below.

**FIGURE 4.**
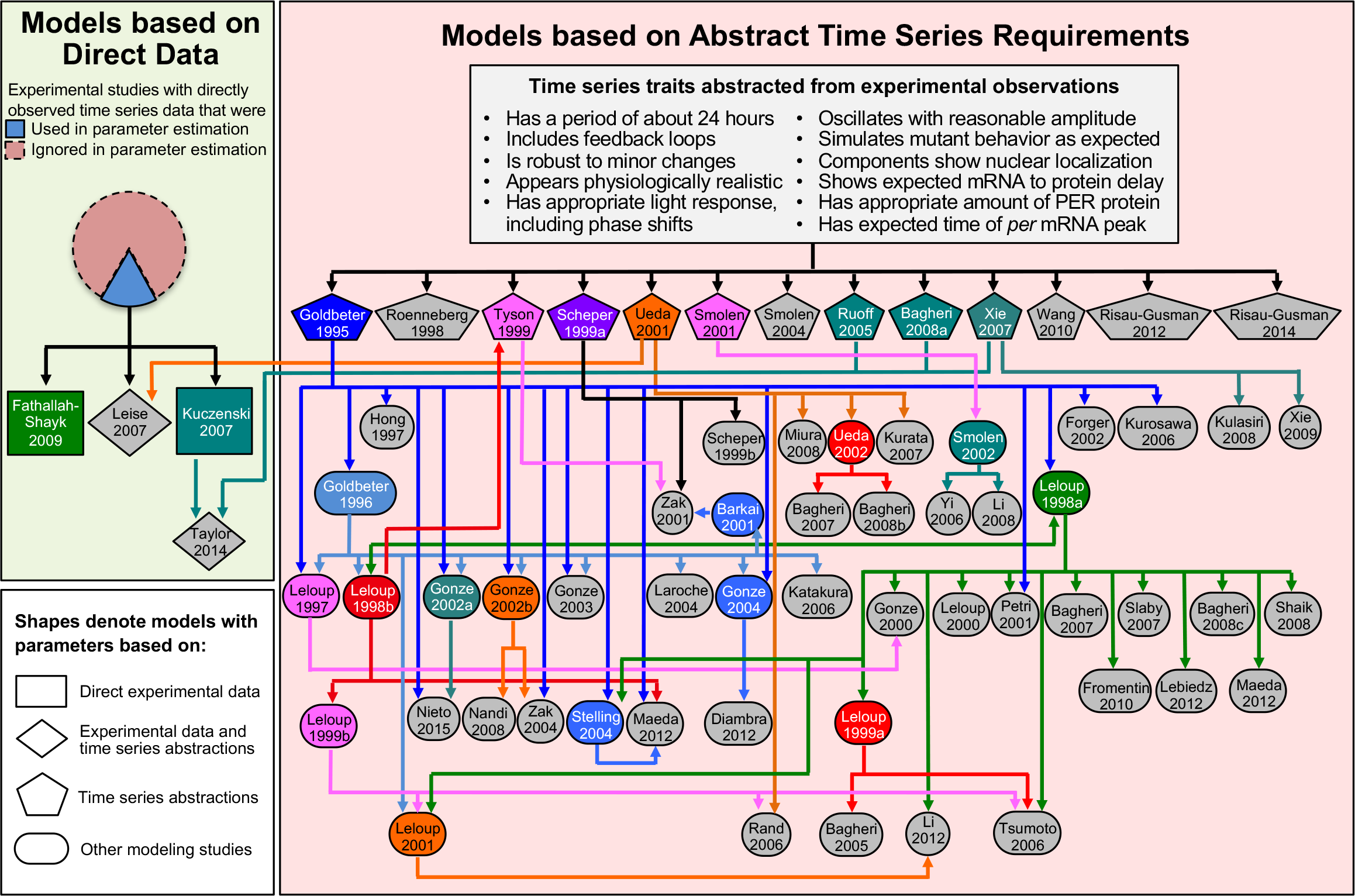
Where do models get their realism from? Overview of data sources reported. Surprisingly few experimentally observed time series are used for estimating parameters in *D. melanogaster* circadian clock simulation studies. We reviewed 66 modeling studies identified here by AuthorYear. Only three of these did estimate parameters directly from full time series observed in experiments (using only 14 of the 86 experimental studies included in FlyClockbase). These three studies (and one that used their results) are denoted on the left by rectangles and rhombuses. The pie chart above them illustrates how many experimental studies were used (blue, solid line, 14) or ignored (pink, dashed line, 72) by modelers. The 13 studies represented by pentagons chose the simpler (but not simple) approach of searching for clock parameter combinations that satisfied some of the abstract time series requirements such as period length, see gray inset box. The 49 ovals represent models with parameters mostly or entirely based on the work of other models. Colored shapes represent models that informed the parameters of other models, where arrows of the same color connect the original and the subsequent models it informs (gray indicates that a model was not yet used for parameterizing other models).

**FIGURE 5.**
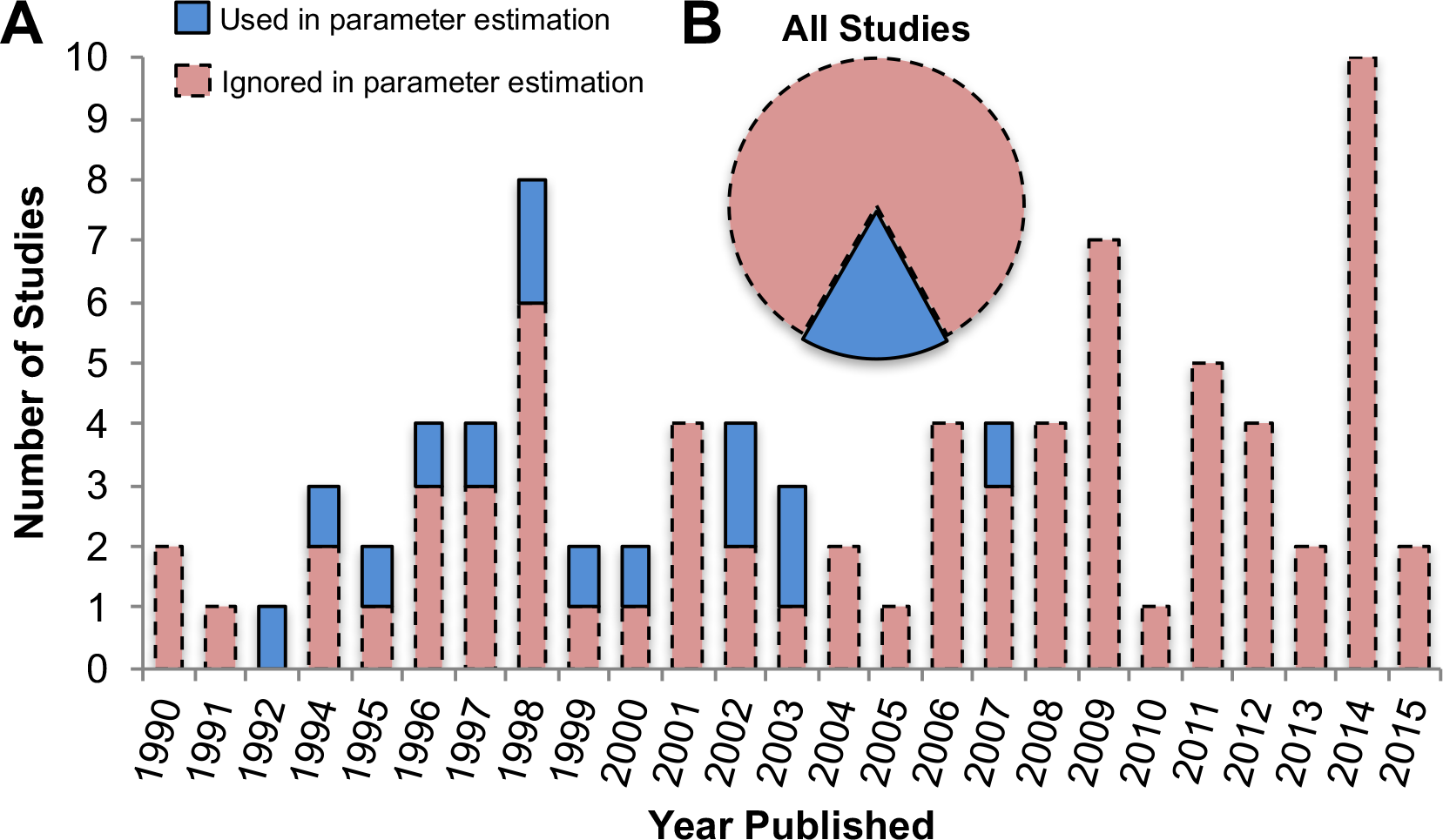
Data accumulation over time: overview of experimental studies available in FlyClockbase and their use for parameter estimation over time. **(A)** All studies in FlyClockbase are displayed based on the year they were published. Most studies have not yet been used to inform model parameters (pink dashed line bars). **(B)** Summary of studies that were later used for parameter estimation are shown as blue solid line bars. These 14 studies are given (Hardin *et al*. 1992; Zeng *et al*. 1994; Sehgal *et al*. 1995; Marrus *et al*. 1996; So and Rosbash 1997; Bae *et al*. 1998; Lee *et al*. 1998; Blau 1999; Bae *et al*. 2000; Kim *et al*. 2002; Shafer *et al*. 2002; Cyran *et al*. 2003; Glossop *et al*. 2003; Kadener *et al*. 2007) and come from the richer collection of 86 studies reported in version variant QQv1 of FlyClockbase: (Hardin *et al*. 1990; Zerr *et al*. 1990; Zwiebel *et al*. 1991b; Hardin *et al*. 1992; Hardin 1994; Sehgal *et al*. 1994; Zeng *et al*. 1994; Sehgal *et al*. 1995; van Gelder *et al*. 1995; Brandes *et al*. 1996; Marrus *et al*. 1996; Qiu and Hardin 1996; van Gelder and Krasnow 1996; Majercak *et al*. 1997; Rouyer *et al*. 1997; So and Rosbash 1997; Stanewsky *et al*. 1997; Bae *et al*. 1998; Cheng *et al*. 1998; Darlington *et al*. 1998; Emery *et al*. 1998; Hamblen *et al*. 1998; Kloss *et al*. 1998; Lee *et al*. 1998; Stanewsky *et al*. 1998; Blau 1999; Ishikawa *et al*. 1999; Bae *et al*. 2000; Rothenfluh *et al*. 2000b; Bao *et al*. 2001; Claridge-Chang *et al*. 2001; Mcdonald *et al*. 2001; Okada *et al*. 2001; Kim *et al*. 2002; Shafer *et al*. 2002; Stanewsky *et al*. 2002; Ueda *et al*. 2002; Akten *et al*. 2003; Cyran *et al*. 2003; Glossop *et al*. 2003; Dissel *et al*. 2004; Majercak *et al*. 2004; Glaser and Stanewsky 2005; Chen *et al*. 2006; Koh *et al*. 2006; Wijnen *et al*. 2006; Yu *et al*. 2006; Fang *et al*. 2007; Kadener *et al*. 2007; Matsumoto *et al*. 2007; Muskus *et al*. 2007; KivimÄe *et al*. 2008; Meissner *et al*. 2008; Peschel 2008; Yoshii *et al*. 2008; Dubruille *et al*. 2009; Kadener *et al*. 2009; Lear *et al*. 2009; Peschel *et al*. 2009; Wulbeck *et al*. 2009; Yoshii *et al*. 2009; Zheng *et al*. 2009; Kula-Eversole *et al*. 2010; Chen *et al*. 2011; Chiu *et al*. 2011; Itoh *et al*. 2011; Lamaze *et al*. 2011; Lim *et al*. 2011; Grima *et al*. 2012; Hughes *et al*. 2012; Kumar *et al*. 2012; Li and Rosbash 2013; Rodriguez *et al*. 2013; Hermann-Luibl *et al*. 2014; Lamba *et al*. 2014; Lee *et al*. 2014a; Oh *et al*. 2014; Pegoraro *et al*. 2014; Shi *et al*. 2014; Simoni *et al*. 2014; Subramanian *et al*. 2014; Weiss *et al*. 2014; Zheng *et al*. 2014; Abruzzi *et al*. 2015; Ma *et al*. 2015).

### FlyClockbase is a new resource enabling studies of circadian clock time series

FlyClockbase includes 403 time series covering 20 different mRNAs and proteins (13 and 7, respectively). An overview of its data model is given in Figure 2, relevant abbreviations are summarized in Table 2. Table 3 shows its number of time series and published studies corresponding to each type of mRNA or protein in FlyClockbase. Figure 5 summarizes publication years from 1990 to 2015 for its 86 experimental studies observing time series of clock components. Figure 5 highlights publication years of the 16% (14/86) experimental wildtype observations used to directly inform 5% (3/66) of all computational model studies of *D. melanogaster* circadian clocks (see Figure 4).

**TABLE 3.** Data in FlyClockbase counting independently observed *D. melanogaster* time series summaries available for each clock component in the Summary Section of FlyClockbase.

| Clock Component | Time Series<br>[*n*] | Studies [*n*] |
| --- | --- | --- |
| *per* mRNA | 89 | 42 |
| PER protein | 77 | 26 |
| *tim* mRNA | 51 | 27 |
| TIM protein | 42 | 16 |
| *clk* mRNA | 33 | 18 |
| *vri* mRNA | 31 | 14 |
| *cry* mRNA | 18 | 10 |
| CRY protein | 16 | 6 |
| *pdp1* mRNA | 15 | 10 |
| CLK protein | 10 | 5 |
| *cwo* mRNA | 5 | 3 |
| PDF protein | 4 | 2 |
| VRI protein | 2 | 2 |
| *jet* mRNA | 2 | 1 |
| *dbt* mRNA | 2 | 2 |
| CYC protein | 2 | 1 |
| *cyc* mRNA | 1 | 1 |
| *sgg* mRNA | 1 | 1 |
| *ck2* mRNA | 1 | 1 |
| PDP1 protein | 1 | 1 |
| **Total** | **403** | **86** |

#### Hypothesis testing

FlyClockbase enables the study of many diverse biological questions, making it impossible to present all corresponding biological results here (see Discussion of using FlyClockbase for hypothesis testing). The analysis of most biological questions, however, requires the computation of relevant *Traits* from the time series stored in FlyClockbase. These *Traits* then need to be combined with the relevant *Attributes* of the corresponding time series to form a row in the table of search results (see Figure 2). We named these *PeakValleyTables*, since we constructed one for each clock component in order to focus exclusively on analyses of the circadian timing of the first peak and the first valley of each time series (defined by the respective maximal and minimal amounts during the first day). After initially establishing FlyClockbase and estimating the variability of all clock components, we limited the scope of this study to the following two biological questions:

i. Comparing the variability of daily peaks and valleys of *per* and *tim* mRNA and protein, are there statistically significant differences in variance across independent time series?
ii. Which differences (if any) occur in the timing of *per* mRNA peak times based on different methods of observation?

We picked these specific questions because the initial release of FlyClockbase contained the largest numbers of time series for these four components (ordered by count we used 89 time series for *per*, 77 PER, 51 *tim*. 42 TIM). Thus, FlyClockbase enables comparisons of variability of independently collected time series of circadian clocks.

#### Error analysis

We conducted our initial retrospective meta-analysis of the questions above after compiling FlyClockbase and refining it to correct all errors in data handling that were known at that time (which produced *ObsMod5*, see Figure 3). Since these initial results surprised us, we wanted to test their reliability. We had no estimates for the frequency of errors that might be expected in FlyClockbase. It was not clear whether there was a faster way of obtaining error estimates for such a complex, manually compiled resource designed to organize scattered data that is heterogeneous, diffuse, and characterized by many complexities, uncertainties, and gaps. We thus opted for the most reliable approach available and rechecked every processing step that could have affected the accuracy of our data sources or every peak and valley timing used in our final analysis (producing *ObsMod6*, see Figure 3). These tests improved our accuracy and generated realistic error rates for users of comparable resources (see below).

**TABLE 4.**
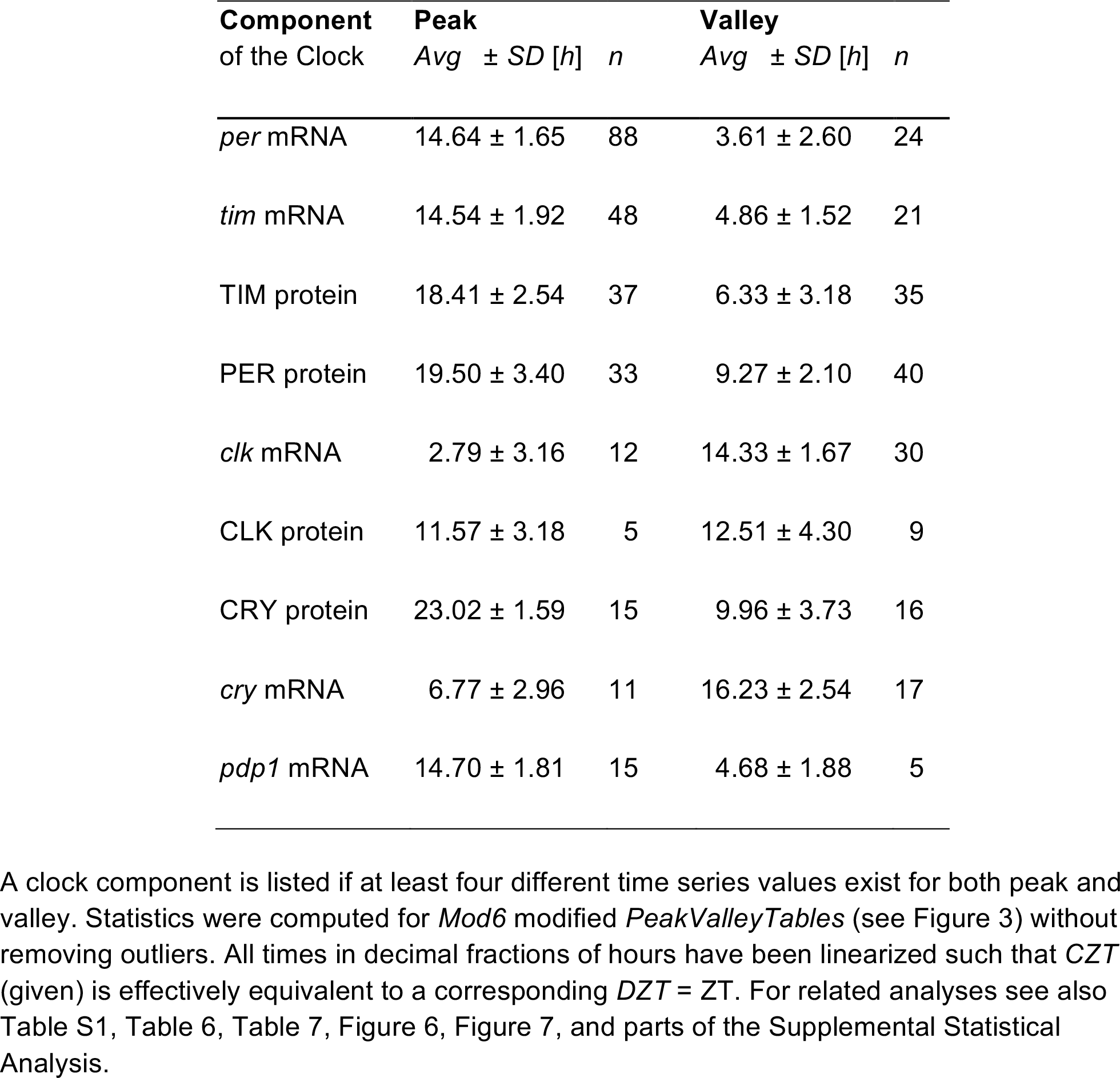
Time series overview: average (*Avg*), standard deviation (*SD)*, and number of time series available (*n*) for daily peak and valley times of some mRNA and proteins in FlyClockbase.

#### Overview of *TimeSeries Traits*

The *PeakValleyTables* described above allowed us to focus on efficient analyses of the timing of extreme amounts. These prominent traits of time series have been used previously to estimate parameters and to compare simulations and experiments (Petri and Stengl 2001; Fathallah-Shaykh *et al*. 2009). Table 4 provides summary statistics based on *ObsMod6 PeakValleyTables* (see Figure 3) after applying outlier-removal method (i) described above. We will later provide an independently computed overview of the mean and *SD* for the peak and valley time of PER protein, TIM protein, *per* mRNA, and *tim* mRNA using outlier-removal method (iii), see Figure 6 below. The amounts of *clk* and *cry* mRNA peak during the day and reach a valley during the night. The components *per* mRNA, *tim* mRNA, *pdp1* mRNA, PER protein, TIM, protein, and CRY protein display the opposite pattern: peaks occur during the night and valleys during the day.

Variability of peak and valley times as measured by standard deviation varied substantially from about one to four hours. This difference motivated us to determine whether these differences in variability were statistically significant and might point to mechanistic causes with implications for the inner workings of the clock. The most frequently observed clock components were *per* and *tim* mRNA and protein, which made them prime candidates for comparing their variability. However, one important task remained before investigating our hypotheses: testing for errors.

#### Why estimate errors?

We had to be reasonably certain that we could rely on the values stored in our *PeakValleyTables*. This was particularly important for detecting significant differences in variability, since random errors in the capture or processing of data are known to easily generate spurious values that substantially impact observable variability. As explored in the Discussion, attempts to detect and correct errors in author-labs and distinguishing them from other potential sources of variability was beyond the scope of our study. Thus, our goal was to be able to confidently exclude errors during the capture and processing of data on our side. We defined as error in this context any result that did not hold up to scrutiny, when rechecked using the rules we had agreed on for processing time series data. These rules were set up after we gained substantial experience with various counter-intuitive aspects of the data, as documented in this study. Thus, our definitions of error were not arbitrary and therefore allow us to contribute below a specific estimate of human error rates to the broader area of human error analysis (see Discussion below).

Databases curated by human experts usually have substantially lower error rates than those that were automatically compiled (Schnoes *et al*. 2009; Koskinen *et al*. 2015). Still, the dangers of error accumulation in heterogeneous collections of data contributed by humans are well known (Carthey *et al*. 2003; Zeeberg *et al*. 2004; Panko and Aurigemma 2010; Panko 2016). It thus appears desirable to increase the number of human error estimates available for biological information resources. The analysis discussed next provided a unique point estimate of the error rates users might want to expect when working with manually compiled resources of similar complexity. A more thorough analysis of errors is beyond the scope of this study (and is likely to be substantially affected by the compiler techniques discussed below). Nevertheless, our results below support our claim that our main findings are probably not the result of a unlucky confluence of statistical flukes generated by errors we could have detected when reanalyzing the same data carefully.

#### Quality control in FlyClockbase and *TraitTables*

In order to detect and reduce the number of errors in the *PeakValleyTables* used for computing variability, we conducted four rounds of identifying values outside of the range defined by the mean ± 1 *SD* of a *Trait* (*Mod2*, *Mod3*, *Mod4*, *Mod5* in Figure 3); this was done for all clock components in FlyClockbase and serves as the basic way of introducing all new data. We used this as the starting point for obtaining the error estimates reported here. Since time series *TraitTables* and search results tables are not strictly part of FlyClockbase (see Figure 2), we separated our error estimates of FlyClockbase from those obtained for *TraitTables* (see Results below) to improve the quality of our estimates. In a unique effort, we then re-examined in-depth all relevant content, attributes and traits of each time series that informed the four most important *PeakValleyTables* (*per* mRNA, *tim* mRNA, PER protein, and TIM protein). We scrutinized each value representing a peak or valley time. This required a substantial effort, which aimed at a twofold goal:

i. removal of errors for improving our estimates of variability differences, and
ii. providing an approximate estimate for the numbers of similar errors that users of FlyClockbase (or similarly complex data resources) might expect in other areas, where such error rates have not yet been determined.

Before presenting our findings, we submit to the reader that the researchers compiling FlyClockbase had been very conscientious, brought a high degree of expertise and enthusiasm to the project, and did their very best to avoid mistakes. Thus we thought conditions were favorable and we were not sure if we would find errors. Unfortunately, aiming to avoid and correct mistakes does not protect against the inevitable occurrence of mistakes at low but predictable rates (see Table 5 and Discussion).

**TABLE 5.** Human error rates for four *Mod5 PeakValleyTables*.

| **Clock Component**<br>TimeSeries counted | **Errors**<br>counted in FlyClockbase | ***Peak TraitTable*** |  | ***Valley TraitTable*** |  |
| --- | --- | --- | --- | --- | --- |
|  |  | **All Peaks**<br>counted | **Errors**<br>counted | **All Valleys**<br>counted | **Errors**<br>counted |
| *per* mRNA (88) | 2 | 87 | 7 | 24 | 6 |
| *tim* mRNA (49) | 1 | 48 | 2 | 20 | 0 |
| TIM protein (43) | 1 | 29 | 4 | 34 | 6 |
| PER protein (57) | 1 | 30 | 12 | 39 | 8 |

#### Basic Null-Hypothesis H_*basic*_

FlyClockbase includes a particularly large number of repeatedly observed time series of *per* and *tim* mRNAs and proteins in wildtype (and wildtype-like) circadian clocks of *D. melanogaster*. These time series were recorded as wildtype control experiments while observing the effects of mutants in order to explore circadian clock functions. We expect corresponding wildtype controls to produce similar time courses that differ only by inevitable stochastic effects – in the absence of experimental complications. Such complications would make flies non-comparable across studies (see Discussion of potential causes, such as variable natural genetic diversity across fly strains, developmental diversity, unknown environmental impact on measurements, and others). We start by ignoring all such potential complications. Our aim is to initially work with the simplest model that still appears somewhat useful. We define a corresponding basic null-hypothesis *H_basic_* to inform our background time series expectations and enable a defined starting point for hypothesis testing with the help of appropriately selected data in FlyClockbase. In light of the types of observations and available calibrations, we define *H_basic_* as:

> *The basic time series type in FlyClockbase is determined by the molecule type observed and its context, which for H_basic_ is defined as the central core clock of wildtype organisms of a defined taxonomic unit (like D. melanogaster) under standard experimental conditions. H_basic_ excludes time series that are (i) from non-LD 12:12 observations, (ii) affected by the presence of mutations known to or intended to alter the clock, (iii) measuring non-central circadian clocks that are independent or merely derived from an organisms’ central clock, and (iv) any complications from genotypes, phenotypes, environments, methods, or more*.
>
> ***For these basic time series types defined above, FlyClockbase reports as H_basic_ the ensembles below containing either dynamically changing amounts of a specified molecule type, or values of corresponding time series traits.***
>
> *These ensembles of the most reliable observations in FlyClockbase are based on time series that satisfy the following conditions: (i) Timed amounts are reported as uncalibrated, relative measurements, which allow comparisons only within each time series, not between time series. (ii) Comparisons of observed amounts indicate all potential outcomes by drawing on as much evidence as possible and extracting as much quality as reasonable from the available data. This requires that comparisons define a method of incorporating such data appropriately. This data includes quantifications of uncertainty in methods of measurement, observational errors, replicates, accuracies of timed amounts, and methods of inference. (iii) When using H_basic_, all relevant details need to be documented, including the data available at the time, its state of refinement (ObsMod+), and the methods used. Documentation requires specifying their stabilizing versioning numbers as defined by the StabilizingZone of the POST system (see Table P1 in Supplemental Material). (iv) Any observations included in either ensemble type exclude outliers using Carling’s method (2000) to enable a focus on typical clocks*.
>
> ***Satisfying all these conditions, H_basic_ assumes that the remaining variability of observed amounts or traits is only caused by the natural stochasticity of discrete molecular events inside of individual cells.***

*H_basic_* is a powerful starting point for exploring clock biology. Next, we will use *H_basic_* to compare typical *H_basic_* behavior of time series traits observed in different clock components. We then relax assumptions of *H_basic_* to illustrate how FlyClockbase can generate a variety of hypotheses about diverse subtleties that might be important for generating high-quality observations of circadian clocks.

### Hypothesis on Peak time variances: PER exceeds TIM

We initially screened for variability in the peak and valley times of all clock components available (see Table 4). This analysis revealed substantial differences between the standard deviations of peak times observed for PER and TIM respectively highlighted in Figure 6. We hypothesized that these differences could have mechanistic causes and might thus point to new insights into clock mechanisms (see Discussion). Before exploring such mechanistic models, we wanted to know, whether our observations were (i) statistically significant and (ii) not caused by low-level errors in data handling during the construction of FlyClockbase in light of corresponding challenges with data quality (Mccallum 2013) as faced by many biological information resources (see Discussion and Supplemental Material for more details).

**FIGURE 6.**
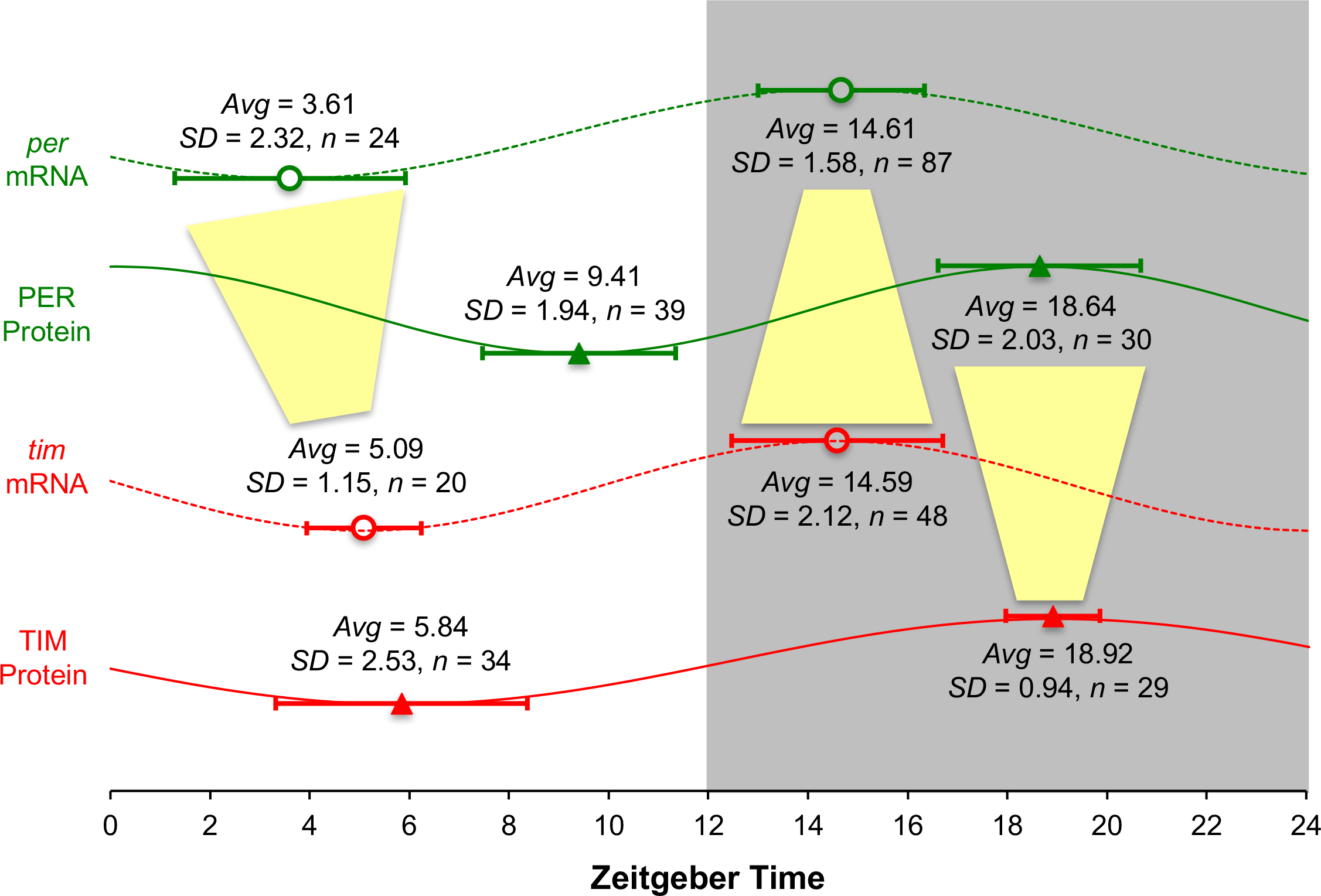
Hypotheses on timing variances in PER and TIM as tested by FlyClockbase data, which extends our understanding of circadian clocks in flies by enabling observations of significant differences in variance between peaks and valleys of different clock components. Here we summarize results from comparing mRNAs and proteins of *per* and *tim*. Yellow funnels highlight changes in respective variances between timing traits of *per* and *tim* that are not negligible (over 5% of 10^5^ bootstraps of available data reject the correspondingly customized null-hypothesis *H_basic_* that variances are equal). For related analyses see also Table 6, Table 7, Figure 7, Table 4, Table S1 and parts of the Supplemental Statistical Analysis. Overall, the summary statistics for the timing of peaks and valleys of the mRNAs and proteins of *per* and *tim* are consistent with our current understanding of the *D. melanogaster* circadian clock. Open circles with dashed curves indicate mRNAs, and filled triangles with solid curves represent proteins; *per* (green) is shown in the top two, and *tim* (red) in the bottom two curves, all adjusted to fit locations of peaks and valleys within one cycle. Averages and *SD*s were calculated for *CZT* times, linearized as described in Materials and Methods, and then back-transformed to *DZT* (if differences existed as in the rare cases where day 1 of a time series was unusable). The statistics shown report summaries for the *n* remaining time series after removing outliers by applying the method of Carling (2000) to the data in the *TraitTables* compiled from the time series observations in FlyClockbase and refined as described for ‘*Mod6*’ in Figure 3. Histograms of the data summarized here are shown in Figure 7 as ‘CoreData’ and statistical results testing the significance of differences are reported in Table 6, the text of the Results Section, and in the Supplemental Statistical Analysis (automatically produced by the R script provided as Supplemental Material). Table 4 and Table S1 present related results from an independent statistical analysis of the same data that was completely implemented using only spread-sheet software. While the results shown in this figure and the underlying outlier analysis were performed in R, the input dataset *Mod6* analyzed by the R script was produced using spread-sheets.

**FIGURE 7.**
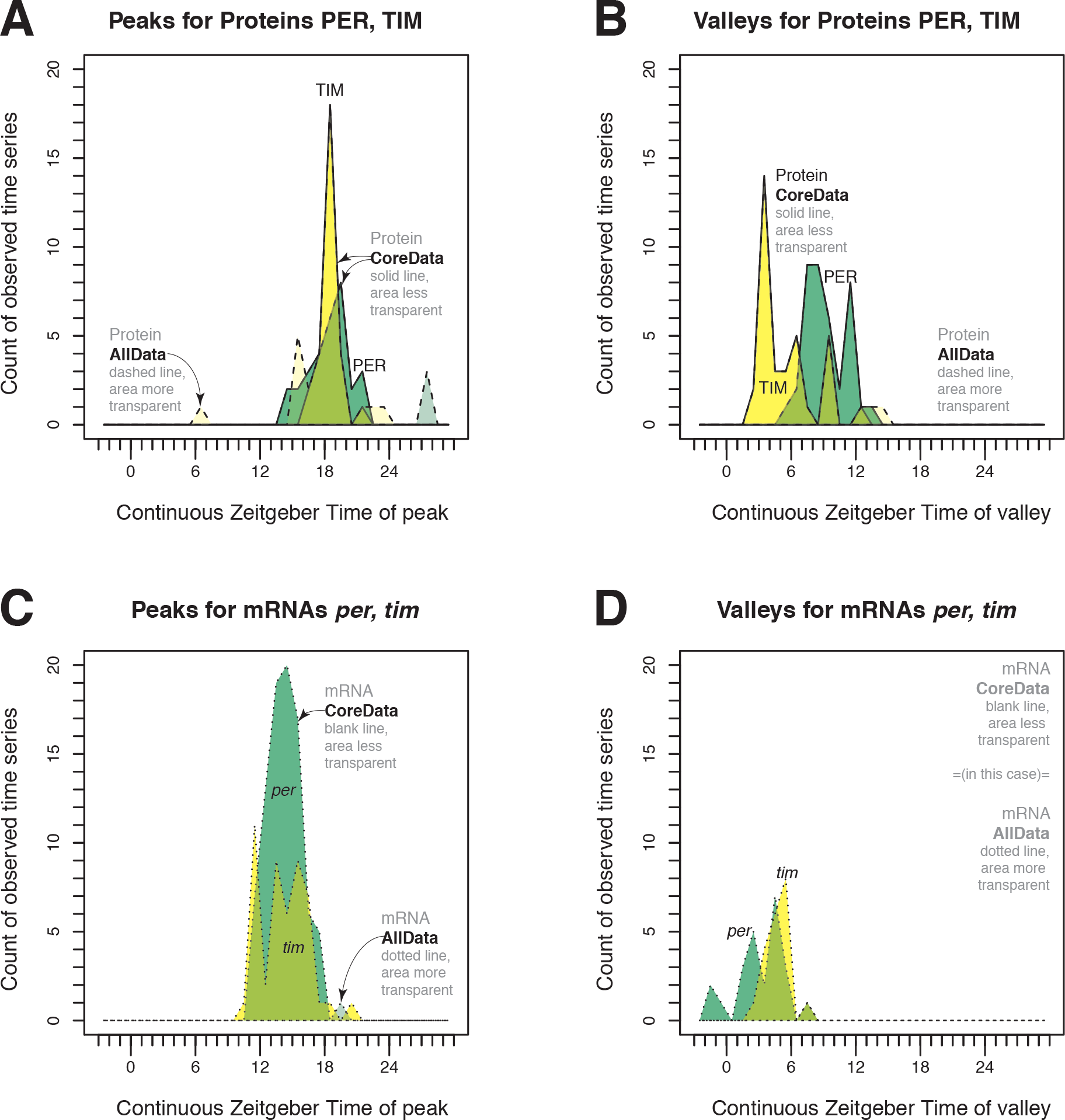
Histograms for *per* and *tim* mRNA and protein time series traits like peaks and valleys as observed in independent experiments show how FlyClockbase enables comparing variability. Here we compare the variability of timing for peaks (A, C) and valleys (B, D) of the proteins (A, B) and mRNAs (C, D) for the genes *per* (green) and *tim* (yellow). For all panels (A-D), we provide overlapping transparent histograms of all data (more transparent, dashed or dotted line) and of the core dataset from which outliers were removed by a boxplot method with Carling’s modification (less transparent, solid or no line). Descriptive summary statistics of the ‘CoreData’ distributions (excluding outliers) are given in Figure 6 and measures of statistical significance of differences are reported in Table 6 of the Results Section, and in the Supplemental Statistical Analysis (the annotated PDF of plots that were auto-generated by the script that also produced the plots in this figure; for an overview of calculations in the script, see Statistical Methods; for details, see R code in Supplemental Material). We assessed the impact of manually identified rare extreme outliers by excluding them from a copy of the dataset (termed “NoXtremes”) for which we repeated all analyses, including R’s automated outlier detection using Carling’s modification; the results are not substantially different and are reported in the Supplemental Statistical Analysis. For other related analyses see also Table 6, Table 7, Figure 6, Table 4, and Table S1.

#### Data quality

Since variances are easily inflated by outliers caused by errors in data handling, we aimed to obtain assessments of peak and valley times that were as accurate as possible given the observations reported by the studies in FlyClockbase. The desire for improving the overall rigor of our variability assessment motivated the various procedures described above for identifying outliers and investigating potential errors that might have artificially inflated estimates of variability. Thus, we manually re-inspected all steps of inference for each peak and valley data point that contributed to our final calculations of timing statistics. We started with the already highly refined dataset *ObsMod5* and created *ObsMod6* by correcting all irregularities from data handling or processing on our side that we could detect in FlyClockbase or our *TraitTables* (see Figure 3). Starting from the figures of publications, we checked the results of all our manual operations that could have modified traits, up to the final values used as input for our R script that produced Figure 7. The statistics of the resulting human error analysis is shown in Table 5.

##### Main analysis with outlier robustness

We decided to carefully investigate the role of outliers when analyzing variance in order to arrive at robust conclusions. We were motivated by the following considerations:

1. *Use of robust statistics*. We aimed to use state-of-the-art statistical methods designed for delivering robust conclusions that minimize the chances that a few odd values have an unduly large impact on the overall conclusions (see Wilcox (2012) for an introduction to robust estimation and hypothesis testing).
2. *Dealing with rare ObsOdd*. As described above, we checked and re-checked all observed peak and valley times that entered our final comparisons between *per* and *tim* components, aiming for the best interpretation of each reported time series. Despite this scrutiny, there were four observed time series that we could not interpret convincingly. All four extreme values pertain to protein peak time, with one extreme for TIM (time series ID 14.1.1, peak at 6.966*h* linearized *CZT*) and three extremes for PER (time series IDs 43.2.1, 43.3.1, and 43.5.1, all with a peak at 28*h* linearized *CZT* and all from one study). All four also showed a clear signal in their original figure that appeared to have been analyzed correctly (based on our reading of the respective studies), yet all four reported times appeared to be clearly distinct from the distribution of times reported by all other similar studies. For example, TIM extreme outlier time series above observed a peak almost exactly 12*h* away from the mean of the equivalent TIM distribution of peak timings without outliers. While we could not find any indication that morning and evening had been flipped in the corresponding study, it is very difficult to exclude such human errors in light of the many challenges that easily frustrate data analyses in spreadsheet software (see Discussion below). We stopped further attempts of re-interpretation, since the original raw data was not publicly available. As revisited in the Discussion, these values could represent any of the following: genuine observations of typical behavior of rare fly clocks, rare behavior of typical fly clocks (at least sustained for the length of the time series), some rare combination of measurement protocol details that conspired to systematically bias observations (which were correctly interpreted), or human errors leading to misinterpretations in the complex chain of time series observation and analysis. We could not find similarly extreme outliers in corresponding observations of mRNA. We consider the overall evidence to be too incomplete, contradictory and therefore inconclusive to determine which process may contribute most. Observing these extreme outliers might raise the distant possibility that apparent wildtype circadian clock systems can exhibit extreme deviations from their normal timing behavior in a few percent of occasions. However, exploring the possibilities of such exotic behaviors is beyond the scope of this study and not possible without a more elaborate error-management for observation and analysis of time series in circadian clock studies. It would also require a substantial number of individual time series observations in the *DetailSection* of FlyClockbase to enable the independent computation of summary statistics. To test whether these handpicked extreme outliers made a difference to our conclusions, we grouped and denoted them in our Statistical Methods Section as ‘outlier removal approach (ii)’. We then created a parallel analysis track in our R script that took every computation on the full dataset (‘*Mod7*’) and repeated it on a manually created copy of the input files (‘*Mod8*’), where this manual outlier removal approach (ii) had been applied. To increase clarity, ‘*Mod7*’ or ‘*Mod8*’ are also labeled, respectively, “*BestEachObs*” or “*BestNoXtrem*” in our R code, and “with Extremes” or “no Extremes” in the titles of the automatically generated plots. (‘*Mod7*’ is essentially identical with ‘*Mod6*’ except for trivial changes to facilitate automated reading from R; the *Mod7* and *Mod8* files are only stored next to our R script that reads them as input, they are not stored in the main time series trait folder). As can be seen in the Supplemental Statistical Analysis online, the removal of these four specific time series does not substantially change our conclusions.
3. We were unsatisfied with the subjective nature of the decisions summarized above as ‘outlier removal method (ii)’. While extreme outliers are reasonably easy to detect, there is a gradual transition to less extreme values, where subjective decisions about the inclusion of particular time series could easily lead to a new set of problems by creating ascertainment biases that are impossible to correct for statistically. Thus, we decided to employ a principled method. After some experimentation, we arrived at outlier removal approach (iii) which has been described elsewhere (Carling 2000; Wilcox 2012); see Statistical Methods.

Our results in Table 6 report our current best analysis of the most reliable data on the variability of *per* and *tim* protein and mRNA peak and valley timing accessible to us in FlyClockbase. For the reasons given above, we decided to exclude outliers as identified by our approach (iii) see Materials and Methods (Carling 2000; Wilcox 2012).

#### Alternative handling of outliers

In the Supplemental Statistical Analysis we compared results after removing outliers using approach (iii) as shown in Table 6 with those obtained from the full dataset to investigate whether removing outliers affects conclusions. Given the extraordinary range of timing variability for the peaks of PER and TIM, it is unsurprising that the difference in variance reported in Table 6 loses statistical significance when outliers are included. In Table 7 we summarize our results of comparing different outlier approaches. Corresponding values for all other results in Table 6 can be extracted from the Supplemental Statistical Analysis.

We conclude from Table 6 that the majority of circadian clocks in *D. melanogaster* are significantly more variable in their timing of PER peaks in comparison to TIM peaks (P<5%). In a minority of cases outliers can introduce such large amounts of variability that indicators of significance are swamped and a loss of significance is perceived (see Table 7 and Discussion). All other comparisons shown in the Supplemental Statistical Analysis confirm this overview. Our observation of significant differences in variability contrasts with the near absence of differences in the average timing of these peaks.

**TABLE 6.** Timing distribution differences tested between *per* and *tim* to compare averages and variances of peak and valley times for proteins and mRNAs after removing outliers.

|  | Point distribution<br>CZT Observed [h] |  | Differences Distribution<br>of CZT point distributions [h] |  |  | Test |
| --- | --- | --- | --- | --- | --- | --- |
|  | *per* | *tim* | Lo 2.5% | Middle 50% | Hi 97.5% | P-Value |
| **Protein peak** |  |  |  |  |  |  |
| *Var* | 4.1377 | 0.89211 | 0.97136 | 3.2456 | 5.6033 | **\* (*P* < 5%)** |
| *Avg* | 18.644 | 18.915 | -0.99996 | -1.2345x10<sup>-5</sup> | 0.99994 | 0.98753 |
| *n* | 91% =30/33 | 78% =29/37 |  |  |  |  |
| **mRNA peak** |  |  |  |  |  |  |
| *Var* | 2.4877 | 4.4906 | -4.1225 | -2.0029 | -0.0035509 | **\* (*P* < 5%)** |
| *Avg* | 14.606 | 14.587 | -0.83733 | 4.3138x10<sup>-6</sup> | 0.99996 | 0.66505 |
| *n* | 99% =87/88 | 100% =48/48 |  |  |  |  |
| **Protein valley** |  |  |  |  |  |  |
| *Var* | 3.7568 | 6.3773 | -6.8437 | -2.6205 | 1.3534 | (*P* > 5%) |
| *Avg* | 9.4058 | 5.8421 | 3.00 | 4.00 | 5.00 | **1.489x10<sup>-7</sup>** |
| *n* | 100% =39/39 | 94% =34/36 |  |  |  |  |
| **mRNA valley** |  |  |  |  |  |  |
| *Var* | 5.3608 | 1.3215 | 0.17418 | 4.0393 | 8.3148 | **\* (*P* < 5%)** |
| *Avg* | 3.606 | 5.0894 | -2.0611 | -1.00 | -1.8909x10<sup>-5</sup> | **0.012835** |
| *n* | 100% =24/24 | 100% =20/20 |  |  |  |  |

**TABLE 7.** Impact of outlier removal approaches on differences in variance of protein peak times (PER – TIM)

| **Outlier removal approach** | **Outliers counted / All data** | **PER-TIM Variance Lo 2.5% CI** | **PER-TIM Variance Mi 50%** | **PER-TIM Variance Hi 97.5% CI** | **Significant at $P = 0.05$** |
| --- | --- | --- | --- | --- | --- |
| No outliers removed | 0/33 (PER)<br>0/37 (TIM) | **-8.308** | 4.7564 | 15.755 | *no* |
| Removal type (ii): Manual extremes | 3/30 (PER)<br>1/36 (TIM) | **-1.7526** | 1.3432 | 4.2201 | *no* |
| Removal type (iii): Carling (2000) | 3/30 (PER)<br>8/29 (TIM) | 0.97136 | 3.2456 | 5.6033 | *yes* |

#### Other peak comparisons

We find differences in the variability of mRNA peak times, albeit interestingly with inverted sign, compared across *per-tim* equivalent parts. This means that significantly lower variability in *per* mRNA peaks precedes significantly larger variability in *PER* protein. Conversely, significantly larger variability in *tim* mRNA precedes significantly lower variability in *TIM* protein. This flip indicates that the differences in variability of the peaks for PER and TIM are not caused by corresponding differences in the variance of their mRNA peaks. We therefore conclude that these differences are caused by mechanisms affecting variances of peaks in the causal reaction networks after the transcription of *per* and *tim*. These differences in variance contrast with non-significant differences in the average peak timing of the same peak time distributions.

#### Valley comparisons

The valleys of PER and TIM occur at significantly different average times, irrespective of how many outliers are included. However, the non-significant differences in variances reported in Table 6 become barely significant at the 5% level when adding the two outliers observed in TIM. When comparing the valleys of *per* and *tim* mRNAs, we found significant differences in average time; *per* valleys precede *tim* valleys, even though averages do not differ among peaks. Valleys of *per* and *tim* also showed significant differences in variance, with *per* being more variable than *tim*. We also report an inversion of variances when compared to their corresponding mRNA peaks. Our results for the valleys of *per* and *tim* mRNAs are not impacted by outliers, since none of our approaches to outlier analysis could identify any outliers among the 24 and 20 observations in FlyClockbase, respectively.

### Comparing methods for measuring *TimeSeries* of *per* mRNA

In addition to comparing different clock components, FlyClockbase can contrast experimental details that differ between independently observed time series of the same clock component. These experimental details are stored as *TimeSeries Attributes* in FlyClockbase and used for extracting corresponding sets of *TimeSeries* or *TimeSeries Traits* for additional statistical analyses. Here we compared *per* mRNA time series recorded by different measurement methods. We chose *per* mRNA because it is the most common time series in FlyClockbase and thus provides the largest statistical power for detecting potential systematic biases. Given the differences in variances between clock components reported above, we wanted to know if such differences could have been produced by using different methods of obtaining time series.

#### Measurement methods

The following five measurement methods were used to collect at least three *per* mRNA time series in FlyClockbase: microarray, nascent-seq and RNA-seq, PCR, RNase protection assays (RPAs), and Northern Blots. Two of the four time series measured with microarrays were outliers, and all four of the time series measured with RNA-seq or nascent-seq were from a single study. These time series were consequently eliminated from our analyses. Many time series did not include a valley time for *per* mRNA, so we focused solely on *per* peak time.

##### Comparing means

As expected, we could not find any significant differences in the averages of peak times observed by different measurement methods when comparing PCR vs Northern Blot, PCR vs RNAse Protection Assay (RPA), and Northern Blot vs RPA (using the Mann-Whitney-U test at the 5% level, see Supplemental Statistical Analysis).

##### Comparing variances

We found significant differences in variance (using *comvar2*) to compare measurement methods as above. Comparing the combined peaks from time series measured with any PCR method to the pool of those measured with RPA or Northern Blot resulted in significantly different variances for both. The 95% CI of differences of variance reported by *comvar2* after 10^5^ bootstraps for RPA and Northern Blots are 0.4413 to 6.817 and 0.4764 to 10.586, respectively (for explanation on extra digits, see Table 6). The histograms of the corresponding distributions are given in Figures 8A and 8B.

**FIGURE 8.**
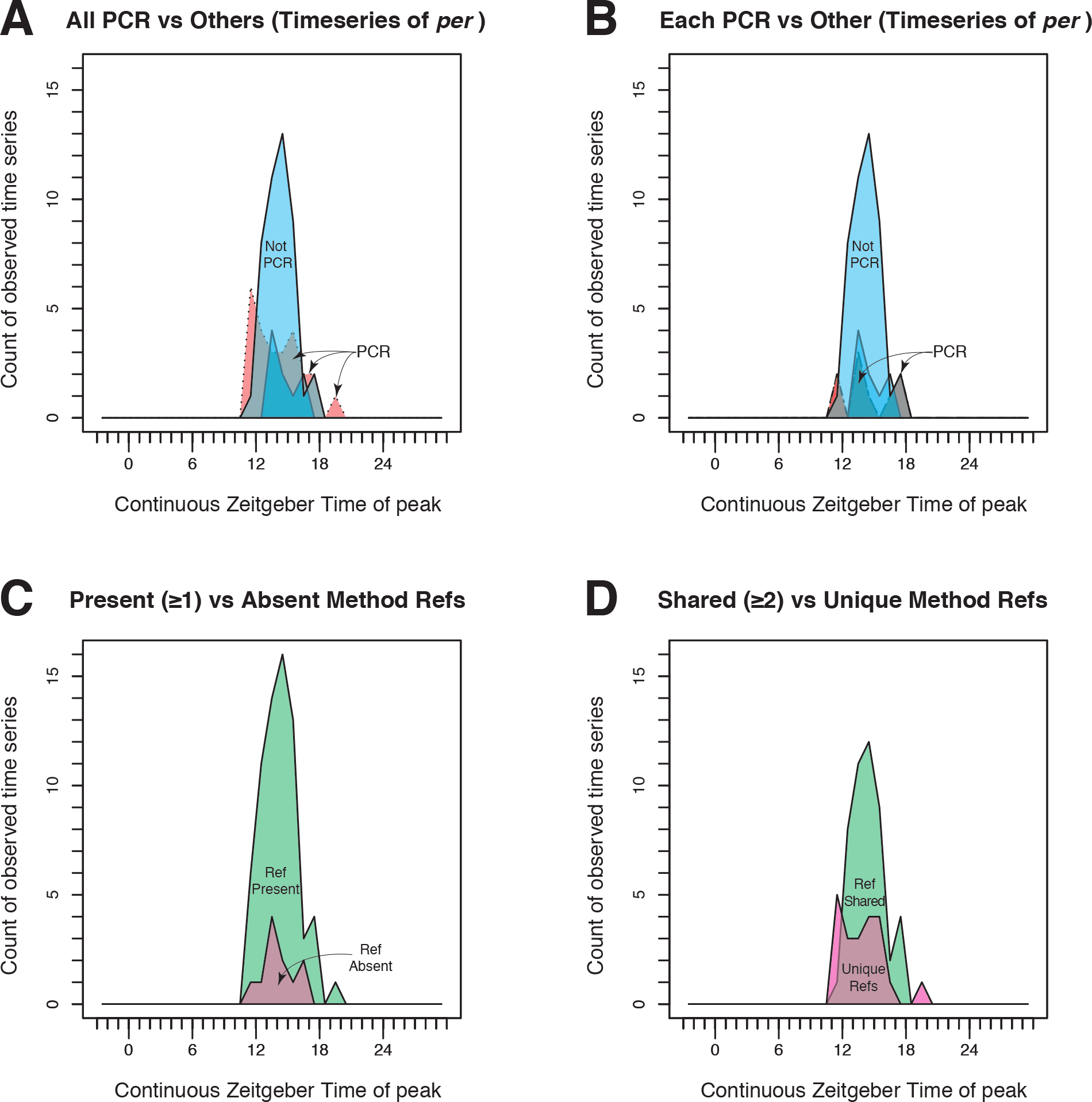
Comparisons of PCR vs non-PCR methods for measuring amounts of per mRNA in time series as enabled by FlyClockbase. Here we compare the variability of peak timing in all available time series of *per* mRNA without grouping by study, in order to contrast distributions of peak times obtained from time series (A) observed by all PCR methods combined (red) vs (blue) all non-PCR methods combined (blue); (B) observed by different PCR methods like qPCR or RT-PCR individually (red) vs (blue) all non-PCR methods combined (blue); (C) observed in studies citing at least one method reference (green) vs (purple) studies with no method reference (purple); and (D) observed in studies sharing two or more method references (green) vs (purple) studies with method references unique among studies contributing to *per* mRNA observations in FlyClockbase (*and* citing at least one method reference, purple). For all panels (A-D), we provide overlapping transparent histograms. Measures of statistical significance of differences are reported in the Results Section and parts of the Supplemental Statistical Analysis. For related data and analyses see also Table 8, Figure 9, and Figure 10.

The differences in variances are strongest for peak times measured with PCR or Northern Blots. Comparing the nine qPCR observations with the nine Northern Blots showed significantly higher variance in qPCR observations (*comvar2*, 10^5^ bootstraps, 95% CI for differences in variance: 0.353 to 7.598). Likewise, the 16 RT-PCR observations in FlyClockbase are more variable than the nine Northern Blots (*comvar2*, 10^5^ bootstraps, 95% CI for differences in variance: 0.46889 to 11.486). As reported above, comparing peaks from PCR to those from RPAs showed significant differences in variance. We expected this pattern to hold also for non-pooled PCR, but found results to be no longer statistically significant. Each type of PCR had a very skewed distribution of differences in variance with a substantial bias similar to that of Northern Blots (see *comvar2* results in the Supplemental Statistical Analysis). However, reduced sample size diminished statistical power, so differences are no longer significant.

#### Method references

We were surprised by the larger variance in PCR results compared to results from non-PCR methods. We speculated that this might not necessarily be inherent to PCR but could be caused by a larger diversity of method protocols. If this were general, we would expect studies with shared method protocol references to report results with less variability than studies which did not share such references. Therefore, in studies that did not cite any method references, we might suspect a larger diversity of method protocol variants, and thus more diverse results.

**FIGURE 9.**
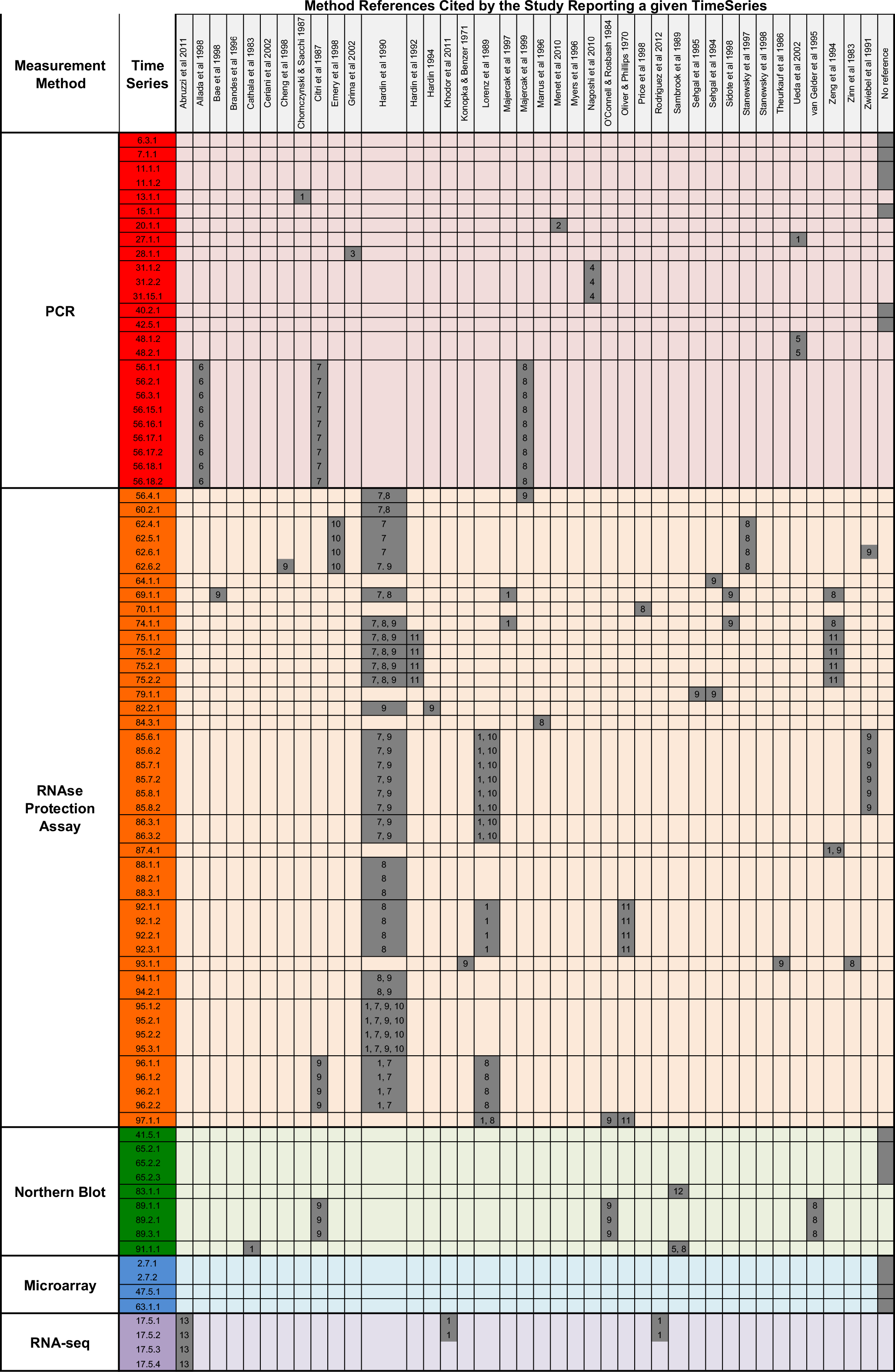
Referenced protocol use similarity matrix for studies reporting *per* mRNA time series, grouping protocols by measurement approach. The vertical axis shows time series IDs of *per* mRNA time series grouped by broader measurement method, and the horizontal axis indicates the experimental protocols references cited. The numbers given in the table for each time series and reference refer to a classification key indicating which of several broader types of methods had been used (1 – RNA extraction; 2 – Expression analysis; 3 – SYBR green detection protocol; 4 – RNA quantification; 5 – RNA purification; 6 – serial dilutions; 7 – controls; 8 – general protocol; 9 – probes; 10 – hybridization, RNase digestion, electrophoretic product separation; 11 – dissection; 12 – transfer to gel, labeling probes, hybridization, washing; 13 – PCR for cDNA library). Boxes shaded in gray represent the references cited by each time series, and the numbers in the gray boxes correspond to the broader types of methods to which this specific part of a protocol reference belongs. Gray boxes shaded in the “no reference” column indicate time series from studies without method references. All time series from the same study measured with a given method are outlined in dark grey. For related data and analyses see also Table 8, Figure 8, Figure 10, and parts of the Supplemental Statistical Analysis. Table 8 provides summaries of counts of time series and studies based on information in this table. The details of experimental protocols are described in their respective references (Oliver and Phillips 1970; Konopka and Benzer 1971; Cathala *et al*. 1983; Zinn *et al*. 1983; Oconnell and Rosbash 1984; Theurkauf *et al*. 1986; Chomczynski and Sacchi 1987; Citri *et al*. 1987; Lorenz *et al*. 1989; Sambrook *et al*. 1989; Hardin *et al*. 1990; Zwiebel *et al*. 1991a; Hardin and Hall 1992; Hardin 1994; Sehgal *et al*. 1994; Zeng *et al*. 1994; Sehgal *et al*. 1995; van Gelder *et al*. 1995; Marrus *et al*. 1996; Myers *et al*. 1996; Majercak *et al*. 1997; Stanewsky *et al*. 1997; Allada *et al*. 1998; Bae *et al*. 1998; Cheng *et al*. 1998; Emery *et al*. 1998; Price *et al*. 1998; Sidote *et al*. 1998; Stanewsky *et al*. 1998; Majercak *et al*. 1999; Ceriani *et al*. 2002; Grima *et al*. 2002; Ueda *et al*. 2002; Menet *et al*. 2010; Nagoshi *et al*. 2010; Abruzzi *et al*. 2011; Khodor *et al*. 2011; Rodriguez *et al*. 2013).

Figure 9 provides detailed information about the method protocol references cited in the studies we used. As before, we focused on the peak times of individual time series as observed by PCR, RPAs, or Northern Blots. Some authors largely followed the methods outlined in other referenced studies, while others only incorporated protocol references for specific aspects such as probe development, the use of controls, or RNA extraction (see classification key in the caption of Figure 9 for a list of some methodological aspects that might be of interest).

Table 8 provides summary counts of the time series represented in the various broader categories of Figure 9 and the number of studies that observed them. Table 8 also provides summary counts of the time series from studies that either share or don’t share references with each other (along with the corresponding numbers of studies; this classification excluded studies with no references). The underlying network of protocol identities is represented in Figure 9, and histograms of the corresponding distributions are given in Figure 8CD.

**TABLE 8.**
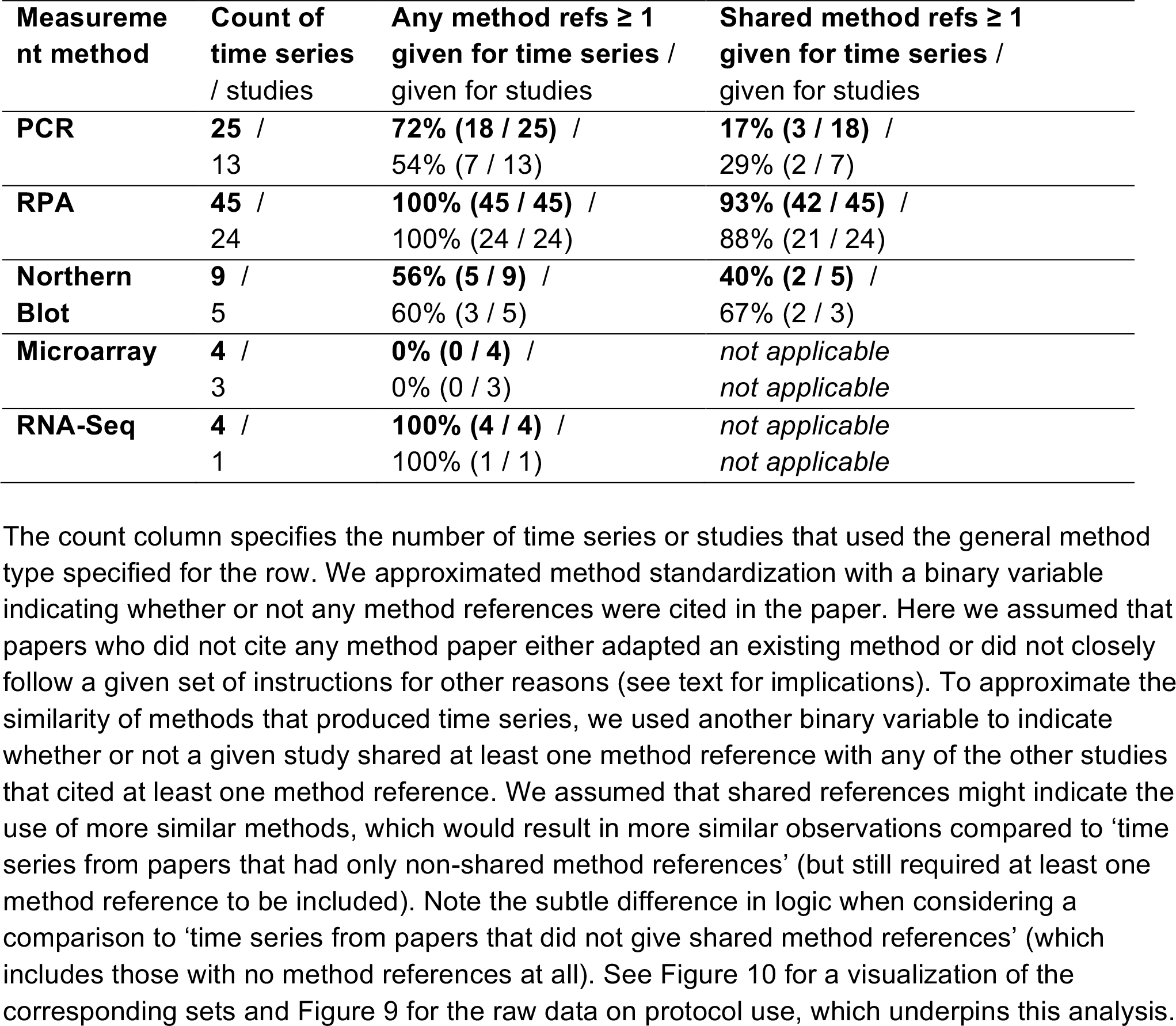
Standardization proxies: overview of potential approximations for quantifying method detail standardization and method detail similarity for independent observations of *per* mRNA.

##### Presence or absence of method references

We grouped all analyzed time series into two categories, which might be nicknamed “*Any Method Ref*” if their studies provided at least one reference for details of the experimental protocols used to observe time series using a given broad type of method, or “*No Method Ref*” if not a single relevant experimental protocol reference was given. Comparing average peak times among these groups showed no significant differences and no detectable indication of bias (Mann-Whitney-U test, 95% CI: -1.00 to 1.00, p = 0.736). While differences in variance were not significant, a clear bias in variance was noticeable in the distribution of differences, possibly indicative of significance if higher sample sizes were available (*comvar2*, 10^5^ bootstraps, 95% CI for differences in variance: -1.385 to 5.6876).

##### Presence or absence of shared method references

We subdivided all available time series with at least one method reference into those with and without shared method references (see Figure 9 and Table 8). Average peak times did not change significantly if method references were or were not shared, but some bias could be seen (Mann-Whitney test, 95% CI for difference in location: -2.00 to 5.4208x10^-5^, *P* = 0.1829). Similarly, variances of peak times showed bias but were not significantly different (*comvar2*, 10^5^ bootstraps, 95% CI for differences in variance: -1.1045 to 6.325, see Supplemental Statistical Analysis).

#### Reduction of statistical power by application of inappropriate logic?

The last groups above had both clear biases in their differences in variance as revealed by *comvar2*, but neither test was significant at the 5% level. Combining both results in a slightly different way might increase relevant sample sizes and significance. As shown in Figure 10, we currently exclude studies with not a single method reference from our analysis of studies with shared references. We initially justified this by assuming that method references must be present for deciding whether studies could have used the same methods at the lab-bench or not. It turns out that this line of reasoning might say more about the need for greater clarify in logic formalisms than about the effects of sharing references or not. We will next make explicit, what we unknowingly implied above; we do so in order to demonstrate the difficulties of creating clear statements in words chosen *adhoc* and aimed at clarifying statements in a limiting logic formalism.

Above we assume implicitly that a study without any method protocol references may or may not have used the exact same methodological procedures as another study without method references. Applying Boolean logic here attempts to force a

> Yes (OK)
>
> No (KO)

answer to the question about the use of a common protocol. This logic fails here, because it does not allow us to specify that we cannot provide any certain answer based on the absence of a reference (we cannot exclude the possibility that two studies may have shared the same protocol). Instead, we might consider the BioBinary logic that FlyClockbase is adopting from Evolvix (see Supplemental Material). This is a step forward, because it can distinguish the cases of

> MIS (‘inapplicable’ because ‘no-method-given’)
>
> OKO (‘somewhere between OK and KO’)

from the other cases (OK/KO), where we can reach relatively more clear decisions because method protocol references are given. Our statistical analysis above can now be rephrased as stating that the absence of a reference is a BioBinary ‘MIS’ (since we formally cannot answer the question due to a missing value). As a result, we would exclude such values, since it is always easy to name multitudes of irrelevant values that clearly should not have any influence on our analysis. This interpretation might appear unsatisfactory at first, but it is still a step forward, because it is explicitly stated and therefore more tangible, which might attract further scrutiny.

If explained clearly, most experimental biologists will probably not hesitate to point out that the chances of using identical experimental protocols in the lab are miniscule unless there is a shared reference to a common protocol (which would probably be referenced). There are simply too many variations that are done most easily. Thus, instead of assuming the absence of evidence (which might allow for shared protocols, even if no references are given), we can assume evidence of absence (since it is rather unlikely that two labs use the same procedures, without sharing references to a common protocols). Given the many variations of PCR that are easily created in the absence of further detailed instruction, it is difficult to see, how labs might accidentally share procedures. Thus, technically, this case must be classified as a BioBinary OKO; however, overwhelming evidence suggests that a KO (not sharing) is much more likely.

##### Result

These considerations suggest that it would be more appropriate to add the time-series with no references to the group of those that have only one (non-shared) reference. This might increase statistical power enough to make tests significant that compare “shared vs not shared references”. Pivotally, these considerations demonstrate the importance of using the right type of logic and type system. It is easy to miss important points in Boolean logic without appropriate visualization. Since logic formalisms are best automatically supported at the compiler level, a step change in speed and quality of many biological analyses could be facilitated by a compiler that correctly map BioBinary and more expressive logic formalisms to data – out of the box. This conclusion is general for the construction of *VBIR*s, and independent from the particular outcomes of any specific tests that might be performed, because the existence of such unresolved difficulties in representing formal logic can easily create bugs that bind large amounts of research time, which could otherwise be dedicated to *VBIR* development.

**FIGURE 10:**
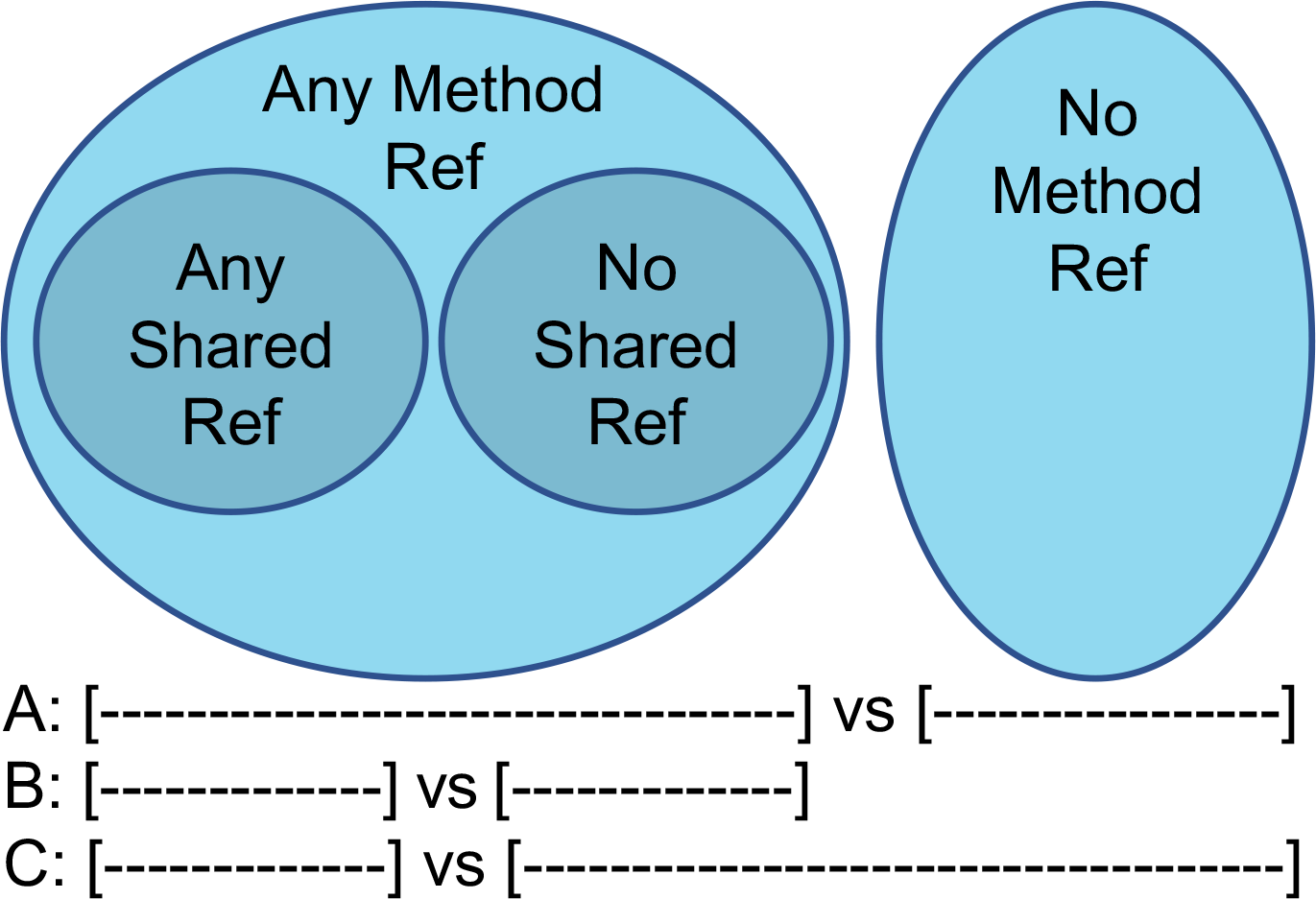
Logic of datasets in statistics: example for an interaction of statistics and formal logic where the logic for selecting datasets unnecessarily reduced the power of a statistical test. Here we illustrate the logic we used for constructing the datasets that group observed time series into the sets used for approximating the degree of method standardization and method similarities that produced the per mRNA time series observations in FlyClockbase. We denote as ‘any’ an individual time series from an individual study with a count of references to experimental protocols, where the count is ≥1; ‘no’ stands for a count of 0. Grouping **A** and **B** indicate test groupings performed by the R script in the Supplemental Material. Grouping **C** indicates the grouping suggested by a more thorough logical analysis performed after completing the statistical tests presented in the Results Section. For related data and analyses see also Table 8, Figure 8, Figure 9, and parts of the Supplemental Statistical Analysis.

#### Comparing methods by study to reduce multiple comparisons

Figure 9 shows that a single study often reports multiple time series, which could bias the results above (based on analyzing each time series individually without grouping into studies). Thus, big studies contributing many time series might exert an unduly large influence on the results. For example, nine of the 25 time series measured with PCR (36%) are from a single study (Majercak *et al*. 2004), but this study is only 1 of 13 (7.7%) that used PCR to measure *per* mRNA. Given the many complexities of this system, the most appropriate way to address this problem is unclear. Treating each time series independently might give large studies too much influence. However, using only a single value from each study (e.g. the mean), irrespective of how many time series it represents, could give small studies too much weight and thus risk adding irrelevant and noisy artificial variance. The latter approach is therefore an extreme approach of countering potential problems with the former. We used this alterative study-based approach to explore the robustness of our conclusions about measurement methods, albeit with the added caution that the loss off statistical power might be too large for reaching clear conclusions.

Indeed, using only a single average value from each study reduces the remaining statistical power so much that no result remains significant (see Supplemental Statistical Analysis). It is worth noting though that some comparisons of variance still showed such a strong bias that the collection of additional observations might result in significance. The comparisons that approached significance were these: Variance between PCR and Northern Blots (*comvar2*, 95% CI: -0.281 to 13.340, with a point estimate for a difference in variance of 4.926 h^2^); Variance between PCR and RPA studies (*comvar2*, 95% CI: -0.161 to 8.426, with a point estimate for a difference in variance of 4.065 h^2^). Such comparisons of non-PCR methods vs PCR-methods were closer to significance than the non-PCR method comparison of Northern Blots vs RPA studies (*comvar2*, 95% CI: -3.492 to 0.538, estimated difference -0.861 *h*^2^). Differences in variance based on shared references produced even weaker signals (but see Figure 10 and Discussion of Logic for a potential boost of statistical power). All corresponding details are recorded in the Supplemental Statistical Analysis.

## DISCUSSION

We discuss the more technical aspects of our work in the context of three broader aims of this study: (i) introducing FlyClockbase and connecting it to our current understanding of circadian clocks; (ii) using FlyClockbase to ask new questions about variability in circadian clock time series, possibly illuminating important aspects of clock mechanisms and methods of observation; (iii) improve and simplify how FlyClockbase, and by extension similar biological information resources (*VBIR*s), are constructed, expanded, and maintained.

### Why learn lessons on data?

It would be easy to finalize this study without the third aim. However, it is impossible to address the timely and relevant topic of organizing biological data without the concrete context of a specific resource like FlyClockbase. This discussion is relevant because the increasingly data-centric nature of biological discovery has resulted in calls for improved access to existing data (NIH *et al*. 2012; Read *et al*. 2015; Wilkinson *et al*. 2016), which is easier said than done (Goldston 2008; Doan *et al*. 2012; Gitelman 2013; Huang and Gottardo 2013; Mccallum 2013). As physical access to data is increasing, the next frontier is defined by the ability to *efficiently* identify datasets of relevance for a given topic. The diversity of biological questions would make any one single resource for all biologists too cumbersome to use. Instead, this aim could be achieved more efficiently by empowering research communities to construct resources for their own contexts, albeit using a shared interoperable infrastructure. This infrastructure will be perceived as useful to the degree it can provide standards that convincingly address common challenges faced by all biologists aiming to construct *VBIRs* for organizing notoriously uncertain biological data. Developing such standards is an enormous task that requires the integration of lessons from many more studies than any single effort such as this one could produce. Thus, our more modest aim here is to highlight lessons we learned while constructing FlyClockbase, hoping they will be useful as the biological community works towards finding more general solutions (NIH *et al*. 2012; Read *et al*. 2015; Wilkinson *et al*. 2016).

### Why do we need many *VBIRs*?

Efficiently constructing many *VBIRs* like FlyClockbase is necessary for integrating biological information at the scale needed for current research. The need is driven by immense challenges, such as mechanistically understanding and curing cancer (NIH *et al*. 2016; Samuels *et al*. 2016), mapping genotypes to phenotypes in personalized medicine and elsewhere (Roden 2011; Mackay *et al*. 2012; Kirk *et al*. 2015; Ashley 2016), or the evolutionary systems biology goal of mechanistically predicting realistic fitness landscapes (Loewe 2016). Irrespective of whether it is possible to realize these broader visions, any serious attempt will require the diligent construction of many interoperable *VBIRs* that connect well to state of the art expertise, and advance biological research in the relevant areas. Thus, we will next examine FlyClockbase in this respect.

### FlyClockbase is consistent with current hypotheses

Overall, time series in FlyClockbase are consistent with general published clock knowledge and with mechanisms currently thought to control the clock. Informative reviews of the clock that draw on previously published experimental studies are given elsewhere (Hardin 2011; Özkaya and Rosato 2012). Conclusions from these reviews are supported by FlyClockbase summary statistics given in Table 4.

#### CLK and other proteins

Increasing amounts of VRI protein between about ZT4 and ZT16 repress *clk* transcription, with an especially pronounced effect after ZT14 (Cyran *et al*. 2003; Glossop *et al*. 2003). Although CLK protein typically functions as a transcriptional activator for *vri*, the formation of the PER/DBT/CLK/CYC complex represses the activity of CLK between approximately ZT16 and ZT4 (Hardin 2011). *vri* transcription is therefore decreased, causing lower levels of VRI and reducing the repression of *clk* transcription by VRI. This allows *clk* mRNA to increase and reach a maximum around dawn (Allada *et al*. 1998; Özkaya and Rosato 2012). *clk* mRNA levels also increase due to the action of the transcriptional activator PDP1 protein, which becomes especially strong around ZT18 (Cyran *et al*. 2003). Time series in FlyClockbase reflect this pattern of *clk* mRNA reaching a maximum in the morning (peak time = 2.79 *DZT* mean ± 3.16 h *SD*), decreasing throughout the day into the early night (valley time = 14.33 *DZT* mean ± 1.67 h *SD*) and then beginning to increase again in the late night. Although *clk* mRNA does show rhythmic expression, the amount of CLK protein is not cyclic (Yu *et al*. 2006). Data from FlyClockbase support this constant expression, as the n=5 peaks and n=9 valleys observed for CLK overlap almost completely. However, there might not be enough observations to fully settle the issue, since the variation for both CLK traits (±3.18h, ±4.30h *SD*) is currently lower than expected for peaks and valleys that are all drawn from one stable uniform distribution (±>6.8h; see: Materials and Methods Section, Peak-Valley Section, Randomizing Time Section).

#### PER and TIM dynamics

PER protein, *per* mRNA, TIM protein, and *tim* mRNA have expression patterns which are generally opposite to that of *clk* mRNA. Transcription of *per* and *tim* begins mid-morning and is promoted by the transcriptional activator CLK (Allada *et al*. 1998; Darlington *et al*. 1998). FlyClockbase shows the peak time of *per* mRNA at mean *DZT* = 14.61 ± 1.58 h *SD* and of *tim* mRNA at mean *DZT* = 14.59 ± 2.12 h *SD* (using outlier removal method (iii) as in Figure 6; for outlier removal method (i) see Table 4). This supports data suggesting mRNA levels increase through the day and into early evening (Allada *et al*. 1998; Hardin 2011). Protein accumulation reportedly lags behind that of mRNA by about six to eight hours (Zwiebel *et al*. 1991b; Hardin 2011), though data from FlyClockbase support a shorter delay of around four to five hours (delay from mean *per* mRNA peak to mean PER peak = 4.03 or 4.84 h, delay from mean *tim* mRNA peak to mean TIM peak = 4.33 or 3.82 h, from Figure 6 or Table 4, respectively). Between approximately ZT18 and ZT4, the PER/DBT/CLK/CYC complex represses the activity of CLK (Lee *et al*. 1998; Lee *et al*. 1999; Bae *et al*. 2000; Hardin 2011). This leads to decreased transcriptional activation of *per* and *tim*, causing a decrease in *per* and *tim* mRNA levels, which is reflected in FlyClockbase as a mean *per* mRNA valley at *DZT* = 3.61 ± 2.32 h *SD* and a mean *tim* mRNA valley at *DZT* = 5.09 ± 1.15 h *SD* (Figure 6; difference in variance significant at 5% level). During the day, TIM protein is degraded in response to light (Naidoo *et al*. 1999; Busza *et al*. 2004; Ozturk *et al*. 2011), as indicated in FlyClockbase by an early mean valley at *DZT* = 5.84 ± 2.53 h *SD*. PER, which is typically stabilized by TIM, is then destabilized and more prone to phosphorylation and subsequent degradation (Gekakis *et al*. 1995; Kloss *et al*. 2001; Merbitz-Zahradnik and Wolf 2015). This finding is consistent with FlyClockbase reports of a late PER mean valley = 9.41 ± 1.94 h *SD* (Figure 6). The PER valley is significantly different from the TIM valley (*P* = 1.489x10^-7^) as determined by the Mann-Whitney-U test. Unless specified otherwise, similar subsequent tests are two-sided and unpaired. To minimize search time for readers, we added a few non-significant digits to many results in the main text to help streamline searches for the context of such results in the Supplementary Statistical Analyses. As the amount of PER decreases, inhibition of CLK-mediated transcription by the PER/CLK/CYC/DBT complex also decreases, and CLK resumes promoting transcription of *per* and *tim*.

### FlyClockbase facilitates hypothesis-driven research

Circadian clocks in *D. melanogaster* are molecular systems of substantial complexity which have been inspiring generations of researchers to construct numerous hypotheses about how they work. FlyClockbase can substantially contribute to various life-stages of a hypothesis.

#### Starting with a blank slate

FlyClockbase can set the stage by integrating existing observations. It is beyond the scope of this paper to discuss the many ways of extending FlyClockbase beyond its current goals, for example, by providing the possibility to add time series from mutants. As FlyClockbase accumulates more observations, its power will increase to help researchers to put new observations into context by comparing them with already integrated data. Such comparisons can inspire new hypotheses and help evaluate them quickly. For example, we discuss below the hypothesis that significant differences in the timing variability of PER and TIM are the result of mechanistic interactions integral to the operation of this clock. This hypothesis grew out of our observation that the difference in variability of the peak time in these proteins was larger than we expected.

#### Using *Attributes* to compare timing variability

Another way of generating new hypotheses using FlyClockbase is to draw on the many attributes stored for time series. This structured information classifies time series in FlyClockbase in rich ways: many groups of column entries are easily combined into hypotheses for identifying significant differences between different genotypes, strains, observation methods, environmental conditions and more. Some hypotheses may not be tested easily, as statistical significance may often require more data. However, FlyClockbase already has enough data for testing some hypotheses, and insufficient data might inspire new experiments for testing important ideas. In this way, FlyClockbase can even become a tool for planning experiments.

#### Existing hypotheses on timing

Testing ideas against the data in FlyClockbase will become increasingly efficient as increasing numbers of experimental results are integrated into FlyClockbase. This results in a win-win for research productivity. The first win is clear if sufficient data exists in FlyClockbase to test a hypothesis (saving time). If the necessary data is not available in FlyClockbase, the second win is highlighting the potential need for new experiments in areas with limited data. Researchers can then decide whether to fill this gap with new experiments or prioritize other research. Again, FlyClockbase can help propose experiments within its scope, which could broaden over time.

#### The strength of FlyClockbase

Whether FlyClockbase will contain enough data critically depends on (i) the ease of adding new studies in a consistent way, and (ii) the effort required for checking the integrity of any data fragment. These two core requirements drive our interests in usability and human error analysis as discussed elsewhere throughout this study. Hence, work with FlyClockbase highlights how various subtle yet time-consuming issues of data organization, automation, usability, and error management that are usually classified as “non-biological” can easily become limiting factors for advancing circadian clock research. Our work on FlyClockbase suggest that it is more efficient to use the rather systematic approach of automating as much as reasonable and produce corresponding *VBIRs* in batches. This enables efficiencies of scale similar to those necessary for completing the human genome project (Lander *et al*. 2001; Venter *et al*. 2001). In similar ways, *VBIRs* could help compare data, evaluate methodologies, extend current knowledge, stimulate new ideas, test hypotheses, and create new routes of inquiry. We will next illustrate how FlyClockbase improves scientific productivity for testing hypotheses in its scope, before returning to practical questions of usability, appropriate data models, and efficient implementation for *VBIR*s.

### Peak timing hypotheses and more: PER variance exceeds TIM variance

We chose to compare PER protein, TIM protein, *per* mRNA, and *tim* mRNA because these components are integral to the circadian clock. They interact with many other clock parts (Figure 1), and null mutants for each gene (*per*^01^ and *tim*^01^) lead to arrhythmicity (Konopka and Benzer 1971; Sehgal *et al*. 1994). Also, Table 4 shows that peak and valley observations of these four components were among the most abundant in FlyClockbase and thus best suited for testing differences for statistical significance.

#### Comparing averages

We first compared the mean peak and valley times for PER and TIM protein and for *per* and *tim* mRNA amounts. Neither the proteins nor the mRNA had significantly different mean peak times. The mean valley time for *tim* mRNA is significantly later than for *per* mRNA (Mann-Whitney-U test, *P* = 0.012835, see Supplementary Statistical Analyses for context; two related tests were shy of significance). It might be reasonable to expect this delay to propagate, such that first the peaks of mRNAs, then the peaks of proteins, and ultimately also the valleys of proteins might show a similar pattern of *tim* preceding *per*. However, this pattern is quickly broken, as the respective pairs of mRNA and protein share essentially mean peak times for TIM and PER that are statistically indistinguishable. Following the cycle to mean valley times for proteins even reverses into the opposite pattern: PER mean valley time is significantly later than TIM mean valley time (see Figure 6; *P* = 1.489x10^-7^ as reported above). Two related tests also indicated significance for all outlier removal approaches tested (see context in Supplementary Statistical Analyses). These timing patterns defy the simplistic expectation of merely propagating delays and suggest mechanistic causes. Demonstrating statistical significance with the help of FlyClockbase suggests that these patterns might be worth simulating in stochastic models that capture causal mechanisms and respect the discrete nature of molecules (and resulting variability).

#### Explaining averages

We propose that the earlier mean valley time for TIM can be explained by the rapid degradation of TIM in response to light (Busza *et al*. 2004). Still, TIM’s peak time at mean *DZT* 18.92 ± 0.94 h *SD* might deserve a closer look. TIM peaks in the middle of the dark period and not at its very end as might be expected if light was solely responsible for degrading TIM.

The TIM degradation pathway is well characterized and begins with the activation of CRY via a light-dependent conformational change (Berndt *et al*. 2007; Ozturk *et al*. 2011; Vaidya *et al*. 2013). This change allows CRY to bind TIM in the nucleus (Ceriani *et al*. 1999; Busza *et al*. 2004). The F-box protein JET then ubiquitinates TIM to promote degradation by the COP9 signalosome (Koh *et al*. 2006; Knowles *et al*. 2009). Following TIM degradation, CRY is also degraded in response to JET-mediated ubiquitination (Peschel *et al*. 2009). In-depth reviews of the TIM degradation pathway are given elsewhere (Hardin 2011; Peschel and Helfrich-FÖrster 2011).

While light-dependent TIM degradation could explain why TIM reaches its valley *before* PER, it cannot account for the observation that *tim* mRNA reaches its valley a bit *after per* mRNA. The discrepancy in valley time cannot be caused by differences in mRNA production, as mean peak times for *per* and *tim* mRNA are not significantly different. We found this irrespective of the test or outlier removal method. Differences in variance barely exceeded 5% significance, albeit only if we remove outliers by approach (iii). See Figure 6 for overview and Supplementary Statistical Analyses for details. Therefore, we suggest considering differences in *per* and *tim* mRNA degradation. These degradation patterns could be influenced by CURLED (see below).

#### Comparing variances

We also compared the variability of peak and valley times for the proteins PER and TIM and for the mRNAs *per* and *tim*. Table 6 reports that peak time is more variable for PER than TIM as indicated by differences in variance that exceed *P* = 0.05, albeit only if we remove outliers using our approach (iii) as described (Carling 2000). Table 7 and the Statistical Methods present the necessary nuances. In short, we are confident that the differences in variance that we observe in most time series are significant, and not easily attributed to:

- *Statistical flukes* for the overwhelming majority of PER/TIM time series reported (see Table 6 and Supplementary Statistical Analyses for details on the robust bootstrap-based tests performed on the 84% relevant protein peak time series that were not removed as outliers by Carling’s approach, i.e. 59 of 70 combined protein peak times of PER or TIM); or
- *Trivial data errors* or inappropriate data handling at our end (see Figure 3 for the substantial effort in checking FlyClockbase for errors that resulted in *Mod5*, which was used as starting point for manually re-checking every single time series that contributed to our observation of PER-TIM differences in variance and resulted in the correction of all errors in *Mod6* as reported in Table 5);

Thus, we reject the explanations above based on our work. In contrast, the potential explanations below for the origins of outliers are more difficult to reject and cannot be tested on a routine basis. It also appears unlikely to us that these explanations contribute more than occasional outliers to typical experimental observations. Therefore *H_basic_*, the default basic null-hypothesis for data from FlyClockbase, recommends above that outliers are removed as described by CARLING (Carling 2000). This method is ‘approach (iii)’ in our Statistical Methods and was chosen after comparing the features of related approaches for outlier removal as reviewed elsewhere (p. 97 in Wilcox, 2012).

#### Outliers

About 16% of all protein peak times (3/33 PER, 8/37 TIM) or 4.3% (14/325) of all peak valley traits in Table 6 have been identified as outliers by Carlings method. Including all outliers exhausts the statistical power that FlyClockbase can currently provide for investigating our differences of variance. As a result, statistical significance collapses (see Table 7; *P* > 5% in our tests). However, this observation is unlikely to affect our conclusion that some systematic biological mechanisms are probably responsible for producing the PER>TIM variance patterns we report. Indeed, the outliers raise intriguing questions about the sources of their variability. We cannot currently distinguish the following potential sources of variability that will be discussed separately.

##### Genetic background differences

A substantial minority of flies that are currently classified as wildtype could have circadian clocks with significant genetic differences. This hypothesis might not be as unlikely as it may appear when only considering core clock genes as shown in Figure 1. Carefully listing *all* genes with potential impact on clock timing quickly reveals much larger mutational targets. Clock models also depend on specific rates of transcription, translation and degradation. These processes are governed by huge molecular machines. Unless otherwise more harmful, mutations that significantly delay or accelerate these machines will affect circadian rhythms. Frequencies are unknown, but such mutations in the genetic background of a clock might occur more often than mutations in core clock genes with similar effects on timing. If true, these clock background mutations could contribute much to the natural genetic variation in fly sleep patterns, which can be substantial (Harbison *et al*. 2009a; Harbison *et al*. 2009b; Harbison *et al*. 2013). In addition, selective pressure on circadian clocks is substantial (Beaver *et al*. 2002; Sharma 2003; Yerushalmi and Green 2009). It generates observable latitudinal clines of allele frequencies (Costa *et al*. 1992; Rosato *et al*. 1996; Sawyer *et al*. 1997; Sharma 2003; Sandrelli *et al*. 2007; Hut *et al*. 2013). Evolutionary importance of individual clock components has been demonstrated for various clock genes (Beaver *et al*. 2002), including *tim* (Sandrelli *et al*. 2007), and *per*, which contains a repetitive region that increases mutation rates for length polymorphisms. The resulting mutational effects are apparently large enough to maintain a latitudinal cline (Costa *et al*. 1992; Rosato *et al*. 1996; Sawyer *et al*. 1997; Kyriacou *et al*. 2007; Weeks *et al*. 2007; Kyriacou *et al*. 2008). Thus, mechanistic differences between the circadian clocks of flies from the wild are likely to exist and may resurface unexpectedly in clock studies.

If relevant and substantial, such differences could greatly complicate construction and parameter estimation in *the* “wildtype *D. melanogaster* circadian clock model”. While numerous models have contributed towards this aim (see Figure 4), there has not yet been a *single* model that integrates *all* known data on the clock of a *single well-defined* natural genotype. This ambitious aim becomes much more complicated if natural variability in clock genes makes time series more variable. Such variability from natural clock variants could undermine the statistical power of parameter estimation methods for constructing a *single* clock model for a well-defined genotype.

Controlled observations of all data in a single line of fly descent is – in theory – an easy way out. However, it might be difficult in practice to observe one fly well enough to match the statistical power of results from many years of research by many labs. Such focus on a single genotype could also generate a rather unusual clock model if one of many rare mutants with large effects is present (Eyre-Walker and Keightley 2007; Harbison *et al*. 2013). Developing models of such precision could advance methods for personalized medicine (Hodson 2016). However, most Drosophila clock researchers will probably prefer less precise clock models that usually match more observations in typical flies. Such general models could be inferred by parameter estimation methods from sets of time series collected in many genotypes by various methods but excluding outlier time series using the systematic approaches we employed (Carling 2000).

##### Environmental or developmental differences for measured flies

Unrecognized environmental factors that vary among measurements might modulate genetically identical circadian clocks. If true, experimental protocols for observing circadian rhythms in Drosophila could be improved to increase accuracy of biological replicates. Given that different authors do not necessarily report the same set of environmental attributes, a first step towards improving experimental protocols might be to develop a standardized set of *TimeSeriesAttributes* for FlyClockbase that improve the precision of reports from ongoing studies. It has been demonstrated that age impacts the clock in flies (Umezaki *et al*. 2012). Environmental factors that affect development in ways that strongly impacts circadian rhythms could be a potential source of outliers.

##### Evolution in different lab environments

Consistent differences in selection can cause flies to follow different evolutionary trajectories and sometimes the results can be observed in the lab over a number of years (Leroi *et al*. 1994; Shabalina *et al*. 1997; Hoffmann *et al*. 2001; Haag-Liautard *et al*. 2007; Keightley *et al*. 2009). The flies generating the data in FlyClockbase might have lived in environments with differences significant enough to trigger some adaptive evolution over a number of years. FlyClockbase does not yet have enough statistical power to detect significant differences between strains – if they exist. For example, our initial internal screening showed no differences between wildtype, color modified strains (*yw* or *w1118*) or other strains. Currently, FlyClockbase does not have dedicated *TimeSeriesAttributes* for characterizing the environmental history of flies from the decade leading up to the measurements. FlyClockbase is ideally positioned for integrating such data, once it becomes available, and the necessary *TimeSeriesAttributes* have been developed. However, not all fly clock studies report the LD environment in which their strains evolved for the previous 250 generations (Kannan *et al*. 2012). We have no specific evidence to support the claim that evolution in the lab produced the outliers we observed. However, some statistically significant evolution of the *D. melanogaster* circadian clock was observed after applying a relevant selective pressure for 80 generations in the lab (Kannan *et al*. 2012). Also, note that *per* contains repetitive nucleotides in its DNA, which result in high mutation rates for repeat polymorphisms with adaptive significance (Rosato *et al*. 1996; Sawyer *et al*. 1997).

##### Human errors

Setting up, performing, or analyzing clock experiments are complex tasks, as are reporting experimental procedures, relevant labels, or analyzed data. Such operations are error-prone (see discussion below) and can make reproducibility a challenge (Bammler *et al*. 2005; Freedman *et al*. 2015a). If all 16% were the result of combining all human errors before publication, then the overall rate would be surprisingly close to the 14% human error rate that we measured in Table 5, and corrected before our final test of the hypothesis that PER variance exceeds TIM variance. Our ability to detect human errors before publication is very limited. Hence, we took published plots and their attributes at face value. We excluded time series that had ambiguities we could not resolve (e.g. from poor plot quality), however, this does not exclude human errors before publication (see Section on human errors in Supplemental Material).

##### Conclusion on outliers

Thus, we have no reason to assume that errors before publication could not have produced some of the 16%. It also seems unlikely that none of the other causes above could have contributed. Distinguishing between the hypotheses above is currently beyond the statistical power of FlyClockbase. However, this might be irrelevant for many of the aims for which FlyClockbase was developed for: a broader understanding of circadian clocks, often intentionally ignoring the details of many special cases. Thus, even if we had perfect knowledge of all potential sources of variability above, we might still want to exclude outliers using a systematic approach such as the one we employed here (Carling 2000).

#### Hypotheses on causes for differences in variance

An obvious explanation for such differences in protein peak times between PER and TIM could be given by similar differences in the mRNAs required for producing these proteins. This short-sighted hypothesis is easily tested using FlyClockbase. It turns out to be demonstrably wrong. As shown in Table 6 and Figure 6, the increased variance of PER relative to TIM cannot be attributed to an overall greater variance of *per* mRNA, because the peak time of *per* mRNA has a significantly lower variance than *tim* mRNA. Therefore, we can rule out carry-over from similar patterns of variance in mRNA peaks.

##### Phosphorylation network size

Here we propose that the increased relative variance of PER can be explained by the larger number of post-translational modifications for PER (relative to those observed for TIM). Post-translational modifications such as phosphorylation play a critical role in in the clock (Weber *et al*. 2011; Risau-Gusman and Gleiser 2012). While the exact nature and mechanisms of these modifications have yet to be fully resolved, there is strong evidence that PER undergoes more post-translational modifications than TIM.

*TIM protein* is phosphorylated by SGG (Ko *et al*. 2010), which promotes nuclear accumulation of PER/TIM complexes (Martinek *et al*. 2001), perhaps by allowing interaction with the nuclear import protein IMPalpha1 (Jang *et al*. 2015s). SGG-dependent TIM phosphorylation has also been implicated in light-induced TIM degradation, likely in conjunction with CRY and JET (Rothenfluh *et al*. 2000a; Bae and Edery 2006; Koh *et al*. 2006; Peschel *et al*. 2009). TIM protein therefore undergoes approximately two to three post-translational modifications.

*PER protein*, however, could be subject to ten or more post-translational modifications. PER is initially phosphorylated by NEMO, which then promotes additional phosphorylation by DBT (Chiu *et al*. 2011; Yu *et al*. 2011). DBT phosphorylates PER multiple times and influences PER stability, nuclear translocation, and SLIMB-induced degradation (Baylies *et al*. 1992; Edery *et al*. 1994; Rothenfluh *et al*. 2000a; Martinek *et al*. 2001; Ko *et al*. 2002; Kim *et al*. 2007; Chiu *et al*. 2008; KivimÄe *et al*. 2008; Ko *et al*. 2010; Chiu *et al*. 2011; Mezan *et al*. 2013). PER is also phosphorylated by CK2A, which promotes nuclear import (Lin *et al*. 2002a; Lin *et al*. 2005; Meissner *et al*. 2008). PP1 and PP2A both work against these kinases to dephosphorylate and stabilize PER (Harms *et al*. 2004; Sathyanarayanan *et al*. 2004; Fang *et al*. 2007; Chiu *et al*. 2008; Garbe *et al*. 2012).

##### Mechanistic phosphorylation network stochasticity hypotheses

Post-translational modifications could be opportunities for increasing the variability of timing. This is especially true if a required molecular type only exists in low copy numbers per cell at some relevant stages of a circadian cycle. As described above, PER protein is at the center of a large network of potential phosphorylation patterns and proteins, which also include dephosphorylations. This network of post-translational modifications dwarfs those observed in TIM protein. A large number of different types of potential modifications will break a large population of PER molecules into much smaller sub-populations, thereby greatly increasing stochasticity. The heterogeneity of this network and the relevance of antagonistic forces (dephosphorylation delays degradation) increase the potential for stochasticity and complicate predictions without detailed stochastic simulations. In comparison, few rather large subpopulations for TIM probably result in copy numbers that are high enough to substantially reduce stochasticity.

*Previous simulations* have highlighted the possibility of additional variability in the time required for growing to a defined level, when amplification starts from smaller amounts. For example, biochemical systems like signal-transduction cascades that amplify very low molecular counts can easily generate differences in variance for times to reach a peak (Loewe *et al*. 2009a; Loewe *et al*. 2009b; Akman *et al*. 2010; Ehlert and Loewe 2014). The inherent stochasticity of circadian clocks might explain the observed variability via various mechanisms. Potential explanations could include systematic differences in the distributions of the low molecular counts at the start of the respective amplifications. If this does not cause all observed differences in variability of PER or TIM peak timing, differences could be further amplified by the nature of the different reaction networks that generate the peaks of PER or TIM.

##### Future simulations

While beyond the scope of this present study, we think that such mechanistic phosphorylation network stochasticity hypotheses are worth exploring in reasonably realistic stochastic simulation models.

#### Expanding hypotheses on CURLED

The inversion of variance differences seen when comparing mRNAs and proteins of PER and RIM suggests that the variability discussed above is probably governed by the post-translational processes described above. However, it is less clear how these processes might explain the significant differences in the variance of valley timing for *per* and *tim* mRNA.

Circadian mRNA degradation might be influenced by CURLED (CU), which is known to affect circadian rhythms. Although *curled* mutants have been known for decades, CU was only recently identified as dNOCTURNIN (NOC), the *D. melanogaster* homolog of the mammalian NOCTURNIN (GrÖnke *et al*. 2009). NOC has been shown to associate with the CCR4-NOT complex, which promotes deadenylation (and subsequent degradation) of mRNA (Temme *et al*. 2010). While NOC is thought to influence circadian gene control, specific NOC targets have yet to be identified (Godwin *et al*. 2013). The gene *noc* produces three transcripts (*nocturnin-RD, nocturnin-RC, and nocturnin-RE*), and NOCTURNIN-RD is rhythmically expressed in DN3s (Nagoshi *et al*. 2010), a subset of dorsal neurons which are part of the circadian circuit and contribute to evening activity (Stoleru *et al*. 2004). NOCTURNIN-RD knockdown mutants have abnormal responses to constant light exposure, suggesting that NOCTURNIN may play a role in circadian light responses (Nagoshi *et al*. 2010). Green *et al*. (Green *et al*. 2007) also noted changes in gene expression in response to a high-fat diet for mutant *Noc*^-/-^ mice, which could implicate NOC in circadian metabolic control. It would be premature to postulate an interaction between NOC and *per* or *tim* mRNA. Instead, we suggest here that NOC and other circadian proteins that influence mRNA degradation might be a fruitful area of investigation, particularly given the connection between NOC, light response, and metabolism.

### Hypothesis: peaks from PCR methods are more variable

The *Attributes* collected for time series in FlyClockbase can be used to compare different groups of time series for a given clock component based on biological, methodological, or other factors. These comparisons can suggest sources of variability that affect future experiments and the interpretation of simulations. To illustrate this possibility, we compared different measurement methods for observing peaks and restricted our analysis to *per* mRNA time series, which produced 88 peaks, the largest number we could extract from FlyClockbase.

#### Methods background

Five methods were used to collect at least three independent time series: (i) Microarrays, (ii) RNA-seq and nascent-seq, (iii) PCR, (iv) RNase protection assays (RPAs), and (v) Northern Blots. Each method provides advantages and disadvantages. Historically, Northern Blots were the first of the five methods to be developed. Although they can provide information about transcript size (Sharkey *et al*. 2004), they have low sensitivity (Vandenbroucke *et al*. 2001) and can only be used to analyze one gene at a time (Fryer *et al*. 2002). RPAs were developed after Northern Blots. They can analyze multiple transcripts (Sharkey *et al*. 2004) and can be used to determine absolute RNA levels (Vandenbroucke *et al*. 2001). However, they might have low reproducibility (Qu and Boutjdir 2007), and RPA is time-intensive, typically requiring about four days (Streit *et al*. 2009). All three of the newer techniques (PCR, microarrays, and RNA-seq) are high-throughput methods (Bustin 2002; Sharkey *et al*. 2004; Mortazavi *et al*. 2008). PCR, RNA-seq and nascent-seq methods do not require previous knowledge of specific genes or sequences to be identified (Fryer *et al*. 2002; Mortazavi *et al*. 2008). While automation can be difficult for PCR (Fryer *et al*. 2002), microarrays are typically automated. Although RNA-seq and nascent-seq are the most newly developed methods, technical variability may be a concern, particularly when using a low number of read counts (Bullard *et al*. 2010; Mcintyre *et al*. 2011).

#### Sensitivity vs reproducibility

When comparing measurement methods, it is important to consider both the sensitivity and reproducibility of each method; methods that have high sensitivity may or may not have high reproducibility. Northern Blots generally have low sensitivity (Gilliland *et al*. 1990; Wang and Brown 1999; Malinen *et al*. 2003). RPA is more sensitive than Northern Blots, but sensitivity remains a challenge, especially when using low amounts of mRNA (Wang and Brown 1999; Vandenbroucke *et al*. 2001). PCR is considerably more sensitive than either RPA or Northern Blots (Wang and Brown 1999; Malinen *et al*. 2003). RNA-seq also shows good sensitivity (Mortazavi *et al*. 2008), but RNA-seq sensitivity depends on normalization techniques (Bullard *et al*. 2010). There is mixed evidence for microarray sensitivity. For example, some researchers found that Northern Blots were slightly more sensitive than microarrays for 14 of 29 assayed genes (Taniguchi *et al*. 2001). Six of the remaining genes, however, were detected by microarrays and not by Northern Blots, suggesting microarrays were more sensitive to these genes (Taniguchi *et al*. 2001). Older microarrays might only have been able to detect changes reliably if they were at least two-fold (Fryer *et al*. 2002), but some newer Microarrays have become at least as sensitive as RNA-seq (Willenbrock *et al*. 2009).

#### Poor reproducibility may be masked by biological variability

In addition to differences in sensitivity, measurement methods vary in reproducibility, and the procedures used in each measurement method point to different potential sources of variability. For example, RNA-seq requires a small portion of the sample RNA to be used to construct a library, and PCR used to create this library can introduce variability through amplification bias (Aird *et al*. 2011). However, biological variability typically outweighs methodological variability for RNA-seq (Bullard *et al*. 2010). A detailed review of the reproducibility of RNA-seq is given elsewhere (Seqc/maqc-iii Consortium 2014). Reports of reproducibility for microarray studies have been mixed. The 2005 Toxicogenomics Research Consortium raised concerns of variability between laboratories and between platforms (Bammler *et al*. 2005), and cross-platform reproducibility issues were echoed elsewhere too (Canales *et al*. 2006). However, simultaneously a large study by the MAQC Consortium found microarray experiments to be reproducible both across platforms and across laboratories (Shi *et al*. 2006). We expected time series measured with RNA-seq and microarrays to show the least variability, but there was not sufficient data to test this hypothesis.

#### Experimental causes for PCR variability

A number of different factors have been shown to influence the variability of PCR experiments (Bustin 2002). For example, different samples can have different amplification efficiencies (Vanguilder *et al*. 2008), and, as noted above, the percentage of GC bases can introduce amplification bias (Aird *et al*. 2011; Orpana *et al*. 2012). Others noted that, although PCR is often thought to be a “gold standard”, extensive tests showed that calibration and selecting appropriate primers and probes can be challenging (Vanguilder *et al*. 2008; Seqc/maqc-iii Consortium 2014). A low quantity of starting material can also influence PCR variability (Vandenbroucke *et al*. 2001; Bustin and Nolan 2004).

#### PCR and Northern Blot accuracy

Despite these challenges, PCR was developed more recently than RPA and Northern Blots, and the latter have largely have largely fallen out of favor, at least partially due to the greater degree of sensitivity afforded by PCR. Northern Blots are also less accurate than PCR (Vanguilder *et al*. 2008) and are considered to have low reproducibility (Qu and Boutjdir 2007). We therefore expected to see greater variability in time series measured with RPA and Northern Blot and less variability in those measured with PCR. However, our analysis revealed that peak values for time series collected with PCR were significantly more variable than those from time series measured with RPA or Northern Blot.

#### Differences between qPCR and RT-PCT?

We subdivided time series measured with PCR into those measured with real-time PCR (“RT-PCR”) and those not measured with RT-PCR (“qPCR”). Neither type of PCR was more variable than the other. Variability did not significantly differ between qPCR and Northern Blot or between qPCR and RPAs, but careful inspection of the bootstrap distributions produced by *comvar2* suggests that this could be merely an issue of statistical power (qPCR has fewer samples than RT-PCR). Accordingly, RT-PCR was significantly more variable than Northern Blot and RPA. Finding significantly higher variability for RT-PCR was also surprising, given that real-time PCR was developed more recently than qPCR, and RT-PCR is considered to be the standard for PCR, as it decreases experimental error by requiring less data processing than qPCR (Vanguilder *et al*. 2008).

#### Analysis of measurement protocol references using FlyClockbase

We attempted to explain our observed increase of variability for time series traits observed with PCR by examining the experimental protocol references cited by each type of mRNA observation method. It is common for the methods section of studies in FlyClockbase to reference the experimental protocol of previously published studies. We hypothesized that differences in these protocol references could explain the increased variability of time series measured with PCR. The time series identifier structure, easy access to references, content of respective studies, and the overall structure of FlyClockbase were instrumental for collecting and organizing information on experimental protocol references, even though this data was not originally recorded. The information on method references for *per* mRNA time series is shown in Figure 9. Table 8 provides overview counts that translate into statistical power when analyzing each method, either based on counts of time series (bold numbers in Table 8), or based on counts of studies (non-bold numbers in Table 8).

#### Shared references are less frequent for PCR studies

Of the three methods we analyzed, some methods generally cited more references than others. While all 45 time series using RPA cited at least one method reference, such references were cited by only about 56% (5/9) of the studies using Northern Blots and about 72% (18/25) of PCR based time series. Some of the method references were cited once, while others were cited more frequently. We defined “shared method references” as references cited by two or more studies which use the same measurement method. Studies using RPA had 93% (42/45) shared method references. 40% (2/5) of studies that used Northern Blots had at least one shared method reference, while only 17% (3/18) of PCR studies had a shared method reference. We hypothesized that decreased variability for time series measured with RPAs and Northern Blots could be attributed to increased number of method references and shared method references. However, our statistical tests found no significant differences between time series with or without method references or shared method references. We therefore suggest that increased variability in time series measured with PCR is not caused by a lack of properly documenting or a lack of using shared protocols but rather stems from actual differences in variability based on measurement method.

### Explanation: large fluctuations from PCR stochasticity

Here we provide a mechanistic explanation for the increased variance of *per* mRNA peak times as measured by PCR (here brief for RT-PCR and qPCR) in comparison to non-PCR methods (here brief for Northern Blot and RPAs). Briefly, repeated replication required by PCR starts with substantial stochasticity at very low copy numbers before reaching its deterministic exponential growth phase. Non-PCR methods for observing *per* mRNA do not require replication and thus have less potential for variability. Thus, non-PCR methods cannot distort peak timings as much as PCR can.

#### Exponential growth causes for PCR variability

As indicated above, many different factors can influence PCR variability, including amplification bias, calibration, primers, probe selection, operator experience, and importantly the overall quantity of starting material. It is easy to compare at length potential reasons for variability in PCR and in other methods. We suggest the following simplified analysis that relies on the fundamentally different behavior of timing errors in exponential growth vs linear growth. Such errors generate the larger variance of PCR-measured peak timings. Our explanation requires three basic assumptions:

i. Real-world individuals cannot be divided without destroying them.
ii. Replication without resource limits inevitably leads to exponential growth.
iii. Timing of later events in an exponentially growing system are easily affected by an earlier or later start of growth.

These assumptions define implicitly a theoretical model of exponential growth that can explain the increased variability mechanistically. We then present evidence suggesting the larger variability of PCR peak times should not come as a surprise.

#### Great sensitivity and poor reproducibility are linked

The goal of PCR is to amplify rare nucleic acids by repeatedly replicating in well-defined rounds. During later stages of growth many molecules are replicated simultaneously. Therefore, any individual replication event will not significantly impact the overall population, as the stochasticity of many individuals cancels out increasingly. Ordinary differential equations work well for such large populations, because their constant violations of basic assumption (i) are negligible here. The contrast of this precisely predictable scenario could not be bigger when compared to the very early stages of a PCR reaction designed to start with low copy numbers for maximal sensitivity. Here basic assumption (i) severely constrains potentially parallel actions, because single molecules cannot be broken up without affecting their functionality and are limited in what they can do simultaneously. This limitation inevitably creates stochastic waiting times that lead to larger or smaller growth delays that generate the initial timing differences at the root of cascades of delay that propagate throughout the exponential growth phase due to assumption (iii).

Amplifying single molecules is a hallmark of PCR’s exceptional sensitivity. It also causes PCR’s reproducibility problems for the reasons just explained, making it extremely sensitive to early rare template numbers. Here timing differences in polymerase access and replication speed can quickly snowball into faster or slower growth, and thereby, lower or higher amounts inferred for the original molecules investigated. These problems are highly relevant for all forms of quantitative PCR, which are designed to operate completely under exponential growth for better quantitation.

Circadian clock cycles with small variations of initial amounts inside of cells, stochastic timing differences, variations in extracted volume, and other factors can easily conspire to modify final amounts inferred by PCR (if stopped before resources become scarce). PCR time series measurements rely on these final amounts to define the PCR end results used for inferring how much may have been present at the beginning. For this to work as a quantitative method, PCR has to be stopped in the middle of exponential growth, implying it will inevitably experience substantial noise from slight variations in the starting conditions under a broad range of circumstances.

#### Observations of the theory

The strong impact of stochastic timing differences in exponentially growing systems is easily demonstrated in stochastic simulations of a very simple exponentially growing population (Ehlert and Loewe 2014). There the initial amount is kept constant for all simulations, making timing differences the only source of stochasticity. The same principles are responsible for translating the stochasticity of low molecule counts at the input of sensitive signaling cascades into a reliably transmitted signal, albeit with variation in the waiting time until the signal is switched on completely (Loewe *et al*. 2009a; Loewe *et al*. 2009b). Thus it is not surprising if experimental measurements show that different researchers with varying PCR expertise can easily generate 100-fold differences in their inferred initial number of molecules at the start of a PCR (Bustin 2002). Such variability might stem from small changes introduced to factors that impact the exponential growth essential to PCR in subtle, but powerful ways; see (Bustin 2002; Bustin and Nolan 2004).

##### How this applies to amount peak timing observed by PCR

As shown by growth mechanism discussed above individual PCR reactions bring individual challenges, which complicates observations of time series. The main reason is that each time point measured by PCR will require an independent PCR reaction probably starting with a low molecular count as obtained from sacrificing an independently running circadian clock. Thus, observing mRNA clock oscillations by some quantitative PCR method will inextricably intertwine two processes of variation that inevitably interfere with each other’s observation in these two ways:

i. Oscillations of the clock itself may exhibit substantial stochasticity depending on molecular amounts involved (Akman *et al*. 2010). This implies that the peak itself as measured by PCR may vary, even if PCR were perfectly precise.
ii. Low initial molecule counts of the templates that start any quantitative PCR reaction cause substantial inherent stochasticity that can substantially affect the final amount of PCR products measured. If this happens, researcher are likely to infer corresponding deterministic changes in the initial molecular counts (Bustin 2002).

Since every single time series observed by any quantifying PCR is inevitably impacted by both, a substantial amount of random noise is added to each independently observed time point. As a result, several time points might falsely appear to be peaks (or the converse).

#### Summary

A low quantity of starting material can influence PCR variability to a very large degree (Vandenbroucke *et al*. 2001; Bustin and Nolan 2004). Given the systematically larger potential for measurement noise in PCR methods caused by the low initial molecule count induced stochasticity, it might even be surprising that PCR is not noisier compared to methods like RPA or Northern Blots that do not require exponential growth.

### Model curation for integrating molecular systems biology data

The process of model curation inherently works towards integrating all data that is relevant and available for a given model of interest. Models may be broadly defined as systems, parts, processes or questions that are being represented from certain perspectives to efficiently find particular types of answers deemed to be interesting. We will next briefly discuss, how this view of model curation can facilitate the integration of knowledge-fragments from molecular systems biology in order to enable the emergence of expertise as represented by well-curated systems biology models (e.g. of circadian clocks) or corresponding relevant sets of real-world observations (e.g. of time series in FlyClockbase). We will first look at more specific followed by more general levels of abstraction before discussing other fundamental aspects of model curation.

#### Related concrete solutions

At a more specific level, there is no shortage of standards, formalisms, approaches, tools and other systems for supporting the application of more abstract categories (like ontologies or models) to concrete problem areas of interest. Examples include the Systems Biology Markup Language for constructing simulation models (Hucka *et al*. 2003; Krause *et al*. 2010), Systems Biology Graphical Notation for visually representing molecular reaction models (Le Novere *et al*. 2009; Moodie *et al*. 2011), UMLS and SNOMED for defining and using medical reference terms with different approaches to synonyms (Major *et al*. 1978; Merrill 2009), and specific ontologies for listing existing entities such as ‘all genes’ in an area of interest (Jonquet *et al*. 2011; Musen *et al*. 2012).

At the most specific level are concrete collections of actual models implemented in one of the formalisms described above. For example, BioBase, which collects and curates published SBML models (Le Novere *et al*. 2006; Chelliah *et al*. 2015). This is closer to the level of FlyClockbase, which collects and curates published time series within its scope. The substantial number of different formalisms for describing models can be very confusing. To get a clearer understanding of essential, non-redundant aspects of model construction it can be useful to consider a more abstract perspective.

#### Related abstract frameworks

Several abstract perspectives exist. An ontology is a list of potentially existing things. A taxonomy is a list of potentially existing species. A type system is a classification of potentially existing types and how they could be used to compose new types. Similarly, a model is a specification of potentially existing elements in the world of the model. At the most general level, ontologies and taxonomies are fundamentally related (Arp *et al*. 2015). The same holds for type systems, the semantic web, and models in general. All these could be described as ‘worlds’, as each of these is like a small description of its world. Unless otherwise specified, they buy into the Closed World Assumption, which implies that nothing else exists or matters except for the details explicitly specified. At this abstract level, worlds are all equivalent to systems that encapsulate detailed statements about the conditional existence of sub-systems, items or events that may be nested or composed from defined structures, capabilities, and/or other properties. Such abstractions enable the detection of isomorphisms that can facilitate the transfer of equivalent solutions across problem domains and hence cut development costs by building on results obtained elsewhere. For example, different elements or types can be grouped into a set in the contexts of taxonomy, ontology, type system, or model construction. They each may use different key words to describe this concept, but its core meaning, i.e. semantics, stays the same. Each of these worlds comes with its own semantic formalism.

It can be challenging to navigate these abstract semantic formalisms for representing the meaning of statements (van Renssen 2005). This resulted in the paradoxical (non-expert) use of ‘semantics’ as synonym for ‘meaningless’ in common language. A semantic model that is genuinely useful to its writer but incomprehensible to its reader is not useful to that reader and thus appears ‘meaningless, resulting in semantic irreproducibility (Loewe 2016). The resulting communication failure is a substantial problem for modeling, programming, giving names and using names (Loewe 2016). FlyClockbase has not been spared; we encountered a broad range of semantic problems caused by naming, from trivial spelling errors (with non-trivial consequences in database searches) to profound research questions about the nature of certain molecules (see discussion of CURLED above). Related questions of naming and nomenclature are of critical importance for biomedical research; correspondingly tools that efficiently map local nomenclature to standard nomenclature have been identified as critically important (NIH *et al*. 2012), and would have made development of FlyClockbase substantially faster (e.g. by helping to manage changes in local names).

#### Baseline: conceptual unity of reality despite diversity of experimental methods

Science builds on the physical unity of reality that is observed by different persons using different methods. This principle is usually so compelling that it is unconsciously assumed. It allows scientists to confidently assume the same conceptual unity for aspects of reality that are challenging to study because they may present a different view when investigated by different approaches. This principle of conceptual unity is the foundation of model curation. For example, let *Q* be the amount of a given protein type in a single cell specified by place and time. Then *Q* itself does not depend on the various methods subsequently used to measure Q. If results from different methods of observation contradict each other, we can confidently search for errors. The confidence is rooted in the conceptual unity of our world, or any cell, or Q.

#### Contradictory biological information

While developing FlyCockbase we repeatedly encountered situations where there were contradictions between different observations that appeared to be of equally high credibility. On some occasions, even substantial efforts on our part to check each credible source of confusion we could think of, did not identify any credible information on what could have gone wrong. Such situations are profoundly confusing and cost substantial amounts of time. Handling such difficult situations defines much of the quality of a *VBIR* and its underlying logic formalism (see also Discussion below and in Supplementary Text).

##### Debugging time limits

We found it important to limit the time we used for attempts to resolve such problems. In this we aimed to be generous yet responsible with our resources. We also started to search for more formal ways of signaling among curators when a particular set of problematic time series already had been investigated sufficiently. The implications of this question for the reproducibility debate are unclear. Should a seemingly solid experimental study be declared ‘irreproducible’ because an apparently rushed, ill-conceived experiment failed to reproduce results? Probably not. Should a seemingly rushed, ill-conceived original study be defended as a valid original observation, despite the apparent inability of thorough, time-consuming attempts to reproduce results? Probably neither. However, where is the line between these two rather extreme scenarios? It is not the task of biological model curators to assess the credibility of experiments by repeating them. Hence, they need other means of assessing the quality and relevance of reported observations for the model they curate. A more differentiated formalized way of communicating various perceived problems could greatly increase the efficiency of curation work by relieving curators of the implied unrealistic obligations to always get to the bottom of all inconsistencies or to invent a reliable taxonomy of resulting errors.

##### Communicating errors

Developing a formalism for communicating clearly how to handle errors efficiently is a complicated problem more closely related to compiler construction than to biological questions. It requires expertise in both areas. We started to search for efficient ways of how to best represent outcomes of error analyses for particular time series. We aimed to formalize such communications with the intent to enable a compiler to exclude certain types of errors from the results of time series searches. It became increasingly clear that binary choices like “error: yes/no” were inappropriately simplistic for many real-world uses of data in biology.

##### Types of problems with data

The discussion above demonstrates that a differentiated approach is necessary for appropriately representing biological data. Not all trustworthy biological expertise is documented by directly observed data and not all data that is available has the quality most researchers would ideally aspire to. Statistical inference and logical deduction from experimental observations are also valuable tools of biological discovery. However, they can only yield conclusions that are as strong as the observations that support them. It is therefore of utmost importance for reliable and reproducible research in biology to represent specific experimental observations and general biological data as they are, including all known limitations and unforeseeable circumstances, confounding variables, or event. From this perspective, almost all data is imperfect to some degree. Imperfection in an imperfect world is not a problem, as long as we are aware of the nature of the imperfections. The current reproducibility crisis reflects in part the complicated nature of reporting the essence of research results concisely, yet without ignoring their limitations or omitting potentially undermining details (Baker 2016). In our study, we have attempted to be as complete and open as possible, e.g. by conducting a human error analysis for the raw data of our most important conclusions and reporting multiple potential variations of our statistics (see Supplemental Statistical Analysis); this has both substantially increased the length of this report and the time to complete it. As can be seen in the overall structure of our R-script that computed our final analysis, such work often requires exploring various alternative analyses. These all initially appear to be equally valid ways of working around a given imperfection of the data. Substantial parts of calculating all useful tests can reasonably be delegated to a compiler for many frequently encountered scenarios – assuming there is a formal way of communicating the type of data imperfection to the compiler.

##### Using imperfect data

For the reasons above, imperfect biological data is extremely valuable. Hence, no need to throw out baby hypotheses with imperfect data bathwater. High-quality model curation considers what can reasonably be learned from an imperfect dataset by describing as many quantitative aspects as reasonable, reflecting ideas from the “New Statistics” (Cumming 2013; Cumming 2014). Often the cutting edge of research is defined by situations where not enough high-quality data is available for a final interpretation. In fact, the value of resources like FlyClockbase is precisely in their ability to synthesize the limits of what is known and highlight hypotheses that merit further experimentation. Ignoring imperfect data in this context would be inappropriate.

##### Imperfection spelled out

Hence, FlyClockbase will require better ways of representing confusing and uncertain real-world data at the current biological cutting edge of research. Data there can be aggregated, biased, contradictory, diffuse, exception-prone, false, generated, gap-ridden, hidden, imprecise, jumbled, limited, missing, modified, noisy, objectionable, problematic, questionable, redundant, scattered, swapped, tangential, transformed, uncertain, veiled, washed, wobbly, xeroxed, or otherwise imperfect. A hallmark of good biologists is their ability to intuitively navigate these difficulties appropriately in their study systems. The rise of big data has led to substantial experience in how to deal with imperfect data (Mccallum 2013).

##### Problem type repository

Developing *VBIRs* like FlyClockbase efficiently depends critically on the ability of biological model curators to describe these intuitions in ways that are formal enough, so that an automated solution can be developed eventually. Biological experiences with rates of identifying new species in an ecosystem where many of them exist (Grove and Stork 2000) indicate that eventually known species will be resampled. The same can be expected for the varied number of data problems that can be observed during the long-term development of *VBIR*s. *VBIR*s would greatly benefit from a central repository for the logic problems associated with imperfect data. Such a repository can substantially cut costs of identifying logic problems and would help in compiler construction, simply by documenting the extent of the problem. It is difficult, even for experienced biologists, to imagine many of the complications of real-world data in the absence of actual research interactions with real-world data. It is near impossible for complier constructers to do so without also being biologists who work with real-world data.

#### The role of logic

Communicating errors in clear ways fuels our interests in exploring logic formalisms beyond classic Boolean systems (see also below). Providing a full formal definition for a chosen logic formalism, alongside all appropriate proofs and consistency checks is clearly work best done by theoretical computer scientists with the corresponding formal training. For the reasons discussed above, we do not think that developing an appropriate logic can be reasonably delegated. As indicated above by the idea of problem type repository, we found that one of the biggest challenges was the need to realize the existence of a given problem. Typically, and trivially, biologists find it easier to specify uncertainties and inconsistencies in real-world observations based on their experience, while logicians find it easier to identify particular contradictions that result from an inappropriately defined logic formalism. Our experience suggests that their combined imagination and expertise will need to be complemented by a slow careful collaborative review of the detailed problems in a sufficiently complex real-world research scenario. To facilitate such collaboration, we have described elsewhere the Flipped Programming Language Design approach (Loewe 2016), which also inspired our discussion below of Figure 11.

These complex efforts to develop a sufficiently expressive logic for problems with biological observations contribute towards answering the next question that is in principle very simple.

#### Simple question: how many molecules of a given type exist at a given time in a given cell?

Modern biology has invested much effort into developing many diverse approaches for investigating intracellular quantities of interest. Such quantities often relate to the simple question of amounts in one of the myriads of forms in which it is posed in biology today. Generating well-defined, credible answers that properly quantify all relevant uncertainties would go a long way towards providing the data required for algorithms aiming to solve the inverse problem (see Models Section). The answers to this problem quantify the uncertainty of parameters, which are needed for simulating models of molecular systems in cells. Such simulations can be seen as devices for extending biologists’ thinking capabilities and enable investigating new areas of biology (see comments on evolutionary systems biology (Loewe 2016)). Unfortunately, it is extremely challenging to answer the conceptually simple question above for real molecules in real cells with reasonable quantifications of uncertainty.

##### Observations in FlyClockbase

Many of the practical challenges of determining such amounts of molecules have been constant companions of our work with FlyClockbase. For example, consider the differences in techniques used to measure amounts of mRNAs or proteins produced by genes such as *per* or *tim*, (see Figure 10 or *TimeSeries AttributeTable* in FlyClockbase for details on methods). While each technique is limited in unique ways, a given quantity of interest can usually be measured in several ways that vary in trade-offs between precision, cost, and other method parameters. As a result, interesting quantities that can be measured in cells have often been measured by dozens of methods, each of which may be implemented by different independent experimental protocols and belong to one of several applicable broader methodological approaches. Each of these may provide different answers to the following practical questions that are highly relevant for model curation. Are amounts of molecules in a single relevant core clock cell of *D. melanogaster* …

- … *absolute counts* (our preferred ideal) or *relative* (usually reported)?
- … from a *single cell* (may be used to infer molecular noise) or from *averaging* over a population of cells (or some other aggregation difficult to disentangle)?
- … *complete and direct raw observations* (enabling independent statistical analyses) or *summaries* of “typical plots” of “the most relevant data” (that can introduce uncontrollable ascertainment biases as observed in other areas, e.g. (Amos *et al*. 2003; Clark *et al*. 2005; Foll *et al*. 2008; Lachance and Tishkoff 2013; Minikel *et al*. 2014))?
- … *appropriately annotated* with all key details for maximizing long-term use in diverse meta-analyses (rare; authors have little guidance on what to report) or *missing annotations* for key details known to exist even if unreported (e.g. fly age, sex), or not reasonably knowable (e.g. fly clocks were disturbed by unexpected and unreported drastic changes in temperature)?
- … *reasonable approximations of reliable results* (as would be expected from diligent analyses of larger and higher-quality data sets if reproduced in the same system or *irreproducible* (recent observations may give reason to pause (Ioannidis *et al*. 2009a; Salanti and Ioannidis 2009; Mobley *et al*. 2013; Freedman *et al*. 2015a; Freedman *et al*. 2015b; Halsey *et al*. 2015))?

The unity of reality implies that similar representational approaches can contribute towards rigorously assessing reproducibility and towards curating heterogeneous and imperfect datasets into an internally consistent *VBIR*: both efforts would benefit from explicitly stating all uncertainties and other problems associated with a given set of observations.

#### Curation efforts for circadian clock research

FlyClockbase is unique in its scope, datasets covered and many other aspects – as far as we can tell. In particular, we are not aware of other circadian clock time series resources or meta-analyses that bring similar numbers of replicate time-series or studies together in order to answer questions about the differences in variances of peak times between different components of the core circadian clock of *D. melanogaster*. However, FlyClockbase is not the first biological information resource sharing observations about circadian oscillations in gene expression. We will next discuss some examples of related efforts; a review of additional bioinformatics resources relevant for clock research can be found elsewhere (Lopes *et al*. 2013; Li *et al*. 2017).

- *CircaDB* (http://circadb.hogeneschlab.org) is a publicly accessible web database storing time series observations that record how gene expression changes in various mammalian tissues throughout the day (Pizarro *et al*. 2013). It has been used for documenting the large extent to which gene expression in mice follows circadian patterns – with interesting implications for drugs that target the products of rhythmically expressed genes and that might benefit from timed dosage (Zhang *et al*. 2014a). While many genes in the mouse clock are homologous to fly clock genes, there were no observations of non-mammalian gene oscillations in *CircaDB* at the end of 2016. A strength of CircaDB is the availability of detailed tissue specific data from mice.
- *CGDB*, the Circadian Gene DataBase (http://cgdb.biocuckoo.org) version 1.0 (as of 2017-01-14) contains information (i) on 1,382 instances where gene expression followed circadian rhythms as observed by techniques like RT-PCR, Northern Blots or *in situ* hybridization; (ii) on 26,582 observations of gene expression found in transcriptome profiling studies to follow circadian rhythms; and (iii) on 44,836 potentially oscillating genes as identified in a search for orthologs of oscillating genes (Li *et al*. 2017). A strength of CGDB is its broad coverage of 148 different animals, plants, or fungi. Of the 27,964 genes with experimental evidence of oscillatory gene expression, 3166 have been observed in *D. melanogaster*. Of these, 14 observations cover all isoforms of *per* and *tim*, but only 5 of these were recorded in LD. The peak and valley times reported in CGDB do not contradict those reported by us here; however, the reported sample size does not have the statistical power to suggest new hypotheses on potential clock mechanisms.
- *Deep (machine) learning* approaches were investigated for their capacity to predict time and to distinguish rhythmic from arrhythmic time series (Agostinelli *et al*. 2016). To this end BioCycle_Real_ was curated from 36 gene expression datasets, including 32 from CircadiOmics (http://circadiomics.igb.uci.edu/) (Patel *et al*. 2012). Except for one from the plant *Arabidopsis thaliana*, all datasets came from mice (Agostinelli *et al*. 2016).
- *SCNseq* (http://wgpembroke.com/shiny/SCNseq/) provides access to temporal transcriptomics of circadian clock controlling cells in the suprachiasmatic nucleus of the mouse brain at unprecedented precision (Pembroke *et al*. 2015).
- *Bioclock* (http://www3.nd.edu/~bioclock/) is a repository of circadian transcriptional profiling data from *Anopheles gambiae* and *Aedes aegypti*, mosquitoes acting as vectors for malaria and yellow fever, respectively (Rund *et al*. 2011; Rund *et al*. 2013; Leming *et al*. 2014).
- *BioDare* (http://biodare.ed.ac.uk/; http://millar.bio.ed.ac.uk/data.htm) is an online service for data-sharing and analysis of circadian time series observations. It’s 10 datasets from *A. thaliana* were used for comparing period estimation methods and other clock research (Zielinski *et al*. 2014).
- dbCRY (http://www.dbcryptochrome.org/) facilitates comparative genomics of crypotchromes, the light-sensing proteins in clocks (Kim *et al*. 2014); see Figure 1.
- *Diurnal 2.0* (http://diurnal.mocklerlab.org/) provides access to observations of circadian genome-wide gene expression patterns observed in several common model plants (Mockler *et al*. 2007).
- *EUCLIS* (http://www.bioinfo.mpg.de/euclis/) is the ‘EU Clock Information System’. It adapted an advanced database architecture from another systems biology project for circadian clock researchers in order to combine modules for experimental data, clock models, and a related digital library (Batista *et al*. 2007; Lopes *et al*. 2013).

*Individual meta-analyses* occasionally integrate different datasets in an effort to increase the statistical power and reliability of conclusions. For example, combining and curating data from five independent microarray studies in *D. melanogaster* confirmed the rhythmical expression of 81 transcripts while also identifying 133 new cycling transcripts (Keegan *et al*. 2007). To arrive at their conclusions, KEEGAN *et al*. had to obtain data directly from the authors of the microarray studies they analyzed as not all necessary data was available online (Claridge-Chang *et al*. 2001; Mcdonald and Rosbash 2001; Ceriani *et al*. 2002; Lin *et al*. 2002b; Ueda *et al*. 2002). In turn, the same happened with their results: “All data used to produce this report are available upon request. Files that contain the individually formatted results from each of the original reports were too numerous and large to be included with this manuscript …” (Keegan *et al*. 2007). Some (non-meta-analysis) studies that generate substantial amounts of new data put in the substantial additional work necessary for making material available online (e.g. see http://biorhythm.rockefeller.edu/ (Claridge-Chang *et al*. 2001)). Merely *storing* complex data in one or more file archives online is usually easy, but organizing and documenting complex datasets for use by independent researchers is not. This requires semantic reproducibility, which can quickly become prohibitively complex (Loewe 2016) if no existing conventions are shared with users. Projects above have used database technology and/or web interfaces as shared conventions facilitating communication; as argued elsewhere in our study, this is neither ideal for all biologists nor for all work in biology. These problems are less acute for studies that can fall back on using public repositories with an appropriate data format. For example, a functional analysis study of fly genes expressed in response to the light-induced resetting of the circadian clock (Adewoye *et al*. 2015) stored most data at the NCBI-maintained Gene Expression Omnibus database (https://www.ncbi.nlm.nih.gov/geo/query/acc.cgi?acc=GSE39578); NCBI GEO offers semantics particularly well suited for describing typical microarray datasets (Barrett *et al*. 2005; Barrett *et al*. 2011; Barrett *et al*. 2013; Clough and Barrett 2016).

##### Generic model organism repositories

Resources like the individual-study repositories above, meta-analyses, and circadian research specific repositories are complemented by generic model organism resources such as FlyAtlas (http://flyatlas.org) for gene expression information (Chintapalli *et al*. 2007; Chintapalli *et al*. 2013a; Chintapalli *et al*. 2013b; Robinson *et al*. 2013) and FlyBase (http://flybase.org/) for genomic and other information (Ashburner and Drysdale 1994; St PIERRE AND Mcquilton 2009; Gramates *et al*. 2017).

##### Why curate circadian data?

Without detracting from the important achievements that continue to be enabled by the resources specified above, some points are worth noting for further discussion. Scarcity is first. We attempted to be as inclusive as possible and added resources far beyond the focus of our study on core clock genes in flies; recent reviews (Lopes *et al*. 2013; Li *et al*. 2017) do not list much more. Yet rhythmic gene expression is pivotal for fitness, health and more. This is underlined by estimates of 10% or 43% of the expressed genome under rhythmic control (Boyle *et al*. 2017) or of all protein coding genes showing circadian expression patterns somewhere in the body (Zhang *et al*. 2014a), respectively. Given this importance of circadian biology, it seems surprising that not more circadian clock related repositories exist. One potential reason is that the tools for managing data are inadequate and discourage many biologists from getting involved. Secondly, the existing resources are scattered, very heterogeneous with respect to their data structures resulting in poor interoperability, and they are haphazardly reorganized (e.g. web addresses move, internal structures are modified).

##### Challenges

Circadian clock researchers have repeatedly stated over the years that the arrival of new data due to experimental advances creates new challenges for data management and data processing (Zielinski *et al*. 2014). These could be addressed by the development of a new infrastructure that standardizes data and software to reduce re-implementation efforts, improve documentation, and increase collaboration by sharing data (Batista *et al*. 2007; Zielinski *et al*. 2014). This need for improved and simplified infrastructure exists for systems biology in general (Cassman 2005) and similar ideas have been echoed in the debate about reproducibility (Buck 2015). The underlying issues have not yet been solved on a broader scale (NIH *et al*. 2012; NIH 2015; NIH 2016; Wilkinson *et al*. 2016). A recent review of additional bioinformatics resources pointed out that tools cannot replace researchers, because it is “often necessary to conduct an evaluation of the results of a data mining effort to determine the degree of reliability” (Lopes *et al*. 2013). Indeed, experiences at UniProt show that “expert curation is by far the most reliable method to report gold-standard information and provide an up-to-date knowledgebase containing experimental information” (The UniProt Consortium 2017). We argue that much of the low-level work of ensuring reproducibility and adherence to formal standards could be handled reliably by a compiler that transparently executes well-defined recurring tasks (see Discussion below and in Supplemental Material).

#### Perspectives on biological model curation

Circadian clocks control rhythmic gene expression for a substantial and important fraction of the genome, approximating half of all genes in mice (Zhang *et al*. 2014a). It is thus difficult to isolate clocks from the rest of the organism they govern. An overall assessment of the impact of circadian clocks might thus require simulating whole cells or even whole organisms. This perspective raises several questions.

##### Will this scale to real cells?

The challenges of reproducibility for systems that are comparatively small are multiplied on much more complex systems, such as the molecular systems biology simulations of whole cells that have started recently at a larger scale (Karr *et al*. 2012; Karr *et al*. 2013; Lee *et al*. 2013; Purcell *et al*. 2013; Sanghvi *et al*. 2013; Karr *et al*. 2014; Karr *et al*. 2015a; Karr *et al*. 2015b). The ultimate aim of such studies is to understand in detail, how a real cell work and evolves over time (Lynch *et al*. 2014). However, the question is, whether our tools will be able to scale in such a way that errors can be kept down and our toolchain remains reliable. Hence, outstanding reproducibility of smaller models and datasets are a prerequisite for any further integration. We have chosen to focus on the simplest possible implementations when developing *VBIRs* to enable durability.

##### Will tool development overwhelm biological goals?

The essential requirement of tools that handle biological data more accurately and with more ease could continue to bind disproportional amount of resources through a lack of coordination (see eg. (Cassman 2005)). Usually only one excellent tool for a given task is needed, not several that are usually good enough but break down in some special cases, which then require completely new implementations. It is encouraging that the accuracy of computational tools in some areas converges towards that of high-quality experiments (e.g. (Lejaeghere *et al*. 2016)); likewise, the development of more precise higher-level abstractions simplifies much of the lower-level programming (e.g. (Arp *et al*. 2015)). However, the need for new and more precise tools is vast, and only few biologists can program well enough to contribute. Thus, support from computer scientists and professional programmers will certainly be needed. However, without the extraordinarily close collaboration described here it will be extraordinarily difficult to develop tools that are efficient enough in real biological research in order to drive adoption. Only then will tool development start to contribute to the overall biological goals. In our analysis, the development of a VBIR compiler is a particularly efficient way of tool development (see Discussion below). The efficiencies from compiler development might help with raising the funding for VBIRs on a more permanent footing (Ember *et al*. 2013). Simultaneously, experimental methods, their limits, and associated errors and biases will require more rigorous analyses in order to contribute towards a more accurate description of the precision associated with the actual observations (e.g. for sequencing errors see (Roberts *et al*. 2013; Robert and Watson 2015), for n-fold gene expression see (Canales *et al*. 2006; Canales 2016), for PCR see (Bustin 2002; Bustin and Nolan 2004; Vanguilder *et al*. 2008), for tests of a parameter estimation method, e.g. see (Daigle *et al*. 2012); many more analyses for other methods are needed).

##### Will biology try to advance too fast for its own good?

The 1970’s saw the rise of systems theory in ecology, albeit arguably too early (Wolkenhauer 2001). Now systems biology has in principle a computational method at its disposal for every single step along the causality chain from genotype and environment to phenotype and fitness (Loewe 2009; Loewe 2012; Loewe 2016). However, what does not exist at the moment is a rigorous and integrated problem management for the full causality chain. Clearly, more uncertain output at more causal calculations will combine with additional uncertainties at more consequential calculations. If this accumulation of uncertainties occurs on the long causal chain from genotype to phenotype, then it is presently not clear, which signal-to-noise ratio is to be expected. This question can only be resolved by an integral management of uncertainties similar to what we propose. Advancing simulations of whole cells or even organism too fast without allowing for appropriate precision to grow in method development and curation might cause rigorous scientists to lose patience, throw out the baby with the bathwater, and thereby cause unnecessary setbacks.

##### Balance

It is neither possible nor necessary to manage either of the extremes above beyond being aware of them, to avoid falling into either trap. The dynamic nature of biological research will then run its course. However, any foreseeable scenario will have a very large need for biological model curation, which will require many well-equipped biologists, as high-quality model curation will always remain a human task. Similarly, in any credible scenario, biological model curators will greatly benefit from support by a well-equipped VBIRs compiler.

### Towards a compiler for advancing FlyClockbase and biology

Working with FlyClockbase has given us ample opportunities to observe first-hand many diverse problems that frequently complicate an otherwise efficient use of computers or formal simulation methods for advancing biology. Below we highlight a few key observations suggesting a greatly increased efficiency for integrating computers in the workflow of FlyClockbase and similar *VBIR*s could be enabled by the construction of a corresponding *VBIR* compiler. This requires paying the moderately increased cost of constructing such a compiler only once. In return, the whole biological community could substantially cut the excessive costs of manually constructing or maintaining *VBIR*s. The substantial costs of *VBIR*s construction conflict with the growing need of compiling thousands of *VBIR*s that integrate in computable form the biological expertise necessary for engaging the grand challenges of our time. Leveraging abstractions developed in computer science for cutting through the complexities of data management with the help of an appropriately designed compiler could greatly reduce the costs of integrating biological expertise in order to address grand challenges more efficiently.

#### History

Similar thoughts about better computing for biological discovery have been recurring since the dawn of computing (Turing 1936), fueling many discussions of chances and challenges in diverse areas and applications, including the following examples: harness the precision of logic for biological discovery (Woodger *et al*. 1937), simulate genetic systems (Crosby 1973), open science (Bartling and Friesike 2014), improve reproducibility (Ioannidis 2005b; Donoho 2009; Huang and Gottardo 2013; Loewe and Keel 2014; Stodden *et al*. 2014; Freedman *et al*. 2015a; James *et al*. 2015; Stodden 2015; Barba 2016; Loewe 2016; Loewe *et al*. 2016), share data (Wilkinson *et al*. 2016), and do so efficiently at larger scales (NIH *et al*. 2012; NIH 2015; NIH 2016). To contribute to this debate that also affects FlyClockbase, the next sections distill the essence of selected key challenges we observed. We connect our observations to relevant research in other disciplines to reduce rediscovery where possible.

#### Unusual approaches to constructing an unusual compiler

We will conclude that many or most of the problems below could be solved efficiently by carefully constructing a corresponding compiler. Its specialty is to facilitate the implementation of best-practice solutions for constructing VBIRs and addressing the many challenges which biologists regularly face when they aim to use computers for advancing their research. Our approach goes far beyond superficial reassignments of responsibility; rather it proposes that broad classes of problems in biology could benefit from computational solutions if the latter are designed with enough time and care for those abstractions that matter for real-world biology. Figure 11 illustrates key aspects of the development process we used. It represents a very unique informal blend of two opposite extremes in software development. The extreme known as ‘agile methods’ advocates for quick iterations that implement tangible improvements. Its successes have made it popular, but it is not without a dark side that can stifle the development of innovations and strategies needed for solving more complex problems (Janes and Succi 2012; Annosi *et al*. 2016). The other extreme approach to software development is known as the ‘water-fall method’. It emphasizes thorough planning of various stages, and clearly separates developing a design from implementing it. As captured in Figure 11, data integration in FlyClockbase followed faster (internal) release cycles. Questions of fundamental importance with implications for formal or theoretical aspects of compiler design followed a much slower timeline. This allows us to focus our implementation resources on the most promising and efficient formalisms, and avoid the need for implementing potential solutions that appear attractive for some time, but are replaced by the need for further improvements. In such situations, overall speed of compiler development probably benefits from manual VBIR curation, since this allows the compiler design the time it needs to mature. Working in a research setting, as we did with FlyClockbase, creates additional challenges, simply from the unpredictability of research. Classifying potential bugs in compiler construction can be seen as a problem similar to the development of the taxonomy of beetles: both exist in exceedingly larger numbers, and continued random sampling eventually leads to re-encountering similar bugs. Constructing a compiler that can deal with biology’s uncertainty and complexity in a stable and reliable way requires a very extensive sampling of these potential bugs (i.e. logical program inconsistencies). We have developed the Flipped Programming Language Design Approach in order to address this problem using repeated rounds of rigorous review of proposed compiler designs from multiple usability and domain experts (Loewe 2016). Our work on FlyClockbase benefitted from this approach and also contributed to its development. It illustrated for us, how developing good abstractions can take a very long time, and how much finding them worth the effort.

**FIGURE 11:**
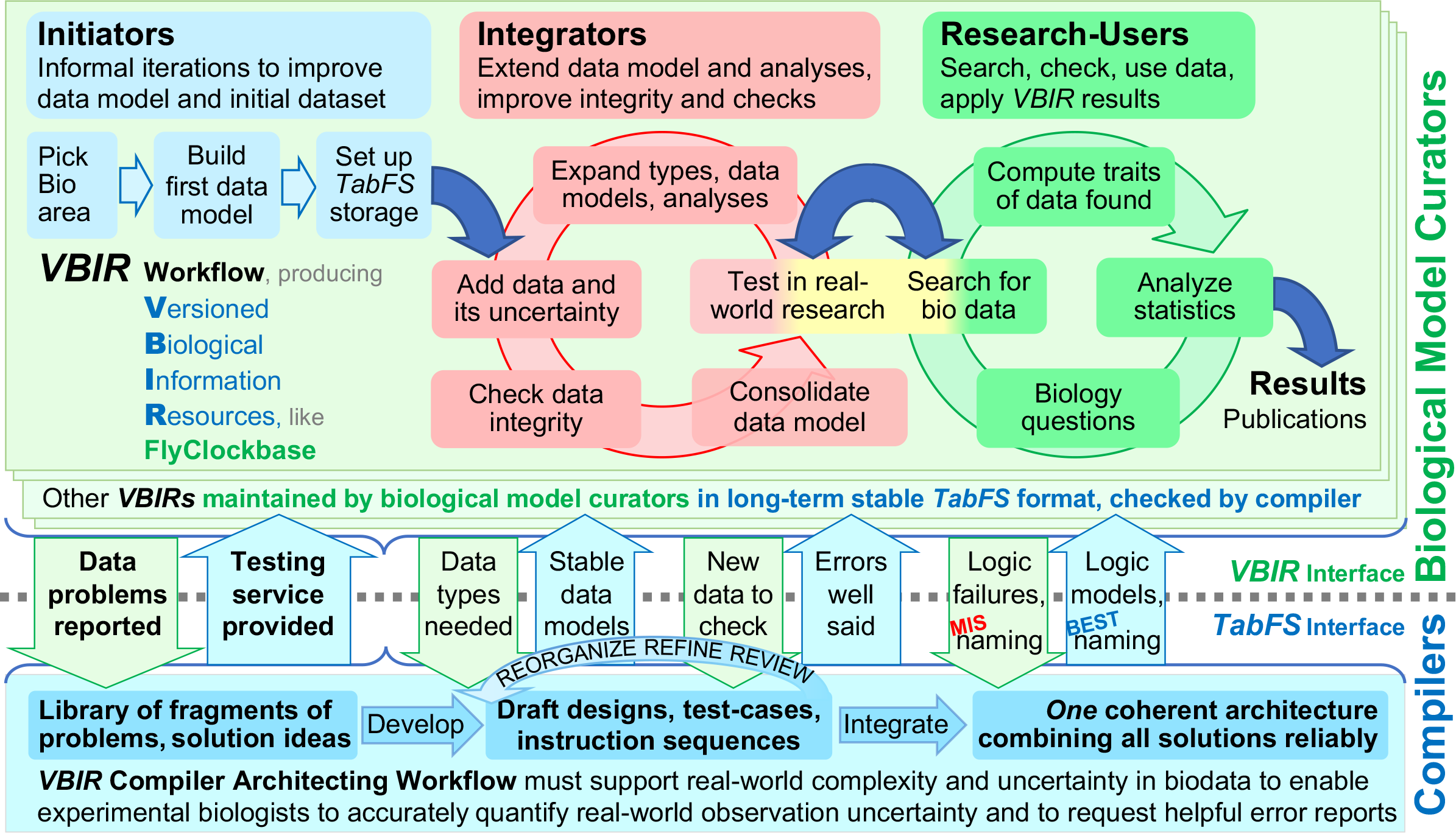
Biological model curator and compiler architect collaboration model for improving the integration of biological data into *VBIR*s while constructing the type system required by a *VBIR* compiler for handling the ubiquitous uncertainty of biological data more precisely. Here we show how biological model curators and compiler builders can collaborate by depicting important aspects of the trans-disciplinary interactions that led to the construction of FlyClockbase, a versioned biological information resource (*VBIR*). We have been using our compiler expertise to inform fundamental decisions about conceptual data structures and file formats in FlyClockbase and TabFS (designed simultaneously, all formal definitions are beyond the scope of this paper). ***Division of work***. We divided work among domains as indicated by the dotted line. Above, biological model curators worked with a focus on constructing and using FlyClockbase in order to understand the circadian clock of *Drosophila*. On the other side was the compiler architect focusing on TabFS data structures and the logic necessary for representing the biological data collected by the curators for FlyClockbase. ***Mutual benefits***. Designs were chosen such that implementing the TabFS compiler logic remained as simple as possible, while enabling formal tests of the integrity of *VBIR* data. In the present study, we developed many of the designs necessary for automation by manually pioneering important tasks. Doing so in a real-life research system of non-trivial complexity was important; this allowed us to explore many real-world details with substantial anticipated impacts on future use. The biological model curators focused on integrating and analyzing time series; this often enabled them to readily provide the compiler architect with detailed advance information about important use-cases and likely problems. The compiler architect could then evaluate potential design options long before implementation could start. This allowed for iterative reviews from several perspectives and with substantial time for analysis. It also resulted in a *VBIR* design and compiler logic that substantially complement each other and are thus prepared for automated testing of *VBIR* data integrity by following a number of simple rules. ***Collaborative exchanges*** of insights at the dotted *VBIR*-*TabFS* compiler interface are exemplified as broad arrows, indicating some important discussion topics (in terms closer to computer science for brevity; our actual discussions were informal and used vocabulary closer to biology). Our trans-disciplinary communication interface required both the high-level aspects of human collaboration and the low-level aspects of technical information exchange. The information flows shown here critically depend on biological model curators with a deep understanding of, and passion for, the details of state-of-the-art biology. Their expertise is essential for defining the boundaries of the system that is being modeled. While working to quantify its uncertainties properly in the *VBIR*, the biologists must be committed to bringing all potential problems of interpretation to the attention of the architect of a compiler, which is being designed to meet the formal needs of the *VBIR*. The architect must be able to select an appropriate mathematical logic formalism for representing the relevant biological problems in meaningful ways. The team needs excellent trans-disciplinary communication skills for efficiently describing, checking, and negotiating the uncounted decisions that collectively generate the systematic organization of an efficient *VBIR*. This requires a high sensitivity for the diffuse difficulties of accurately capturing computationally the uncertainties and contradictions in biological observations encountered while constructing a *VBIR*. Such biological problems require an appropriate logic formalism and thus need to be seen by the compiler architect, irrespective of perceived severity. ***Traps***. Avoiding do-it-yourself analyses of ‘simple problems’ by non-computational biologists is important for protecting against deceptive simplicity. It may not be possible to solve such problems on desirable timelines; still, compiler developers have better chances of spotting the dangerous costly bugs they can cause, which helps to identify solutions (that may already exist for other reasons). For example, quantifying uncertainty with the cutting-edge logic formalism known as ‘fuzzy plurivaluationism’ enables the representation of both semantic indeterminacy *and* various degrees of truth (Smith 2008); ‘BioBinaries’ extend Boolean logic in similar ways (see Supplemental Material). It often takes a trained eye to spot the need for a richer logic and it can be particularly hard to hunt logic errors that omit the possibility of some options (Panko 2016). Finding an appropriate logic formalism is pivotal, because it is impossible to compute without assumptions about logic. Computers always produce logical answers, no matter how flawed their logic might be when interpreted by humans in a real-world context. Such problems are most efficiently addressed at the compiler level by enforcing an ‘appropriate’ type system. Naturally, related discussions revolve around how to define ‘appropriate common sense’. ***Progress in defining type systems*** used for representing a biological system must be driven by those biologists who study the respective biology; to help them capture type defining ideas, we use the TabFS storage system of nested folders and tables which biologists can easily navigate using common operating systems and spreadsheet tools (see main text). ***Cost of constructing VBIRs without an appropriate compiler***. In contrast, without the appropriate logic and the equivalent of TabFS built into a corresponding *VBIR* compiler, it will still be necessary for each *VBIR* to construct, find or develop corresponding tools if quality is important. Unfortunately, economies of scale will be missing in this case, resulting in greatly reduced quality at greatly increased cost. Thus, developing *VBIRs* on a shoe-string budget will either limit *VBIRs* to comparatively simple datasets (thereby excluding much of biology) or is likely to trigger many rushed decisions about underpinning logic. We found that correspondingly simplistic type systems easily frustrate non-computing biologists by expecting them to coerce observed data and its uncertainties into an inappropriate logic (demanding a precision not provided by the data). This results in biases that are often near-impossible to detect, quantify, or exclude (e.g. by testing whether coercion indeed occurred). Thus, representing biological data in programming languages that do not readily support an appropriate logic can easily generate misleading code. It is possible in theory to work around such deficiencies. However, in practice, such logic mismatches often trigger huge costs to be paid after indeterminate times. This is particularly true if such logic has to be developed without the necessary time, logic expertise and compiler tools. As many professional programmers know from their own experience, “premature optimization is the root of all evil” (Knuth 1974). This principle readily translates to premature choices of a logic formalism for representing a biological system, prematurely chosen type systems, the premature optimization of implementation speed and others that create numerous problems in computational biology. It is thus conceivable that a substantial contribution towards the notoriously difficult task of funding *VBIR* development might come from the construction of a compiler that simplifies these complex decisions by offering solutions that can represent biological information and uncertainty more easily. ***Conclusions for FlyClockbase***. Accordingly, we have been designing the FlyClockbase type system for maximizing simplicity and radical openness, allowing all users to easily suggest necessary expansions (simply by changing cells in table text files of TabFS). Such suggestions still require review by a *VBIR* architect with expertise in formal type systems. However, more eyes are likely to identify more type-system challenges and might thus inspire better solutions. As indicated in the figure, many solutions are fragmented at first, and larger designs emerge later. Still, the ultimate integration goal for each *VBIR* is to find a single coherent architecture that is “as simple as possible, but not simpler” (Einstein and Calaprice 2011).

#### Cost of not constructing a *VBIR* compiler

Research on circadian clocks in flies can be used to illustrate some of the cost to biology if no *VBIR* compiler is available. As explained above, time series observations are extremely valuable for inferring mechanistic models of clocks in flies. Yet, in the last 25 years, the vast majority of models of the core *D. melanogaster* circadian clock have been based on *abstract* clock features, such as the response to light, the period and presence of oscillations. We conducted an extensive search for such models, and only three of the 66 models identified used specific experimental time series to inform parameters (Figure 4). Even if combined, these models only used a small fraction of all studies with time series data they could have used (Figure 5). Specifically, parameters study A (Fathallah-Shaykh *et al*. 2009), B (Leise and Moin 2007), and C (Kuczenski *et al*. 2007) were based on time series from, respectively, one study (Kadener *et al*. 2007), three studies (Lee *et al*. 1998; Bae *et al*. 2000; Shafer *et al*. 2002), and 11 studies (Hardin *et al*. 1992; Zeng *et al*. 1994; Sehgal *et al*. 1995; So and Rosbash 1997; Bae *et al*. 1998; Lee *et al*. 1998; Blau 1999; Bae *et al*. 2000; Kim *et al*. 2002; Cyran *et al*. 2003; Glossop *et al*. 2003).

The sparse use of directly observed experimental evidence such as time series is understandable in light of the many challenges that complicate the integration of messy biological real-world observations into the abstract mathematical models that are often extremely simplified to facilitate their mathematical analysis. In addition to such conceptual problems, deceptively simple problems such as the storage and organization of very heterogeneous, imprecise, noisy and contradictory experimental datasets can easily create insurmountable practical challenges for directly using experimental time series data to inform parameters in models.

FlyClockbase substantially lowers this barrier by providing a nucleus for collecting, organizing, and curating relevant time series and their many potentially informative *Attributes*. If increasing numbers of experimental time series are deposited in FlyClockbase and its organizational structures keep pace with this growth, then future modeling studies could be structured in a way that enables the automated improvement of some types of models in response to the submission of new data. Such data handling capabilities are likely to enable the investigation of new biological aspects of circadian clocks that are beyond practical limits of the complexity manageable by current tools. If a reasonably well-working VBIRs compiler had been available for a long time, then the substantially lower barrier to the development of a resource like FlyClockbase would most likely have resulted in a more comprehensive use of hard won experimental data in theoretical models. Even where datasets have been compiled and published under open access, an unstructured way of storing them can very quickly make it prohibitively complicated to keep them up-to-date on the longer-term (e.g. (White *et al*. 2013; Supp *et al*. 2015a)). Such problems also pose challenges for citizen science projects (e.g. (Loewe 2007; Supp *et al*. 2015b)). Even if computational results are fully structured from one perspective, the lack of appropriate data structures for analysis from another perspective, can create prohibitive barriers for some research (Loewe 2002).

#### Counter intuitive challenges and other work

However, before such a vision of biological research can move closer to reality, a number of counterintuitive challenges will need to be addressed. Since an appropriate discussion is beyond the scope of this paper and more details are given in the Supplemental Material, we will merely touch on the tips of several icebergs below by discussing a few illustrative examples. Despite great and sustained progress in logic research, much remains to be done to improve the expressive power of known systems of logic (Smith 2008).

##### More biological precision requires handling more imprecision more precisely

The ultimate aim of FlyClockbase is to improve the precision of circadian clock models by making all relevant data easily available for parameter estimation tool. Accomplishing this goal goes far beyond compiling the data. It requires entirely new approaches for dealing with uncertainty, imprecision, contradictions, gaps, and numerous exceptions created by the astonishing diversity of methods used to observe biological systems. These challenges belong to the many predictably unpredictable surprises that will be encountered by any efforts for constructing *VBIR*s of substantial complexity, such as FlyClockbase. Accurately describing biological observations made in this often-confusing context is a substantial challenge we encountered while developing FlyClockbase. We found that it is not enough to ‘describe the issues in words’; this would merely create additional free-text repositories with unstructured information, maybe with a bit more focus than a corresponding collection of PDF-files with the full text of the study. Such free texts could not have enabled us to search efficiently for time series. We found it extraordinarily useful to have key information in a more structured form, e.g., to compare mRNA measurement methods (see above). However, such structure must not come at the expense of the ability to efficiently represent newly encountered imprecision or data. We found that a working biologist with sufficient domain expertise is the best expert for choosing how to handle newly encountered information: ignore, describe in unstructured comments, or create corresponding *Columns* in an *AttributeTable*. Without a substantially sophisticated and extendable system for dealing with imprecisions, very little information will become available for automated processing in more coordinated ways. Ignoring such problems may be reasonable in some cases, but eventually, the inability to handle such imprecisions correctly will artificially narrow distributions and create illusory precision that wrongly rejects simulations as unrealistic and can unnecessarily complicate parameter searches. Conversely, allowing for too broad a margin of error can easily result in a misleading model caused by biologically unrealistic parameters. Thus, appropriately managing errors and uncertainties in observed time series is one important key to improving mechanistic models of circadian clocks informed by the real-world time series in FlyClockbase.

##### Logic in gene regulatory networks

The classical Boolean logic of compilers and gene regulatory networks share an unexpected connection if the input, output, and every step in between are well approximated by just two states (Karlebach and Shamir 2008). Thus, compilers could provide unexpected help for modeling gene regulatory networks. If provided with the right details, compilers could also automatically detect situations where gene regulation becomes stochastic due to low molecule counts in a cell (Macneil and Walhout 2011). The help of compilers that automatically analyze complex logic constructs correctly could prove essential for understanding how complicated binary gene regulatory networks behave if under the control of the daily rhythms of a circadian clock (Lowrey and Takahashi 2004; Doherty and Kay 2010; Zhang *et al*. 2014a). Logic modelling of various genetics problems has a long history (Cotterman 1983; Opitz 1983; Crow 2001) and a bright future helping us to understand many diverse aspects of gene networks and their modular structures (Mitra *et al*. 2013; Le Novere 2015; Saez-Rodriguez *et al*. 2016).

#### Formal systems of logic are usually not logical enough for biology

This paradox is easily resolved by contrasting the complexity of biology with the simplicity typical for formal logic systems. Errors of omission in formal logic limit its ability to express corresponding biological statements efficiently. Omissions have been found to be among those types of errors that are most difficult to detect (Panko 2016).

Our own work on FlyClockbase confirms the substantial frequency and cost associated with errors of omission. Table 5 reports a substantial discrepancy between two types of error rates observed during our exhaustive in-depth re-check of each time series that could in principle affect our main conclusions regarding the variances of peak timing that differ between PER and TIM. Some types of errors could be characterized as ‘simpler errors’ like obvious swaps or typos in FlyClockbase itself. As expected (Panko 2016), these simpler errors occurred at much lower rates (affecting cells of spreadsheets at rates just below 1%). In contrast, we detected a bit over 10% of all time series when re-checking our meta-analysis for systematic errors such as inadvertently omitting agreed-upon steps from routine analyses by trained curators. Much work in our study went into ensuring that important rules were indeed implemented in all applicable cases.

The unlimited potential of omissions to confound biological results repeatedly creates ‘important biological investigations’ aiming to determine whether a given biological conclusion might have been compromised by faulty logic (see our own examples above, where we excluded too much data). While these investigations can be essential for progress in biology, execution often involves excessive, tedious, ‘non-biological’ work towards finding elusive ‘needles in haystacks.’ This metaphor easily takes on ever more complicating levels of nesting when the logical ‘needles’ in question actually consist of errors of omission; finding them can be as challenging as identifying entirely new logical blind spots for the first time. These challenges were felt throughout the development of FlyClockbase from start to finish on numerous occasions, which were too many to track beyond a few illustrative examples.

##### Data quality, plot quality, and task completion

One specific time series figure seemed such a perfect interpretation nightmare that preventing publication of figures like it might be counted as a donation towards supporting FlyClockbase. This figure provided the initial illusion that it shouldn’t be too difficult to unambiguously decide which data point belonged to each of its different time series. However, when actually attempting to extract the values, it slowly became clear that achieving unambiguity was impossible because the figure had been irreversibly degraded to the point where its semantic reproducibility was no longer complete (Loewe 2016). The reason was an unlucky combination requiring *all* the following factors controlled by different entities: (i) the authors’ decision to combine all these time series into a single plot, (ii) miniscule, similar plot symbols, chosen by the authors or the plotting software, (iii) a very low scan quality for the figure, chosen by the publisher, and (iv) substantial overlaps of some of the time series, chosen by nature.

##### Cost

Aiming to collect all relevant data, we did not want to allow for arbitrary decisions that excluded plots due to a curator’s fleeting perceptions of potential difficulties. It took several independent rounds of revisiting by the three most experienced curators and repeated discussions among them before all agreed on the irresolvable nature of this figure’s ambiguities. In total, we spent more than four hours trying to resolve ambiguities (excluding time for finding and initial digitizing). In contrast, it might only have required 10 min of the person producing the figure to choose plot settings that would have completely eliminated more than four hours of work for us. From this we concluded: (i) FlyClockbase needs a reliable mechanism to help future curators avoid such known time killers (e.g.: ‘KK’ in POST Loewe 2016) in the absence of substantial new information. (ii) Efforts such as FlyClockbase need to find principled ways for protecting their limited time resources against irresolvable ambiguous plots or dataset, without delegating decisions on the inclusion of data to a moment’s fleeting perceptions of a single curator. (iii) It might pay huge dividends across all sciences if a targeted effort could improve the clarity of plots produced by typical default settings.

##### Implications for Logic

This example highlights the recurring observation that managing challenging tasks like the one discussed above might benefit from two different dedicated *BioBinary* values, one for ‘progress of the work’, one for ‘results of the work’. Using the *OKScale* in FlyClockbase, a ‘Progress BioBinary’ could store: Progress: *KO* (not started), *OKO* (working), *OK* (done), *MIS* (incomplete because problems occurred). Similarly, a ‘Result BioBinary’ could store: Result: *KO* (has errors), *OKO* (intermediate), *OK* (completed), *MIS* (missing).

#### How to detect logic errors in FlyClockbase

The detection of logic errors can be greatly accelerated by open discussions that invite outsiders to share their observations freely. This could greatly improve the quality of FlyClockbase if this could be made efficient. One of the most notorious bugs is error by omission. This is equally true for omissions in typical program source code as it is in the analysis of biological observations. Omissions are hard to find anywhere (Panko 2016) and can affect the reproducibility of results (Huang and Gottardo 2013) at great cost to science (Freedman *et al*. 2015a). Clearly, a well-defined formal system of logic that is capable of handling biology’s complexities would be a great asset for FlyClockbase and VBIRs in general. It’s formal axioms and rules would exclude many options as impossible, thereby helping researchers to save time by avoiding many fruitless investigations and focusing attention on options of likely interest. Pivotally, such a logic would achieve these aims by providing a conceptual base-line for establishing the few potentially real omissions that need to be considered, while keeping infinities of useless speculations and contradictions from interfering with productive research. If correct, such a logic would be extremely useful for FlyClockbase. However, to the degree that it is misleading, its adoption would waste critical research capacity and lower the chances of uncovering such problems, because people in general rarely question an adopted logic, even if misleading. Thus, using *any* formal logic for understanding complex real-world biology is a double-edged sword, which cannot be avoided when studying mechanisms, including those of clocks in flies.

The idea that a formal logic for biology could facilitate biological research was first expressed only a year after Turing defined the essence of computing (Turing 1936; Woodger *et al*. 1937). Yet, general mutual inspiration aside, it has been difficult to develop a more general formal logic that “makes better logical sense in biology” Most researchers strongly prefer to collaborate on much more specific questions they understand comparatively well, and experts in logic do not often engage with the uncertainty of biological observations. Hence, it has been much easier for most researchers to produce successful special-purpose computing tools for biology, than to arrive at more general solutions. For example, a sorely needed general-purpose programming language designed by biologists for biologists is not available despite all research in bioinformatics, computational biology, and systems biology so far. A credible effort to produce such a language, requires experienced experimental biologists as prime partners on the very same table, where expert logicians design the formal aspects of a logic for biology in numerous iterations. The expressivity of such a logic needs to be tested by its ability to represent actual real-world wet-lab or field-expedition observations. Working on FlyClockbase as described in Figure 11 provided us with such a rare opportunity.

Identifying omissions in the logic of a complex system does not necessarily provide the right resolution and exceedingly many partial workaround solutions are usually found much faster. Such quick-fixes offer immediate relief, albeit at the cost of increasing accidental and historic complexity inessential to a system’s function (Raymond 2003). Without mechanisms for removal, the accumulation of such special case stop-gaps will eventually increase the complexity of a system until it collapses under its own rules. At this point, new potential users will no longer be able or willing to invest the time needed for learning how to navigate the system’s idiosyncrasies. FlyClockbase will not be able to escape this eventual fate, if its data model is not carefully guarded against these problems. Inessential complexity creates numerous difficulties in many contexts, which include defining programming languages, logics, or type systems in computer science (Pierce 2002), rules of operator precedence (Razali *et al*. 2015-10-26), designing data models, databases, or data integration frameworks (De Tre *et al*. 2004; Doan *et al*. 2012), maintaining a user-friendly organization for large libraries, information resources or hard drives, and constructing ontologies, taxonomies, modeling frameworks, or query languages (van Renssen 2005; Zeigler and Hammonds 2007; Rubinson 2014; Arp *et al*. 2015; Hazber *et al*. 2015). With a bit of abstraction, the shared root of these problems might be summarized by asking: “What is the best description of a complex world with all its possibilities for nesting, linking, traveling, and communicating?” The relevance of such logic research for FlyClockbase, is that it greatly simplifies managing the complexities of consistently handling inconsistent biological data in FlyClockbase.

#### Error handling in the face of uncertainty

No *VBIR* of sufficient complexity will be free of errors. This certainly also applies to FlyClockbase. The question is, how to handle errors. Problems with tracing the identity, availability, accuracy, precision, and reliability of data have been the topic of numerous investigations in various contexts, some of which involve big data (e.g. see (Reason and Mycielska 1982; Reason 1990; Reason and Hobbs 2003; Goldston 2008; Doan *et al*. 2012; Gitelman 2013; Grimes *et al*. 2013; Mccallum 2013; Reason 2013; Blankenberg *et al*. 2014; Reason 2015)).

##### Opportunities

FlyClockbase presented us with excellent opportunities for exploring numerous important issues for complex VBIRs aiming to integrate data that is imperfect in some form, such as being incomplete, uncertain, contradictory, erroneous or scattered across a wide range of sources. Any of these conditions occur frequently in biology. It is beyond the scope of this study to explore all such challenges faced by every biologist, whether she’s aware or not; in the Supplemental Material we describe a few of the insights gleaned from our work on FlyClockbase. These could be summarized as follows.

##### Challenges

Any information resource of substantial biological interest will quickly grow to a complexity at which it will inevitably accumulate a substantial amount of human errors that are difficult to detect by human users. Many independent repeats of biological information are typically associated with large amounts of genuine biological variability. In many current biological databases, it can be difficult to distinguish such genuine variability from artificial variability that is easily caused by human errors of various well-known types. Such errors span a broad range of different complexities and corresponding frequencies. For example, simple typos or label swaps usually occur at low rates such as 1%, see (Panko 2016). Simple logic errors occur at substantially higher rates, especially in spreadsheets (Panko 1998; Panko and Aurigemma 2010; Panko 2013; Panko 2016). However, errors of omission are usually the hardest to find (Panko 2016). This is especially true, when an omission has become part of a logic formalism. This is one reason, why it is so important to use good approaches to represent *Null* (White *et al*. 2013), and why it can be dangerous to confuse different types of *Null* (Waraporn and Porkaew 2008; Hoare 2009; Thalheim and Schewe 2011). FlyClockbase has experienced *Null*-confusion already. Entries in a *DZT* column define an hour of the day. We initially allowed *DZT*=0*h* as a valid time, excluded 24*h*, and defined ‘absence’ as ‘*NotGiven*’, a particular type of *‘Nothing*’. However, understanding ‘*Nothing*’ correctly is difficult. Hence, it is unsurprising when biological model curators occasionally allow well-known intuitive algebraic properties of addition to affect their views of ‘Null’. As a result, ‘*0 apples*’ is correctly interpreted as ‘*no apples*’; yet it may fuel the erroneous idea of equating ‘time not observed’ and ‘adding zero to a list of hours. In this case, some *DZT*=0 values are correct and some are not, but checking correctness is complicated and expensive. This test becomes trivial, if 24*h* is included as a valid time, and 0*h* is defined as always invalid. For more details on such challenges, see also discussion of the BioBinary data type in the Supplemental Material.

##### Trans-disciplinary solutions

Several non-biological areas of research and technology, such as computer science, space flight, and nuclear reactor safety have developed sophisticated approaches for detecting and correcting potential human errors (NASA *et al*. 2001-09-30; NASA *et al*. 2006-07; NASA *et al*. 2011; Panko 2016). While designers of biological information resources can learn much from the decades of research that informed the development of human error analysis tools in those areas, it is less straight forward how these insights could be applied to improve the quality of biological information available to most biologists. A source of concern is the substantial complexity of many human error analysis frameworks (Reason and Mycielska 1982; Reason 1990; NASA *et al*. 2001-09-30; Reason and Hobbs 2003; NASA *et al*. 2006-07; Goldston 2008; NASA *et al*. 2011; Gitelman 2013; Grimes *et al*. 2013; Mccallum 2013; Reason 2013; Blankenberg *et al*. 2014; Reason 2015; Panko 2016). Most of these frameworks will handle the complexity of biological data, but require near prohibitive research and implementation efforts that make integration into grass roots *VBIR* projects such as FlyClockbase not efficient if started by biologists. However, that does not imply that sophisticated approaches cannot contribute to solutions, even if VBIRs curators do not bring the expertise necessary for implementing a framework. To see how this might work requires a look at an advanced area in computer science that is not readily accessible to many: compiler construction.

#### Error analyses could be amortized across *VBIRs* by compilers

As argued above, appropriate error analyses for a single *VBIR* are not feasible. However, our experience with developing FlyClockbase suggests that a substantial number of essential tasks are recurrent when compiling any *VBIR* of comparable complexity.

##### Efficiency

Thus, the most efficient solution to improving the quality of *VBIR*s without exploding costs is to develop an automated compiler that can test for all known VBIR problems and that supports a programming language that integrates biology expertise (Loewe 2016). Programmers frequently say that it is important to use the right tool for a given programming task. Despite numerous biology-oriented libraries for non-biological programing languages (e.g. (Stajich *et al*. 2002)) no general-purpose programming language exists yet for supporting typical complier-style consistency analyses for general complex *biological* datasets like *VBIR*s. We will not repeat here the substantial number reasons why such a language would be helpful and why current (non-biological) programming languages are insufficient (see Supplemental Material and additional reasons discussed in Loewe *et al*. (2017)).

##### Examples

Such a compiler could address tasks such as the following. There is a need for handling missing data, inapplicable data and similar cases by choosing appropriate representations that distinguish these cases instead of lumping them together as ‘NA’ or the value zero (e.g. (Candan *et al*. 1997; White *et al*. 2013)). All biological measurements will always come as imprecise ranges, not as precise values. Measurement methods for a given observation are usually heterogeneous and need some description. Observations can be made in may be compared between various *MethodRealms*. like *in vitro, in vivo*, or *in silico*. Comparisons between wildtypes and mutants are frequent. Synonyms are almost ubiquitous. It is easy to continue this list with many other aspects of biological interest. In addition, there are data processing basics, such as the ability to read in all tables of FlyClockbase and produce a report of all inconsistencies and errors that require human attention. The arrival of big data has brought substantial experience with questions of data hygiene (Goldston 2008; Howe *et al*. 2008; Krishnamurthy *et al*. 2011; Gitelman 2013; Mccallum 2013; Schutt and O’neil 2013; Mahmood 2016; Zweig 2016). Most of this expertise is also essential for correctly and efficiently handling data in *VBIR*s. For all features like those above and all error types detected, a solution only needs to be implemented once for simultaneously improving the reliability of all *VBIR*s.

### PopGen predictions on FlyClockbase survival and success

Most new versioned biological information resources (*VBIR*s) such as FlyClockbase face a dizzying array of potential paths into the future, not unlike newly mutated alleles in a population. As population geneticists have learned, all this complexity can be boiled down to two essential outcomes (Kimura 1962): all alleles are either kept or lost eventually. To explore other useful aspects of this analogy, we will abstract a few brief lessons from population genetics that also apply to collections of information.

#### The stage

If seen in such a general way, a newly arisen DNA-allele could be compared to a newly published *VBIR* similar to FlyClockbase or a newly developed tool in bioinformatics (thereby accessing a broader pool of historic precedents). Both alleles and *VBIRs* contain new information, stored in DNA or on computer hard drives respectively. Both are part of their ecosystems, which belong to different realms. An allele exists in carbon-based organisms that compete for natural resources in a population where the allele may be kept indefinitely. Omitting replication details allows for simplification. One could think of alleles as replicators based on DNA; similarly, memes were originally defined as replicators in mindspace (Dawkins 1976). Thus like FlyClockbase, each *VBIR*. can be seen as a meme that competes for the ‘mindshare’ of humans potentially interested in a given topic. Technically, memes are units of information that usually spread through communication and compete for the limited attention of individuals and communities, irrespective of their success of replication (Dawkins 1976; Lynch *et al*. 1989; Gleeson *et al*. 2014; Dawkins 2016; He *et al*. 2016). These generic features result in mechanisms similar to those of population genetics, which we use here to derive informal expectations for the future of FlyClockbase and similar *VBIRs* (a formal theory is beyond our scope). We do so hoping to avoid the most likely outcome, the complete loss of FlyClockbase, by aiming to increase the chances that FlyClockbase will be kept in the population of useful *VBIRs*. We next reinterpret concepts like aging, death, growth and reproduction from the perspective of *VBIR*s; Incomplete Fitness Traits (IFT) like these combine with a given environmental context to define fitness in biological evolution (Loewe 2016). Even without a quantitative meme model, we expect qualitatively similar outcomes when translating IFTs to the realm of *VBIR*s memes. In many cases this will suffice to make decisions that increase the chances of survival for FlyClockbase.

#### Aging and death

*VBIRs* are aging if they degrade without the time and energy investments necessary for maintaining their semantic reproducibility (Loewe 2016); they are on their deathbed when nobody wants to use them anymore, and are buried once nobody can remember them. Potential causes of death vary with age and include (i) being locked into remaining an exploratory toy ‘too simple’ for any real use, (ii) being ‘too simple in comparison’ from a lack of features that could have helped fight competing *VBIRs* and win over their human users, (iii) having become ‘too complicated’ for real-world users after years of accumulating inessential complexity (Raymond 2003), and (iv) many other causes from internal specifics to external generics (such as political decisions).

#### Growth and reproduction

*VBIRs* can grow in various respects, some helpful, some harmful, and some hard to assess. We use ‘growth’ here only in a narrow sense for helpful traits like features required by users. In contrast, we denote as ‘aging’ the growth of harmful traits like inessential complexity, whereas the reduction of such complexity can be seen as growth (e.g. by simplifying an interface to save user time). Likewise, the loss of useful features can be seen as aging caused by semantic irreproducibility. For example, this could be caused by incompatible changes in required software packages. Here growth always affects the quality of the best implementation of a *VBIR* type, in contrast to reproduction, which could be seen as increasing its mindshare through favorable communication and/or copying the *VBIR* data to new servers (presumably to win new voluntary users). Thus, growth and reproduction in this sense are likely to help a *VBIR* to spread and increase its fitness.

#### Speciation and merging

The same is not usually true for processes comparable to speciation. The ‘forking’ of a *VBIRs* or any other software or data collection into two independent lines of development is often perceived as an unwelcome increase in complexity by users without a stake in the details (e.g. Python 2 vs Python 3). This implies that the reduction of independent lines of development should be welcomed, but reality is more nuanced. A reduction from merging without loss of features is positive.

#### Extinction

However, sometimes it is impossible to save all features due to mutual incompatibility or other constraints; this might be comparable to extinction, where good features are irredeemably lost to global mindshare. If occurring to all development lines of a *VBIR* (e.g. due to catastrophic environmental changes such as ‘loss of funding’), then the loss is usually tragic, even if the *VBIR* is preserved as a fossil on cutting-edge archives of its time (like floppy disks, CDs, bioinformatics journals, websites, and various open source repositories). As in real life, software fossils are rarely revived, an act that would require extra-ordinary semantic reproducibility as defined elsewhere (Loewe 2016). Semantic reproducibility is very difficult to achieve, as seen and further discussed in the source code for the statistical analyses in this study and the discussion of the ‘DISCOVARCY’ documentation style (see Table D1 in the Supplemental Material). In both cases, it is much more likely to lose fossils to changing environments and random damage than to revive them successfully. Furthermore, chances of successful reactivation drop dramatically in both cases, as bacteria are easier to revive than dinosaurs, and old algorithms for merely sorting numbers are reused more easily than the software systems that put a man on the moon (though we do not wish to imply that either is possible). Extinction can happen to any *VBIR*, no matter how well known. Some planning can usually ensure preservation of a fossil form; ideally a tombstone will inform would-be users where the fossil is archived (see Supplementary Material).

#### Horizontal gene transfer

As we watched the evolution of FlyClockbase we witnessed a number of remarkable exchanges of information. Our experiences have played out in the conceptual arena defined by Figure 11: we started as initiators, completed the substantial integration work presented here, and have used FlyClockbase for research purposes (see Results). At the same time one of us has been deeply engaged with developing the compiler architecture for the Evolvix modeling language, aiming to meet particular requirements of biology. This combination of aims has enabled a substantial flow of critical design information that has benefitted all sides. Compiler architects benefit from first-hand exposure to challenging practical problems in logic and data modeling in the domain of their target audience, while biologists are kept from computationally short-sighted quick-fixes that otherwise could easily wreck a *VBIR* on the longer-term. Such collaborations are powerful opportunities for uncovering and clarifying misconceptions on all sides and at all levels; in our experience, they greatly improve the conceptual quality and robustness of resulting solutions, but come at the expense of the rate at which some more tangible results can be produced. Combining these costs with those of human error analyses (see above) when developing reliable *VBIRs* increases costs substantially, often prohibitively.

*Practically*, we advocate that *VBIRs* do not reinvent the wheel of reliability independently. This unnecessary reimplementation work is expensive and substantially increases costs of developing and maintaining a *VBIR*. A thorough analysis of historic sources of funding for various existing *VBIRs* has exposed a lack of support for this critical work that integrates, consolidates, and checks the quality of data in *VBIR*s (Ember *et al*. 2013). An overview of these essential tasks in the context of FlyClockbase is given in Figure 11. Here we suggest that much of these costs could disappear if the initiators, integrators, and researchers working with a *VBIR* would have efficient means of passing on their formal needs for data representation and analysis to the architects of an integrative compiler. From their integrative perspective, these architects could then provide solutions that are compatible and interoperable for many *VBIR*s. Support for such a versatile open source compiler-building project that serves the *VBIRs* community well would not nearly be as expensive as independently solving this problem repeatedly. Experience indicates that well-maintained tools do get used; such a project could hence substantially contribute towards closing the critical funding gap highlighted by a thorough analysis elsewhere (Ember *et al*. 2013). Here is not the space to provide a reasonable overview of the many aspects of working towards an integrated compiler architecture. Informed by experiences with FlyClockbase, the tips of several icebergs are touched in Figure 11. It lists important needs of various contributors, and specifies several types of lessons learned by *VBIR* contributors and services provided by the compiler and its construction team envisioned here. This work generally occurs in three broad stages of integration: combining fragmented insights gleaned from work on FlyClockbase, investigating broader designs, and integrating solutions into a single coherent architecture. The high-level analogy of aging and growth in *VBIRs* plays out on the background summarized by Figure 11. The values of such *IFT*s governing the evolutionary trajectory of *VBIR* meme evolution are determined by the hundreds of small implementation decisions necessary for arriving at an overall coherent *VBIR* organization perceived as elegant, expressive, useful, efficient, and overall simple enough to be worth a user’s while. Such simplicity is pivotal for engaging anonymous users with a *VBIR* (or any other meme), especially since many are suffering from information overload, data smog, and the resulting paradox of choice (Shenk 1997; Schwartz 2004). Physics is not the only discipline where theory should be “as simple as possible, but not simpler” (Einstein and Calaprice 2011).

#### Potential predictions

So, what can we learn from population genetics to improve the long-term usefulness of FlyClockbase? We do not aim to exhaustively list all general lessons but rather present several possibilities of likely interest in the bigger picture that we expect based on population genetics theory. This will allow readers to connect additional dots between more detailed requirements and solutions presented above as well as in the Supplemental Material. As we review the following potential paths into the future we interchangeably use the terms ‘FlyClockbase’, ‘new allele’, ‘bioinformatics tool’, and *VBIR* to reduce repetition.

**Loss is likely** for all new information. Population genetics theory shows that most newly arisen alleles are lost very quickly by the random sampling that occurs between generations (Kimura 1962). All alleles have to navigate this hurdle, regardless of how beneficial they might otherwise be. Observing bioinformatics research quickly reveals a similar pattern: on the web very many tools start out (and fizzle out), professional researchers ensure that at least one peer-reviewed publication exists (but lack the time to keep websites and tools from breaking), enthusiastic programmers will keep tools working (but are happy with little documentation), good software engineers understand the value of organization and documentation (but usually do not work in biology). All new tools and resources face an intimidating phalanx of these and similar dilemmas, which made us think hard about all possible avenues for simplifying the overall system while increasing flexibility. First lesson: FlyClockbase is no exception and faces the same challenges. It may sound strange to discuss death in the context of a birth that we believe is to be celebrated. However, ignorance is not a good defense against child mortality.

#### Loss is fast and ‘child mortality’ matters

Alleles that have just arisen by mutation and new bioinformatics tools that have just been published also share another important detail: they will probably be lost very soon. Except for extremely harmful alleles, initial survival for good and bad alleles depends almost entirely on the individual that carries them. Therefore, FlyClockbase must travel as light as possible if it is to survive. Like other *VBIR*s, it must be able to fit it into the life of a single publicly known person who can act as a synchronizing point of contact for coordinating further work (even if not done by that person). Such public maintainers of *VBIR*s are probably extremely busy and will have very little time and energy left for high-maintenance solutions. This excludes the use of many great database technologies that unfortunately shower their users regularly with recommended updates with various degrees of compatibility and urgency. Without a highly-automated process, such updates would prohibitively increase the rate of aging for FlyClockbase or the energy required to maintain it. Accordingly, we have been developing approaches to simplify life with *VBIRs* like FlyClockbase, but much more remains to be done. Our various strategies for simplifying are discussed above and in the Supplementary Material. Laurence Loewe has agreed to be the first public maintainer of FlyClockbase and will post updates to the GitHub website given at the beginning of our description of FlyClockbase.

#### Fossilization is usually deadly

An easy way of avoiding the loss of a project from a broken hard drive is to submit it to a public repository such as Github. This ensures a form of travelling light, as everything stays in place, if a maintainer does nothing (maintenance cost is near zero, and mostly a thought and a password). This establishes a minimalistic baseline, as the mere existence of data (or ancient code, see above) does not differ much from fossils, which are awkward to access, dry and brittle to work with, and for all practical purposes impossible to revive. Lesson three: If nobody continues to work with the code, then chances are that it has already fossilized. Thus, we next review steps that are likely to facilitate future work with FlyClockbase.

### Next practical steps for FlyClockbase

In order to raise the chances of survival and success as described above, we are working towards implementing the following practical steps that improve the organization of FlyClockbase and move it towards increased stability.

#### Reorganize files, define versioning policy and simplify folder structure

It is very frustrating to work with a project where everything can move (and break) at a moment’s notice. Nascent resources never really know what awaits them, and FlyClockbase has not been different. As a result, our time series data has seen more profound reorganizations of its storage space than any of us had anticipated. Some of this additional work was due to the fact that we were simultaneously developing crucial technological underpinnings, such as TabFS (Figure 11) and the POST system (Loewe 2016). We also did not have a stepwise guide on *VBIR* construction with an overview roadmap from an expert, which could have further reduced the work. However, as indicated in Figure 11, the initial phases of a *VBIR* will always be special: each *VBIR*, by definition, is ill-defined at its inception and negligibly small. As it starts growing, it is restructured, renamed, and reorganized many times while in its ‘embryonic’ form. While guidance helps, some messy aspects of initiating a *VBIR* are probably impossible to avoid if the freedom is retained to develop any *VBIR* supporting any research. Figure 11 makes a clear distinction between this early more informal stage and the subsequent iterations managed by integrators. While much of the work of integrators is also done at the initiator stage, *VBIR* publication marks a milestone. It is an excellent opportunity for internal restructuring and cleanup that should not be dismissed lightly, as reorganizing will never be as easy again. This is particularly true if the versioning system changes. Long-term resources require stabilizing versioning systems right from the start to reduce inessential complexity and confusion. FlyClockbase will build on the *StablizingZone* (Loewe 2016) of the POST system. These needs motivated us to delay publication of FlyClockbase while still under review.

#### Use a public distributed version control system to be efficient

The use of Git for version control is rising and services like http://github.com allow open source projects to be published ‘at no cost’. Not having to pay for leaving code in a published state increases chances of avoiding ‘death by negligence’ for many *VBIRs*. More importantly, using Git allows *VBIR* collaborators to essentially cut the huge costs of manually performing the search and merge operations regularly required for close research collaborations. We have experienced enough of these complex operations to appreciate the huge value provided by Git and have decided to use it for FlyClockbase (currently in a closed repository using http://gitolite.com/gitolite/). However, using Git is not free of costs. At first these seem reasonable: learn how to use Git and avoid advanced moves that get ‘the rest of us’ into serious trouble (including loss of data). However, in our experience, Git idiosyncrasies and the complexities of version trees pose such formidable barriers for most biological users, that tool adoption requires a large activation energy, even when using excellent graphical user interface software (albeit developed for programmers). We have found an approach for getting biologists to work with reasonably well with Git. It currently requires determination, detailed instructions, an expert who performs all operations except the very simplest, and who happily explains everything again until users follow the instructions (cleaning up the mess, if they do not). Given the outstanding efficiency of Git, not just for FlyClockbase, motivated us to explore how to hide our simplified Git workflow behind scripts called when users ‘hit a button’. While our design requires more development and testing, our internal results so far suggest that it will be more than worth the effort develop this for FlyClockbase (and reuse for other *VBIR*s). We highlight all this because many biologists seem unable to imagine how much more efficient the development of a *VBIRs* can be if Git works as it should. Conversely, many Git users seem unable to imagine why some biologists prefer to explore every non-Git option first, irrespective of cost. We found that some of these ‘attractive’ alternatives can easily turn into complexity traps or create serious bottlenecks for development. This is in particular true for the prevalent mode of distributing supporting material for journal articles, which allows reading data from files without the ability to write back. Such immutability is good for ensuring well-defined versions, but unless there are files that can actually be updated in place, it will be very difficult to efficiently work with the data. Complications can abound when manually merging two sets of text files, each with changes that accumulated in separately evolving lines of revision descent. Since not all combinations have been tested together, some could trigger prohibitive integration problems comparable to the severity of Dobzhansky-Muller Incompatibilities known from evolutionary genetics (Coyne and Orr 2004). In this analogy, using a system like Git for regularly merging all new changes into a single main line of revision descent is comparable to keeping all individuals in one large population. This efficiently prevents the accumulation of the source-code equivalent of Dobzhansky-Muller Incompatibilities and is desirable for improving one *VBIR* that serves a single purpose. Thus, typical archival data storage is not ideal for FlyClockbase and similar *VBIRs*; we therefore chose Git and aim to mitigate its less than ideal aspects.

#### Develop TabFS

For decades, there has been no shortage of databases, file formats, file systems, and other types of storage - all with unique strengths and weaknesses. There is no universal agreement on how to best store complex data transparently. Text-based formats that distribute data across folders in a file system provide instant and continuous access to content that is easy to read and write for humans. This flexibility does not depend on any special tools that could break. However, such transparency benefits are balanced by the need to ensure consistency in the presence of notoriously inconsistent human users. While binary formats increase speed and consistency, they complicate *VBIR* development and create costly dependencies on special tools for reading or writing *any* data. As *VBIR* development requires biological model curators to easily modify the data model of a *VBIR*, we decided against using existing excellent binary technologies such as ProtocolBuffers (https://developers.google.com/protocol-buffers/) and HDF5 (https://www.hdfgroup.org/hdf5/). For these and other reasons detailed in the Supplemental Material we decided that *VBIR*s and TabFS require human readable text-based file formats. Appropriately reviewing these is beyond the scope this paper, but some recurring patterns provide food for thought. For example, the text-based ‘eXtensible Mark-up Language’ (XML) and the representation independent ‘Abstract Syntax Notation One’ (ASN.1) are both widely used, formally defined (https://www.w3.org/XML/ - https://www.ncbi.nlm.nih.gov/Structure/asn1.html) and demonstrate the following possibilities for data storing file format standards:

1. it is possible to define broadly applicable standards that maintain a very simple and stable core set of features (encouraging simplicity in TabFS);
2. combining a few built-in data types with arbitrary nesting and repeating of user defined data types can inspire multitudes of specific extensions (suggesting TabFS will need to help users navigate diverse complex *VBIRs* code contributions to reduce complexity and unnecessary reinvention);
3. fierce competitors are capable of adopting shared standards resulting in win-win-win outcomes for both competitors and the general public; see the use of ASN.1 in telecommunication (and also by NCBI, see link above; the research benefits of a system that significantly simplifies the sharing of complex biological data are undisputed; developing FlyClockbase across the Win-Mac divide together with the Project Organization Stabilizing Tool (POST) system (Loewe 2016) by using the process in Figure 11 led to major TabFS design ideas that could simplify current challenges enough to motivate grass-roots adoption; details beyond scope here);
4. the rise of simple text-based XML could not quench the need for even simpler text-based file formats that are popular for simplicity where it matters (suggesting TabFS needs to provide comparable ease of use);
5. simplistic file formats are insufficient for representing many types of biological complexity and will therefore never be adopted universally (suggesting TabFS must handle arbitrarily complex data in elegant ways).

Comma Separated Value files (CSVs) and the equivalent tab-delimited table files of TabFS are still particularly convenient file formats of choice due to their simplicity and extraordinary broad interoperability. Research collaborations frequently share data across very different systems. Thus, a file format that can easily be read and written everywhere remains competitive against faster rivals that do not work everywhere. Unfortunately, CSVs store only values, but cannot store types and cannot directly describe arbitrary data structures. Therefore, all additional information requires extensions that are rarely standardized. Recent text-based standards like JSON (see http://json.org) or YAML (see http://yaml.org) cover many use-cases, but have not replaced CSVs in many contexts.

##### Tables in their simplest form

The two-dimensional layout of CSVs is particularly well suited for time series, arrays, and other frequent forms of biological data. CSVs are easy to read and write with spreadsheet tools that are widely used among biologists. Many experimental biologists would not hesitate to use such tools for modifying sets of CSVs but would avoid equivalent tasks in SQL databases. This fundamental usability advantage of text-based tables motivated our data storage choices for FlyClockbase. The downside to this flexibility is the lack of formally defined computational expressivity that is powerful enough to represent all the needs of *VBIR*s. Our numerous searches have brought many interesting file formats to our attention, but none approaches the simplicity and usability of CSVs while also providing a stable international standard with the features necessary for efficient *VBIR*s development. This gap surprised us.

##### TabFS specification

We plan to fill this important *VBIR* tool gap by developing a definition and implementation of TabFS. The TabFS specification aims to define precisely a completely open and customizable, easily accessible and usable, extremely simple and stable, maximally versatile and expressive storage system for long-term use in *VBIR*s such as FlyClockbase. Here *long-term* indicates the requirement to be long-term backwards compatible as defined by the ‘*TrustedTested*’ (TT) level in the POST system defined elsewhere (Loewe 2016). To achieve these goals, some aspects of computational speed will receive a lower priority in TabFS, as speed of *VBIR* development is more important for TabFS than speed of execution. Practically, TabFS builds on the stability of standard file systems and uses tables and other fragments in files and folders to implement well-defined conventions for storing the necessary nuances required for *VBIR*s development. A major design aim is to keep the raw convenience and efficiency of tab-delimited table text files (hence the name TabFS). Such files are easily edited by spreadsheet tools familiar to many biologists and readily imported and exported by many other systems. TabFS is developed in the context of *VBIR* development as described in Figure 11 and uses the flipped programming language design approach presented elsewhere (Loewe 2016). Many *VBIRs* share similar problems, some of which are typical for biology. Solving them once in a reusable way can greatly contribute to the reproducibility and the sustainability of domain specific resources of digital data (Ember *et al*. 2013).

#### Define a type system for TabFS

Work towards defining each essential data type for TabFS in general and *VBIRs* in particular will need to continue in parallel to developing TabFS itself. Substantial overlap in development is essential for ensuring that TabFS provides all important capabilities for making high-level *VBIRs* development efficient while minimizing overall system complexity. Establishing a stable core of TabFS first will greatly shrink the complexity of developing a stable and consistent type system for recurrent tasks in both TabFS and general *VBIRs* development. The same mechanisms will later be used by developers of any specific *VBIR* to define a type system for their particular area that can then be enforced with the same mechanisms that protect the integrity of TabFS or general *VBIR* types. Since type systems are conceptually equivalent to ontologies at a high level (Arp *et al*. 2015), such work can be structured in work-stages that are familiar to biologists since the start of taxonomy: observe, describe, define. Practically:

1. *Observing* which types of folders, files, or fragments are useful for developing and maintaining a *VBIR* is only possible in the context of a real *VBIR* with real research problems, such as FlyClockbase. Pure thought or toy projects cannot reveal enough real-world nuisances and nuances for developing a high-quality *VBIR* type-system. The next step for the resulting list of observed entities is:
2. *Describing* at epic length in human readable text every detail about, why and how exactly each folder, file, or fragment is stored and used by expert biological model curators provides a solid foundation for the final step of explaining all this to computers:
3. *Defining* each type formally, which results in a checklist for determining the integrity of this *VBIR* type and for detecting all known errors.

Initially, such checklists are best developed and refined by expert users willing to accept a temporary slowdown caused by the need to document and check every step of their work, including the simplest ones (tedious for humans, essential preparation for computers). Once sufficiently detailed, these checks can be automated, enabling experts to focus their energy on complications that computers cannot currently handle correctly. Such a style of collaboration with *VBIR* compilers would allow all parties to focus on what they do best. Machines mindlessly repeat mind-numbing instruction sequences. In contrast, experts focus on activities where humans excel: apply expertise and common sense to check the integrity of computational results, think creatively about new tasks for the *VBIR* compiler, and expand the *VBIR* by exploring interesting hypotheses. Once this *VBIR* has matured enough to answer the interesting questions in its field, start a new *VBIR*.

For the first several *VBIRs* most contributions to such a compiler will probably focus on the basics of defining and referencing various types of memory devices, such as folders, files, file names, tables and fragments of these. A well-defined type system will greatly simplify the implementation of the consistency checks that are essential for maintaining the integrity of FlyClockbase.

#### Automate TabFS checks to help expand the biology of FlyClockbase

Developing FlyClockbase, TabFS and a *VBIR* compiler for ensuring the long-term stability of *VBIR*s can be greatly facilitated by a code library implementing a storage interface for TabFS instances. Detecting formal errors, enforcing rules and limitations, ensuring the full execution of all aspects of a TabFS or *VBIR* task, and performing other jobs can then be delegated to such a storage library and will no longer consume precious development or research time. These new liberties can then be invested in expanding the reach of FlyClockbase by adding the latest biological studies, new and old mutants, and many other aspects. Additions require defining new columns or new values for the controlled lists of existing columns. Carefully reviewing anticipated usage reduces clutter in the name-spaces of FlyClockbase. This is pivotal, since column names become immutable once pronounced ‘*TrustedTested*’ as defined (Loewe 2016). The ability of FlyClockbase to disentangle the long-term need for stability and the short-term freedom required for *VBIR* innovation will critically depend on the early introduction of a well-thought out stabilizing version number system for FlyClockbase, lest it be killed by inessential complexity on the long run.

### Conclusion

This study contributes important foundations to our overall goal of improving the reproducibility, reliability, and relevance of biological data analyses, starting with observations of the *D. melanogaster* circadian clock. To this end, we aim to automate as many repetitive tasks as possible by providing computational tools that can be efficiently used by experimental biologists. Ideally, this will inspire increased adoption of computational tools and empower biologists to expand their thinking capabilities to investigate new questions. This will be required to meet current grand challenges from personalizing medicine to predicting mechanistic fitness landscapes in evolutionary systems biology (Loewe 2016). Such types of problems often require the analysis of innumerable smaller computational models, which is impossible without highly automated information processing to cut through the associated cognitive complexity.

#### FlyClockbase as a *VBIR*

The resource we compiled might be able to serve as an example for a versioned biological information resource that is organized in a radically simple way by being completely accessible as tables of text. It also exemplifies what a ‘small model’ in a grand challenge context might look if comparable in size to our clock model (see Figure 1) with similar amounts of time series or other experimental data. We expect such data to be as scattered as it was for FlyClockbase. Experience with time series in FlyClockbase suggests that many other datasets are probably also likely to contain a mix of broad general trends and numerous statements that remain incomplete, imprecise and contradictory. To successfully handle this avalanche of challenges in biology, we have been analyzing observations and models of the fly circadian clock. Simultaneously we have been collecting instances, where automation by a compiler could greatly increase the efficiency of integrating biological knowledge-fragments and maintaining the integrity of a VBIR in face of common uncertainties in biological data.

#### Designing a compiler for biological data

The design of such a compiler is greatly improved in our experience, when developed simultaneously and in close collaboration with biological model curators who regularly expose compiler designers to the many imperfections of biological data. The seemingly perfect abstractions of compiler type systems need to meet the messy observations made in biology, and conversely, biological observations need to become more organized by learning from the abstraction techniques developed in computer science. Such trans-disciplinary communication is possible in our experience (see Figure 11 for an overview of the process). Consequently, our work in this study drills deep in distant areas from different disciplines, both basic and applied. The volume of relevant material forced us repeatedly to refer to Supplemental Material, the Evolvix BEST Names study (Loewe 2016), or simply limit scope (usually indicated). A brief overview of the relevant research areas might illustrate these challenges for compiler construction.

#### Trans-disciplinary aspects

The seemingly disparate areas of enquiry in this study are deeply connected by our desire to improve the reproducibility and reliability of models in computational molecular systems biology. We study:

i. *the molecular genetics* of gene regulatory networks in *Drosophila* circadian clocks (reviewed in Figure 1),
ii. *the statistics of robust differences* in variance among observed time series traits (Figure 5),
iii. *the applied mathematics of simulating* time series from Continuous Time Markov Chain models (Figure 4 lists models, leaving simulation for later),
iv. *the behavior of modelers*, namely how they prefer to parameterize their models (Figure 4),
v. *the human-computer interactions* that help to reduce data smog and information overload by improving visualization and organization in plots, in models, in and data structures (Figure 1,2,6,7,9),
vi. *the statistics of detecting human errors* in spreadsheets, data analysis, logic, and source code (Figure 3, Table 5, Discussion, Supplementary Material),
vii. *the data science of reproducibility* for improving reliability, semantic, statistic, and other reproducibility of publishable research results from the early investigative stages (see Supplemental Material, Table P1 and the ‘DISCOVARCY’ Documentation Style), and
viii. *the computer science of compilers and programming languages* as needed for supporting the development of other biological information resources like FlyClockbase. This requires addressing a broad range of topics, including mathematical logic, type theory, arithmetic, syntax, semantics, memory organization, naming, and others. Figure 11 provides an overview of the types of interactions we have observed between biological model curators and a compiler architect while developing FlyClockbase.

Thus, we touched the tips of many icebergs and often needed to limit our scope. Much of this tension was caused by our desire to build a compiler that understands the imprecisions and complexities of biology and supports the efficient construction of high-quality *VBIR*s. We have pursued this goal by constructing such a *VBIR* and performing manually all tasks that we would like to delegate; this gave us the opportunity to reflect on the nature of the tasks and the quality of the outcome. This reduces the speed of both: compiler construction and *VBIR* construction, but simultaneously greatly increases quality. As argued by our analogy to aspects of population genetics theory, such increases in quality can be pivotal for the survival of a *VBIR* like FlyClockbase, which can easily be killed by small increases of inessential complexity. In this study, we provided a broad overview of this tandem work. We have removed from this paper all aspects that can also stand on their own. For example, readers of this journal might be less interested in a formally complete description of the data structures that comprise Evolvix and the nuances of data models that contribute towards long-term stability. We endeavored to keep in the main text only those computational aspects that are most important for navigating the broader concepts used in FlyClockbase or that convey a general overview of our approach to reducing the cost of maintaining digital resources with the help of a compiler designed for this purpose. There is no reason why such a compiler could not be used by individual researchers collecting their own data, some of which they might want to share later. Therefore, our work presented here could also be seen from the following points of view.

#### View on gene expression variability

The most direct purpose of our study is to use FlyClockbase to generate and analyze hypotheses about circadian clocks in *D. melanogaster*. We analyzed patterns of circadian variability across diverse independent studies of fruit flies, accumulating the largest number of time series for this purpose to date (to our knowledge). We have used the statistical power of FlyClockbase to detect consistent differences in the variance of peak times for the important clock proteins PER and TIM. This led us to hypothesize that these differences have mechanistic causes that are worth investigating with the methods of computational molecular systems biology (out of scope here). Our detailed analysis of variances in the peaks of PER and TIM and the potential causes for outliers (see above) suggests the removal of outliers by default using the method of Carling (2000) to focus more efficiently on estimating what typical clocks usually do (without suppressing natural variability in time series). Similarly, FlyClockbase can be used to compare the accuracy of different observation methods (Figure 8) and many other *Attributes*. An important contribution of FlyClockbase towards simulations of fly clock models of gene expression variability is its rich set of over 400 wildtype time series that can be used - in principle - to improve estimates for circadian clock parameters. Such estimates might change the rather sobering observation that most clock modelers do not use most experimental observations when deciding on the parameter values for their simulations (see Figure 4). A study using state-of-the-art inference methods for obtaining the best possible clock model has been moved beyond the scope of this paper but could start immediately.

#### View on simplifying *VBIRs* development

The broader purpose of our study is to develop, describe, and use FlyClockbase as a real-world testing ground for designing an extraordinarily reliable yet simple system for long-term backwards-compatible data integration. We also explored how to annotate, name, reference, identify, store, query, retrieve, and analyze the imperfect and complex biological data and its translation into well-defined computational concepts. Developing these capabilities is essential for the long-term mission of programming languages like Evolvix that aim to provide built-in support for biological research. This goal requires unusual amounts of direct user feedback from experimental biologists to the language designers, as described elsewhere (Loewe 2016). Since computers and their computations are ultimately abstract, software engineers have come to value the input of so called ‘domain experts’ without whom it would be impossible to develop efficient and reliable non-trivial systems. Such feedback is easier to provide in engineering and other technical scenarios where domain experts and software engineers tend to speak a similar language. However, such a shared language does not usually exist in biology where the ‘domain experts’ are experimental biologists who often are not used to expressing their expertise in a form easily understood by software engineers. It is an important goal of Evolvix to fill that gap and enable the best experimental biologists to express their expertise in a form that is readily translatable into computable models. Simplifying the construction of VBIRs is an essential component of this larger goal and critically important for evolutionary systems biology (Loewe 2016).

#### View on Evolutionary System Biology

The ultimate long-term purpose of FlyClockbase is to substantially contribute towards implementing the vision of mechanistic simulations in evolutionary systems biology as detailed elsewhere (Loewe 2009; Loewe 2012; Loewe 2016). Evolutionary systems biology aims to quantify fitness landscapes by mapping genotypes (via realistic fitness causality networks) to phenotypes and ultimately fitness. Since circadian clocks have a large impact on fitness, their behavior is of direct evolutionary importance (Beaver *et al*. 2002; Beaver *et al*. 2003; Dodd *et al*. 2005; Loewe and Hillston 2008; Akman *et al*. 2010; Beaver *et al*. 2010). Constructing a high-quality model of a circadian clock in *D. melanogaster* could thus provide the opportunity to explore many mutant options *in silico* (Loewe and Hillston 2008) and thus bring us closer to the goal of quantifying fitness landscapes of interest (Loewe 2009; Loewe 2012; Loewe 2016). To enable this vision, myriads of models on the scale of FlyClockbase will need to be constructed, connected and analyzed both individually and in various combinations. Most of today’s tools do not manage imprecision with the high degree of precision that is needed for integrating models at such a scale. To address these problems, we need the *VBIR*s automation discussed above and other new approaches to biological model curation.

#### Biological model curation

The substantial needs for biological model curation illustrated in this study highlight a challenge faced by biology as a discipline. Researchers have accumulated very large amounts of biological data that is currently scattered across the scientific literature in forms that are difficult to access efficiently (or become completely inaccessible as lab notebooks are being thrown out or primary data is lost from hard drives). In FlyClockbase we integrated scattered data from across the literature. The substantial amount of work involved forced us to acknowledge, that it is not possible to engage in the integration of biological information at this scale without a substantial investment of time. Even if *VBIR*s construction is eventually simplified to the highest possible degree by the most user-friendly compiler and *VBIR*s construction environment imaginable, the need for model curation in biology will not become trivial. On the contrast, such a compiler could motivate a new generation of biologists to actually revisit and integrate data that has long been ignored, because using it without compiler support would have been too tedious. This possibility will likely boost interest in a currently unusual avenue to biological research that is not well represented in the biological job market of today.

##### Status quo

For a long time, most biology undergraduates have been aiming to work at the bench in a wet-lab. Biologists overly focused on wet-lab work might undervalue the importance of biological model curation by underestimating the intellectual efforts it requires. However, what use is experimental data if it remains inaccessible? While biological model curation does not generate new data *per se*, it makes existing experimental observations accessible in integrated forms. The resulting information repositories, such as GeneBank, are prime sources of data used by computational biologists. The rising importance of computational modeling and bioinformatics in biology is now recognized well enough so that students in these areas can readily self-identify and point to labs, role models and career paths. Such computational professions require substantial training in formal methods, quantitative approaches and computational tools – usually not easily understood by experimental biologists who dedicate their career to investigating a particular system in great detail. Conversely, many computational, mathematical, and other programming biologists struggle to develop enough dedication for a career committed to studying a single biological system. The time they take to develop their computational expertise takes away from the time they have to develop their biological intuitions to the level required for high-quality biological model curation.

##### A growing avenue to biological research

Work on biological model curation which was integral to obtaining the results we presented alerted us to a rising need for the integration of biological data. As shown by the new biological insights presented in this study, biological model curation is as essential to biological research as bioinformatics algorithm development, original lab observations, and field data collecting. It does not stand behind lab experiments or computational work in its potential for contributing new biological insights. The low entry bar to model curation should not be mistaken for a lacking ability to advance the cutting edge of science. Each major avenue of biological research has trivial activities that do not speak to its potential for biological innovation. Pipetting samples into tubes does not reflect the complexities of experimental biology. Defining the initial values for a few variables in a program does not reflect the potential for innovations from computational biology. Similarly, the simple activity of comparing a few numbers from a few studies in a spreadsheet does not reflect the importance of biological model curation for progress towards addressing grand scientific challenges. In our experience, in depth biological model curation for non-trivial questions requires a substantial amount of attention that will not realistically leave much room for additional work on the side, whether in wet-lab or in computation. The FlyClockbase work present here demanded the undivided attention of several researchers and integrators. Model curation work is easy to scale up or down, but significant new findings still require dedicated resources – as everywhere in research.

##### What it takes to do biological model curation

While biological model curators are still rare, their work has more history that commonly known (see Introduction on biocurators). Biological model curators must have sufficient interests in the wet-lab work necessary for generating the observations they curate to know about typical pitfalls, but they typically do not work at the bench. They must be sufficiently aware of the strengths and weaknesses of relevant modeling approaches and extract the most relevant information from the scientific literature, but they do not need to be expert programmers. Most importantly, they need a passion for ‘their’ system to the point where they want to know everything about it, irrespective of the method used to observe it. This will enable them to accumulate enough expertise for learning about the strengths and weaknesses of different methods of observation and for developing an intuition about the quality of a given data set. Such expertise is essential for helping to improve the overall reproducibility of statistical processing pipelines by improving quality of relevant input data, as recently called for (Leek and Peng 2015).

##### On the shoulders of giants

We aimed to stand on the shoulders of giants in fly clock research. This would have been impossible without the biological contributions from the high-quality model curation work that resulted in FlyClockbase. To enable more biologists to stand on the shoulders of their giants we have been working towards capturing our experiences with FlyClockbase in the definitions of *VBIR*s. We expect that constructing a corresponding *VBIR*s complier will greatly accelerate the integration of the biological expertise required to meet the grand challenges of our time. One of these is to understand the long causality chain that starts with the daily rhythms of core clocks and ends with detailed mechanisms for the changes in health and fitness caused by the daily rhythms of the thousands of genes under circadian control.

## Acknowledgements

We thank Jerry Yin, John Hogenesch and Chris Bradfield for stimulating discussions of circadian clocks, methods of how to observe them in mice and flies, and how to interpret aspects of these observations and of our findings. We thank John Yin, Millard Susman and Anthony D. Pietsch for many discussions on more general aspects of this work and numerous comments on earlier versions of this manuscript. We thank Cecilia Moog, Jocelyn Meyer, Sarah L. Northey, Ginger Ann Contreras, Coco Contreras. Kimberli Ward, and Martha Loewe for helping to refine some sections of the text, as well as reviewing BioBinary definitions and suggesting substantial relevant improvements.

We thank Alyssa Hotz, Claire Nusbaum, Devanie Tucker, Noah Waters, and Keegan Reilly for stimulating discussions of various aspects of circadian clocks and their importance and all other members of the Loewe Lab who have in various ways contributed to innumerable discussions of requirements for a programming language that can handle the uncertainties and complexities of biology.

We are very grateful to Ben Liblit, David Page, Ines Dutra, David Anderson, and Michael Ferris for numerous comments over the years on a wide range of technical, formal, and mathematical details that have shaped in several ways the design of FlyClockbase and/or aspects of analyses performed in this study.

We are particularly grateful to Seth A. Keel for setting up the initial Git repository of FlyClockbase and helping with numerous Git-troubles. Most importantly, he contributed critically to the stability of FlyClockbase by highlighting the flexibility-vs-performance trade-off in databases-vs-filesystems: he made us realize how databases usually compromise flexibility, long-term stability, and low maintenance costs in order to gain performance in searching for answers to prepared types of questions. In contrast, for research we need maximal flexibility, reliability, accessibility, recoverability – all at minimal complexity and costs. Seth showed that these traits are more important than faster answers to more restricted, recurrent questions, since question types in research are unpredictable.

## Funding

We thank the Welton Family Foundation for an early grant to K.S. that helped to start this work, the National Science Foundation for NSF CAREER Award 1149123 to L.L. that supported most of this work, and the Wisconsin Institute for Discovery at the University of Wisconsin-Madison for general support. Any opinions, findings, and conclusions or recommendations expressed in this material reflect those of the corresponding author and do not necessarily reflect the views of NSF or other funders or contributors to this work.

## Author contributions

### Conception and design

K.S. created, designed, developed, tested, and repeatedly improved most aspects of FlyClockbase, *PeakValleyTables*, and their initial statistical pilot analyses that indicated which results were interesting for our first analysis, eventually leading to the analyses presented here. K.S. was supported by J. Dr. and E.N. in many practical aspects, including substantial re-organizations of FlyClockbase; she was also supported by L.L. in theoretical, modeling, and various data layout design aspects, as well as in questions of computational modeling, and managing connections to Evolvix. After L.L. set the initial stage by formulating the overarching goal to work towards creating the best possible circadian clock model for *D. melanogaster*. K.S. took the idea and in several iterations developed the biological model in Fig. 1 which drove many biological aspects of FlyClockbase and of this paper; L.L. conceived the approach for investigating error rates and potential blind spots of poor reliability in FlyClockbase.

L.L. conceived, designed, and has been continually developing Evolvix, TabFS, POST, and various other concepts in Evolvix that also underpin FlyClockbase and governed its technology choices. In his design work he was substantially supported by K.S. who conducted countless usability reviews and provided numerous suggestions for improving, clarifying and simplifying many diverse aspects of Evolvix, TabFS, logic systems and operator precedence problems from a user’s perspective. Her feedback included conceptual suggestions, architectural aspects of item-type-context interplays with BioBinaries to indicate TS analysis integrity, numerous ideas for syntax and name choices that improve the consistency and visual elegance of many important punctuation symbol choices for Evolvix, TabFS, and FlyClockbase. L.L. conceived and designed the overall strategy of mixing disciplines by combining in-depth molecular system biology model curation work integrating all expertise and observations about a system, with research relevant for compiler construction. The latter includes in-depth analyses of the limits of some formal logic systems and the practical aspects of developing an architecture that better supports biology by developing a readable syntax and naming-support for facilitating VBIR curatoration.

### Experiments and acquisition of data

K.S. collected and organized all data (as described above, supported by J.Dr. and E.N.). J.Da wrote a script that greatly facilitated many early explorations by K.S. To measure human error rates, L.L. developed a strategy in close collaboration with K.S. and J.Dr., who both executed this strategy (with assistance from E.N.) by thoroughly searching for human errors that – despite all caution – had persisted into *Mod5* (resulting in *Mod6* after correction; see text).

### Analysis and interpretation of data

Substantial statistical consulting and numerous very helpful discussions came from B.H. and J.Da. The initial exploratory statistical analysis was done by K.S. (with consulting support from B.H.). B.H. provided an important independent statistical analysis that was instrumental in prioritizing questions while designing the final statistical analysis. L.L. independently developed and implemented the final statistical analysis performed by over 12K lines of R source code available as Supplemental Material with the input files used. L.L. collated the output with comments into a single PDF denoted as Supplemental Statistical Analysis in the text. B.H. also contributed towards various practical aspects of exploring the data and helped defining the approach that turned into the “linearization” of *CZT* discussed in this study; B.H.’s regular practical and statistical input has been extremely helpful for K.S. and L.L. by providing innumerable valuable pointers that helped navigate many complex decisions about how to best analyze the data presented. B.H. thereby substantially shaped the work and increased the overall statistical realism and rigor of this study. K.S. designed and implemented the workflow for increasingly refining the *PeakValleyTables*.

### Drafting of manuscript, revising content

K.S. wrote the first draft of the whole paper, created Figures 1-6+9, substantially refined many sections (Introduction, all sections directly discussing biological parts, Materials and Methods, Results, Discussion) and provided detailed feedback on most parts written or revised by L.L. In addition, K.S. helped adjust the overall paper structure and figure representations based on discussions with L.L. K.S. created several items of Supplemental Material, summarizing various results in tables, and worked with L.L. to streamline the overall presentation of results in order to reduce complexity for readers.

L.L. wrote the sections describing the data model of FlyClockbase (including its Supplemental Materials and substantial revisions to the FlyClockbase overview Fig. 2). L.L. greatly expanded K.S.’s texts on statistics and modeling, as used in this study (robust statistics, see R script, Supplemental Statistical Materials; parameter estimation for solving inverse problems). LL substantially edited the presentation of the Results and Discussion, and added Figures 7,8,10,11, and Tables 2,8. L.L. motivated and greatly improved the error analysis within the limitations of this study. L.L. wrote all sections on biological model curators, and computational aspects that build links to VBIRs, TabFS, Evolvix, reproducibility, logic, complier construction, the ‘DISCOVARCY’ documentation style, and broader links to biological modeling and evolutionary systems biology. L.L. provided biological consulting thought-out this project.

## 1. FlyClockbase design processes in detail

An overview of biological and data integrity aspects of FlyClockbase, as well as a description of relevant data types and their units is given in the Models section of the main text. Here we present practical ‘behind the scenes’ aspects of information management work. Although information management work is not “biological”, it is of substantial importance for the usability, reproducibility, and maintainability of FlyClockbase. Our paper stands by itself without these details. However, given the importance of reproducibility in computational biology; the enormous struggle to design systems that facilitate a desirable level of reproducibility over longer periods of time; as well as our interest in this topic, we decided to take the opportunity to reflect on some of the choices we made, and our subsequent experiences.

### FlyClockbase development is intertwined with Evolvix

Many aspects of FlyClockbase are linked with our work on developing reliable general-purpose programming capabilities for Evolvix. Evolvix is a modeling language that aims to facilitate accurate modeling for investigating biological systems and to provide long-term backwards compatibility for computational biologists to build on each other’s work in a more direct way (reducing time on re-implementing models). A prototype of Evolvix is available for download at (http://evolvix.org). The prototype download efficiently simulates pure mass-action models described in a declarative programming paradigm, and efficiently collects the most important time series data points during a specified simulation run (thwarting the origin of potentially large simulation results datasets before they become difficult to handle). This Evolvix prototype has been used for simulating models in research (Ehlert and Loewe 2014). Our practical experiences with this prototype in the context of a more complex modeling study (Ehlert and Loewe 2014) demonstrated the need for adding general-purpose programming capabilities to further simplify the construction of many modeling scenarios (debugging across several different languages as done currently is not an efficient use of research time). To overcome these challenges, we are developing an extension for Evolvix that implements a general-purpose programming language, designed to simplify general programming as much as possible, in order to stay true to the original vision of Evolvix, i.e. to make accurate modeling easier. This effort turned out to be highly unusual and required us to develop a new approach to programming language design, which we described elsewhere as Flipped Programing Language Design Approach (Loewe *et al*. 2017). Briefly, this approach ensures that unexperienced users of Evolvix can provide useful input in how the language is designed in order to ensure overall simplicity is maximized too, while experts work to maximize expressivity.

### FlyClockbase and its challenges to programming languages

Many real-world modeling problems in biology share characteristics that simultaneously characterize corresponding general-purpose programming challenges. The biggest computational obstacles to efficient computation in biology might be in its unique mix of diversity, complexity, uncertainty and vastness. In combination, these efficiently frustrate many computational abstractions that work well for less intense combinations of these complicating traits. A particular challenge is often the unpredictability and subtleness of such problems and exceptions; thus, there is no substitute for covering ground “on the ground” by walking detail by detail through specific biological examples while paying particular attention to cognitive dissonances that indicate a poor fit between the biological realm and the realm of current programming approaches. When certain types of programming bugs are eventually observed repeatedly, a programming ecologist might take solace in the observation that continued sampling in complex ecosystems will eventually yield a finite count for the number of types in a species richness problem (Dopfer *et al*. 2008; Zhang and Stern 2009; Magnussen *et al*. 2010), even if that number is very large as in the case of bugs (Grove and Stork 2000; Hamilton *et al*. 2010). FlyClockbase has provided us with the outstanding opportunity to observe at close range many instances that frustrate the efficient use of biological observations for computational purposes. Taken alone these issues are rarely large or complicated, but combined can be challenging (“glass of water” vs “tsunami”). This is a characteristic of scattered big data that often become relevant only in much larger datasets (to accumulate enough exceptions for triggering complexity problems). The data collected for FlyClockbase presented numerous challenges regarding the diversity and uncertainty of time series data. These challenges of FlyClockbase have inspired various important requirements for Evolvix development, aiming to simplify such work in the future (see Discussion of main text for some examples, but reporting most results would be beyond the scope of this study).

### Evolvix development follows a unique approach

The unique programming challenges of biology have inspired the rather unconventional Flipped Programming Language Design Approach for developing Evolvix (Loewe *et al*. 2017). This design process for Evolvix is heavily front-loaded and emphasizes the coherent, user-friendly integration of functionally mature sets of features that are chosen with a view to long-term stability. The aim of these priorities is to reduce idiosyncrasies and confusion for users, as well as inessential complexity and costly busy-work for developers, especially on the long term. The slightly higher short-term development efforts required for facilitating the necessary review processes are negligible in comparison to the benefit of developing Evolvix in a way that generates a long-term backwards-compatible language.

### Long-term goals of Evolvix

Evolvix aims to provide general-purpose programming capabilities designed by biologists for biologists to help with facing the complexities of biology. Given that inessential complexity is at the heart of many inefficiencies and problems in computational biology, it is mandatory for Evolvix to find the most efficient way of cutting as much programming complexity as possible. Efficiency will increase the ease of handling additional biological complexity. Architects of languages or compiler designs can only consciously address problems that they have noticed or that have been brought to their attention. Once such problems are openly visible, it is much easier to propose solutions. Once a solution has been proposed, it is much easier to critique it. The Flipped Programming Language Design approach used by Evolvix facilitates a communication line between a language architect that can still change the design of a language, and potential future users, who would otherwise just keep to themselves the many reasons that motivate them to classify a programing language as too complicated for them. Evolvix aims to break that cycle, where (otherwise clever) users prefer to not acquire computational skills that are well within their reach, simply because they are confused by some language-specific idiosyncrasy. To get passed this barrier to semantic reproducibility requires work towards debugging what has been termed the Code2Brain Interface (Loewe *et al*. 2017).

### How FlyClockbase helped Evolvix

In programming language design, as elsewhere, problems that remain hidden rarely go away and accidentally solving them is unlikely. We have used this perspective to turn these problems into a treasure trove of inspiration for designing innovative Evolvix programming language features. Working through the fly clock time series data that we integrated into FlyClockbase has presented us with a rich set of subtleties, semi-regularities, exceptions, uncertainties, contradictions, and other imperfections of real-world biological data. The decision to face these difficulties has enabled the development of designs for general-purpose programming language features that fit biological research problems much more naturally than solutions developed in ‘non-biological’ general-purpose programming languages (e.g. BEST Names for handling synonyms as described elsewhere (Loewe *et al*. 2017)). Many of the general results from our substantial engagement in this trans-disciplinary process remain beyond the scope of this study. However, the study that introduces FlyClockbase is the most natural place for presenting some important aspects of the process that helped us develop key concepts for representing the confusing complexity of biological data that we encountered in our work with FlyClockbase. An overview is described in Figure 11 of the main text and its respective comments. We have included illustrative examples of insight gained from this process below and in the Discussion.

### How Evolvix helped FlyClockbase

Conversely, the intertwined development process has motivated the adoption by FlyClockbase of numerous approaches originally developed to meet Evolvix requirements. An unduly short list of examples include the BEST Names concept, (Loewe and Keel 2014; Loewe *et al*. 2016), the Project Organization Stabilizing Tool (POST) system (Loewe *et al*. 2017), the insights we used to assign a special role to standard filesystems for storing data (which eventually stimulated the development of TabFS as described in the main text Discussion as the storage structure underpinning FlyClockbase), BioBinaries, and the stabilizing versioning system briefly described next.

### Stabilizing Versioning

To facilitate development, versioned variants of FlyClockbase are numbered according to the stabilizing versioning system developed for Evolvix and described as part of the “*StabilizingZone*” of the POST system (see online material of Loewe *et al*. (2017)). The term “version variants” is meant to highlights the diversity of types of variants that might change an instance of FlyClockbase in the stabilizing versioning system that it uses. For example, let us consider the version variants of this document when initially submitted to reviewers, and at the moment of initial publication as peer-reviewed study.

#### QQv1

The version variant associated with this submission is QQv1. This is the Brief Evolvix way of pointing to the meaning of the *StabilityCode* for “*QualityQuest*” and indicate that a substantial milestone has been reached that is known in *QQ* as *Version 1* and is now waiting for evaluation by various types of reviewers (including usability-reviewers and subject-matter expert reviewers for all relevant disciplines). There might be many or few, small or big, new versions, releases, and/or patches on the QQ level. How many depends on the number necessary for incorporating the feedback that is necessary for taking the next (more public) step in a responsible manner.

#### RRv1

The version variant RRv1 is associated with the first reviewed release that is intended for some productive use (albeit without particular stability guarantees yet). This is the Brief Evolvix way of saying “*ReviewedRelease*” and indicate that someone with the authority to release this variant has reviewed all pertinent issues and deemed the overall maturity to be sufficient for public release. For this study, this point will be reached, when FlyClockbase accompanies the final publication of this study in the form of a corresponding public release of FlyClockbase on Github.

#### Why stabilizing versioning?

This versioned variant numbering system was selected to encourage responsible steps towards enabling long-term backwards compatibility for FlyClockbase and to enable it to be a ***V**BIR*s, i.e. a truly *Versioned* Biological Information Resource (vs being only a BIRs, where “Current” is the only well-defined state or versioning is incomplete). Stabilizing versioning alleviates the tension between repeated rounds of rigorous review as required for long-term stability and the flexibility to quickly experiment with risky ideas as required for innovation. Some of these ideas have to be tried out before it is possible to reasonably decide what to do with them (many of them should never appear in a system aiming for stability). The concept of a *StablizingZone* has been developed to alleviate a similar tension in Evolvix. It builds on the *StabilityCodes* that are part of the POST system and are shown in Table P1.

**TABLE P1.**
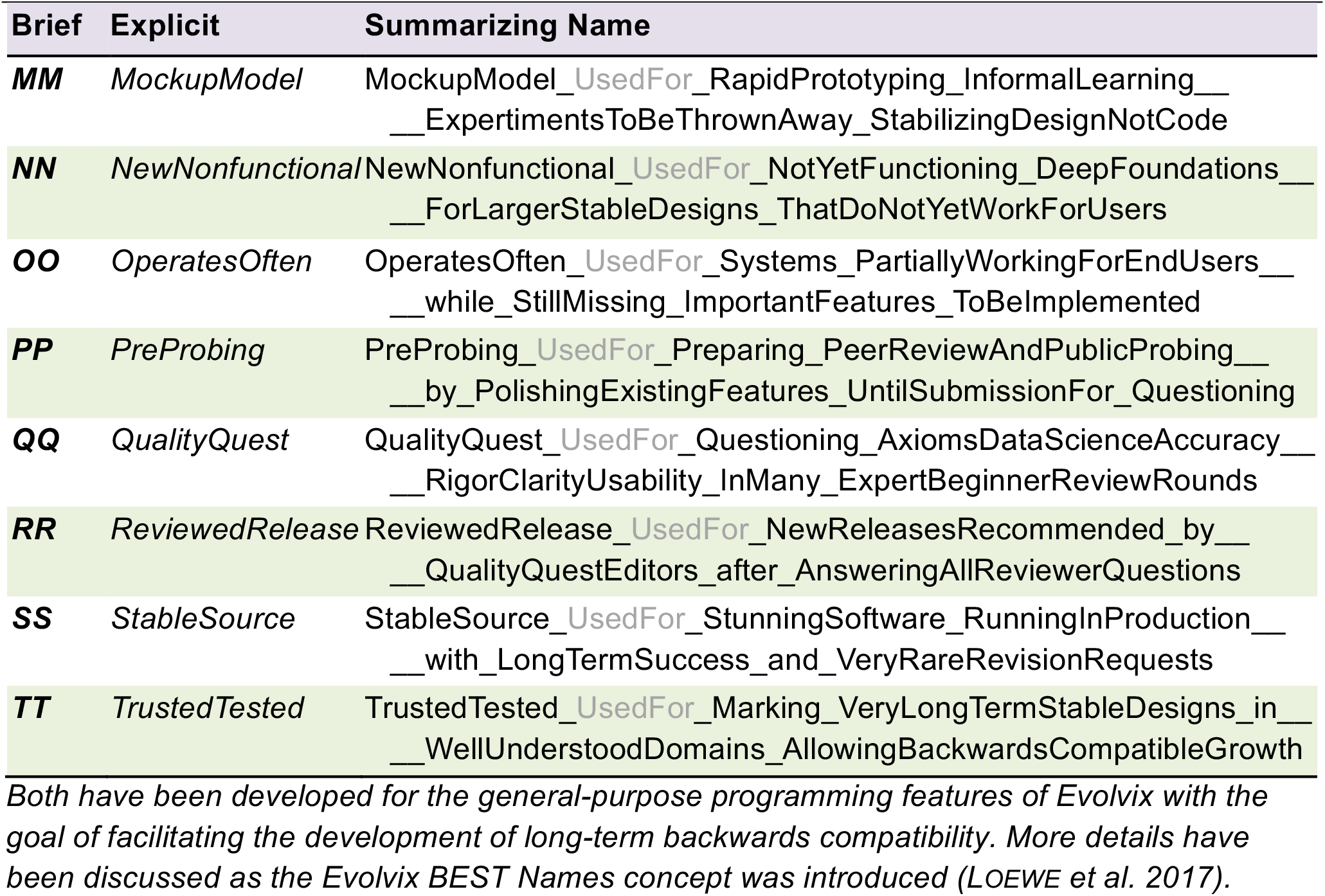
The *StablizingZone* of the *Project Organization Stabilizing Tool* (POST) system is defined by the *StabilityCodes MM* to *TT*, presented by their BEST Names.

#### The role of TabFS in the stability of FlyClockbase

FlyClockbase depends on the stability of its underpinning storage infrastructure. As argued below, we decided against traditional databases, because the constant stream of upgrades easily imposes prohibitive burdens of IT administration work on those biologists who are likely to initiate and maintain *VBIR*s. To resolve this, FlyClockbase development has also been intertwined with the development of what we call ‘TabFS’, an extremely thin and transparent file system that sits on top of a standard file system. This study is not the place to appropriately describe and define TabFS; suffice to say it is being designed to have the abilities of a fully functional file system with additional features that simplify some work in computational biology. These features are combined with a priority on radical openness allowing the local user in the host operating system to change anything and everything (e.g. this allows a biological model curator to easily change the type system of a *VBIR*). These desirables of *TabFS* come at the price of having to deal with inconsistent states that are more likely to be caused by users in *TabFS* than in other file systems. Resolving this will require additional checks and redundancies to enable recovery of consistency when users generate errors. Thus, the biggest trade-off in this setup is a slower performance. However, a fundamentally important and conscious design decision for *TabFS* is to trade short-term performance for increased long-term durability. Practically, *TabFS* stores everything in tab-delimited tables of text stored in the folders of a standard file system. It can be thought of as an organized equivalent of Comma-Separated-Value (CSV) files, albeit with strict formal rules that enable compiler checks.

#### How does stabilizing versioning benefit FlyClockbase?

The decision to buy into the stability of major standard file systems does not mean that problems of storage incompatibility cannot affect FlyClockbase. To efficiently organize FlyClockbase we need to abstract recurring themes, and these abstractions need to be developed, which takes time. FlyClockbase bundles these storage abstractions into TabFS; hence any changes in TabFS that are not backward compatible will threaten the data stored in FlyClockbase if not handled appropriately. This is where stabilizing versioning is expected to be extremely helpful: it indicates to users how reliable a potential upgrade is likely to be. It also indicates to developers and reviewers shared expectations about the quality of the code they write or read. Thus, proper versioning is essential for stability of FlyClockbase as a VBIR. Reporting more on our progress towards this goal is beyond the scope of this study.

#### Types in FlyClockbase

In addition to using TabFS, FlyClockbase has also been developing its own type system and content that requires coordinated changes and can hence benefit from versioning.

#### Test case for Evolvix

Without detracting from its biological focus, FlyClockbase provides an important real-world testing ground for developing approaches to enable long-term backwards compatibility in Evolvix (Loewe *et al*. 2017) serving longer-term research goals in Evolutionary Systems Biology (Loewe 2016).

### Why not a conventional database?

FlyClockbase is designed at a time when data science is of growing importance for biomedical research (see (NIH *et al*. 2012)). We have considered the many virtues of diverse well-developed database management systems (from SQL to NoSQL to NewSQL). These guarantee consistency of data, simplify complex searches via special query languages, and increase speed of access for growing amounts of data that is appropriately organized (e.g. (Jurney 2013; Mahmood 2016)).

Unfortunately, these advantages usually come at a price that includes the hidden or deferred costs of:

i. becoming increasingly dependent on a specific stack of software that is often growing in complexity,
ii. complicating installation or transport across platforms,
iii. becoming too difficult for non-specialists to tentatively add new data and datatypes,
iv. complicating the migration from an old (often rigid) database schema to one with new features (and complexity, but usually equally fragile).

Combined with poor documentation and sloppy naming, such problems can easily degrade the semantic reproducibility (Loewe *et al*. 2017) and, hence, the long-term usability of any data collection that builds on top of such tools. The Introduction of the main text discusses more aspects. As a result, pre-clinical biomedical research in the US has been estimated to invest about $7Bn/yr in studies that were deemed to contain irreproducible data analysis or reporting (Freedman *et al*. 2015). Such an environment is hardly conducive to efficient biology, computational or otherwise; as a consequence, the critical importance for reproducibility of research results has been recognized across many disciplines (Donoho 2009; Goecks *et al*. 2010; Karr *et al*. 2012; NIH *et al*. 2012; Stodden *et al*. 2014; James *et al*. 2015; Karr *et al*. 2015a; Karr *et al*. 2015b; Kenall *et al*. 2015; Poldrack and Poline 2015; Stodden 2015; Loewe 2016; Loewe *et al*. 2017). As argued in the Discussion of the main text, simple text files have many desirable features. We decided to carefully build on those.

### Logic challenges posed by missing data

Another notorious difficulty in many “standard” databases is how to indicate the absence of particular data, which can be interpreted as different types of zero. The difference between “not available”, “not applicable” and various other types of “null” is widely recognized (Zaniolo 1984; Candan *et al*. 1997; De Tre *et al*. 2004; Waraporn and Porkaew 2008; Bosc and Pivert 2010; Hernich *et al*. 2011; Thalheim and Schewe 2011; Hartmann and Link 2012; Lifschitz *et al*. 2012; Martinez *et al*. 2013; Mirza 2015), yet standard databases struggle to provide automated support beyond an unspecified, potentially ambiguous ‘*NA*’ (may stand for *NotAvailable*, *NotApplicable*, *NotAllowable*, …). Further, they build on the Closed World Assumption (that all relevant data is always fully specified) and aim to represent everything in the stark true-false dichotomy of Boolean logic. Current research in logic has provided many examples for the crippling effects of such black-white limitations when confronted with a more nuanced world of color (Smith 2008). These problems surface very quickly, when attempting to reconcile the crisp true-false dichotomies of a world of Boolean variables with biological observations with a certainty that is less than clear-cut. To quickly allow for the identification of such biologically important cases without losing the advantages of the crisp semantics of Boolean logic, the following “OKScale” of four alternative states for a value of the type ‘BioBinary’ has been designed for Evolvix (see p.16, online material of (Loewe *et al*. 2017)). FlyClockbase is adopting the *BioBinary* type to facilitate clearly marking the states described in Table P2.

**TABLE P2.**
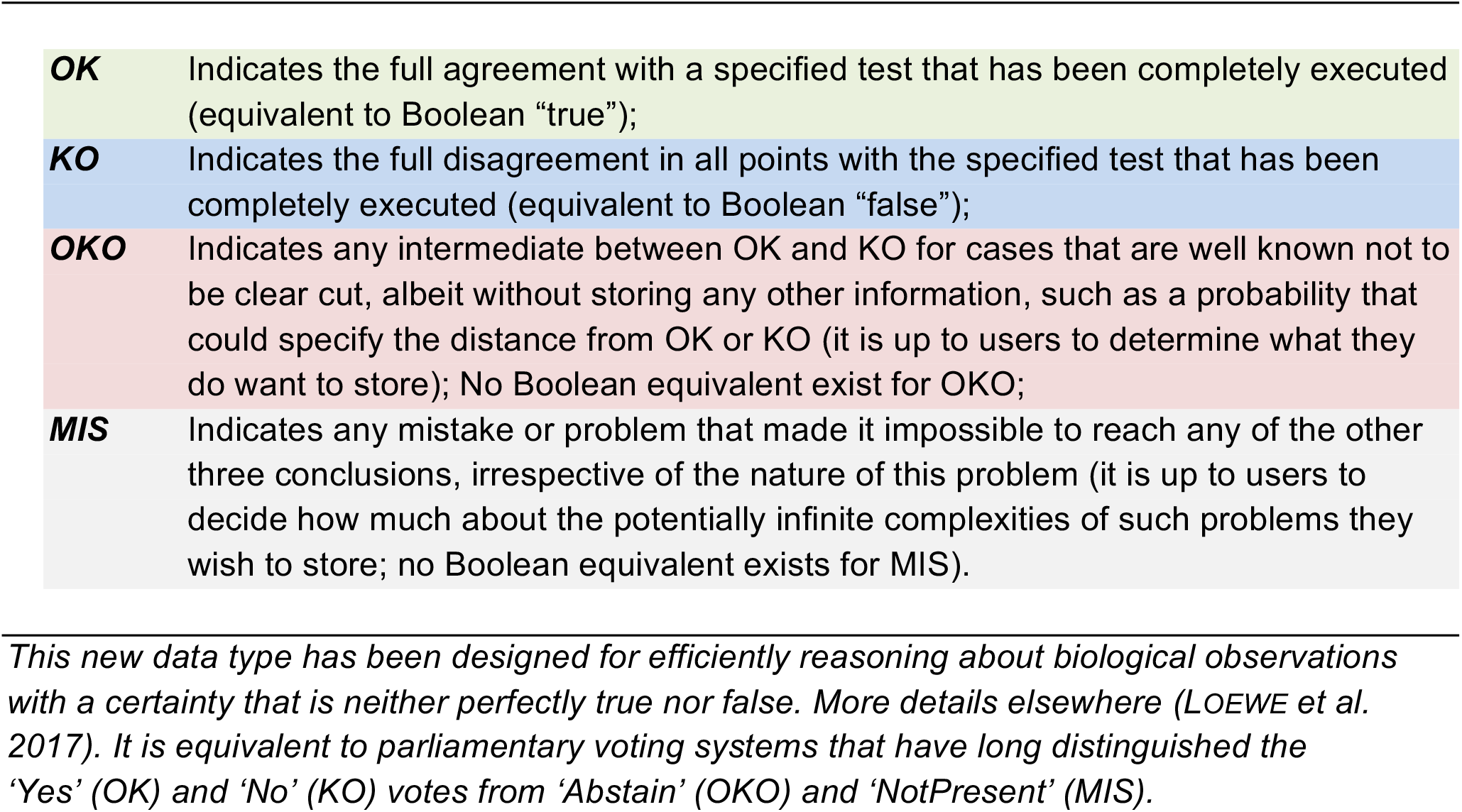
The *BioBinary* data type for cases in biology that are less than clear-cut.

The basic features of the *BioBinary* do not eliminate the need to find a way of representing the infinitely many more detailed types of OKO, KO, and MIS, which might be used to represent details about the infinitely many stages of incompleteness, zero, null-hypotheses, or potential mistakes and contradictions that could be encountered when analyzing biological data. It is extremely difficult to anticipate which of the more nuanced types of information beyond these four will be encountered while actually analyzing biological observations. Maximizing the flexibility for describing such exceptions, while providing useful guidelines to educate FlyClockbase users about important subtleties, improves the quality of the reported data and reduces the frequency of misleadingly strong statements that overstate their claim. Such statement can easily be generated by if the logic formalism of a system does not allow recording a weaker statement. Our most important goal is to have a logical formalism that is neither overly restrictive (which would prohibit the representation of certain observations), nor as unstructured and flexible as a blank page (which would make it very difficult to conduct any useful logic analyses). We found it helpful to work with lists of controlled vocabularies and only to add freeform comments for special cases, which could eventually add to controlled vocabulary if repetitive. The decisions about which columns are controlled, which are free, which entries make it into a controlled vocabulary list, and when a free-form comment will be extracted in order to populate a new column, and many more such decisions strongly depend on the subject matter and the content of a VBIRs. We therefore decided that biological model curators would need the full ability to change any of the types in a VBIRs as they see fit to represent the observations they curate. Since VBIRs are Versioned, all these decisions can and should be properly reviewed to improve consistency.

### Permissions, backups and the reliability of data storage

Beginners and experts, all have full administrative (“root”) access to almost every aspect in FlyClockbase. The very open approach of FlyClockbase raises the question of how to guarantee the integrity of its use. While in theory a user can compromise FlyClockbase quickly, the chances of such an event with lasting consequences are much smaller than these unrestricted settings might suggest. When viewed in context:

i. few users of FlyClockbase *aim* to compromise their installation; moderate hiding of critical infrastructure or clear indications warning about the potential of breaking sensitive parts is usually sufficient to prevent mistakes; the latest working release of FlyClockbase is never in danger if a release manager keeps backups;
ii. distributed development of FlyClockbase poses the same challenges as all distributed information processing, namely, the merge/cache-invalidation/naming problem. It can be solved in principle by distributed version control systems (like Git), or by using cloud synchronized folders in combination with strictly following a version variant naming scheme to prevent loss of data (see Fig. 1BC in Loewe *et al*. 2017). We tested both approaches (Git and cloud-style automatic synchronization). We found both to be less inviting than they could be if a set of reliable scripts were to automate repetitive tasks in a way that reliably excludes loss of data (and without adding to the cognitive load of the user).

While it may seem that widely known database technologies are free from these problems, they in-turn come with independent problems of their own. For example, FlyClockbase may give ordinary users too much administrative power. Many other databases resolve this problem by locking down access. Yet, in turn these databases require special expertise or permissions for exploring new avenues of recording a richer spectrum of biologically relevant difficulties with contradictory scattered big data. Even experienced database architects may use natural language for the very early initial stages of database development. We aim to give model curators the ability to do exactly this ‘*in situ*’.

#### On some challenges in logic work

Capturing details important for describing biological observations in a consistent way can be challenging research in logic formalisms, because of the vagueness of some biological data (Smith 2008). Most biological model curators are not likely to have the training in formal methods to define the new formalisms that remain at the end of this research. As argued above and in Figure 11 (main text), curators can still contribute a lot by providing examples and detailed expertise on how to interpret their various challenging aspects, which is impossible for non-biologists to do. To facilitate the practical work necessary for enabling this type of discussion, a number of smaller problems need to be resolved:

- *Form of recording*. Any persistent record of observations in logic needs to encode the observations of principles or bugs in some formal way that can also be read by other persons. In a world of infinite resources, it does not matter how these observations are encoded, as long as their semantics is completely reproducible (Loewe *et al*. 2017). However, for most biologists engaged with model curation, patience with logic formalisms is a limited resource. Approaches used in practice for communicating logic problems ought to be as simple to use as possible. A blank page or an arbitrary text description work well, written or spoken; it is the task of a person with more understanding in formalisms to translate such fragmented insights into a more coherent formal picture. Additional burdens, such as transforming descriptions of logic problems into a particular logic formalism, are probably counterproductive. Many curators might be tempted to indicate that they have nothing to report, simply to avoid the difficult task to re-encode their observations, e.g. in a relational database formalism. While certainly possible, it does not mean that this is advisable; in principle, these problems could also be expressed in the form of Gödel numbers (GÖDEL 1931), but this would only add huge amounts of inessential complexity. Thus, the simplest possible storage medium is probably best.
- *Durability of recording*. For the sake of argument, let us assume that there was a very straightforward way of recording all such statements in a database. Then such work can become unnecessarily complicated, if the database system requires frequent complex updates that are triggered by external database developers aiming to deliver ongoing improvements like new features, security, performance or more. If such upgrading becomes excessive, paper and pencil might become increasingly attractive.
- *Size and complexity*. Documenting progress in debugging or developing logic does not usually generate large amounts of data, but the few cases that are there are important and complicated. Hence, an uncomplicated, distraction free storage medium is desirable.
- *Speed of copying*. For efficient communication, it is desirable to find a storage medium that is easily copied through communication channels. This is where non-electronic approaches have their greatest drawbacks.

#### Trade-offs

To solve these challenges in a practical way, let us consider the following advantages and disadvantages of paper, file systems and databases.

- *Paper*, for example, does not suddenly refuse to operate as expected because it needs an upgrade – a plus for stability. Its durability is legendary and generally measured on the scale of centuries. However, the speed of writing, reading, and copying is slow and error prone.
- *Dedicated databases*, relational or otherwise, when designed for fast retrieval of certain types of answers are hard to beat in speed for that particular application. Yet that speed will drop to zero, if the latest essential upgrade is not applied and all data in the system becomes inaccessible (see tombstone example below).
- *Filesystems* in contrast, offer the best of both worlds: almost the reliability of paper (assuming that there will always be electricity and hardware that enables access to the file system and perpetuation of backups); almost the flexibility of paper (assuming corresponding types of files); almost the speed of databases (assuming the data is organized so that the speed of reading or writing approaches the limit supported by the hardware). The big question for file systems is whether a way of organization can be found that keeps all important data appropriately organized.

##### Long-term stability in VBIRs data structures will enable new biology

*We suggest that a compiler could help curators to efficiently maintain long-term stable VBIRs in consistent formal states so that these VBIRs can be used as foundations, which are solid enough to build on them for the long term. This will eventually make it feasible to construct more advanced VBIRs on top of more basic ones*.

*If foundational VBIRs describe causal genotypic or environmental information and more advanced VBIRs describe more consequential molecular, cellular, physiological, or other phenotypic information, then VBIRs enable the implementation of a full fitness-causality network that maps the genotypes and environments of an organism to its phenotypes in a transparent mechanistic way that connects well to the latest updates of all data that is available – if all relevant data and data structures are well curated and appropriately versioned for stability*.

### Tombstone example

Here we illustrate with an example the very real long-term danger to biological research posed by database technology that is not long-term backwards compatible. The TIGR Gene Indices were first published at http://www.tigr.org/tdb/tgi (Quackenbush *et al*. 2000). After their introduction, the TIGR Gene Index databases quickly became a well-known tool for biological discovery (Lee and Quackenbush 2003; Pertea *et al*. 2003; Lee *et al*. 2005). They are now no longer available online as documented on their tombstone:

http://compbio.dfci.harvard.edu/tgi/

In case the tombstone itself disappears eventually, here we paraphrase its 2017 report of some vital statistics about TGI:

*Supported by NIH, DOE and NSF 1998-2010, the relevant TGI papers were cited >2000 times. When the tombstone was written the TGI website still received >7 million hits per year (assuming the actual number on the tombstone was a typo). When funding ended in 2010, the team continued to maintain the website, but the hardware and software required behind the scenes began to fail. Effective July 15, 2014 operations had to be suspended, because there were not sufficient funds to maintain it properly. The software powering TGI (DFCI Gene Indices Software Tools) and the data sets it used was ‘fossilized’ to*

> *ftp://occams.dfci.harvard.edu/pub/bio/tgi/software/*
>
> *ftp://occams.dfci.harvard.edu/pub/bio/tgi/data/*.

It is not up to us to comment on TGI’s science or its funding history. Neither is relevant to our main point: extinction is a real risk for *VBIR*s and fossilization to some archive is not a real life-saver. The haphazard nature of funding for biological information repositories is well known and a significant source of concern (Ember *et al*. 2013). Less obvious is the impact of a stable VBIRs compiler for TGI. Imagine the software behind TGI would use appropriate abstractions and thus not fail. Imagine it could continue to operate reliably on different hardware, including that of users. Imagine the software would be long-term backwards compatible. Imagine it could help many biologists to contribute to curation of TGI. Imagine a whole community would annotate, improve, deprecate, or otherwise edit various aspects of TGI in order to preserve its benefits or point to improved successor tools. Imagine other biological research codes had built on long-term stable parts of TGI and could all continue to operate simply by copying TGI to a local hard drive. Would that make a difference?

## 2. Human error analyses: approaches, challenges, efficiencies

Problems with tracing the identity, availability, accuracy, precision, and reliability of data have been the topic of numerous investigations and recently received renewed attention in the context of assessing potential uses of scattered biological data sources and big data (e.g. see (Reason and Mycielska 1982; Reason 1990; Reason and Hobbs 2003; Goldston 2008; Gitelman 2013; Grimes *et al*. 2013; Mccallum 2013; Reason 2013; Blankenberg *et al*. 2014; Reason 2015)).

FlyClockbase presented us with an excellent opportunity to explore numerous issues related to the quality, accuracy, and reliability of complex collections of information that aim to integrate data that can be incomplete, uncertain, contradictory, erroneous and scattered across a wide range of sources. Much expertise in biology currently exists in such less-than-well-defined states that can be difficult or impossible to process computationally. Thus, we expect that some of the lessons we learned while compiling FlyClockbase become increasingly important as biologists expand their use of computational resources throughout their work. While it is beyond the scope of this study to appropriately review all corresponding challenges and potential solutions we encountered, a few observations of common challenges of general interest are reviewed next.

### Finding capable curators

Compiling nontrivial resources of biological information requires a substantial and sustained effort from researchers who bring a certain set of skills. They need to (i) possess the background expertise necessary to understand the importance of the information they compile, and (ii) be motivated to collect information from various scattered representations into a single, less idiosyncratic representation that constitutes the new resource. New resource creation, as listed above, usually requires more effort than experimental or computational biologists are willing to dedicate. This might require researchers with a special interest in high-quality information collection regarding the respective system. Like airplane pilots, curators will need training in appropriate error handling.

### Errors are inevitable

A broad body of research has shown that human errors in any non-trivial data processing are inevitable and that error rates increase with the complexity of a task (Reason 1990; Reason and Hobbs 2003; Reason 2013; Reason 2015; Panko 2016). This is true in particular for the use of spreadsheet software (Panko 1998; Panko and Sprague 1998; Panko and Aurigemma 2010), for which rates for various types of error have been measured. Our own results confirm that some errors (e.g. simpler data entry) occur with lower rates than other more complex errors, that stem from occasionally forgetting non-obvious steps in a complicated procedure (see Table 4). We can confirm previous findings, showing that even the most motivated researchers with substantial training and background expertise will inevitably err when transferring information in complicated scenarios where many complex details could go awry (see also (Lesgold *et al*. 1988; Galletta *et al*. 1993; Galletta *et al*. 1997; Panko and Sprague 1998)).

### Errors are hard to detect for humans

Proofreading software is well known for not catching all spelling errors. It is also well known, that not all humans catch all spelling errors. Yet spelling errors are the simplest errors. All others are more difficult to detect in comparison. Errors in logic are particularly difficult; however, the worst category of errors are errors of omission (Panko 2016).

Given the various complex operations required for integrating and analyzing data with FlyClockbase, FlyClockbase has competitive error rates, as found when re-visiting parts of the database (see Table 5). However, manually extracting even simple time series traits (e.g. peak-valley times) has proven more error prone than we expected. The complexity of traits can rise substantially as increasing numbers of important conditions and special cases need to be observed for extracting a valid trait. Eventually complex decisions, when integrating less-than-perfectly fitting new observations into biological information resources, might easily generate very complex error scenarios. This is mainly because such work requires many decisions about which types in the data and in a *VBIR* are functionally equivalent and when effective differences are large enough to justify the introduction of new categories. The complexity of such offline decisions could rival the complexity (after preparation and training) of online decisions in mission-critical scenarios that have been analyzed elsewhere, from cardiac surgery (Carthey *et al*. 2001; Reason 2005) to space flight (NASA *et al*. 2011; Hooey *et al*. 2014). In many of these scenarios, human failure is catastrophic; correspondingly sophisticated human reliability analysis methods have been developed to ensure that human error is exceedingly unlikely - if not preventable, in principle (Reason 1990; NASA *et al*. 2001-09-30; Reason and Hobbs 2003; NASA *et al*. 2006-07; NASA *et al*. 2011; Reason 2013; Reason 2015).

### Human error analysis and prevention methods are difficult to develop

Many of the methods that are appropriate for investigating and preventing human error in contexts such as space flight, nuclear power plants, or aviation (NASA *et al*. 2006-07; NASA *et al*. 2011) are much too complex and costly for applying them towards improving the reliability of biological information resources like FlyClockbase. This is true even if the complexity of problems generated by resources such as FlyClockbase rivals or exceeds the complexity of navigating a space ship.

The huge immediate and catastrophic danger and costs of errors in navigating space ships and running nuclear power stations make it obvious and easy to justify the large effort required for in-depth analyses of human errors. In contrast, adding data to *VBIRs* seems harmless; it certainly is if only the immediate impact is considered, and it probably also is on the long term. However, in biomedical research there is a small chance that errors in *VBIRs* are not as harmless as they seem if they remain uncorrected. The following scenario is difficult to exclude categorically in bio-medical research. Imagine that some researcher integrates data into some medically relevant *VBIR* and makes an unnoticed error that remains in that *VBIR* for a long time and even replicates across repositories. If that error prevents the discovery an important new medical cure by leading a researcher to the wrong conclusions, then cost and risk calculations are no longer trivial.

It is not clear how to resolve this problem. If each *VBIR* produced its own error analyses, costs would quickly spiral out of control and few resources would remain for actual biological research.

Error analyses require a substantial level of biological expertise, yet are perceived as ‘tedious’ and ‘non-biological’ (by experimentalists) or ‘non-automatable’ (by computationally oriented biologists). Thus, recruiting experts for such error analyses can be exceedingly difficult or prohibitive, despite their importance for modern biology. The following potential ‘solutions’ are also likely to be counter-productive on the long term:

i. *Omitting error analyses*. While cost-effective during the initial stages of compiling a new resource, ignoring the potential for errors by not engaging in error analyses will result in a growing number of databases with questionable quality. Whenever databases have been analyzed, some rate of errors has been found (e.g. (Jones *et al*. 2007; Hutchins *et al*. 2010; Weil *et al*. 2015)). It should not usually require open letters or other drastic measures (e.g. (Drake *et al*. 2008; Allison *et al*. 2016)) to possibly get errors corrected. The current state of software tools makes post-publication changes very difficult and most databases do not offer an easy way to submit proposed changes for review and efficient inclusion by administrators. While this explains the reluctance to update the latest available information, it is not conducive to advancing the integrity of scientific data. If no resolution to this error crisis can be found on the long-term, the resulting loss of quality will lead to a loss of reliability, reproducibility, and ultimately to a loss of trust in ‘data-based’ results. These costs are not immediately obvious to the authors of a new resource, but they will eventually manifest somewhere in the scientific community, either as irreproducible results, misguided research directions, or unnecessary ‘confusion-tax’ for the users of a resource.
ii. *Automating database operations*. Some types of errors are greatly reduced by the use of automated database technology (e.g. leveraging SQL to keep indices up to date and reduce redundancy,). This is particularly effective for types of errors that occur in large numbers, thus making it easy to automate detection. *Heterogeneity*. However, the highly heterogeneous nature of many advanced biological information resources can easily generate as many potentially relevant exceptions as entries. Relevant exceptions not addressed by the built-in logic of a database management system are at high-risk of causing errors, since errors by omission (of a special condition) are particularly difficult to detect (e.g. see discussions of *BioBinary*). Such errors can be even more difficult to detect in automated systems, if they are not already detected automatically. Automation might tempt researchers to become over-reliant on computational results they no longer understand (especially if implemented by non-biological experts). The substantial computer science literature on compiler construction demonstrates that the use of automation via software does not protect against the innumerable types of errors that can be added to source code and can corrupt results (if not caught by a compiler; see (Parsons 1992; Aho 2007; Cooper 2012; Grune 2012)). This suggests the following interesting conclusion: *Automation trades some human errors for others*. For example, inconsistency errors that often occur in data that is entered manually are avoided by automation through programming; get programming can easily generate inconsistencies of its own that humans would not produce. Since about 2% - 5% of all lines of source code have been estimated to contain some errors (Panko 2016), it is far from obvious how automatically handling the many special cases in resources like FlyClockbase can make additional analyses of various human errors superfluous.
iii. *No retrospective meta-analyses*. Not engaging with data analyses that trigger the need for error analyses robs biology of important perspectives. This would be equivalent to a call for stopping such otherwise non-controversial biological research. The impact on integrative work and relevant modeling efforts could be devastating. Hence, this options is not attractive.

While currently no ideal solutions exist, raised awareness of the problem is likely to eventually contribute to the development of appropriate solutions. As argued above in the context of compiler design: seeing the problem in the first place is often the most difficult step. Maybe compilers could also help by automating many tedious aspects of these error analyses.

### Error analyses could be amortized across resources by compilers

As argued above, developing appropriate error analyses for a single *VBIR* is not feasible. However, our experience with developing FlyClockbase suggests that a substantial number of essential tasks are recurrent and of comparable complexity across the development of a groups of *VBIR*s. Examples include the need for handling missing data, inapplicable data, imprecise ranges (rather than precise values), heterogeneous method descriptions, comparisons of wild types to mutants, and many other aspects of biological interest. Thus:

> *The most efficient solution to improving the quality of VBIRs without exploding costs is to develop an automated compiler that can automatically test for all known and formally described cases of errors and supports a programming language that integrates biology expertise to simplify the description of test cases by biologists*.

Programmers frequently say that it is important to use the right tool for a given programming task, yet no such tool exists for the compilation of complex *biological* datasets. We will not repeat here the substantial number of reasons why such a language would be helpful and why current (non-biological) programming languages are insufficient (see Online Material and additional reasons discussed in (Loewe *et al*. 2017)).

### Efficiencies of scale

In the long term, it would be more cost efficient to construct a compiler that can read in all tables of FlyClockbase (or other *VBIR*s) and produce a report of all inconsistencies and errors that require human attention (as well as produce tables with all requested search results). Thus, certain types of errors would be detected automatically. Error detection modules and other solutions would be implemented only once and would simultaneously improve the reliability of all VBIRs construction efforts.

For those who wonder why we did not directly implement such a compiler, we would like to point out that - as so often is the case in computational biology - the overall time for obtaining a given biological result by manual ‘compilation’ is usually much shorter than the time it would take to implement a corresponding compiler. This certainly was the case here. For example, compiling the time series data of wildtype flies and checking the integrity of PER and TIM data to reasonable reliability was completed *much* faster manually, then if we had attempted to automate it by constructing a compiler for this. This cost calculation changes very quickly if large amounts of integration work are anticipated over long amounts of time in FlyClockbase or other *VBIR*s. There seems to be little debate over the need for curating biological data, especially as it keeps growing (Bourne and Mcentyre 2006; Burkhardt *et al*. 2006; Salimi and Vita 2006; Howe *et al*. 2008; Burge *et al*. 2012).

### Tool development and funding strategy development are connected tasks

Some development of tools beyond the scope of a single project already started, addressing the large amount of text mining done by biocurators (Wei *et al*. 2013; Rak *et al*. 2014; Singhal *et al*. 2016). However, developing clear and successful funding strategies has remained a challenge (Ember *et al*. 2013; Orchard and Hermjakob 2015; Rodriguez-Esteban 2015; Reiser *et al*. 2016). The inefficiencies of present-day curation work in biology are substantial and might be sufficient to convince most funders to invest elsewhere. Our experiences have shown that many of these inefficiencies could be alleviated in principle by appropriately constructed tools. By today’s standards it was unimaginably laborious to sequence a few genes in 1977, or a single human genome in 2001 (Sanger *et al*. 1977a; Sanger *et al*. 1977b; Lander *et al*. 2001; Venter *et al*. 2001; Hayden 2014; Sheridan 2014; Telenti *et al*. 2016). Given the lack of appropriate tools, the lack of funding for corresponding work might be less surprising. Funding for sequencing human genomes in 1977 was negligible, if not $0.

## 3. DISCOVARCY Documentation Style Questions for Coders

The attitude of some programmers towards documentation can be summarized as:

> ***The source code is the documentation*.**

Indeed, the agile approach to software development tries to avoid documentation, which can be costly to write, and is constantly out of date for fast developing code bases unless it is maintained with a large amount of effort (e.g. see comments in (Henry 2013)). As a corollary, one might conclude that documenting scientific research code is a waste of time, since by definition such code is moving fast^1^. Unfortunately, such practice results in semantic irreproducibility (Loewe *et al*. 2017), which hampers research (Freedman *et al*. 2015). On the other extreme, again, it is neither reasonable nor efficient to demand that all research codes are refactored and documented excessively. For the most part, research codes are indeed only used very few times.

### Source code as electronic lab note book for in silico research

For example, consider the highly specialized *R*-script, which was written especially for this study and that might be seen as a ‘unique research application’. It is likely to serve only the purpose of re-running the analyses of our study, facilitating computational reproducibility of our study’s results. While writing that code, there was no mandate to use best practices for software engineering – unless it would help get the job done faster or more reliably. Hence, no refactoring for longer-term use was done for increasing the quality of the source code.

### Trade-offs in code writing

Good writing usually requires multiple rounds of revision and similarly, good code requires substantial refactoring. Depending on the occasion, this may be effort well spent, or time wasted. Our supplementary code for this study lives in this tension, and so do many other research computing codes written for specific unique analyses. How can such code be made more readable in order to help other researchers benefit from the work of the authors, and not waste their time on inessential complexities? This question is at the heart of semantic reproducibility (Loewe *et al*. 2017). Table D1 introduces the DISCOVARCY questions on documentation style. The acronym DISCOVARCY highlights typical problem areas that can sometimes easily be improved once awareness increases. The acronym is on purpose designed to illustrate the many transitions small and large that are between worst and best practice. Some of the distinguishing characteristics are easy to change while programming, but prohibitively costly later (e.g. choosing summarizing variable names). On other occasions, it is prohibitively costly to refactor a whole code base merely to make it easier to understand. Such cost-quality related tensions highlighted by DISCOVARCY are linked to a fundamental trade-off faced by every innovator aiming to produce features reproducible by others. The trade-off forces numerous choices between the following ideals:

- *Reproducibility* (much time for tests and documenting, little time for new features);
- *New features* (little time for tests and documenting, much time for new features).

Every scientist needs to navigate the contrast between these ideals that is particularly sharp in research, where the goal is to make innovative new discoveries, while enabling others also to reproduce these innovations. The same tensions exist for software developers and programmers, only that reproducibility comes with a human and a machine aspect. For machines, reproducibility is the repeated execution on machines; this links the diversity of machines included in this claim directly to the cost of reproducibility. The human aspect of reproducibility implies that a human being other than the developer can read and understand the code correctly, an ability defined by the quality of the Code2Brain interfaces involved (Loewe *et al*. 2017). As with machines, producing something that is more widely understood is usually more expensive. Much of the work needed for improving clarity and documentation of research codes could be delegated to a compiler that can easily translate between Brief Names preferred by developers and more verbose alternatives more easily understood by newcomers. The Evolvix BEST Names concept has been developed in order to facilitate such translation (Loewe and Keel 2014; Loewe 2016; Loewe *et al*. 2016; Loewe *et al*. 2017). Documenting code has a long history (Knuth and Levy 1994) and requires more attention than this study can offer here.

### DISCOVARCY in brief

The documentation style in Table D1 is based on the notion that awareness can allow researchers to exploit unique coding time opportunities for writing inexpensive imperfect comments that nevertheless greatly increase the real-world readability of their code. The DISCOVARCY acronym highlights strategies that allow code writers to easily reduce common challenges to code readers. Briefly,

> **Describing Design** beats *deduction*,
>
> **Info Included** beats *inference*,
>
> **Source Simplicity** beats *secrecy*,
>
> **Code Clarity** beats *complexity*,
>
> **Offline-Online Overview Offers** beat *online odysseys*,
>
> **Vetted Variables** beat *vagueness*,
>
> **Argued Axioms** beat *arbitrary assumptions*,
>
> **Relevant Restraints** beat *random restrictions*,
>
> **Collected Comments** beat *cancelled comments*, and
>
> **Your Yield** from using this code should not cost *years* of learning about it.

**TABLE D1:**
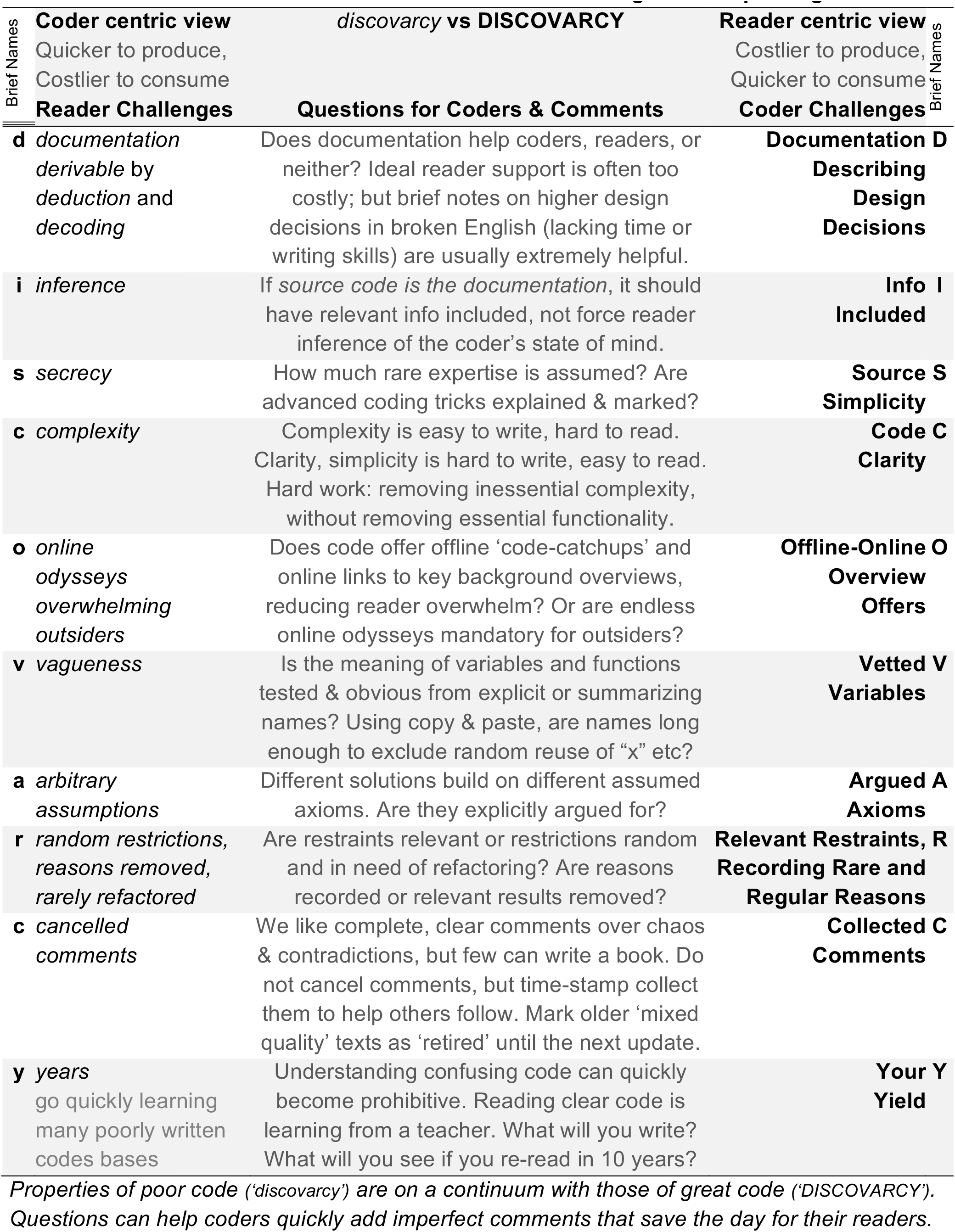
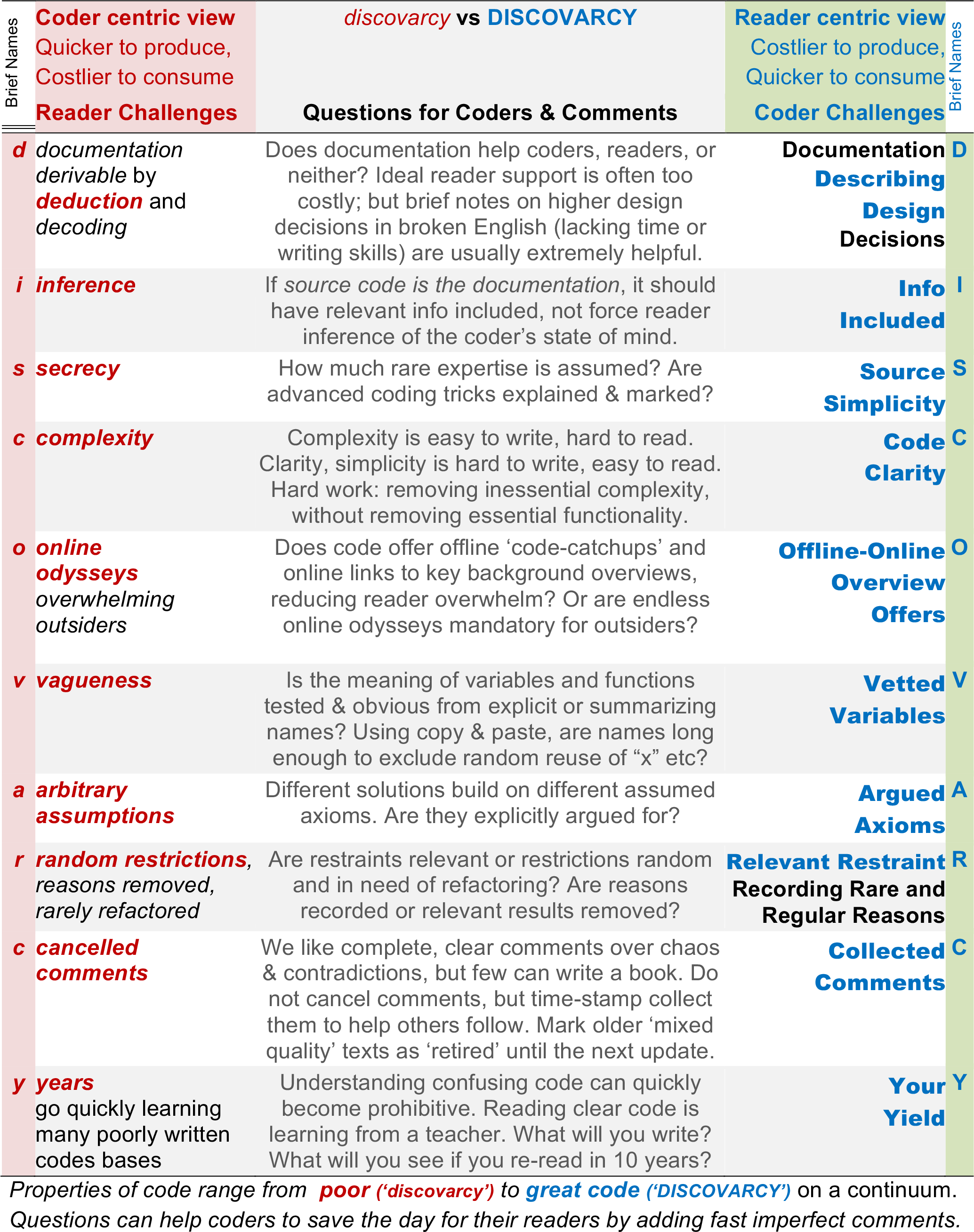
The **DISCOVARCY Documentation Style** raises awareness for causes that make source code **hard to read** and offers efficient strategies for **improving it**.

## 4. Example of an error a compiler could have caught by type checking: Effect of linearizing clock times on distributions of *Peaks* and *Valleys*

Please see Methods in the main text for motivating the linearization of times and for details on how it works. To illustrate the effect in the context of real data analysis, we selected types of mRNA and protein from FlyClockbase that are affected to varying degrees. Their observed peak and valley times are in Table S1, which compares their respective *ObsRaw* (non-linearized) and *ObsMod* (linearized) times.

### Recall Example Context

For observing the impact of linearizing time, we remember that the time measurements in *ObsRaw TimeSeries* belong to the time type *DailyZT*, or *DZT*. If this information about types is understood by a compiler, then the compiler will know that such time measurements are recorded in time units of decimalized *Hours* and that they are reset to 0*h* at the dawn of each day, which will reoccur every 24*h* and increase a separate day counter by +1. If we ignore the day counter as usually done in circadian biology research, then *DZT* is a cyclical time type, which simultaneously is also a cyclical real number type. In contrast, all *ObsMod TimeSeries* are measured in linear time without resets, as defined by *ContinuousZT, or CZT*. A compiler that understands this type information will know how to convert *CZT* to *DZT* and back (by adjusting the additional number of days that have occurred; see main text). If the compiler understands the assumptions required for computing valid summary statistics, it can check if those assumptions are met.

For example, using the well-known equations for inferring the mean of a Normal distribution requires that all values exist on a linear real number line, before attempting to calculate averages. Using values of a fundamentally different type (e.g. with the value ‘`MyText`’) would result in a compiler error (“`Cannot use ‘MyText’ when averaging`”). For time measurements of type *CZT*, the decision is easy, because the definition 0 < *CZT* ≤ infinity can never generate cyclical time. For DZT time types, the decision is not as easy, because definitions like 0 < *DZT* ≤ 24*h* do not imply the end of time once the 24*h* mark has passed. Flies, clocks and the rest of the world simply move on and will generate new data points. A sloppy *DZT* implementation will ignore the day-counter and act as if controlled by a time-loop – albeit with no effect on the linear time experienced by flies. The logical contradiction between the existence of a time-loop as assumed by such programs absence of such a time-loop in real life does not always matter. For example, if comparing only events from a *single* day, nothing can go wrong. However, some time series in FlyClockbase are longer than 24*h*, which can generate profound confusion, as explained next.

### Problem in principle

To illustrate the problem, let us consider an artificial dataset of *DZT* peak times for which a mean of 23*h* has been inferred for the first day and where the observed min and max has been 22.1*h* and 23.9*h*, respectively. Despite a circular number type, calculating a mean of 23*h* is appropriate; no data is included from another day; hence, all times are effectively linear. Let us assume that the next peak added occurs 90 minutes after 23*h*. If measured in *CZT*, this new peak would add 24.5*h* to an expected mean of 23*h*; the time remains linear and the mean remains valid. But expressed as *DZT*, this new peak would appear as a big outlier at 0.5*h* in simplistic comparisons with a mean of 23h (which implicitly drop a day). This type mismatch would misrepresent a peak of +1.5*h* after 23*h* as -22.5*h*. an erroneously large distance with the wrong sign. Such errors will bias inferred means in the wrong direction and greatly inflate inferred variances, possibly obscuring genuine biological signals. If undetected, these errors may substantially bias biological conclusions at unspecified cost to both reproducibility and biology.

### Problem in practice

Initially we were not aware of these logical subtleties in handling different types of time. Calculating averages and variances requires a linear scale, a condition that is *completely* met out-of-the-box for all *Peak* or *Valley* times of some core clock components reported in Table S1. In these cases, time-linearizing does not affect summary statistics, since all observations already belong to the same day (e.g. TIM Protein *Peak*, 18.41 ± 2.54*h*). However, if some observations belong to the previous or next day, a change in summary statistics can be observed (e.g. PER protein *Peak: Raw* 16.77 ± 5.48 to *Mod* 19.51 ± 3.40 *h SD*; see also the other average values in bold). To interpret SDs, it is instructive to consider random times of a day.

### Uninformative DZTs

As reported in the main text, *SD* > 6*h* is on the order of the *SD* of a uniform distribution across the day (deterministic 1h intervals from 0 to 24*h* result in *SD* ± 7.36*h* around 12*h* and drawing 1,000 corresponding uniformly distributed values does not fundamentally change the results (*SDs*: 6.81, 6.92, 7.02 *h*). Informal tests reducing the sample size to 20 presented a similar picture, albeit with substantially more noise (as we would expect for sample sizes comparable to the number of time series for a component in FlyClockbase).

### Detecting excessive noise

We can use the random *DZT* results to detect some cases, where a clock signal is known to exist, but measurement or data handling problems have obscured the signal dramatically (so it looks like random noise). Indeed, Table S1 presents such cases. Peaks of Raw *clk* mRNA (*SD* ± 8.59*h*), PER protein (*SD* ± 5.99*h*), and *per* mRNA valley (*SD* ± 6.22*h*) all show this problem, but there are also enough other cases to clearly demonstrate that not all time series are affected.

**TABLE S1.** The impact of linearizing circadian time on summary statistics like the *Peak* (A) and *Valley* (B) traits of time series.

| **A** |  | **Peak** |  |  |  | **Peak** |  |  |  |  |
| --- | --- | --- | --- | --- | --- | --- | --- | --- | --- | --- |
| **Clock Component** | $n_{Raw}$ | Median | | | | Avg ± SD | | | | |
| | $n_{Mod}$ | *Raw*<br>*Circ* | *Mod*<br>*Linear* | *Mod*<br>*Maj* | *Mod*<br>*Odd* | *Raw*<br>*Circ* | *Mod*<br>*Linear* | *Mod*<br>*Maj* | *Mod*<br>*Odd* | $n_{Maj}$<br>$n_{Odd}$ |
| *clk* mRNA | 12 | 6.00 | 3.56 | 3.56 | 5.03 | **8.79** | 2.79 | 3.56 | 3.02 | 4 |
| | 12 | | | | | **$\pm 8.59$** | $\pm 3.16$ | $\pm 1.12$ | $\pm 3.73$ | 8 |
| *per* mRNA | 88 | 14.9 | 14.9 | 15.0 | 12.35 | 14.62 | 14.64 | 14.72 | 14.48 | 68 |
| | 88 | | | | | $\pm 1.68$ | $\pm 1.65$ | $\pm 1.01$ | $\pm 2.87$ | 22 |
| *tim* mRNA | 48 | 14.83 | 14.83 | 15.0 | 12.0 | 14.67 | 14.54 | 14.85 | 14.14 | 30 |
| | 48 | | | | | $\pm 2.09$ | $\pm 1.92$ | $\pm 0.98$ | $\pm 2.66$ | 22 |
| PER protein | 36 | 18.93 | 19.0 | 19.0 | 16.87 | **16.61** | 17.72 | 18.61 | 19.71 | 30 |
| | 36 | | | | | **$\pm 5.99$** | $\pm 4.79$ | $\pm 2.10$ | $\pm 5.90$ | 10 |
| TIM protein | 39 | 19.0 | 19.0 | 19.0 | 19.0 | 18.03 | 18.65 | 18.50 | 18.65 | 35 |
| | 39 | | | | | $\pm 3.50$ | $\pm 2.83$ | $\pm 1.35$ | $\pm 4.29$ | 11 |

| **B** |  | **Valley** |  |  |  | **Valley** |  |  |  |  |
| --- | --- | --- | --- | --- | --- | --- | --- | --- | --- | --- |
| **Clock Component** | $n_{Raw}$ | Median | | | | Avg ± SD | | | | |
| | $n_{Mod}$ | *Raw*<br>*Circ* | *Mod*<br>*Linear* | *Mod*<br>*Maj* | *Mod*<br>*Odd* | *Raw*<br>*Circ* | *Mod*<br>*Linear* | *Mod*<br>*Maj* | *Mod*<br>*Odd* | $n_{Maj}$<br>$n_{Odd}$ |
| *clk* mRNA | 30 | 14.97 | 14.97 | 14.97 | 14.50 | 14.33 | 14.33 | 14.58 | 14.15 | 17 |
| | 30 | | | | | $\pm 1.67$ | $\pm 1.67$ | $\pm 0.69$ | $\pm 2.12$ | 17 |
| *per* mRNA | 24 | 5.0 | 4.5 | 5.0 | 4.75 | **6.61** | 3.61 | 4.20 | 3.65 | 20 |
| | 24 | | | | | **$\pm 6.22$** | $\pm 2.60$ | $\pm 1.36$ | $\pm 3.40$ | 10 |
| *tim* mRNA | 21 | 5.0 | 5.0 | 5.0 | 5.0 | 4.86 | 4.86 | 5.13 | 4.84 | 18 |
| | 21 | | | | | $\pm 1.52$ | $\pm 1.52$ | $\pm 0.81$ | $\pm 1.37$ | 11 |
| PER protein | 47 | 9.0 | 9.0 | 8.68 | 12.0 | 9.01 | 9.27 | 8.68 | 10.08 | 32 |
| | 47 | | | | | $\pm 2.17$ | $\pm 2.10$ | $\pm 0.92$ | $\pm 3.06$ | 15 |
| TIM protein | 36 | 4.0 | 4.0 | 4.0 | 10 | 6.19 | 6.19 | 4.83 | 9.66 | 29 |
| | 36 | | | | | $\pm 3.26$ | $\pm 3.26$ | $\pm 1.51$ | $\pm 3.98$ | 10 |

### Solution by linearization

Close inspection indicated the problem (see above). We used these insights to devise a linearization procedure. The manual effort required for detecting the problem in the first place and then developing a solution consumed substantial amounts of research time. Other researchers will have to retrace or re-invent our work at substantial cost if they aim to avoid this problem. If a solution could be made available in the form of a compiler, then others could use and build on our results with almost no effort.

### How a simple VBIR compiler can help

Compilers can easily detect type-mismatches such as “*DZT*” vs “*CZT*” and plainly refuse any operations that require any mix of types. While this would mean that our data could not be analyzed with automated help, it would also guarantee that the root problem of type mismatch is brought to the attention of researchers.

### How a more advanced VBIR compiler can help

Sophistication beyond plain checks of type labels requires dedicated code addressing specific problem using domain specific biological expertise. For example, a *VBIR* compiler with definitions for a well-developed type system for circadian time series analysis could automatically check for potential *DZT* vs *CZT* confusion when calculating statistics beyond day 1, while allowing day 1 to be calculated if all data points are indeed from that day. Such a compiler would use a test for matching types to deduce the implicit drop of days from inspecting the type mismatch in combination with the actual data available at days beyond the first and explore remaining safe options. If user-friendly enough, it would create an understandable error diagnosis and alert the biological model curator in charge. If intelligent enough, such a compiler will offer a list of potential solutions from which the biologist in charge can choose efficiently how to address this problem and newcomers could request additional information. If analyses could only be performed by lucky accident (because *this* dataset allowed it, but others might not), then this compiler could produce a warning to that effect, so curators can anticipate potential problems as new data arrives.

### Other examples

It would be easy to create a long and detailed list of cases where appropriate support by a compiler could have speeded up our work with FlyClockbase. Examples include the lack of support for expressing ranges or three-point-estimates for parameter values or measurements with substantial uncertainty, the lack of warnings when spreadsheet parsing or calculating algorithms come across values of the wrong type, the lack of appropriate number systems that guarantee correct handling of cyclical and linear numbers and many more.

### Compilers that directly assist in biological research

A sufficiently advanced compiler would be like a helpful assistant^2^, finding errors like needles in haystacks of potential problems, cutting debugging time, and preventing untold additional problems on the way. Such a compiler will be useful to the degree that it can provide diagnoses and abstractions for detailed and difficult challenges. These challenges need to be solved first by a human, who can then build/teach a compiler how to solve the problem.

### Expected cost reductions

Once the compiler will then be able to routinely perform the corresponding work the cost savings for *VBIR* development and teaching become substantial (the compiler can easily point to additional resources and more as demonstrated by the integrated help function of the *R* statistical programming language. If statistics has its language in *R*, why does biology not have its own language? The efficiencies gained would allow the scientists of today to enable the next generation of scientists tomorrow to get much faster to stand on the shoulders of giants. The compiler would be like an aerial cableway, which gets others faster to a place of uncluttered overviews.

## Notes

1 It is important to distinguish such fast-moving research code from other, more mature scientific code that implements important algorithms which have become important research tools and are thus expected to behave reliably.

2 See http://elm-lang.org/blog/compilers-as-assistants for an exposition of what it might mean if compilers actually operated as assistants. This problem is inextricably linked to questions of syntax design. For example, see the importance of operator priorities [Razali et al., 2015, Operators and Precedence in Programming Languages.] and parsing which impacts the flexibility and quality of error messages that a compiler can produce [see Grune & Jacobs, 2008, *Parsing techniques: a practical guide*. Springer, New York.]

## LITERATURE CITED IN SUPPLEMENTAL MATERIAL

Aho, A. V. L., Monica S.; Sethi, Ravi; Ullman, Jeffrey D., 2007 Compilers: Principles, Techniques, and Tools. Pearson Education, Boston, MA.

Allison, D. B., A. W. Brown, B. J. George and K. A. Kaiser, 2016 Reproducibility: A tragedy of errors. Nature 530: 27–29.

Blankenberg, D., J. E. Johnson, J. Taylor, A. Nekrutenko and G. Team, 2014 Wrangling Galaxy’s reference data. Bioinformatics 30: 1917–1919.

Bosc, P., and O. Pivert, 2010 Modeling and Querying Uncertain Relational Databases: A Survey of Approaches Based on the Possible Worlds Semantics. International Journal of Uncertainty Fuzziness and Knowledge-Based Systems 18: 565–603.

Bourne, P. E., and J. McEntyre, 2006 Biocurators: Contributors to the world of science. Plos Computational Biology 2: 1185–1185.

Burge, S., T. K. Attwood, A. Bateman, T. Z. Berardini, M. Cherry et al., 2012 Biocurators and Biocuration: surveying the 21st century challenges. Database-the Journal of Biological Databases and Curation.

Burkhardt, K., B. Schneider and J. Ory, 2006 A biocurator perspective: Annotation at the research collaboratory for structural bioinformatics protein data bank. Plos Computational Biology 2: 1186–1189.

Candan, K. S., J. Grant and V. S. Subrahmanian, 1997 A unified treatment of null values using constraints. Information Sciences 98: 99–156.

Carthey, J., M. R. de Leval and J. T. Reason, 2001 The human factor in cardiac surgery: errors and near misses in a high technology medical domain. Ann Thorac Surg 72: 300–305.

Cooper, K. D. T., Linda, 2012 Engineering A Compiler. Morgan Kaufmann, Burlington, MA.

de Tre, G., R. de Caluwe and H. Prade, 2004 Null values revisited in prospect of data integration. Semantics of a Networked World: Semantics for Grid Databases 3226: 79–90.

Donoho, D. M., Arian; Rahman, Inam; Shahram, Morteza; Stodden, Victoria, 2009 15 Years of Reproducible Research in Computational Harmonic Analysis. Computing in Science and Engineering 2009: 8–18.

Dopfer, D., W. Buist, Y. Soyer, M. A. Munoz, R. N. Zadoks et al., 2008 Assessing genetic heterogeneity within bacterial species isolated from gastrointestinal and environmental samples: How many isolates does it take? Applied and Environmental Microbiology 74: 3490–3496.

Drake, V. J., B. Frei and L. M. Bouley, 2008 An error in the US Department of Agriculture nutrient database results in vitamin A values that are 6 times too high. Am J Clin Nutr 87: 1067–1068; author reply 1068-1069.

Ehlert, K., and L. Loewe, 2014 Lazy Updating of hubs can enable more realistic models by speeding up stochastic simulations. J Chem Phys 141: 204109.

Ember, C., R. Hanisch, G. Alter, H. Berman, M. Hedstrom et al., 2013 Sustaining Domain Repositories for Digital Data: A White Paper, pp. http://datacommunity.icpsr.umich.edu/sites/default/files/WhitePaper_ICPSR_SDRDD_121113.pdf in Workshop on Sustaining Domain Repositories for Digital Data, June 24-25, 2013, ICPSR, University of Michigan, Alfred P. Sloan Foundation.

Freedman, L. P., I. M. Cockburn and T. S. Simcoe, 2015 The economics of reproducibility in preclinical research. PLoS Biol 13: e1002165.

Galletta, D. F., D. Abraham, M. El Louadi, W. Lekse, Y. A. Pollailis et al., 1993 An Empirical Study of Spreadsheet Error Performance. Journal of Accounting, Managment, and Information Technology 3: 79–95.

Galletta, D. F., K. S. Hartzel, S. Johnson and J. L. Joseph, 1997 Spreadsheet Presentation and Error Detection: An Experimental Journal. Journal of Management Information Systems 13: 45–63.

Gitelman, L., 2013 "Raw data" is an oxymoron. The MIT Press, Cambridge, Massachusetts; London, England.

Gödel, K., 1931 Über formal unentscheidbare Sätze der Principia Mathematica und verwandter Systeme, I. Monatshefte für Mathematik und Physik 38: 173–198.

Goecks, J., A. Nekrutenko, J. Taylor and T. Galaxy, 2010 Galaxy: a comprehensive approach for supporting accessible, reproducible, and transparent computational research in the life sciences. Genome Biol 11: R86.

Goldston, D., 2008 Big data: Data wrangling. Nature 455: 15–15.

Grimes, M. L., W. J. Lee, L. Van der Maaten and P. Shannon, 2013 Wrangling Phosphoproteomic Data to Elucidate Cancer Signaling Pathways. Plos One 8.

Grove, S. J., and N. E. Stork, 2000 An inordinate fondness for beetles. Invertebrate Taxonomy 14: 733–739.

Grune, D., and C. J. H. Jacobs, 2008 Parsing techniques: a practical guide. Springer, New York.

Grune, D. e. a., 2012 Modern Compiler Design. Springer Science, New York, NY.

Hamilton, A. J., Y. Basset, K. K. Benke, P. S. Grimbacher, S. E. Miller et al., 2010 Quantifying Uncertainty in Estimation of Tropical Arthropod Species Richness. American Naturalist 176: 90–95.

Hartmann, S., and S. Link, 2012 The Implication Problem of Data Dependencies over SQL Table Definitions: Axiomatic, Algorithmic and Logical Characterizations. Acm Transactions on Database Systems 37.

Hayden, E. C., 2014 Technology: The $1,000 genome. Nature 507: 294–295.

Henry, P. L., 2013 "Ultimately, the source code is the documentation for developers" quoted in: "Lessons from the Agile trenches: The ups and downs of Agile development". ibm.com/developerWorks: http://www.ibm.com/developerworks/library/a-agiletrenches/a-agiletrenches-pdf.pdf.

Hernich, A., L. Libkin and N. Schweikardt, 2011 Closed World Data Exchange. Acm Transactions on Database Systems 36.

Hooey, B. L., M. Aurisicchio, R. Bracewell and D. C. Foyle, 2014 Evidence-Based Error Analysis: Supporting the Design of Error-Tolerant Systems. Human-Computer Interaction: Applications and Services, Pt Iii 8512: 401–412.

Howe, D., M. Costanzo, P. Fey, T. Gojobori, L. Hannick et al., 2008 Big data: The future of biocuration. Nature 455: 47–50.

Hutchins, L. N., Y. Ding, J. P. Szatkiewicz, R. Von Smith, H. Yang et al., 2010 CGDSNPdb: a database resource for error-checked and imputed mouse SNPs. Database (Oxford) 2010: baq008.

James, D., N. Wilkins-Diehr, V. Stodden, D. Colbry, C. Rosales et al., 2015 Standing Together for Reproducibility in Large-Scale Computing: Report on reproducibility@XSEDE, pp. http://arxiv.org/abs/1412.5557 in Arxiv: Distributed, Parallel, and Cluster Computing (cs.DC), arXiv:1412.5557v2 [cs.DC].

Jones, C. E., A. L. Brown and U. Baumann, 2007 Estimating the annotation error rate of curated GO database sequence annotations. BMC Bioinformatics 8: 170.

Jurney, R., 2013 Agile data science. O’Reilly Media, Beijing; Sebastopol, CA.

Karr, J. R., J. C. Sanghvi, D. N. Macklin, M. V. Gutschow, J. M. Jacobs et al., 2012 A whole-cell computational model predicts phenotype from genotype. Cell 150: 389–401.

Karr, J. R., K. Takahashi and A. Funahashi, 2015a The principles of whole-cell modeling. Curr Opin Microbiol 27: 18–24.

Karr, J. R., A. H. Williams, J. D. Zucker, A. Raue, B. Steiert et al., 2015b Summary of the DREAM8 Parameter Estimation Challenge: Toward Parameter Identification for Whole-Cell Models. PLoS Comput Biol 11: e1004096.

Kenall, A., S. Edmunds, L. Goodman, L. Bal, L. Flintoft et al., 2015 Better reporting for better research: a checklist for reproducibility. Gigascience 4: 32.

Knuth, D. E., and S. Levy, 1994 The CWEB system of structured documentation: version 3.0. Addison-Wesley Pub. Co., Reading, Mass.

Lander, E. S., L. M. Linton, B. Birren, C. Nusbaum, M. C. Zody et al., 2001 Initial sequencing and analysis of the human genome. Nature 409: 860–921.

Lee, Y., and J. Quackenbush, 2003 Using the TIGR gene index databases for biological discovery. Curr Protoc Bioinformatics Chapter 1: Unit 1 6.

Lee, Y., J. Tsai, S. Sunkara, S. Karamycheva, G. Pertea et al., 2005 The TIGR Gene Indices: clustering and assembling EST and known genes and integration with eukaryotic genomes. Nucleic Acids Res 33: D71–74.

Lesgold, A., H. Rubinson, P. Feltovich, R. Glaser, D. Klopfer et al., 1988 Expertise in a Complex Skill: Diagnosing X-Ray Pictures. in The Nature of Expertise., edited by R. G. Micheline Chi and M. Farr. Lawrence Erlbaum, Hillsdale, NJ.

Lifschitz, V., K. Pichotta and F. K. Yang, 2012 Relational theories with null values and non-herbrand stable models. Theory and Practice of Logic Programming 12: 565–582.

Loewe, L., 2016 Systems in Evolutionary System Biology. Encyclopedia of Evolutionary Biology ed. 1, vol. 4: 297–318 (http://evolutionarysystemsbiology.org/pdf/Loewe-2016-evosysbio.pdf).

Loewe, L., and S. Keel, 2014 BEST Names for Semantic Units to Support Reproducible Modeling, pp. 4 pages, Workshop: "reproducibility@XSEDE", XSEDE14 Annual Conference https://www.xsede.org/documents/659353/703287/xsede659314_loewe.pdf in *2014-07-14* (https://www.xsede.org/web/reproducibility), Atlanta, GA, USA.

Loewe, L., K. S. Scheuer and S. Keel, 2016 Evolvix Concept: BEST Names for Semantic Units Increase Semantic Reproducibility and Help Combine Complex Biological Models, pp. https://sites.google.com/site/dslabhs2016/proceeding-papers in Workshop on Data Science, Learning, and Applications to Biomedical & Health Sciences, January 7 - 8, 2016. NY Academy of Sciences, New York Academy of Sciences.

Loewe, L., K. S. Scheuer, S. A. Keel, V. Vyas, B. Liblit et al., 2017 Evolvix BEST Names for semantic reproducibility across code2brain interfaces; with 74 pages online material at http://dx.doi.org/10.1111/nyas.13192 and updates at http://evolvix.org/naming; http://evolvix.org/post. Annals of the New York Academy of Sciences 1387: 124–144.

Magnussen, S., B. Smith, C. Kleinn and I. F. Sun, 2010 An urn model for species richness estimation in quadrat sampling from fixed-area populations. Forestry 83: 293–306.

Mahmood, Z., 2016 Data science and big data computing: frameworks and methodologies. Springer Berlin Heidelberg, New York, NY.

Martinez, M. V., C. Molinaro, J. Grant and V. S. Subrahmanian, 2013 Customized Policies for Handling Partial Information in Relational Databases. Ieee Transactions on Knowledge and Data Engineering 25: 1254–1271.

McCallum, Q. E., 2013 Bad Data Handbook: Mapping the world of data problems. O’Reilly, Sebastopol, CA.

Mirza, G. A., 2015 Null Value Conflict: Formal Definition and Resolution. 2015 13th International Conference on Frontiers of Information Technology (Fit): 132–137.

NASA, F. Chandler, T., Y. H. Chang, James, A. Mosleh, J. Marble, L. et al., 2006-07 Human Reliability Analysis Methods Selection Guidance for NASA. NASA/OSMA, NASA, Washington, DC.

NASA, K. Leiden, K. R. Laughery, J. Keller, J. French et al., 2001-09-30 A Review of Human Performance Models for the Prediction of Human Error. National Aeronautics and Space Administration, System-Wide Accident Prevention Program, Ames Research Center, Moffett Field, CA, http://citeseerx.ist.psu.edu/viewdoc/download?doi=10.1.1.468.6969&rep=rep1&type=pdf

NASA, M. Stamatelatos and H. Dezfuli, 2011 Probabilistic Risk Assessment Procedures Guide for NASA Managers and Practitioners (2nd Edition), NASA Headquarters, Washington, DC.

NIH, D. DeMets, L. Tabak, R. Altman, D. Botstein et al., 2012 Data and Informatics Working Group Report to the Advisory Committee to the Director, pp., http://acd.od.nih.gov/DataandInformaticsWorkingGroupReport.PDF.

Orchard, S., and H. Hermjakob, 2015 Shared resources, shared costs–leveraging biocuration resources. Database (Oxford) 2015.

Panko, R. R., 1998 Applying code inspection to spreadsheet testing. Effective Utilization and Management of Emerging Information Technologies: 358–362.

Panko, R. R., 2016 Human Error Website. Accessed 2016-10-31: http://panko.shidler.hawaii.edu/HumanErr/Index.htm

Panko, R. R., and S. Aurigemma, 2010 Revising the Panko-Halverson taxonomy of spreadsheet errors. Decision Support Systems 49: 235–244.

Panko, R. R., and R. H. Sprague, 1998 Hitting the wall: errors in developing and code inspecting a ‘simple’ spreadsheet model. Decision Support Systems 22: 337–353.

Parsons, T. W., 1992 Introduction to Compiler Construction. Computer Science Press, New York.

Pertea, G., X. Huang, F. Liang, V. Antonescu, R. Sultana et al., 2003 TIGR Gene Indices clustering tools (TGICL): a software system for fast clustering of large EST datasets. Bioinformatics 19: 651–652.

Poldrack, R. A., and J. B. Poline, 2015 The publication and reproducibility challenges of shared data. Trends Cogn Sci 19: 59–61.

Quackenbush, J., F. Liang, I. Holt, G. Pertea and J. Upton, 2000 The TIGR gene indices: reconstruction and representation of expressed gene sequences. Nucleic Acids Res 28: 141–145.

Rak, R., R. T. Batista-Navarro, A. Rowley, J. Carter and S. Ananiadou, 2014 Text-mining-assisted biocuration workflows in Argo. Database (Oxford) 2014.

Razali, N., J. Noble and S. Marshall, 2015-10-26 Operators and Precedence in Programming Languages. PLATEAU’ 15: 53–56 (http://dx.doi.org/10.1145/2846680.2846690).

Reason, J., 2005 Safety in the operating theatre - Part 2: human error and organisational failure. Qual Saf Health Care 14: 56–60.

Reason, J. T., 1990 Human error. Cambridge University Press, Cambridge England; New York.

Reason, J. T., 2013 A life in error: from little slips to big disasters. Ashgate, Surrey, UK England.

Reason, J. T., 2015 Organizational accidents revisited. Ashgate, Burlington, VT.

Reason, J. T., and A. Hobbs, 2003 Managing maintenance error: a practical guide. Ashgate, Aldershot, Hampshire, England; Burlington, VT, USA.

Reason, J. T., and K. Mycielska, 1982 Absent-minded?: the psychology of mental lapses and everyday errors. Prentice-Hall, Englewood Cliffs, N.J.

Reiser, L., T. Z. Berardini, D. Li, R. Muller, E. M. Strait et al., 2016 Sustainable funding for biocuration: The Arabidopsis Information Resource (TAIR) as a case study of a subscription-based funding model. Database (Oxford) 2016.

Rodriguez-Esteban, R., 2015 Biocuration with insufficient resources and fixed timelines. Database (Oxford) 2015.

Salimi, N., and R. Vita, 2006 The biocurator: Connecting and enhancing scientific data. Plos Computational Biology 2: 1190–1192.

Sanger, F., G. M. Air, B. G. Barrell, N. L. Brown, A. R. Coulson et al., 1977a Nucliotide sequence of bacteriophage phi X174 DNA. Nature 265: 687–695.

Sanger, F., S. Nicklen and A. R. Coulson, 1977b DNA sequencing with chain-terminating inhibitors. Proceedings of the National Academy of Sciences of the U.S.A. 74: 5463–5467.

Sheridan, C., 2014 Illumina claims $1,000 genome win. Nat Biotechnol 32: 115.

Singhal, A., R. Leaman, N. Catlett, T. Lemberger, J. McEntyre et al., 2016 Pressing needs of biomedical text mining in biocuration and beyond: opportunities and challenges. Database (Oxford) 2016.

Smith, N. J. J., 2008 Vagueness and Degrees of Truth. Oxford University Press, Oxford, Great Britain.

Stodden, V., 2015 Reproducing Statistical Results. Annu. Rev. Stat. Appl. 2: 1–19.

Stodden, V., F. Leisch and R. D. Peng (Editors), 2014 Implementing Reproducible Research. CRC Press, Boca Raton, FL.

Telenti, A., L. C. Pierce, W. H. Biggs, J. di Iulio, E. H. Wong et al., 2016 Deep sequencing of 10,000 human genomes. Proc Natl Acad Sci U S A 113: 11901–11906.

Thalheim, B., and K. D. Schewe, 2011 NULL ‘Value’ Algebras and Logics. Information Modelling and Knowledge Bases Xxii 225: 354–367.

Venter, J. C., M. D. Adams, E. W. Myers, P. W. Li, R. J. Mural et al., 2001 The sequence of the human genome. Science 291: 1304–1351.

Waraporn, N., and K. Porkaew, 2008 Semantics of null values in subqueries. Imecs 2008: International Multiconference of Engineers and Computer Scientists, Vols I and Ii: 400–405.

Wei, C. H., H. Y. Kao and Z. Lu, 2013 PubTator: a web-based text mining tool for assisting biocuration. Nucleic Acids Res 41: W518–522.

Weil, G., C. Motamed, A. Eghiaian, M. L. Guye and J. L. Bourgain, 2015 The use of a clinical database in an anesthesia unit: focus on its limits. J Clin Monit Comput 29: 163–167.

Zaniolo, C., 1984 Database Relations with Null Values. Journal of Computer and System Sciences 28: 142–166.

Zhang, H. M., and H. Stern, 2009 Sample Size Calculation for Finding Unseen Species. Bayesian Analysis 4: 763–791.

## LITERATURE CITED IN MAIN TEXT

2016 Advances in mathematical modeling, optimization and optimal control. Springer Science+Business Media, New York, NY.

Aarts, A. A., J. E. Anderson, C. J. Anderson, P. R. Attridge, A. Attwood et al., 2015 Estimating the reproducibility of psychological science. Science 349.

Abruzzi, K., X. Chen, E. Nagoshi, A. Zadina and M. Rosbash, 2015 RNA-seq profiling of small numbers of Drosophila neurons. Meth Enzymol 551: 369–386.

Abruzzi, K. C., J. Rodriguez, J. S. Menet, J. Desrochers, A. Zadina et al., 2011 Drosophila CLOCK target gene characterization: implications for circadian tissue-specific gene expression. Genes & development 25: 2374–2386.

Ades, A. E., N. J. Welton, D. Caldwell, M. Price, A. Goubar et al., 2008 Multiparameter evidence synthesis in epidemiology and medical decision-making. J Health Serv Res Policy 13 Suppl 3: 12–22.

Adewoye, A. B., C. P. Kyriacou and E. Tauber, 2015 Identification and functional analysis of early gene expression induced by circadian light-resetting in Drosophila. BMC Genomics 16: 570.

Agostinelli, F., N. Ceglia, B. Shahbaba, P. Sassone-Corsi and P. Baldi, 2016 What time is it? Deep learning approaches for circadian rhythms. Bioinformatics 32: i8–i17.

Aimo, L., R. Liechti, N. Hyka-Nouspikel, A. Niknejad, A. Gleizes et al., 2015 The SwissLipids knowledgebase for lipid biology. Bioinformatics 31: 2860–2866.

Aird, D., M. G. Ross, W. S. Chen, M. Danielsson, T. Fennell et al., 2011 Analyzing and minimizing PCR amplification bias in Illumina sequencing libraries. Genome Biol 12: R18.

Akman, O., M. L. Guerriero, L. Loewe and C. Troein, 2010 Complementary approaches to understanding the plant circadian clock. Proceedings 3rd Workshop ‘’From Biology To Concurrency and back’’ - FBTC 2010 published in Electronic Proceedings in Theoretical Computer Science 19: 1–19 (http://arxiv.org/abs/1002.4661v1001).

Akten, B., E. Jauch, G. K. Genova, E. Y. Kim, I. Edery et al., 2003 A role for CK2 in the Drosophila circadian oscillator. Nature neuroscience 6: 251–257.

Allada, R., N. E. White, W. V. So, J. C. Hall and M. Rosbash, 1998 A mutant Drosophila homolog of mammalian Clock disrupts circadian rhythms and transcription of period and timeless. Cell 93: 791–804.

Altman, D. G., S. M. Gore, M. J. Gardner and S. J. Pocock, 1983 Statistical guidelines for contributors to medical journals. Br Med J (Clin Res Ed) 286: 1489–1493.

Amos, W., C. M. Hutter, M. D. Schug and C. F. Aquadro, 2003 Directional evolution of size coupled with ascertainment bias for variation in Drosophila microsatellites. Mol Biol Evol 20: 660–662.

Anderson, D. F., 2007 A modified next reaction method for simulating chemical systems with time dependent propensities and delays. J Chem Phys 127: 214107.

Anderson, D. F., A. Ganguly and T. G. Kurtz, 2011 Error Analysis of Tau-Leap Simulation Methods. Annals of Applied Probability 21: 2226–2262.

Annosi, M. C., M. Magnusson, A. Martini and F. P. Appio, 2016 Social Conduct, Learning and Innovation: An Abductive Study of the Dark Side of Agile Software Development. Creativity and Innovation Management 25: 515–535.

Arp, R., B. Smith and A. D. Spear, 2015 Building ontologies with Basic Formal Ontology. Massachusetts Institute of Technology, Cambridge, Massachusetts http://ontology.buffalo.edu/smith/.

Ascher, U. M., and L. R. Petzold, 1998 Computer Methods for Ordinary Differential Equations and Differential-Algebraic Equations. SIAM, Philadelphia.

Ashburner, M., and R. Drysdale, 1994 FlyBase–the Drosophila genetic database. Development 120: 2077–2079.

Ashley, E. A., 2016 Towards precision medicine. Nat Rev Genet 17: 507–522.

Bae, K., and I. Edery, 2006 Regulating a Circadian Clock’s Period, Phase and Amplitude by Phosphorylation: Insights from Drosophila. Journal of Biochemistry 140: 609–617.

Bae, K., C. Lee, P. Hardin and I. Edery, 2000 dCLOCK is present in limiting amounts and likely mediates daily interactions between the dCLOCK-CYC transcription factor and the PER-TIM complex. Journal of Neuroscience 20: 1746–1753.

Bae, K., C. Lee, D. Sidote, K. Y. Chuang and I. Edery, 1998 Circadian regulation of a Drosophila homolog of the mammalian Clock gene: PER and TIM function as positive regulators. Mol Cell Biol 18: 6142–6151.

Bagheri, N., M. J. Lawson, J. Stelling and F. J. Doyle, 2008a Modeling the Drosophila melanogaster Circadian Oscillator via Phase Optimization. Journal of biological rhythms 23: 525–537.

Bagheri, N., J. Stelling and F. J. Doyle, 2007 Circadian phase entrainment via nonlinear model predictive control. International Journal of Robust and Nonlinear Control 17: 1555–1571.

Bagheri, N., S. R. Taylor, K. Meeker, L. R. Petzold and F. J. Doyle, 2008b Synchrony and entrainment properties of robust circadian oscillators., pp. S17–28 in J R Soc Interface.

Baker, M., 2016 1,500 scientists lift the lid on reproducibility. Nature 533: 452–454.

Bammler, T., R. P. Beyer, S. Bhattacharya, G. A. Boorman, A. Boyles et al., 2005 Standardizing global gene expression analysis between laboratories and across platforms. Nat Methods 2: 351–356.

Banko, Z., and J. Abonyi, 2015 Mixed dissimilarity measure for piecewise linear approximation based time series applications. Expert Systems with Applications 42: 7664–7675.

Bao, S., J. Rihel, E. Bjes, J. Y. Fan and J. L. Price, 2001 The Drosophila double-timeS mutation delays the nuclear accumulation of period protein and affects the feedback regulation of period mRNA. Journal of Neuroscience 21: 7117–7126.

Barba, L. A., 2016 The hard road to reproducibility. Science 354: 142–142.

Barkai, N., and S. Leibler, 2000 Circadian clocks limited by noise. Nature 403: 267–268.

Barrett, T., T. O. Suzek, D. B. Troup, S. E. Wilhite, W. C. Ngau et al., 2005 NCBI GEO: mining millions of expression profiles–database and tools. Nucleic Acids Res 33: D562–566.

Barrett, T., D. B. Troup, S. E. Wilhite, P. Ledoux, C. Evangelista et al., 2011 NCBI GEO: archive for functional genomics data sets–10 years on. Nucleic Acids Res 39: D1005–1010.

Barrett, T., S. E. Wilhite, P. Ledoux, C. Evangelista, I. F. Kim et al., 2013 NCBI GEO: archive for functional genomics data sets–update. Nucleic Acids Res 41: D991–995.

Bartling, S., and S. Friesike, 2014 Opening science. Springer, New York, http://book.openingscience.org/

Bateman, A., 2010 Curators of the world unite: the International Society of Biocuration. Bioinformatics 26: 991.

Batista, G. E. A. P. A., X. Wang and E. J. Keogh, 2011 A Complexity-Invariant Distance Measure for Time Series.

Batista, R. T., D. B. Ramirez, R. D. Santos, M. C. del Rosario and E. R. Mendoza, 2007 EUCLIS–an information system for circadian systems biology. IET Syst Biol 1: 266–273.

Bauer, P., A. Thorpe and G. Brunet, 2015 The quiet revolution of numerical weather prediction. Nature 525: 47–55.

Baylies, M. K., L. B. Vosshall, A. Sehgal and M. W. Young, 1992 New Short-Period Mutations of the Drosophila Clock Gene Per. Neuron 9: 575–581.

Beaver, L. M., B. O. Gvakharia, T. S. Vollintine, D. M. Hege, R. Stanewsky et al., 2002 Loss of circadian clock function decreases reproductive fitness in males of Drosophila melanogaster., pp. 2134–2139 in P Natl Acad Sci Usa.

Beaver, L. M., L. A. Hooven, S. M. Butcher, N. Krishnan, K. A. Sherman et al., 2010 Circadian clock regulates response to pesticides in Drosophila via conserved Pdp1 pathway., pp. 513–520 in Toxicol. Sci.

Beaver, L. M., B. L. Rush, B. O. Gvakharia and J. M. Giebultowicz, 2003 Noncircadian regulation and function of clock genes period and timeless in oogenesis of Drosophila melanogaster., pp. 463–472 in J Biol Rhythms.

Beersma, D. G. M., 2005 Why and how do we model circadian rhythms? Journal of Biological Rhythms 20: 304–313.

Bell-Pedersen, D., V. M. Cassone, D. J. Earnest, S. S. Golden, P. E. Hardin et al., 2005 Circadian rhythms from multiple oscillators: Lessons from diverse organisms. Nature Reviews Genetics 6: 544–556.

Berndt, A., T. Kottke, H. Breitkreuz, R. Dvorsky, S. Hennig et al., 2007 A Novel Photoreaction Mechanism for the Circadian Blue Light Photoreceptor Drosophila Cryptochrome. Journal of Biological Chemistry 282: 13011–13021.

Bishop, C. M., 2006 Pattern recognition and machine learning. Springer, New York.

Blau, J., 1999 Cycling vrille Expression Is Required for a Functional Drosophila Clock. Cell 99: 661–671.

Boos, D. D., and L. A. Stefanski, 2011 P-Value Precision and Reproducibility. American Statistician 65: 213–221.

Bourne, P. E., and J. McEntyre, 2006 Biocurators: Contributors to the world of science. PLOS Computational Biology 2: 1185–1185.

Boyle, G., K. Richter, H. D. Priest, D. Traver, T. C. Mockler et al., 2017 Comparative Analysis of Vertebrate Diurnal/Circadian Transcriptomes. PLOS ONE 12: e0169923.

Brandes, C., J. D. Plautz, R. Stanewsky, C. F. Jamison, M. Straume et al., 1996 Novel features of drosophila period Transcription revealed by real-time luciferase reporting. Neuron 16: 687–692.

Brodland, G. W., 2015 How computational models can help unlock biological systems. Seminars in Cell & Developmental Biology 47–48.

Broman, K. W., 1999 Cleaning genotype data. Genet Epidemiol 17 Suppl 1: S79–83.

Broman, K. W., M. P. Keller, A. T. Broman, C. Kendziorski, B. S. Yandell et al., 2015 Identification and Correction of Sample Mix-Ups in Expression Genetic Data: A Case Study. G3 (Bethesda) 5: 2177–2186.

Brooksbank, C., G. Cameron and J. Thornton, 2005 The European Bioinformatics Institute’s data resources: towards systems biology. Nucleic Acids Res 33: D46–53.

Buck, S., 2015 Solving reproducibility. Science 348: 1403.

Bullard, J. H., E. Purdom, K. D. Hansen and S. Dudoit, 2010 Evaluation of statistical methods for normalization and differential expression in mRNA-Seq experiments. Bmc Bioinformatics 11.

Burkhardt, K., B. Schneider and J. Ory, 2006 A biocurator perspective: Annotation at the research collaboratory for structural bioinformatics protein data bank. PLOS Computational Biology 2: 1186–1189.

Bussieck, M. R., and A. Meeraus, 2004 General algebraic modeling system (GAMS). Modeling Languages in Mathematical Optimization 88: 137–157.

Bustin, S. A., 2002 Quantification of mRNA using real-time reverse transcription PCR (RT-PCR): trends and problems. Journal of Molecular Endocrinology 29: 23–39.

Bustin, S. A., and T. Nolan, 2004 Pitfalls of quantitative real-time reverse-transcription polymerase chain reaction. J Biomol Tech 15: 155–166.

Busza, A., M. Emery-Le, M. Rosbash and P. Emery, 2004 Roles of the two Drosophila CRYPTOCHROME structural domains in circadian photoreception. Science 304: 1503–1506.

Buzbas, E. O., and N. A. Rosenberg, 2015 AABC: approximate approximate Bayesian computation for inference in population-genetic models. Theor Popul Biol 99: 31–42.

Canales, B. K., 2016 Comment On: Prospective evaluation of urinary metabolic indices in severely obese adolescents after weight loss surgery. Surg Obes Relat Dis 12: 367–368.

Canales, R. D., Y. Luo, J. C. Willey, B. Austermiller, C. C. Barbacioru et al., 2006 Evaluation of DNA microarray results with quantitative gene expression platforms. Nat Biotechnol 24: 1115–1122.

Carling, K., 2000 Resistant outlier rules and the non-Gaussian case. Computational Statistics & Data Analysis 33: 249–258.

Carter, S. J., H. J. Durrington, J. E. Gibbs, J. Blaikley, A. S. Loudon et al., 2016 A matter of time: study of circadian clocks and their role in inflammation. Journal of Leukocyte Biology 99: 549–560.

Carthey, J., M. R. de Leval, D. J. Wright, V. T. Farewell, J. T. Reason et al., 2003 Behavioural markers of surgical excellence. Safety Science 41: 409–425.

Cassman, M., 2005 Barriers to progress in systems biology. Nature 438: 1079.

Cathala, G., J. F. Savouret, B. Mendez, B. L. West, M. Karin et al., 1983 A Method for Isolation of Intact, Translationally Active Ribonucleic-Acid. DNA-a Journal of Molecular & Cellular Biology 2: 329–335.

Ceriani, M. F., T. K. Darlington, D. Staknis, P. Más, A. A. Petti et al., 1999 Light-dependent sequestration of TIMELESS by CRYPTOCHROME. Science 285: 553–556.

Ceriani, M. F., J. B. Hogenesch, M. Yanovsky, S. Panda, M. Straume et al., 2002 Genome-wide expression analysis in Drosophila reveals genes controlling circadian behavior. Journal of Neuroscience 22: 9305–9319.

Chelliah, V., N. Juty, I. Ajmera, R. Ali, M. Dumousseau et al., 2015 BioModels: ten-year anniversary. Nucleic Acids Res 43: D542–548.

Chen, K. F., N. Peschel, R. Zavodska, H. Sehadova and R. Stanewsky, 2011 QUASIMODO, a Novel GPI-Anchored Zona Pellucida Protein Involved in Light Input to the Drosophila Circadian Clock. Current biology: CB 21: 719–729.

Chen, W.-F., J. Majercak and I. Edery, 2006 Clock-gated photic stimulation of timeless expression at cold temperatures and seasonal adaptation in Drosophila., pp. 256–271 in J Biol Rhythms.

Cheng, Y., B. Gvakharia and P. E. Hardin, 1998 Two alternatively spliced transcripts from the Drosophila period gene rescue rhythms having different molecular and behavioral characteristics., pp. 6505–6514 in Mol Cell Biol.

Chintapalli, V. R., M. Al Bratty, D. Korzekwa, D. G. Watson and J. A. Dow, 2013a Mapping an atlas of tissue-specific Drosophila melanogaster metabolomes by high resolution mass spectrometry. PLOS ONE 8: e78066.

Chintapalli, V. R., J. Wang and J. A. Dow, 2007 Using FlyAtlas to identify better Drosophila melanogaster models of human disease. Nat Genet 39: 715–720.

Chintapalli, V. R., J. Wang, P. Herzyk, S. A. Davies and J. A. Dow, 2013b Data-mining the FlyAtlas online resource to identify core functional motifs across transporting epithelia. BMC Genomics 14: 518.

Chiu, J. C., H. W. Ko and I. Edery, 2011 NEMO/NLK phosphorylates PERIOD to initiate a time-delay phosphorylation circuit that sets circadian clock speed., pp. 357–370 in Cell.

Chiu, J. C., J. T. Vanselow, A. Kramer and I. Edery, 2008 The phospho-occupancy of an atypical SLIMB-binding site on PERIOD that is phosphorylated by DOUBLETIME controls the pace of the clock. Genes & development 22: 1758–1772.

Chomczynski, P., and N. Sacchi, 1987 Single-Step Method of Rna Isolation by Acid Guanidinium Thiocyanate Phenol Chloroform Extraction. Analytical Biochemistry 162: 156–159.

Chouhan, N. S., R. Wolf, C. Helfrich-Forster and M. Heisenberg, 2015 Flies remember the time of day. Curr Biol 25: 1619–1624.

Chylek, L. A., L. A. Harris, J. R. Faeder and W. S. Hlavacek, 2015 Modeling for (physical) biologists: an introduction to the rule-based approach. Physical Biology 12.

Cirelli, C., 2009 The genetic and molecular regulation of sleep: from fruit flies to humans. Nature Reviews Neuroscience 10: 549–560.

Citri, Y., H. V. Colot, A. C. Jacquier, Q. Yu, J. C. Hall et al., 1987 A family of unusually spliced biologically active transcripts encoded by a Drosophila clock gene., pp. 42–47 in Nature.

Claridge-Chang, A., H. Wijnen, F. Naef, C. Boothroyd, N. Rajewsky et al., 2001 Circadian regulation of gene expression systems in the Drosophila head. Neuron 32: 657–671.

Clark, A. G., M. J. Hubisz, C. D. Bustamante, S. H. Williamson and R. Nielsen, 2005 Ascertainment bias in studies of human genome-wide polymorphism. Genome Res 15: 1496–1502.

Clough, E., and T. Barrett, 2016 The Gene Expression Omnibus Database. Methods Mol Biol 1418: 93–110.

Collins, F. S., and L. A. Tabak, 2014 Policy: NIH plans to enhance reproducibility. Nature 505: 612–613.

Coogan, A. N., A. L. Baird, A. Popa-Wagner and J. Thome, 2016 Circadian rhythms and attention deficit hyperactivity disorder: The what, the when and the why. Progress in Neuro-Psychopharmacology & Biological Psychiatry 67: 74–81.

Cooper, K. D. T., Linda, 2012 Engineering A Compiler. Morgan Kaufmann, Burlington, MA.

Costa, R., A. A. Peixoto, G. Barbujani and C. P. Kyriacou, 1992 A latitudinal cline in a Drosophila clock gene., pp. 43–49 in Proc Biol Sci.

Cotterman, C. W., 1983 Relationship and probability in Mendelian populations. Am J Med Genet 16: 393–440.

Coyne, J. A., and H. A. Orr, 2004 Speciation. Sinauer Associates, Sunderland, Mass.

Crosby, J. L., 1973 Computer simulation in genetics. Wiley, London, New York,.

Crow, J. F., 2001 Shannon’s brief foray into genetics. Genetics 159: 915–917.

Csillery, K., M. G. Blum, O. E. Gaggiotti and O. Francois, 2010a Approximate Bayesian Computation (ABC) in practice. Trends Ecol Evol 25: 410–418.

Csillery, K., M. G. B. Blum, O. E. Gaggiotti and O. Francois, 2010b Approximate Bayesian Computation (ABC) in practice. Trends in Ecology & Evolution 25: 410–418.

Cumming, G., 2013 The New Statistics: A How-To Guide. Australian Psychologist 48: 161–170.

Cumming, G., 2014 The New Statistics: Why and How. Psychological Science 25: 7–29.

Curran-Everett, D., 2009 Explorations in statistics: confidence intervals. Advances in Physiology Education 33: 87–90.

Cusick, M. E., H. Yu, A. Smolyar, K. Venkatesan, A. R. Carvunis et al., 2009 Literature-curated protein interaction datasets. Nat Methods 6: 39–46.

Cyran, S. A., A. M. Buchsbaum, K. L. Reddy, M.-C. Lin, N. R. J. Glossop et al., 2003 vrille, Pdp1, and dClock form a second feedback loop in the Drosophila circadian clock. Cell 112: 329–341.

Daigle, B. J., Jr., M. K. Roh, L. R. Petzold and J. Niemi, 2012 Accelerated maximum likelihood parameter estimation for stochastic biochemical systems. BMC Bioinformatics 13: 68.

Darlington, T. K., K. Wager-Smith, M. F. Ceriani, D. Staknis, N. Gekakis et al., 1998 Closing the circadian loop: CLOCK-induced transcription of its own inhibitors per and tim. Science 280: 1599–1603.

Dawkins, R., 1976 The selfish gene. Oxford University Press, Oxford.

Dawkins, R., 2016 The selfish gene: 40th anniversary edition. Oxford University Press, New York, NY.

Debray, T. P., K. G. Moons, G. van Valkenhoef, O. Efthimiou, N. Hummel et al., 2015 Get real in individual participant data (IPD) meta-analysis: a review of the methodology. Res Synth Methods 6: 293–309.

Demidenko, E., 2016 The p-Value You Can’t Buy. American Statistician 70: 33–38.

Deng, Z., S. Arsenault, C. Caranica, J. Griffith, T. Zhu et al., 2016 Synchronizing stochastic circadian oscillators in single cells of Neurospora crassa. Sci Rep 6: 35828.

Diambra, L., and C. P. Malta, 2012 Modeling the emergence of circadian rhythms in a clock neuron network. PLOS ONE 7: e33912.

Ding, H., G. Trajcevski, P. Scheuermann, X. Wang and E. J. Keogh, 2008 Querying and Mining of Time Series Data Experimental Comparison of Representations and Distance Measures.

Dissel, S., V. Codd, R. Fedic, K. J. Garner, R. Costa et al., 2004 A constitutively active cryptochrome in Drosophila melanogaster. Nature neuroscience 7: 834–840.

DiStefano, J. J., 2013 Dynamic systems biology modeling and simulation. Elsevier, Academic Press, Amsterdam.

Doan, A., A. Halevy and Z. G. Ives, 2012 Principles of data integration. Morgan Kaufmann, Waltham, MA.

Dodd, A. N., N. Salathia, A. Hall, E. Kevei, R. Toth et al., 2005 Plant circadian clocks increase photosynthesis, growth, survival, and competitive advantage. Science 309: 630–633.

Doherty, C. J., and S. A. Kay, 2010 Circadian control of global gene expression patterns., pp. 419–444 in Annu Rev Genet.

Doi, S. A., 2014 Evidence synthesis for medical decision making and the appropriate use of quality scores. Clin Med Res 12: 40–46.

dos Santos, G., A. J. Schroeder, J. L. Goodman, V. B. Strelets, M. A. Crosby et al., 2015 FlyBase: introduction of the Drosophila melanogaster Release 6 reference genome assembly and large-scale migration of genome annotations. Nucleic Acids Research 43: D690–D697.

Drager, A., and B. O. Palsson, 2014 Improving collaboration by standardization efforts in systems biology. Front Bioeng Biotechnol 2: 61.

Drummond, G. B., and S. L. Vowler, 2011 Show the data, don’t conceal them. Journal of Physiology-London 589: 1861–1863.

Drummond, G. B., and S. L. Vowler, 2013 Statistical Reporting Guidelines. Type I: families, planning and errors. Exp Physiol 98: 3–6.

Dubruille, R. R., A. A. Murad, M. M. Rosbash and P. P. Emery, 2009 A constant light-genetic screen identifies KISMET as a regulator of circadian photoresponses. Audio, Transactions of the IRE Professional Group on 5: e1000787–e1000787.

Dumbell, R., O. Matveeva and H. Oster, 2016 Circadian Clocks, Stress, and Immunity. Frontiers in Endocrinology 7.

Edery, I., L. J. Zwiebel, M. E. Dembinska and M. Rosbash, 1994 Temporal phosphorylation of the Drosophila period protein., pp. 2260–2264 in P Natl Acad Sci Usa.

Edwards, A. W. F., 1992 Likelihood. John Hopkins University Press.

Efthimiou, O., T. P. Debray, G. van Valkenhoef, S. Trelle, K. Panayidou et al., 2016 GetReal in network meta-analysis: a review of the methodology. Res Synth Methods 7: 236–263.

Einstein, A., and A. Calaprice, 2011 The ultimate quotable Einstein. Princeton University Press, Princeton, N.J.

Emery, P., W. V. So, M. Kaneko, J. C. Hall and M. Rosbash, 1998 CRY, a Drosophila clock and light-regulated cryptochrome, is a major contributor to circadian rhythm resetting and photosensitivity. Cell 95: 669–679.

Erguler, K., and M. P. Stumpf, 2011 Practical limits for reverse engineering of dynamical systems: a statistical analysis of sensitivity and parameter inferability in systems biology models. Mol Biosyst 7: 1593–1602.

Eyre-Walker, A., and P. D. Keightley, 2007 The distribution of fitness effects of new mutations. Nat Rev Genet 8: 610–618.

Fang, Y., S. Sathyanarayanan and A. Sehgal, 2007 Post-translational regulation of the Drosophila circadian clock requires protein phosphatase 1 (PP1). Genes & development 21: 1506–1518.

Fathallah-Shaykh, H. M., 2010 Dynamics of the Drosophila Circadian Clock: Theoretical Anti-Jitter Network and Controlled Chaos. PLOS ONE 5.

Fathallah-Shaykh, H. M., J. L. Bona and S. Kadener, 2009 Mathematical Model of the Drosophila Circadian Clock: Loop Regulation and Transcriptional Integration. Biophysical Journal 97: 2399–2408.

Foll, M., M. A. Beaumont and O. Gaggiotti, 2008 An approximate Bayesian computation approach to overcome biases that arise when using amplified fragment length polymorphism markers to study population structure. Genetics 179: 927–939.

Forger, D., M. Drapeau, B. Collins and J. Blau, 2005 A new model for circadian clock research? Molecular systems biology 1: 2005.0014.

Freedman, L. P., I. M. Cockburn and T. S. Simcoe, 2015a The economics of reproducibility in preclinical research. PLOS Biol 13: e1002165.

Freedman, L. P., M. C. Gibson, S. P. Ethier, H. R. Soule, R. M. Neve et al., 2015b Reproducibility: changing the policies and culture of cell line authentication. Nat Methods 12: 493–497.

Fryer, R. M., J. Randall, T. Yoshida, L. L. Hsiao, J. Blumenstock et al., 2002 Global analysis of gene expression: Methods, interpretation, and pitfalls. Experimental Nephrology 10: 64–74.

Garbe, D. S., Y. Fang, X. Zheng, M. Sowcik, R. Anjum et al., 2012 Cooperative interaction between phosphorylation sites on PERIOD maintains circadian period in Drosophila. PLOS Genetics 9: e1003749–e1003749.

Gekakis, N., L. Saez, A. M. Delahaye-Brown, M. P. Myers, A. Sehgal et al., 1995 Isolation of timeless by PER protein interaction: defective interaction between timeless protein and long-period mutant PERL., pp. 811–815 in Science.

Gerard, C., D. Gonze and A. Goldbeter, 2009 Dependence of the period on the rate of protein degradation in minimal models for circadian oscillations. Philosophical Transactions of the Royal Society A: Mathematical, Physical and Engineering Sciences 367: 4665–4683.

Gibson, R., B. Alako, C. Amid, A. Cerdeno-Tarraga, I. Cleland et al., 2016 Biocuration of functional annotation at the European nucleotide archive. Nucleic Acids Res 44: D58–66.

Gillespie, D. T., 1977 Exact Stochastic Simulation of Coupled Chemical-Reactions. Abstracts of Papers of the American Chemical Society 173: 128–128.

Gillespie, D. T., 2007 Stochastic simulation of chemical kinetics. Annu Rev Phys Chem 58: 35–55.

Gillespie, D. T., 2008 Simulation Methods in Systems Biology. Formal Methods for Computational Systems Biology 5016: 125–167.

Gillespie, D. T., A. Hellander and L. R. Petzold, 2013 Perspective: Stochastic algorithms for chemical kinetics. Journal of Chemical Physics 138: 170901.

Gilliland, G., S. Perrin, K. Blanchard and H. F. Bunn, 1990 Analysis of Cytokine Messenger-Rna and DNA - Detection and Quantitation by Competitive Polymerase Chain-Reaction. Proceedings of the National Academy of Sciences of the United States of America 87: 2725–2729.

Glaser, F. T., and R. Stanewsky, 2005 Temperature synchronization of the Drosophila circadian clock. Current biology: CB 15: 1352–1363.

Gleeson, J. P., J. A. Ward, K. P. O’Sullivan and W. T. Lee, 2014 Competition-induced criticality in a model of meme popularity. Phys Rev Lett 112: 048701.

Glossop, N., J. Houl, H. Zheng, F. Ng, S. Dudek et al., 2003 VRILLE feeds back to control circadian transcription of Clock in the Drosophila circadian oscillator. Neuron 37: 249–261.

Glynn, E. F., J. Chen and A. R. Mushegian, 2006 Detecting periodic patterns in unevenly spaced gene expression time series using Lomb-Scargle periodograms. Bioinformatics 22: 310–316.

Godwin, A. R., S. Kojima, C. B. Green and J. Wilusz, 2013 Kiss your tail goodbye: The role of PARN, Nocturnin, and Angel deadenylases in mRNA biology. Biochimica Et Biophysica Acta-Gene Regulatory Mechanisms 1829: 571–579.

Goldbeter, A., 1995 A Model for Circadian Oscillations in the Drosophila Period Protein (Per). Proceedings. Biological sciences / The Royal Society 261: 319–324.

Goldbeter, A., 2002 Computational approaches to cellular rhythms. Nature 420: 238–245.

Goldston, D., 2008 Big data: Data wrangling. Nature 455: 15–15.

Gonze, D., 2011 Modeling circadian clocks: From equations to oscillations. Central European Journal of Biology 6: 699–711.

Gonze, D., J. Halloy and A. Goldbeter, 2002a Deterministic versus stochastic models for circadian rhythms. Journal of Biological Physics 28: 637–653.

Gonze, D., J. Halloy and A. Goldbeter, 2002b Robustness of circadian rhythms with respect to molecular noise. Proceedings of the National Academy of Sciences of the United States of America 99: 673–678.

Gonze, D., J. Halloy and A. Goldbeter, 2004 Emergence of coherent oscillations in stochastic models for circadian rhythms, pp. 221–233 in Physica a-Statistical Mechanics and Its Applications.

Gonze, D., J. Halloy, J.-C. Leloup and A. Goldbeter, 2003 Stochastic models for circadian rhythms: effect of molecular noise on periodic and chaotic behaviour. Comptes Rendus Biologies 326: 189–203.

Goodwin, B. C., 1964 A statistical mechanics of temporal organization in cells. Symp Soc Exp Biol 18: 301–326.

Goodwin, B. C., 1965 Oscillatory behavior in enzymatic control processes. Adv Enzyme Regul 3: 425–438.

Gramates, L. S., S. J. Marygold, G. D. Santos, J. M. Urbano, G. Antonazzo et al., 2016 FlyBase at 25: looking to the future. Nucleic Acids Res.

Gramates, L. S., S. J. Marygold, G. D. Santos, J. M. Urbano, G. Antonazzo et al., 2017 FlyBase at 25: looking to the future. Nucleic Acids Res 45: D663–D671.

Green, C. B., N. Douris, S. Kojima, C. A. Strayer, J. Fogerty et al., 2007 Loss of Nocturnin, a circadian deadenylase, confers resistance to hepatic steatosis and diet-induced obesity. Proceedings of the National Academy of Sciences of the United States of America 104: 9888–9893.

Grima, B., A. Dognon, A. Lamouroux, E. Chélot and F. Rouyer, 2012 CULLIN-3 Controls TIMELESS Oscillations in the Drosophila Circadian Clock. PLOS biology 10: e1001367.

Grima, B., A. Lamouroux, E. Chélot, C. Papin, B. Limbourg-Bouchon et al., 2002 The F-box protein Slimb controls the levels of clock proteins Period and Timeless. Nature 420: 178–182.

Grimes, M. L., W. J. Lee, L. Van der Maaten and P. Shannon, 2013 Wrangling Phosphoproteomic Data to Elucidate Cancer Signaling Pathways. PLOS ONE 8.

Grönke, S., I. Bickmeyer, R. Wunderlich, H. Jäckle and R. P. Kühnlein, 2009 Curled encodes the Drosophila homolog of the vertebrate circadian deadenylase Nocturnin., pp. 219–232 in Genetics.

Grune, D. e. a., 2012 Modern Compiler Design. Springer Science+Business Media, New York, NY.

Haag-Liautard, C., M. Dorris, X. Maside, S. Macaskill, D. L. Halligan et al., 2007 Direct estimation of per nucleotide and genomic deleterious mutation rates in Drosophila. Nature 445: 82–85.

Halsey, L. G., D. Curran-Everett, S. L. Vowler and G. B. Drummond, 2015 The fickle P value generates irreproducible results. Nature Methods 12: 179–185.

Hamblen, M. J. M., N. E. N. White, P. T. P. Emery, K. K. Kaiser and J. C. J. Hall, 1998 Molecular and behavioral analysis of four period mutants in Drosophila melanogaster encompassing extreme short, novel long, and unorthodox arrhythmic types. Genetics 149: 165–178.

Harbison, S. T., M. A. Carbone, J. F. Ayroles, E. A. Stone, R. F. Lyman et al., 2009a Co-regulated transcriptional networks contribute to natural genetic variation in Drosophila sleep. Nat Genet 41: 371–375.

Harbison, S. T., T. F. Mackay and R. R. Anholt, 2009b Understanding the neurogenetics of sleep: progress from Drosophila. Trends Genet 25: 262–269.

Harbison, S. T., L. J. McCoy and T. F. Mackay, 2013 Genome-wide association study of sleep in Drosophila melanogaster. BMC Genomics 14: 281.

Hardin, P., and J. C. Hall, 1992 Circadian cycling in the levels of protein and mRNA from Drosophila melanogaster’s period gene, pp. 155–169 in Molecular Genetics of Biological Rhythms, edited by M. W. Young.

Hardin, P. E., 1994 Analysis of period mRNA cycling in Drosophila head and body tissues indicates that body oscillators behave differently from head oscillators., pp. 7211–7218 in Mol Cell Biol.

Hardin, P. E., 2011 Molecular Genetic Analysis of Circadian Timekeeping in Drosophila, pp. 141–173. Elsevier.

Hardin, P. E., J. C. Hall and M. Rosbash, 1990 Feedback of the Drosophila period gene product on circadian cycling of its messenger RNA levels. Nature 343: 536–540.

Hardin, P. E., J. C. Hall and M. Rosbash, 1992 Circadian oscillations in period gene mRNA levels are transcriptionally regulated., pp. 11711–11715 in P Natl Acad Sci Usa.

Harms, E., S. Kivimae, M. W. Young and L. Saez, 2004 Posttranscriptional and posttranslational regulation of clock genes. Journal of Biological Rhythms 19: 361–373.

Hazber, M. A. G., R. X. Li, X. W. Gu, G. D. Xu and Y. H. Li, 2015 Semantic SPARQL query in a relational database based on ontology construction. 2015 11th International Conference on Semantics, Knowledge and Grids (Skg): 25–32.

He, S., X. Zheng and D. Zeng, 2016 A Model-Free Scheme for Meme Ranking in Social Media. Decis Support Syst 81: 1–11.

Hermann-Luibl, C., T. Yoshii, P. R. Senthilan, H. Dircksen and C. Helfrich-Förster, 2014 The Ion Transport Peptide Is a New Functional Clock Neuropeptide in the Fruit Fly Drosophila melanogaster. Journal of Neuroscience 34: 9522–9536.

Heyde, C. C., 1997 Quasi-likelihood and its application: a general approach to optimal parameter estimation. Springer, New York.

Hirschman, L., G. A. Burns, M. Krallinger, C. Arighi, K. B. Cohen et al., 2012 Text mining for the biocuration workflow. Database (Oxford) 2012: bas020.

Hoare, C. A. R., 2009 Null References: The Billion Dollar Mistake. Presentation at QCon 2009-08-25: https://www.infoq.com/presentations/Null-References-The-Billion-Dollar-Mistake-Tony-Hoare; https://www.lucidchart.com/techblog/2015/2008/2031/the-worst-mistake-of-computer-science/; https://valbonne-consulting.com/computing-pioneers/tony-hoare-1934/

Hodson, R., 2016 Precision medicine. Nature 537: S49.

Hoffmann, A. A., R. Hallas, C. Sinclair and L. Partridge, 2001 Rapid loss of stress resistance in Drosophila melanogaster under adaptation to laboratory culture. Evolution 55: 436–438.

Hong, C. I., and J. J. Tyson, 1997 A proposal for temperature compensation of the circadian rhythm in Drosophila based on dimerization of the PER protein. Chronobiology International 14: 521–529.

Huang, E. P., X. F. Wang, K. R. Choudhury, L. M. McShane, M. Gonen et al., 2015 Meta-analysis of the technical performance of an imaging procedure: guidelines and statistical methodology. Stat Methods Med Res 24: 141–174.

Huang, Y., and R. Gottardo, 2013 Comparability and reproducibility of biomedical data. Brief Bioinform 14: 391–401.

Hucka, M., A. Finney, H. M. Sauro, H. Bolouri, J. C. Doyle et al., 2003 The systems biology markup language (SBML): a medium for representation and exchange of biochemical network models. Bioinformatics 19: 524–531.

Hughes, M. E. M., G. R. G. Grant, C. C. Paquin, J. J. Qian and M. N. M. Nitabach, 2012 Deep sequencing the circadian and diurnal transcriptome of Drosophila brain. Genes & development 22: 1266–1281.

Hurley, J. M., J. J. Loros and J. C. Dunlap, 2016 The circadian system as an organizer of metabolism. Fungal Genetics and Biology 90: 39–43.

Hut, R. A., S. Paolucci, R. Dor, C. P. Kyriacou and S. Daan, 2013 Latitudinal clines: an evolutionary view on biological rhythms. Proceedings of the Royal Society B-Biological Sciences 280.

Ince, D. C., L. Hatton and J. Graham-Cumming, 2012 The case for open computer programs. Nature 482: 485–488.

Ioannidis, J. P., 2005a Molecular bias. Eur J Epidemiol 20: 739–745.

Ioannidis, J. P., 2005b Why most published research findings are false. PLOS Med 2: e124.

Ioannidis, J. P., D. B. Allison, C. A. Ball, I. Coulibaly, X. Cui et al., 2009a Repeatability of published microarray gene expression analyses. Nat Genet 41: 149–155.

Ioannidis, J. P., G. Thomas and M. J. Daly, 2009b Validating, augmenting and refining genome-wide association signals. Nat Rev Genet 10: 318–329.

Ishikawa, T., A. Matsumoto, T. Kato, S. Togashi, H. Ryo et al., 1999 DCRY is a Drosophila photoreceptor protein implicated in light entrainment of circadian rhythm., pp. 57–65 in Genes Cells.

Itoh, T. Q., T. Tanimura and A. Matsumoto, 2011 Membrane-bound transporter controls the circadian transcription of clock genes in Drosophila., pp. 1159–1167 in Genes Cells.

Janes, A. A., and G. Succi, 2012 The dark side of agile software development, pp. 215–228 in Proceedings of the ACM international symposium on New ideas, new paradigms, and reflections on programming and software. ACM, Tucson, Arizona, USA.

Jang, A. R., K. Moravcevic, L. Saez, M. W. Young and A. Sehgal, 2015 Drosophila TIM binds importin alpha1, and acts as an adapter to transport PER to the nucleus. PLOS Genet 11: e1004974.

Janssens, A. C., A. M. Gonzalez-Zuloeta Ladd, S. Lopez-Leon, J. P. Ioannidis, B. A. Oostra et al., 2009 An empirical comparison of meta-analyses of published gene-disease associations versus consortium analyses. Genet Med 11: 153–162.

Jasny, B. R., G. Chin, L. Chong and S. Vignieri, 2011 Data replication & reproducibility. Again, and again, and again …. Introduction. Science 334: 1225.

Jaynes, E. T., and G. L. Bretthorst, 2003 Probability theory: the logic of science. Cambridge University Press, Cambridge, UK; New York, NY.

Jin, L., 2011 A data-driven test to compare two or multiple time series. Computational Statistics & Data Analysis 55: 2183–2196.

Johnson, R. T., and K. Dickersin, 2007 Publication bias against negative results from clinical trials: three of the seven deadly sins. Nature Clinical Practice Neurology 3: 590–591.

Johnson, V. E., 2013 Revised standards for statistical evidence. Proceedings of the National Academy of Sciences of the United States of America 110: 19313–19317.

Jonquet, C., P. Lependu, S. Falconer, A. Coulet, N. F. Noy et al., 2011 NCBO Resource Index: Ontology-Based Search and Mining of Biomedical Resources. Web Semant 9: 316–324.

Kadener, S., J. S. Menet, K. Sugino, M. D. Horwich, U. Weissbein et al., 2009 A role for microRNAs in the Drosophila circadian clock. Genes Development 23: 2179–2191.

Kadener, S., D. Stoleru, M. McDonald, P. Nawathean and M. Rosbash, 2007 Clockwork Orange is a transcriptional repressor and a new Drosophila circadian pacemaker component. Genes & development 21: 1675–1686.

Kamp, P.-H., 2011 The most expensive one-byte mistake: Did Ken, Dennis, and Brian choose wrong with NUL-terminated text strings? ACM Queue - The Bikeshed 9: 1–4 (comments at http://queue.acm.org/detail.cfm?id=2010365).

Kannan, N. N., K. M. Vaze and V. K. Sharma, 2012 Clock accuracy and precision evolve as a consequence of selection for adult emergence in a narrow window of time in fruit flies Drosophila melanogaster., pp. 3527–3534 in J Exp Biol.

Karlebach, G., and R. Shamir, 2008 Modelling and analysis of gene regulatory networks. Nature Reviews Molecular Cell Biology 9: 770–780.

Karr, J. R., N. C. Phillips and M. W. Covert, 2014 WholeCellSimDB: a hybrid relational/HDF database for whole-cell model predictions. Database (Oxford) 2014.

Karr, J. R., J. C. Sanghvi, D. N. Macklin, A. Arora and M. W. Covert, 2013 WholeCellKB: model organism databases for comprehensive whole-cell models. Nucleic Acids Res 41: D787–792.

Karr, J. R., A. H. Williams, J. D. Zucker, A. Raue, B. Steiert et al., 2015b Summary of the DREAM8 Parameter Estimation Challenge: Toward Parameter Identification for Whole-Cell Models. PLOS Comput Biol 11: e1004096.

Katakura, Y., and H. Ohmori, 2006 Synchronized Control for Circadian Rhythm Based on the Central Clock Model of Drosophila. Audio, Transactions of the IRE Professional Group on: 2358–2361.

Keegan, K. P., S. Pradhan, J. P. Wang and R. Allada, 2007 Meta-analysis of Drosophila circadian microarray studies identifies a novel set of rhythmically expressed genes. PLOS Comput Biol 3: e208.

Keightley, P. D., U. Trivedi, M. Thomson, F. Oliver, S. Kumar et al., 2009 Analysis of the genome sequences of three Drosophila melanogaster spontaneous mutation accumulation lines. Genome Res 19: 1195–1201.

Khodor, Y. L., J. Rodriguez, K. C. Abruzzi, C.-H. A. Tang, M. T. Marr et al., 2011 Nascent-seq indicates widespread cotranscriptional pre-mRNA splicing in Drosophila. Genes & development 25: 2502–2512.

Kidd, P. B., M. W. Young and E. D. Siggia, 2015 Temperature compensation and temperature sensation in the circadian clock. Proc Natl Acad Sci U S A 112: E6284–6292.

Killeen, P. R., 2005 An alternative to null-hypothesis significance tests. Psychol Sci 16: 345–353.

Kim, E. Y., K. Bae, F. S. Ng, N. R. J. Glossop, P. E. Hardin et al., 2002 Drosophila CLOCK protein is under posttranscriptional control and influences light-induced activity., pp. 69–81 in Neuron.

Kim, E. Y., H. W. Ko, W. Yu, P. E. Hardin and I. Edery, 2007 A DOUBLETIME kinase binding domain on the Drosophila PERIOD protein is essential for its hyperphosphorylation, transcriptional repression, and circadian clock function. Molecular and cellular biology 27: 5014–5028.

Kim, S., R. I. Dogan, A. Chatr-Aryamontri, C. S. Chang, R. Oughtred et al., 2016 BioCreative V BioC track overview: collaborative biocurator assistant task for BioGRID. Database-the Journal of Biological Databases and Curation.

Kim, Y. M., J. Choi, H. Y. Lee, G. W. Lee, Y. H. Lee et al., 2014 dbCRY: a Web-based comparative and evolutionary genomics platform for blue-light receptors. Database (Oxford) 2014: bau037.

Kimura, M., 1962 On the probability of fixation of mutant genes in a population. Genetics 47: 713–719.

Kirk, P. D., A. C. Babtie and M. P. Stumpf, 2015 Systems biology (un)certainties. Science 350: 386–388.

Kivimäe, S., L. Saez and M. W. Young, 2008 Activating PER repressor through a DBT-directed phosphorylation switch. PLOS biology 6: e183.

Kloss, B., J. L. Price, L. Saez, J. Blau, A. Rothenfluh et al., 1998 The Drosophila clock gene double-time encodes a protein closely related to human casein kinase Iepsilon. Cell 94: 97–107.

Kloss, B. B., A. A. Rothenfluh, M. W. M. Young and L. L. Saez, 2001 Phosphorylation of PERIOD Is Influenced by Cycling Physical Associations of DOUBLE-TIME, PERIOD, and TIMELESS in the Drosophila Clock. Neuron 30: 8–8.

Knowles, A., K. Koh, J.-T. Wu, C.-T. Chien, D. A. Chamovitz et al., 2009 The COP9 Signalosome Is Required for Light-Dependent Timeless Degradation and Drosophila Clock Resetting. Journal of Neuroscience 29: 1152–1162.

Knuth, D. E., 1974 Structured programming with go to statements. Computing Surveys 6: 261–301.

Ko, H. W., J. Jiang and I. Edery, 2002 Role for Slimb in the degradation of Drosophila Period protein phosphorylated by Doubletime. Nature 420: 673–678.

Ko, H. W., E. Y. Kim, J. Chiu, J. T. Vanselow, A. Kramer et al., 2010 A hierarchical phosphorylation cascade that regulates the timing of PERIOD nuclear entry reveals novel roles for proline-directed kinases and GSK-3beta/SGG in circadian clocks. Journal of Neuroscience 30: 12664–12675.

Koh, K., X. Zheng and A. Sehgal, 2006 JETLAG resets the Drosophila circadian clock by promoting light-induced degradation of TIMELESS. Science 312: 1809–1812.

Konopka, R. J., and S. Benzer, 1971 Clock mutants of Drosophila melanogaster., pp. 2112–2116 in P Natl Acad Sci Usa.

Koskinen, P., P. Toronen, J. Nokso-Koivisto and L. Holm, 2015 PANNZER: high-throughput functional annotation of uncharacterized proteins in an error-prone environment. Bioinformatics 31: 1544–1552.

Kotsifakos, A., V. Athitsos and P. Papapetrou, 2016 Query-sensitive distance measure selection for time series nearest neighbor classification. Intelligent Data Analysis 20: 5–27.

Kraft, P., E. Zeggini and J. P. Ioannidis, 2009 Replication in genome-wide association studies. Stat Sci 24: 561–573.

Krause, F., J. Uhlendorf, T. Lubitz, M. Schulz, E. Klipp et al., 2010 Annotation and merging of SBML models with semanticSBML. Bioinformatics 26: 421–422.

Krishnamurthy, B., W. Willinger, P. Gill and M. Arlitt, 2011 A Socratic method for validation of measurement-based networking research. Computer Communications 34: 43–53.

Kuczenski, R. S., K. C. Hong, J. Garcia-Ojalvo and K. H. Lee, 2007 PERIOD-TIMELESS interval timer may require an additional feedback loop., pp. e154 in PLOS Comput Biol.

Kula-Eversole, E., E. Nagoshi, Y. Shang, J. Rodriguez, R. Allada et al., 2010 Surprising gene expression patterns within and between PDF-containing circadian neurons in Drosophila. Proceedings of the National Academy of Sciences of the United States of America 107: 13497–13502.

Kulasiri, D., and Z. Xie, 2008 Modelling Circadian Rhythms in Drosophila and Investigation of VRI and PDP1 Feedback Loops Using a New Mathematical Model. Mathematical Modelling of Natural Phenomena 3: 1–26.

Kumar, S. S., D. D. Chen and A. A. Sehgal, 2012 Dopamine acts through Cryptochrome to promote acute arousal in Drosophila. Genes & development 26: 1224–1234.

Kurata, H., T. Tanaka and F. Ohnishi, 2007 Mathematical identification of critical reactions in the interlocked feedback model. PLOS ONE 2: e1103–e1103.

Kurosawa, G., A. Mochizuki and Y. Iwasa, 2002 Comparative study of circadian clock models, in search of processes promoting oscillation. Journal of theoretical biology 216: 193–208.

Kurtz, T. G., 1972 Relationship between Stochastic and Deterministic Models for Chemical Reactions. Journal of Chemical Physics 57: 2976-&.

Kyriacou, C. P., A. A. Peixoto and R. Costa, 2007 A cline in the Drosophila melanogaster period gene in Australia: neither down nor under. Journal of Evolutionary Biology 20: 1649–1651.

Kyriacou, C. P., A. A. Peixoto, F. Sandrelli, R. Costa and E. Tauber, 2008 Clines in clock genes: fine-tuning circadian rhythms to the environment. Trends in Genetics 24: 124–132.

Lachance, J., and S. A. Tishkoff, 2013 SNP ascertainment bias in population genetic analyses: why it is important, and how to correct it. Bioessays 35: 780–786.

Laibe, C., and N. Le Novere, 2007 MIRIAM Resources: tools to generate and resolve robust cross-references in Systems Biology. BMC Syst Biol 1: 58.

Lamaze, A. A., A. A. Lamouroux, C. C. Vias, H.-C. H. Hung, F. F. Weber et al., 2011 The E3 ubiquitin ligase CTRIP controls CLOCK levels and PERIOD oscillations in Drosophila. EMBO Reports 12: 549–557.

Lamba, P., D. Bilodeau-Wentworth, P. Emery and Y. Zhang, 2014 Morning and Evening Oscillators Cooperate to Reset Circadian Behavior in Response to Light Input. CellReports 7: 601–608.

Law, A. M., and W. D. Kelton, 2000 Simulation modeling and analysis. McGraw-Hill, Boston.

Le Novere, N., 2015 Quantitative and logic modelling of molecular and gene networks. Nat Rev Genet 16: 146–158.

Le Novere, N., B. Bornstein, A. Broicher, M. Courtot, M. Donizelli et al., 2006 BioModels Database: a free, centralized database of curated, published, quantitative kinetic models of biochemical and cellular systems. Nucleic Acids Res 34: D689–691.

Le Novere, N., M. Hucka, H. Mi, S. Moodie, F. Schreiber et al., 2009 The Systems Biology Graphical Notation. Nature Biotechnology 27: 735–741.

Lear, B. C., L. Zhang and R. Allada, 2009 The neuropeptide PDF acts directly on evening pacemaker neurons to regulate multiple features of circadian behavior. PLOS biology 7: e1000154–e1000154.

Lebiedz, D., M. Rehberg and D. Skanda, 2012 Robust optimal design of synthetic biological networks. Methods Mol Biol 813: 45–55.

Lee, C., K. Bae and I. Edery, 1998 The Drosophila CLOCK protein undergoes daily rhythms in abundance, phosphorylation, and interactions with the PER-TIM complex. Neuron 21: 857–867.

Lee, C., K. Bae and I. Edery, 1999 PER and TIM inhibit the DNA binding activity of a Drosophila CLOCK-CYC/dBMAL1 heterodimer without disrupting formation of the heterodimer: a basis for circadian transcription., pp. 5316–5325 in Mol Cell Biol.

Lee, E., E. H. Jeong, H.-J. Jeong, E. Yildirim, J. T. Vanselow et al., 2014a Phosphorylation of a Central Clock Transcription Factor Is Required for Thermal but Not Photic Entrainment. PLOS Genetics 10: e1004545.

Lee, R., J. R. Karr and M. W. Covert, 2013 WholeCellViz: data visualization for whole-cell models. BMC Bioinformatics 14: 253.

Lee, X. J., C. C. Drovandi and A. N. Pettitt, 2014b Model choice problems using approximate Bayesian computation with applications to pathogen transmission data sets. Biometrics.

Leek, J. T., and R. D. Peng, 2015 Statistics: P values are just the tip of the iceberg. Nature 520: 612.

Leise, T. L., and E. E. Moin, 2007 A mathematical model of the Drosophila circadian clock with emphasis on posttranslational mechanisms. Journal of theoretical biology 248: 48–63.

Lejaeghere, K., G. Bihlmayer, T. Bjorkman, P. Blaha, S. Blugel et al., 2016 Reproducibility in density functional theory calculations of solids. Science 351: aad3000.

Leloup, J.-C., 2009 Circadian clocks and phosphorylation: Insights from computational modeling. Central European Journal of Biology 4: 290–303.

Leloup, J. C., and A. Goldbeter, 1997 Temperature compensation of circadian rhythms: Control of the period in a model for circadian oscillations of the per protein in Drosophila. Chronobiology International 14: 511–520.

Leloup, J. C., and A. Goldbeter, 1998a A model for circadian rhythms in Drosophila incorporating the formation of a complex between the PER and TIM proteins. Journal of biological rhythms 13: 70–87.

Leloup, J. C., and A. Goldbeter, 1998b Modeling circadian oscillations of the PER and TIM proteins in Drosophila. Biological Clocks 1152: 81–88.

Leloup, J. C., and A. Goldbeter, 2000 Modeling the molecular regulatory mechanism of circadian rhythms in Drosophila. Bioessays 22: 84–93.

Leloup, J. C., and A. Goldbeter, 2001 A molecular explanation for the long-term suppression of circadian rhythms by a single light pulse., pp. R1206–1212 in Am J Physiol Regul Integr Comp Physiol.

Leming, M. T., S. S. Rund, S. K. Behura, G. E. Duffield and J. E. O’Tousa, 2014 A database of circadian and diel rhythmic gene expression in the yellow fever mosquito Aedes aegypti. BMC Genomics 15: 1128.

Lerner, I., O. Bartok, V. Wolfson, J. S. Menet, U. Weissbein et al., 2015 Clk post-transcriptional control denoises circadian transcription both temporally and spatially. Nat Commun 6: 7056.

Leroi, A. M., A. K. Chippindale and M. R. Rose, 1994 Long-term laboratory evolution of a genetic life-history trade-off in Drosophila melanogaster. 1. The role of genotype-by-environment interaction. Evolution 48: 1244–1257.

Leugering, G. n., 2012 Constrained optimization and optimal control for partial differential equations. Birkhäuser, Basel Switzerland; New York.

Lew, M. J., 2012 Bad statistical practice in pharmacology (and other basic biomedical disciplines): you probably don’t know P. British Journal of Pharmacology 166: 1559–1567.

Lewis, J., C. E. Breeze, J. Charlesworth, O. J. Maclaren and J. Cooper, 2016 Where next for the reproducibility agenda in computational biology? BMC Syst Biol 10: 52.

Li, Q., and X. Lang, 2008 Internal noise-sustained circadian rhythms in a Drosophila model., pp. 1983–1994 in Biophys J.

Li, S., K. Shui, Y. Zhang, Y. Lv, W. Deng et al., 2017 CGDB: a database of circadian genes in eukaryotes. Nucleic Acids Res 45: D397–D403.

Li, Y., and M. Rosbash, 2013 Accelerated degradation of perS protein provides insight into light-mediated phase shifting., pp. 171–182 in J Biol Rhythms.

Lim, C., B. Y. Chung, J. L. Pitman, J. J. McGill, S. Pradhan et al., 2007 clockwork orange encodes a transcriptional repressor important for circadian-clock amplitude in Drosophila. Current biology: CB 17: 1082–1089.

Lim, C., J. Lee, C. Choi, V. L. Kilman, J. Kim et al., 2011 The novel gene twenty-four defines a critical translational step in the Drosophila clock., pp. 399–403 in Nature.

Lin, J.-M., V. L. Kilman, K. Keegan, B. Paddock, M. Emery-Le et al., 2002a A role for casein kinase 2alpha in the Drosophila circadian clock. Nature 420: 816–820.

Lin, J.-M., A. Schroeder and R. Allada, 2005 In vivo circadian function of casein kinase 2 phosphorylation sites in Drosophila PERIOD. Journal of Neuroscience 25: 11175–11183.

Lin, Y., M. Han, B. Shimada, L. Wang, T. M. Gibler et al., 2002b Influence of the period-dependent circadian clock on diurnal, circadian, and aperiodic gene expression in Drosophila melanogaster. Proceedings of the National Academy of Sciences of the United States of America 99: 9562–9567.

Linné, C. v., 1758 Caroli Linnæi … Systema naturæ per regna tria naturæ, secundum classes, ordines, genera, species, cum characteribus, differentiis, synonymis, locis. impensis L. Salvii, Holmiæ,.

Loewe, L., 2002 Global computing for bioinformatics. Briefings in Bioinformatics 3: 377–388.

Loewe, L., 2007 Evolution@home: observations on participant choice, work unit variation and low-effort global computing. Software Practice & Experience 37: 1289–1318.

Loewe, L., 2009 A framework for evolutionary systems biology. BMC Systems Biology 3: 27.

Loewe, L., 2012 How evolutionary systems biology will help understand adaptive landscapes and distributions of mutational effects, pp. 399–410 in Evolutionary Systems Biology, edited by O. Soyer. Springer, New York, NY.

Loewe, L., and J. Hillston, 2008 The distribution of mutational effects on fitness in a simple circadian clock. Lecture Notes in Bioinformatics 5307: 156–175.

Loewe, L., and S. Keel, 2014 BEST Names for Semantic Units to Support Reproducible Modeling, pp. 4 pages, https://www.xsede.org/documents/659353/703287/xsede659314_loewe.pdf in 2014-07-14 Workshop: "reproducibility@XSEDE", XSEDE14 Annual Conference (https://www.xsede.org/web/reproducibility), Atlanta, GA, USA.

Loewe, L., S. Moodie and J. Hillston, 2009a Quantifying the implicit process flow abstraction in SBGN-PD diagrams with Bio-PEPA. EPTCS 6: 93–107 http://arxiv.org/abs/0910.1410.

Loewe, L., S. Moodie and J. Hillston, 2009b Technical Report: Defining a textual representation for SBGN Process Diagrams and translating it to Bio-PEPA for quantitative analysis of the MAPK signal transduction cascade. Technical Report, School for Informatics, University of Edinburgh EDI-INF-RR-1334: http://www.inf.ed.ac.uk/publications/report/1334.html

Loftus, G. R., 1993 A Picture Is Worth 1000 P-Values - on the Irrelevance of Hypothesis-Testing in the Microcomputer Age. Behavior Research Methods Instruments & Computers 25: 250–256.

Long, D. M., M. R. Blake, S. Dutta, S. D. Holbrook, J. Kotwica-Rolinska et al., 2014 Relationships between the circadian system and Alzheimer’s disease-like symptoms in Drosophila. PLOS ONE 9: e106068.

Lopes, R. d. S., N. M. Resende, A. C. Honorio-Franca and E. L. Franca, 2013 Application of bioinformatics in chronobiology research. ScientificWorldJournal 2013: 153839.

Lorenz, L. J., J. C. Hall and M. Rosbash, 1989 Expression of a Drosophila mRNA is under circadian clock control during pupation., pp. 869–880 in Development.

Lorsch, J. R., F. S. Collins and J. Lippincott-Schwartz, 2014 Cell Biology. Fixing problems with cell lines. Science 346: 1452–1453.

Lowrey, P. L., and J. S. Takahashi, 2004 Mammalian circadian biology: elucidating genome-wide levels of temporal organization. Annu Rev Genomics Hum Genet 5: 407–441.

Lynch, A., G. M. Plunkett, A. J. Baker and P. F. Jenkins, 1989 A Model of Cultural-Evolution of Chaffinch Song Derived with the Meme Concept. American Naturalist 133: 634–653.

Lynch, M., M. C. Field, H. V. Goodson, H. S. Malik, J. B. Pereira-Leal et al., 2014 Evolutionary cell biology: Two origins, one objective. Proceedings of the National Academy of Sciences of the United States of America 111: 16990–16994.

Ma, L., J. Ma and K. Xu, 2015 Effect of Spaceflight on the Circadian Rhythm, Lifespan and Gene Expression of Drosophila melanogaster., pp. e0121600 in PLOS ONE.

Mackay, T. F., S. Richards, E. A. Stone, A. Barbadilla, J. F. Ayroles et al., 2012 The Drosophila melanogaster Genetic Reference Panel. Nature 482: 173–178.

Macneil, L. T., and A. J. Walhout, 2011 Gene regulatory networks and the role of robustness and stochasticity in the control of gene expression. Genome Res 21: 645–657.

Majercak, J., W.-F. Chen and I. Edery, 2004 Splicing of the period gene 3’-terminal intron is regulated by light, circadian clock factors, and phospholipase C., pp. 3359–3372 in Mol Cell Biol.

Majercak, J., D. Kalderon and I. Edery, 1997 Drosophila melanogaster deficient in protein kinase A manifests behavior-specific arrhythmia but normal clock function.

Majercak, J., D. Sidote, P. E. Hardin and I. Edery, 1999 How a circadian clock adapts to seasonal decreases in temperature and day length., pp. 219–230 in Neuron.

Major, P., B. J. Kostrewski and J. Anderson, 1978 Analysis of the semantic structures of medical reference languages: part 2. Analysis of the semantic power of MeSH, ICD and SNOMED. Med Inform (Lond) 3: 269–281.

Malinen, E., A. Kassinen, T. Rinttila and A. Palva, 2003 Comparison of real-time PCR with SYBR Green I or 5’-nuclease assays and dot-blot hybridization with rDNA-targeted oligonucleotide probes in quantification of selected faecal bacteria. Microbiology 149: 269–277.

Mandrioli, D., and M. Pradella, 2015 Programming Languages shouldn’t be "too Natural". ACM SIGSOFT Software Engineering Notes 40: 1–4.

Marrus, S. B., H. Zeng and M. Rosbash, 1996 Effect of constant light and circadian entrainment of perS flies: evidence for light-mediated delay of the negative feedback loop in Drosophila. EMBO Journal 15: 6877–6886.

Martinek, S., S. Inonog, A. S. Manoukian and M. W. Young, 2001 A role for the segment polarity gene shaggy/GSK-3 in the Drosophila circadian clock. Cell 105: 769–779.

Marygold, S. J., M. A. Crosby, J. L. Goodman and C. FlyBase, 2016 Using FlyBase, a Database of Drosophila Genes and Genomes. Methods Mol Biol 1478: 1–31.

Masri, S., K. Kinouchi and P. Sassone-Corsi, 2015 Circadian clocks, epigenetics, and cancer. Current Opinion in Oncology 27: 50–56.

Matsumoto, A., M. Ukai-Tadenuma, R. G. Yamada, J. Houl, K. D. Uno et al., 2007 A functional genomics strategy reveals clockwork orange as a transcriptional regulator in the Drosophila circadian clock. Genes & development 21: 1687–1700.

Mavelli, F., 2012 Stochastic simulations of minimal cells: the Ribocell model. BMC Bioinformatics 13 Suppl 4: S10.

Mazumdar, M., S. Banerjee and H. L. Van Epps, 2010 Improved reporting of statistical design and analysis: guidelines, education, and editorial policies. Methods Mol Biol 620: 563–598.

McClung, C. A., 2013 How Might Circadian Rhythms Control Mood? Let Me Count the Ways.…. Biological Psychiatry.

McDonald, M. J., and M. Rosbash, 2001 Microarray analysis and organization of circadian gene expression in Drosophila. Cell 107: 567–578.

McDonald, M. J., M. Rosbash and P. Emery, 2001 Wild-type circadian rhythmicity is dependent on closely spaced E boxes in the Drosophila timeless promoter. Mol Cell Biol 21: 1207–1217.

McIntyre, L. M., K. K. Lopiano, A. M. Morse, V. Amin, A. L. Oberg et al., 2011 RNA-seq: technical variability and sampling. Bmc Genomics 12.

McKenzie, J. E., G. Salanti, S. C. Lewis and D. G. Altman, 2013 Meta-analysis and The Cochrane Collaboration: 20 years of the Cochrane Statistical Methods Group. Syst Rev 2: 80.

McNutt, M., 2014 Journals unite for reproducibility. Science 346: 679.

Meissner, R.-A., V. L. Kilman, J.-M. Lin and R. Allada, 2008 TIMELESS is an important mediator of CK2 effects on circadian clock function in vivo. Journal of Neuroscience 28: 9732–9740.

Menet, J. S., K. C. Abruzzi, J. Desrochers, J. Rodriguez and M. Rosbash, 2010 Dynamic PER repression mechanisms in the Drosophila circadian clock: from on-DNA to off-DNA. Genes & development 24: 358–367.

Merbitz-Zahradnik, T., and E. Wolf, 2015 How is the inner circadian clock controlled by interactive clock proteins?: Structural analysis of clock proteins elucidates their physiological role. FEBS Lett 589: 1516–1529.

Merrill, G. H., 2009 Concepts and synonymy in the UMLS Metathesaurus. J Biomed Discov Collab 4: 7.

Mesnard, O., and L. A. Barba, 2016 Reproducible and replicable CFD: it’s harder than you think. Computing in Science and Engineering: https://arxiv.org/abs/1605.04339.

Mezan, S., R. Ashwal-Fluss, R. Shenhav, M. Garber and S. Kadener, 2013 Genome-wide assessment of post-transcriptional control in the fly brain., pp. 49 in Front Mol Neurosci.

Millis, S. R., 2003 Statistical practices: The seven deadly sins. Child Neuropsychology 9: 221–233.

Minikel, E. V., I. Zerr, S. J. Collins, C. Ponto, A. Boyd et al., 2014 Ascertainment bias causes false signal of anticipation in genetic prion disease. Am J Hum Genet 95: 371–382.

Mitchell, C. S., A. Cates, R. B. Kim and S. K. Hollinger, 2015 Undergraduate Biocuration: Developing Tomorrow’s Researchers While Mining Today’s Data. J Undergrad Neurosci Educ 14: A56–65.

Mitra, K., A. R. Carvunis, S. K. Ramesh and T. Ideker, 2013 Integrative approaches for finding modular structure in biological networks. Nature Reviews Genetics 14: 719–732.

Miura, S., T. Shimokawa and T. Nomura, 2008 Stochastic simulations on a model of circadian rhythm generation. Bio Systems 93: 133–140.

Mobley, A., S. K. Linder, R. Braeuer, L. M. Ellis and L. Zwelling, 2013 A Survey on Data Reproducibility in Cancer Research Provides Insights into Our Limited Ability to Translate Findings from the Laboratory to the Clinic. PLOS ONE 8.

Mockler, T. C., T. P. Michael, H. D. Priest, R. Shen, C. M. Sullivan et al., 2007 The DIURNAL project: DIURNAL and circadian expression profiling, model-based pattern matching, and promoter analysis. Cold Spring Harb Symp Quant Biol 72: 353–363.

Molina-Rodríguez, M., and V. Álvarez, 2016 Los ritmos circadianos en cancer y la cronoterapia. Iatreia 29.3: 301.

Moodie, S., S. Moodie, N. Le Novere, E. Demir, H. Mi et al., 2011 Systems Biology Graphical Notation: Process Description language Level 1. Nature Precedings.

Mori, U., A. Mendiburu and J. A. Lozano, 2016 Similarity Measure Selection for Clustering Time Series Databases. Ieee Transactions on Knowledge and Data Engineering 28: 181–195.

Mortazavi, A., B. A. Williams, K. Mccue, L. Schaeffer and B. Wold, 2008 Mapping and quantifying mammalian transcriptomes by RNA-Seq. Nature Methods 5: 621–628.

Moura Neto, F. D., and A. n. J. Silva Neto, 2013 An introduction to inverse problems with applications. Springer, Heidelberg; New York.

Murtaugh, P. A., 2014 In defense of P values. Ecology 95: 611–617.

Musen, M. A., N. F. Noy, N. H. Shah, P. L. Whetzel, C. G. Chute et al., 2012 The National Center for Biomedical Ontology. J Am Med Inform Assoc 19: 190–195.

Muskus, M. J., F. Preuss, J.-Y. Fan, E. S. Bjes and J. L. Price, 2007 Drosophila DBT lacking protein kinase activity produces long-period and arrhythmic circadian behavioral and molecular rhythms., pp. 8049–8064 in Mol Cell Biol.

Myers, M. P., K. Wager-Smith, A. Rothenfluh-Hilfiker and M. W. Young, 1996 Light-Induced Degradation of TIMELESS and Entrainment of the Drosophila Circadian Clock. Science, New Series 271: 1736–1740.

Nagoshi, E., K. Sugino, E. Kula, E. Okazaki, T. Tachibana et al., 2010 Dissecting differential gene expression within the circadian neuronal circuit of Drosophila., pp. 60–68 in Nat Neurosci.

Naidoo, N., W. Song, M. Hunter-Ensor and A. Sehgal, 1999 A role for the proteasome in the light response of the timeless clock protein. Science 285: 1737–1741.

Najafi, A., G. Bidkhori, J. H. Bozorgmehr, I. Koch and A. Masoudi-Nejad, 2014 Genome scale modeling in systems biology: algorithms and resources. Curr Genomics 15: 130–159.

Nakagawa, S., and I. C. Cuthill, 2007 Effect size, confidence interval and statistical significance: a practical guide for biologists. Biological Reviews 82: 591–605.

NCI-NHGRI Working Group on Replication in Association Studies, S. J. Chanock, T. Manolio, M. Boehnke, E. Boerwinkle et al., 2007 Replicating genotype-phenotype associations. Nature 447: 655–660.

Ng, A., B. Bursteinas, Q. Gao, E. Mollison and M. Zvelebil, 2006 Resources for integrative systems biology: from data through databases to networks and dynamic system models. Brief Bioinform 7: 318–330.

NIH, 2015 National Institutes of Health Plan for Increasing Access to Scientific Publications and Digital Scientific Data from NIH Funded Scientific Research. 2015-02: https://grants.nih.gov/grants/NIH-Public-Access-Plan.pdf.

NIH, 2016 Strategies for NIH Data Management, Sharing, and Citation. Request for Information (RFI) Notice Number NOT-OD-17-015: http://grants.nih.gov/grants/guide/notice-files/NOT-OD-17-015.html.

NIH, Blue Ribbon Panel, Tyler Jacks, E. Jaffee, D. Singer et al., 2016 Cancer Moonshot Blue Ribbon Panel Report 2016-10-17, pp. https://www.cancer.gov/research/key-initiatives/moonshot-cancer-initiative/blue-ribbon-panel/blue-ribbon-panel-report-2016.pdf. National Institutes of Health,.

Nordstokke, D. W., and B. D. Zumbo, 2007 A Cautionary Tale About Levene’s Tests for Equal Variances. Journal of Educational Research & Policy Studies 7: 1–14.

Nuzzo, R., 2014 Scientific method: statistical errors. Nature 506: 150–152.

Oconnell, P., and M. Rosbash, 1984 Sequence, Structure, and Codon Preference of the Drosophila Ribosomal Protein-49 Gene. Nucleic Acids Research 12: 5495–5513.

Ogawa, Y., K. Arakawa, K. Kaizu, F. Miyoshi, Y. Nakayama et al., 2008 Comparative study of circadian oscillatory network models of Drosophila. Artificial life 14: 29–48.

Oh, Y., S.-E. Yoon, Q. Zhang, H.-S. Chae, I. Daubnerová et al., 2014 A homeostatic sleep-stabilizing pathway in Drosophila composed of the sex peptide receptor and its ligand, the myoinhibitory peptide., pp. e1001974 in PLOS Biol.

Okada, T., T. Sakai, T. Murata, K. Kako, K. Sakamoto et al., 2001 Promoter analysis for daily expression of Drosophila timeless gene. Biochemical and Biophysical Research Communications 283: 577–582.

Oliver, D., and J. Phillips, 1970 Technical Note. Drosophila Information Service.

Onitilo, A. A., 2014 Is it time for the Cochrane Collaboration to reconsider its meta-analysis methodology? Clin Med Res 12: 2–3.

Opitz, J. M., 1983 Cotterman and combinatorial genetics. Am J Med Genet 16: 389–392.

Oreskes, N., K. Shraderfrechette and K. Belitz, 1994 Verification, Validation, and Confirmation of Numerical-Models in the Earth-Sciences. Science 263: 641–646.

Orpana, A. K., T. H. Ho and J. Stenman, 2012 Multiple Heat Pulses during PCR Extension Enabling Amplification of GC-Rich Sequences and Reducing Amplification Bias. Analytical Chemistry 84: 2081–2087.

Özkaya, Ö., and E. Rosato, 2012 The circadian clock of the fly: a neurogenetics journey through time., pp. 79–123 in Adv Genet.

Ozturk, N. N., C. P. C. Selby, Y. Y. Annayev, D. D. Zhong and A. A. Sancar, 2011 Reaction mechanism of Drosophila cryptochrome. Proceedings of the National Academy of Sciences of the United States of America 108: 516–521.

Packer, M., 2016 Data sharing: lessons from Copernicus and Kepler. BMJ 354: i4911.

Panko, R. R., 2013 Users and Uses Research: Understanding "The Familiar Unknown". Journal of Organizational and End User Computing 25: Iv–Xi.

Parekh, P. K., A. R. Ozburn and C. A. McClung, 2015 Circadian clock genes: effects on dopamine, reward and addiction. Alcohol 49: 341–349.

Patel, V. R., K. Eckel-Mahan, P. Sassone-Corsi and P. Baldi, 2012 CircadiOmics: integrating circadian genomics, transcriptomics, proteomics and metabolomics. Nature Methods 9: 772–773.

Patil, P., P. O. Bachant-Winner, B. Haibe-Kains and J. T. Leek, 2015 Test set bias affects reproducibility of gene signatures. Bioinformatics 31: 2318–2323.

Pegoraro, M., S. Noreen, S. Bhutani, A. Tsolou, R. Schmid et al., 2014 Molecular evolution of a pervasive natural amino-acid substitution in Drosophila cryptochrome. PLOS ONE 9: e86483.

Pembroke, W. G., A. Babbs, K. E. Davies, C. P. Ponting and P. L. Oliver, 2015 Temporal transcriptomics suggest that twin-peaking genes reset the clock. Elife 4.

Peschel, N., 2008 New insights into circadian photoreception and the molecular regulation of the resetting of Drosophilas circadian clock. 1–163.

Peschel, N., K. F. Chen, G. Szabo and R. Stanewsky, 2009 Light-dependent interactions between the Drosophila circadian clock factors cryptochrome, jetlag, and timeless. Curr Biol 19: 241–247.

Peschel, N., and C. Helfrich-Förster, 2011 Setting the clock – by nature: Circadian rhythm in the fruitfly Drosophila melanogaster. Febs Letters 585: 1435–1442.

Petri, B., and M. Stengl, 2001 Phase response curves of a molecular model oscillator: implications for mutual coupling of paired oscillators., pp. 125–141 in J Biol Rhythms.

Pierce, B. C., 2002 Types and programming languages. MIT Press, Cambridge, Mass.

Pierce, B. C., 2005 Advanced topics in types and programming languages. MIT Press, Cambridge, Mass.

Pittendrigh, C., and G. Victor, 1957 An Oscillator Model for Biological Clocks in Rhythmic and Synthetic Processes in Growth, edited by D. Rudnick. Princeton University Press, Priceton, NJ.

Pizarro, A., K. Hayer, N. F. Lahens and J. B. Hogenesch, 2013 CircaDB: a database of mammalian circadian gene expression profiles. Nucleic Acids Res 41: D1009–1013.

Plautz, J. D., M. Straume, R. Stanewsky, C. F. Jamison, C. Brandes et al., 1997 Quantitative analysis of Drosophila period gene transcription in living animals. Journal of biological rhythms 12: 204–217.

Plowman, A. B., 2008 BIAZA statistics guidelines: toward a common application of statistical tests for zoo research. Zoo Biol 27: 226–233.

Pogue-Geile, K. L., J. Lyons-Weiler and D. C. Whitcomb, 2006 Molecular overlap of fly circadian rhythms and human pancreatic cancer. Cancer Letters 243: 55–57.

Preussner, M., and F. Heyd, 2016 Post-transcriptional control of the mammalian circadian clock: implications for health and disease. Pflugers Archiv-European Journal of Physiology 468: 983–991.

Price, J. L., J. Blau, A. Rothenfluh, M. Abodeely, B. Kloss et al., 1998 double-time Is a Novel Drosophila Clock Gene that Regulates PERIOD Protein Accumulation. Cell 94: 83–95.

Purcell, O., B. Jain, J. R. Karr, M. W. Covert and T. K. Lu, 2013 Towards a whole-cell modeling approach for synthetic biology. Chaos 23: 025112.

Qiu, J., and P. E. Hardin, 1996 per mRNA cycling is locked to lights-off under photoperiodic conditions that support circadian feedback loop function., pp. 4182–4188 in Mol Cell Biol.

Qu, Y., and M. Boutjdir, 2007 RNase Protection Assay for Quantifying Gene Expression Levels in Methods in Molecular Biology - Cardian Gene Expression: Methods and Protocols, edited by J. Zhang and G. Rokosh. Humana Press Inc., Totowa, N.J.

Raymond, E. S., 2003 The Art of Unix Programming. Addison-Wesley (Pearson), Boston, MA, USA. http://www.catb.org/~esr/writings/taoup/.

Read, K. B., J. R. Sheehan, M. F. Huerta, L. S. Knecht, J. G. Mork et al., 2015 Sizing the Problem of Improving Discovery and Access to NIH-Funded Data: A Preliminary Study. PLOS ONE 10: e0132735.

Reason, J. T., 1990 Human error. Cambridge University Press, Cambridge England; New York.

Reason, J. T., 2015 Organizational accidents revisited. Ashgate, Burlington, VT.

Refinetti, R., G. C. Lissen and F. Halberg, 2007 Procedures for numerical analysis of circadian rhythms. Biol Rhythm Res 38: 275–325.

Richier, B., C. Michard-Vanhee, A. Lamouroux, C. Papin and F. Rouyer, 2008 The clockwork orange Drosophila protein functions as both an activator and a repressor of clock gene expression. J Biol Rhythms 23: 103–116.

Risau-Gusman, S., and P. M. Gleiser, 2012 Modelling the effect of phosphorylation on the circadian clock of Drosophila. Journal of theoretical biology 307: 53–61.

Risau-Gusman, S., and P. M. Gleiser, 2014 A Mathematical Model of Communication between Groups of Circadian Neurons in Drosophila melanogaster. Journal of biological rhythms 29: 401–410.

Robert, C., and M. Watson, 2015 Errors in RNA-Seq quantification affect genes of relevance to human disease. Genome Biol 16: 177.

Robert, C. P., J. M. Cornuet, J. M. Marin and N. S. Pillai, 2011 Lack of confidence in approximate Bayesian computation model choice. Proc Natl Acad Sci U S A 108: 15112–15117.

Roberts, A., L. Schaeffer and L. Pachter, 2013 Updating RNA-Seq analyses after re-annotation. Bioinformatics 29: 1631–1637.

Robinson, S. W., P. Herzyk, J. A. Dow and D. P. Leader, 2013 FlyAtlas: database of gene expression in the tissues of Drosophila melanogaster. Nucleic Acids Res 41: D744–750.

Roden, D. M., 2011 Personalized medicine and the genotype-phenotype dilemma. J Interv Card Electrophysiol 31: 17–23.

Rodriguez, J., C.-H. A. Tang, Y. L. Khodor, S. Vodala, J. S. Menet et al., 2013 Nascent-Seq analysis of Drosophila cycling gene expression. Proceedings of the National Academy of Sciences of the United States of America 110: E275–E284.

Roenneberg, T., and M. Merrow, 1998 Molecular circadian oscillators: An alternative hypothesis. Journal of Biological Rhythms 13: 167–179.

Rosato, E., A. A. Peixoto, A. Gallippi, C. P. Kyriacou and R. Costa, 1996 Mutational mechanisms, phylogeny, and evolution of a repetitive region within a clock gene of Drosophila melanogaster., pp. 392–408 in J Mol Evol.

Rosato, E., E. Tauber and C. P. Kyriacou, 2006 Molecular genetics of the fruit-fly circadian clock. European Journal of Human Genetics 14: 729–738.

Rothenfluh, A., M. W. Young and L. Saez, 2000a A TIMELESS-independent function for PERIOD proteins in the Drosophila clock. Neuron 26: 505–514.

Rothenfluh, A. A., M. M. Abodeely, J. L. J. Price and M. W. M. Young, 2000b Isolation and analysis of six timeless alleles that cause short- or long-period circadian rhythms in Drosophila. Genetics 156: 665–675.

Rouyer, F., M. Rachidi, C. Pikielny and M. Rosbash, 1997 A new gene encoding a putative transcription factor regulated by the Drosophila circadian clock., pp. 3944–3954 in EMBO J.

Rubinson, C., 2014 Ontological Considerations When Modeling Missing Data With Relational Databases. Social Science Computer Review 32: 769–780.

Rund, S. S., J. E. Gentile and G. E. Duffield, 2013 Extensive circadian and light regulation of the transcriptome in the malaria mosquito Anopheles gambiae. BMC Genomics 14: 218.

Rund, S. S., T. Y. Hou, S. M. Ward, F. H. Collins and G. E. Duffield, 2011 Genome-wide profiling of diel and circadian gene expression in the malaria vector Anopheles gambiae. Proc Natl Acad Sci U S A 108: E421–430.

Ruoff, P., M. K. Christensen and V. K. Sharma, 2005 PER/TIM-mediated amplification, gene dosage effects and temperature compensation in an interlocking-feedback loop model of the Drosophila circadian clock. Journal of theoretical biology 237: 41–57.

Ruoff, P., and L. Rensing, 1996 The temperature-compensated goodwin model simulates many circadian clock properties. Journal of Theoretical Biology 179: 275–285.

Ruoff, P., L. Rensing, R. Kommedal and S. Mohsenzadeh, 1997 Modeling temperature compensation in chemical and biological oscillators. Chronobiology International 14: 499–510.

Ruoff, P., M. Vinsjevik, C. Monnerjahn and L. Rensing, 1999 The Goodwin Oscillator: On the Importance of Degradation Reactions in the Circadian Clock. Journal of biological rhythms 14: 469–479.

Rutila, J. E., V. Suri, M. Le, W. V. So, M. Rosbash et al., 1998 CYCLE is a second bHLH-PAS clock protein essential for circadian rhythmicity and transcription of Drosophila period and timeless. Cell 93: 805–814.

Saez-Rodriguez, J., J. C. Costello, S. H. Friend, M. R. Kellen, L. Mangravite et al., 2016 Crowdsourcing biomedical research: leveraging communities as innovation engines. Nature Reviews Genetics 17: 470–486.

Salanti, G., and J. P. Ioannidis, 2009 Synthesis of observational studies should consider credibility ceilings. J Clin Epidemiol 62: 115–122.

Salavaty, A., 2015 Carcinogenic effects of circadian disruption: an epigenetic viewpoint. Chinese Journal of Cancer 34.

Salimi, N., and R. Vita, 2006 The biocurator: Connecting and enhancing scientific data. PLOS Computational Biology 2: 1190–1192.

Salsburg, D. S., 1985 The Religion of Statistics as Practiced in Medical Journals. American Statistician 39: 220–223.

Sambrook, J., E. Fritsch and T. Maniatis, 1989 Molecular Cloning: A Laboratory Manual. Cold Spring Harbor Laboratory Press, Cold Spring Harbor, New York.

Samuels, S., B. Balint, H. von der Leyen, P. Hupe, L. de Koning et al., 2016 Precision medicine in cancer: challenges and recommendations from an EU-funded cervical cancer biobanking study. Br J Cancer.

Sandrelli, F., E. Tauber, M. Pegoraro, G. Mazzotta, P. Cisotto et al., 2007 A molecular basis for natural selection at the timeless locus in Drosophila melanogaster. Science 316: 1898–1900.

Sanghvi, J. C., S. Regot, S. Carrasco, J. R. Karr, M. V. Gutschow et al., 2013 Accelerated discovery via a whole-cell model. Nat Methods 10: 1192–1195.

Sathyanarayanan, S., X. Zheng, R. Xiao and A. Sehgal, 2004 Posttranslational regulation of Drosophila PERIOD protein by protein phosphatase 2A. Cell 116: 603–615.

Savalei, V., and E. Dunn, 2015 Is the call to abandon p-values the red herring of the replicability crisis? Frontiers in Psychology 6.

Sawyer, L. A., J. M. Hennessy, A. A. Peixoto, E. Rosato, H. Parkinson et al., 1997 Natural variation in a Drosophila clock gene and temperature compensation., pp. 2117–2120 in Science.

Scheper, T. O., D. Klinkenberg, C. Pennartz and J. van Pelt, 1999a A mathematical model for the intracellular circadian rhythm generator. Journal of Neuroscience 19: 40–47.

Scheper, T. O. T., D. D. Klinkenberg, J. J. van Pelt and C. C. Pennartz, 1999b A model of molecular circadian clocks: multiple mechanisms for phase shifting and a requirement for strong nonlinear interactions. Journal of biological rhythms 14: 213–220.

Schnoes, A. M., S. D. Brown, I. Dodevski and P. C. Babbitt, 2009 Annotation error in public databases: misannotation of molecular function in enzyme superfamilies. PLOS Comput Biol 5: e1000605.

Schutt, R., and C. O’Neil, 2013 Doing data science. O’Reilly Media, Beijing; Sebastopol.

Schwartz, B., 2004 The paradox of choice: why more is less. Ecco, New York.

Scribner, E. Y., and H. M. Fathallah-Shaykh, 2011 Modeling of regulatory networks: theory and applications in the study of the Drosophila circadian clock. Methods in enzymology 487: 39–71.

Sehgal, A., J. L. Price, B. Man and M. W. Young, 1994 Loss of Circadian Behavioral Rhythms and per RNA Oscillations in the Drosophila Mutant timeless. Science, New Series 263: 1603–1606.

Sehgal, A., A. Rothenfluh-Hilfiker, M. Hunter-Ensor, Y. Chen, M. P. Myers et al., 1995 Rhythmic expression of timeless: a basis for promoting circadian cycles in period gene autoregulation., pp. 808–810 in Science.

Sephton, S., and D. Spiegel, 2003 Circadian disruption in cancer: a neuroendocrine-immune pathway from stress to disease? Brain Behavior and Immunity 17: 321–328.

SEQC/MAQC-III Consortium, 2014 A comprehensive assessment of RNA-seq accuracy, reproducibility and information content by the Sequencing Quality Control Consortium. Nat Biotechnol 32: 903–914.

Shabalina, S. A., L. Y. Yampolsky and A. S. Kondrashov, 1997 Rapid decline of fitness in panmictic populations of Drosophila melanogaster maintained under relaxed natural selection. Proceedings Of the National Academy Of Sciences Of the United States Of America 94: 13034–13039.

Shafer, O. T., M. Rosbash and J. W. Truman, 2002 Sequential nuclear accumulation of the clock proteins period and timeless in the pacemaker neurons of Drosophila melanogaster. Journal of Neuroscience 22: 5946–5954.

Sharkey, F. H., I. M. Banat and R. Marchant, 2004 Detection and quantification of gene expression in environmental bacteriology. Applied and Environmental Microbiology 70: 3795–3806.

Sharma, A., S. Tiwari and M. Singaravel, 2016 Circadian rhythm disruption: health consequences. Biological Rhythm Research 47: 191–213.

Sharma, V. K., 2003 Adaptive significance of circadian clocks. Chronobiology International 20: 901–919.

Sharpe, D., 2013 Why the Resistance to Statistical Innovations? Bridging the Communication Gap. Psychological Methods 18: 572–582.

Shenk, D., 1997 Data smog: surviving the information glut. Harper Edge, San Francisco, Calif.

Shi, L. M., L. H. Reid, W. D. Jones, R. Shippy, J. A. Warrington et al., 2006 The MicroArray Quality Control (MAQC) project shows inter- and intraplatform reproducibility of gene expression measurements. Nature Biotechnology 24: 1151–1161.

Shi, M., Z. Yue, A. Kuryatov, J. M. Lindstrom, A. Sehgal et al., 2014 Identification of Redeye, a new sleep-regulating protein whose expression is modulated by sleep amount. eLife 3: e01473.

Sidote, D., J. Majercak, V. Parikh and I. Edery, 1998 Differential effects of light and heat on the Drosophila circadian clock proteins PER and TIM. Molecular and Cellular Biology 18: 2004–2013.

Simoni, A., W. Wolfgang, M. P. Topping, R. G. Kavlie, R. Stanewsky et al., 2014 A mechanosensory pathway to the Drosophila circadian clock., pp. 525–528 in Science.

Smolen, P., 2002 A detailed predictive model of the mammalian circadian clock.

Smolen, P., D. Baxter and J. Byrne, 2002 A Reduced Model Clarifies the Role of Feedback Loops and Time Delays in the Drosophila Circadian Oscillator. Biophysical Journal 83: 2349–2359.

Smolen, P., D. A. Baxter and J. H. Byrne, 2000a Mathematical modeling of gene networks. Neuron 26: 567–580.

Smolen, P., D. A. Baxter and J. H. Byrne, 2000b Modeling transcriptional control in gene networks - Methods, recent results, and future directions. Bulletin of Mathematical Biology 62: 247–292.

Smolen, P., D. A. Baxter and J. H. Byrne, 2001 Modeling circadian oscillations with interlocking positive and negative feedback loops. Journal of Neuroscience 21: 6644–6656.

Smolen, P., P. E. Hardin, B. S. Lo, D. A. Baxter and J. H. Byrne, 2004 Simulation of Drosophila circadian oscillations, mutations, and light responses by a model with VRI, PDP-1, and CLK. Biophysical Journal 86: 2786–2802.

So, W. V., and M. Rosbash, 1997 Post-transcriptional regulation contributes to Drosophila clock gene mRNA cycling. EMBO Journal 16: 7146–7155.

Sonawani, A., P. Nilawe, R. S. Barai and S. Idicula-Thomas, 2010 ClotBase: a knowledgebase on proteins involved in blood coagulation. Blood 116: 855–856.

Soranno, P. A., E. G. Bissell, K. S. Cheruvelil, S. T. Christel, S. M. Collins et al., 2015 Building a multi-scaled geospatial temporal ecology database from disparate data sources: fostering open science and data reuse. Gigascience 4: 28.

Soubeyrand, S., F. Carpentier, F. Guiton and E. K. Klein, 2013 Approximate Bayesian computation with functional statistics. Statistical Applications in Genetics and Molecular Biology 12.

St Pierre, S., and P. McQuilton, 2009 Inside FlyBase: biocuration as a career. Fly (Austin) 3: 112–114.

Stainforth, D. A., T. Aina, C. Christensen, M. Collins, N. Faull et al., 2005 Uncertainty in predictions of the climate response to rising levels of greenhouse gases. Nature 433: 403–406.

Stajich, J. E., D. Block, K. Boulez, S. E. Brenner, S. A. Chervitz et al., 2002 The Bioperl toolkit: Perl modules for the life sciences. Genome Res 12: 1611–1618.

Stanewsky, R., 2003 Genetic analysis of the circadian system in Drosophila melanogaster and mammals. Journal of neurobiology 54: 111–147.

Stanewsky, R., M. Kaneko, P. Emery, B. Beretta, K. Wager-Smith et al., 1998 The cryb mutation identifies cryptochrome as a circadian photoreceptor in Drosophila., pp. 681–692 in Cell.

Stanewsky, R., K. S. Lynch, C. Brandes and J. C. Hall, 2002 Mapping of elements involved in regulating normal temporal period and timeless RNA expression patterns in Drosophila melanogaster., pp. 293–306 in J Biol Rhythms.

Stanewsky, R. R., C. F. C. Jamison, J. D. J. Plautz, S. A. S. Kay and J. C. J. Hall, 1997 Multiple circadian-regulated elements contribute to cycling period gene expression in Drosophila. EMBO Journal 16: 5006–5018.

Stanton-Geddes, J., C. G. de Freitas and C. D. Dambros, 2014 In defense of P values: comment on the statistical methods actually used by ecologists. Ecology 95: 637–642.

Sterne, J., 2003 Commentary: Null points - has interpretation of significance tests improved? International Journal of Epidemiology 32: 693–694.

Stodden, V., 2015 Reproducing Statistical Results. Annu. Rev. Stat. Appl. 2: 1–19.

Stodden, V., M. McNutt, D. H. Bailey, E. Deelman, Y. Gil et al., 2016 Enhancing reproducibility for computational methods. Science 354: 1240–1241.

Stoleru, D. D., Y. Y. Peng, J. J. Agosto and M. M. Rosbash, 2004 Coupled oscillators control morning and evening locomotor behaviour of Drosophila. Nature 431: 862–868.

Streit, S., C. W. Michalski, M. Erkan, J. Kleeff and H. Friess, 2009 Northern blot analysis for detection and quantification of RNA in pancreatic cancer cells and tissues. Nat Protoc 4: 37–43.

Stumpf, M. P., 2014 Approximate Bayesian inference for complex ecosystems. F1000Prime Rep 6: 60.

Subramanian, P., V. Prasanna, J. J. Jayapalan, P. S. Abdul Rahman and O. H. Hashim, 2014 Role of Bacopa monnieri in the temporal regulation of oxidative stress in clock mutant (cryb) of Drosophila melanogaster., pp. 37–44 in J Insect Physiol.

Sullivan, D. E., J. L. Gabbard, Jr., M. Shukla and B. Sobral, 2010 Data integration for dynamic and sustainable systems biology resources: challenges and lessons learned. Chem Biodivers 7: 1124–1141.

Sun, Y. Q., J. Y. Li, J. X. Liu, B. Y. Sun and C. Chow, 2014 An improvement of symbolic aggregate approximation distance measure for time series. Neurocomputing 138: 189–198.

Sunnaker, M., A. G. Busetto, E. Numminen, J. Corander, M. Foll et al., 2013 Approximate Bayesian Computation. PLOS Computational Biology 9.

Supp, S. R., D. N. Koons and S. K. M. Ernest, 2015a Using life history trade-offs to understand core-transient structuring of a small mammal community. Ecosphere 6.

Supp, S. R., F. A. La Sorte, T. A. Cormier, M. C. W. Lim, D. R. Powers et al., 2015b Citizen-science data provides new insight into annual and seasonal variation in migration patterns. Ecosphere 6.

Taniguchi, M., K. Miura, H. Iwao and S. Yamanaka, 2001 Quantitative assessment of DNA microarrays - Comparison with Northern blot analyses. Genomics 71: 34–39.

Tarantola, A., 2005 Inverse problem theory and methods for model parameter estimation. Society for Industrial and Applied Mathematics, Philadelphia, PA.

Tarantola, A., 2006 Popper, Bayes and the inverse problem. Nature Physics 2: 492–494.

Tarantola, A., and B. Valette, 1982 Inverse Problems = Quest for Information. Journal of Geophysics-Zeitschrift Fur Geophysik 50: 159–170.

Taroni, F., A. Biedermann and S. Bozza, 2016 Statistical hypothesis testing and common misinterpretations: Should we abandon p-value in forensic science applications? Forensic Science International 259.

Temme, C., L. B. Zhang, E. Kremmer, C. Ihling, A. Chartier et al., 2010 Subunits of the Drosophila CCR4-NOT complex and their roles in mRNA deadenylation. Rna-a Publication of the Rna Society 16: 1356–1370.

Tharyan, P., 1998 The relevance to meta-analysis, systematic reviews and the cochrane collaboration to clinical psychiatry. Indian J Psychiatry 40: 135–148.

The UniProt Consortium, 2017 UniProt: the universal protein knowledgebase. Nucleic Acids Res 45: D158–D169.

Theurkauf, W. E., H. Baum, J. Y. Bo and P. C. Wensink, 1986 Tissue-Specific and Constitutive Alpha-Tubulin Genes of Drosophila-Melanogaster Code for Structurally Distinct Proteins. Proceedings of the National Academy of Sciences of the United States of America 83: 8477–8481.

Toni, T., D. Welch, N. Strelkowa, A. Ipsen and M. P. Stumpf, 2009a Approximate Bayesian computation scheme for parameter inference and model selection in dynamical systems. J R Soc Interface 6: 187–202.

Toni, T., D. Welch, N. Strelkowa, A. Ipsen and M. P. H. Stumpf, 2009b Approximate Bayesian computation scheme for parameter inference and model selection in dynamical systems. Journal of the Royal Society Interface 6: 187–202.

Trafimow, D., and M. Marks, 2015 Editorial: banning null hypothesis significance testing procedures. Basic and Applied Social Psychology 37: 1–2.

Tressoldi, P. E., D. Giofre, F. Sella and G. Cumming, 2013 High Impact = High Statistical Standards? Not Necessarily So. PLOS ONE 8.

Turing, A. M., 1936 On computable numbers, with an application to the Entscheidungsproblem. Proc. Lond. Math. Soc. 42: 230–265.

Tyson, J. J., C. I. Hong, C. D. Thron and B. Novak, 1999 A simple model of circadian rhythms based on dimerization and proteolysis of PER and TIM., pp. 2411–2417 in Biophys J.

Ueda, H. R., M. Hagiwara and H. Kitano, 2001 Robust oscillations within the interlocked feedback model of Drosophila circadian rhythm. Journal of theoretical biology 210: 401–406.

Ueda, H. R., A. Matsumoto, M. Kawamura, M. Iino, T. Tanimura et al., 2002 Genome-wide transcriptional orchestration of circadian rhythms in Drosophila. Journal of biological chemistry 277: 14048–14052.

Umezaki, Y., T. Yoshii, T. Kawaguchi, C. Helfrich-Förster and K. Tomioka, 2012 Pigment-dispersing factor is involved in age-dependent rhythm changes in Drosophila melanogaster., pp. 423–432 in J Biol Rhythms.

Vaidya, A. T., D. Top, C. C. Manahan, J. M. Tokuda, S. Zhang et al., 2013 Flavin reduction activates Drosophila cryptochrome. Proceedings of the National Academy of Sciences of the United States of America 110: 20455–20460.

Van Gelder, R. N., H. Bae, M. J. Palazzolo and M. A. Krasnow, 1995 Extent and character of circadian gene expression in Drosophila melanogaster: identification of twenty oscillating mRNAs in the fly head., pp. 1424–1436 in Curr Biol.

Van Gelder, R. N., and M. A. Krasnow, 1996 A novel circadianly expressed Drosophila melanogaster gene dependent on the period gene for its rhythmic expression., pp. 1625–1631 in EMBO J.

van Gend, C., and J. L. Snoep, 2008 Systems biology model databases and resources. Essays Biochem 45: 223–236.

van Renssen, A. S. H. P., 2005 Gellish: A Generic Extensible Ontological Language - Design and Application of a Universal Data Structure. Delft University Press, Delft, Netherlands.

Vandenbroucke, I. I., J. Vandesompele, A. De Paepa and L. Messiaen, 2001 Quantification of splice variants using real-time PCR. Nucleic Acids Res 29.

VanGuilder, H. D., K. E. Vrana and W. M. Freeman, 2008 Twenty-five years of quantitative PCR for gene expression analysis. Biotechniques 44: 619–626.

Vaux, D. L., 2012 Know when your numbers are significant. Nature 492: 180–181.

Vilar, J. M. G., H. Y. Kueh, N. Barkai and S. Leibler, 2002 Mechanisms of noise-resistance in genetic oscillators. Proceedings of the National Academy of Sciences of the United States of America 99: 5988–5992.

Wager-Smith, K., and S. A. Kay, 2000 Circadian rhythm genetics: from flies to mice to humans. Nature Genetics 26: 23–27.

Wang, J., and T. Zhou, 2010 A computational model clarifies the roles of positive and negative feedback loops in the Drosophila circadian clock. Physics Letters A 374: 2743–2749.

Wang, T., and M. J. Brown, 1999 mRNA quantification by real time TaqMan polymerase chain reaction: Validation and comparison with RNase protection. Analytical Biochemistry 269: 198–201.

Wasserstein, R. L., and N. A. Lazar, 2016 The ASA’s Statement on p-Values: Context, Process, and Purpose. American Statistician 70: 129–131.

Weber, F. F., D. D. Zorn, C. C. Rademacher and H.-C. H. Hung, 2011 Post-translational timing mechanisms of the Drosophila circadian clock. Audio, Transactions of the IRE Professional Group on 585: 1443–1449.

Weeks, A. R., S. W. McKechnie and A. A. Hoffmann, 2007 Robust clines and robust sampling: a reply to Kyriacou et al., pp. 1652–1654 in J Evol Biol.

Weiss, R., O. Bartok, S. Mezan, Y. Malka and S. Kadener, 2014 Synergistic interactions between the molecular and neuronal circadian networks drive robust behavioral circadian rhythms in Drosophila melanogaster. PLOS Genet 10: e1004252.

White, E. P., E. Baldridge, Z. T. Brym, K. J. Locey, D. J. McGlinn et al., 2013 Nine simple ways to make it easier to (re)use your data. Ideas in Ecology and Evolution 6: 1–10.

Wierling, C., R. Herwig and H. Lehrach, 2007 Resources, standards and tools for systems biology. Brief Funct Genomic Proteomic 6: 240–251.

Wijnen, H., F. Naef, C. Boothroyd, A. Claridge-Chang and M. W. Young, 2006 Control of daily transcript oscillations in Drosophila by light and the circadian clock. PLOS Genetics 2: e39–e39.

Wilcox, R. R., 2002 Comparing the variances of two independent groups. Br J Math Stat Psychol 55: 169–175.

Wilcox, R. R., 2012 Introduction to Robust Estimation & Hypothesis Testing 3rd Edition. Academic Press (Elsevier), Amsterdam.

Wilkinson, D. J., 2012 Stochastic modelling for systems biology. CRC Press/Taylor & Francis, Boca Raton.

Wilkinson, M. D., M. Dumontier, I. J. Aalbersberg, G. Appleton, M. Axton et al., 2016 The FAIR Guiding Principles for scientific data management and stewardship. Sci Data 3: 160018.

Wilkinson, R. D., 2013 Approximate Bayesian computation (ABC) gives exact results under the assumption of model error. Stat Appl Genet Mol Biol 12: 129–141.

Willenbrock, H., J. Salomon, R. Sokilde, K. B. Barken, T. N. Hansen et al., 2009 Quantitative miRNA expression analysis: comparing microarrays with next-generation sequencing. RNA 15: 2028–2034.

Wolkenhauer, O., 2001 Systems biology: the reincarnation of systems theory applied in biology? Brief Bioinform 2: 258–270.

Woodger, J. H., A. Tarski and W. F. Floyd, 1937 The axiomatic method in biology. The University Press, Cambridge Eng.

Wooley, J. C., and H. S. Lin, 2005 Computational Modeling and Simulation as Enablers for Biological Discovery in Catalyzing Inquiry at the Interface of Computing and Biology, edited by J. C. Wooley and H. S. Lin, Washington (DC).

Woolston, C., 2015 Psychology journal bans P values. Nature 519: 9–9.

Wu, Q. Q., K. Smith-Miles and T. H. Tian, 2014 Approximate Bayesian computation schemes for parameter inference of discrete stochastic models using simulated likelihood density. Bmc Bioinformatics 15.

Wulbeck, C., E. Grieshaber and C. Helfrich-Forster, 2009 Blocking endocytosis in Drosophila’s circadian pacemaker neurons interferes with the endogenous clock in a PDF-dependent way. Chronobiol Int 26: 1307–1322.

Xie, Z., and D. Kulasiri, 2007 Modelling of circadian rhythms in Drosophila incorporating the interlocked PER/TIM and VRI/PDP1 feedback loops. Journal of theoretical biology 245: 290–304.

Xie, Z., D. Kulasiri, S. Samarasinghe and J. Qian, 2010 An Unbiased Sensitivity Analysis Reveals Important Parameters Controlling Periodicity of Circadian Clock. Biotechnology and Bioengineering 105: 250–259.

Xu, K., X. Zheng and A. Sehgal, 2008 Regulation of feeding and metabolism by neuronal and peripheral clocks in Drosophila., pp. 289–300 in Cell Metab.

Yerushalmi, S., and R. M. Green, 2009 Evidence for the adaptive significance of circadian rhythms. Ecology Letters 12: 970–981.

Yi, M., Y. Jia, Q. Liu, J. Li and C. Zhu, 2006 Enhancement of internal-noise coherence resonance by modulation of external noise in a circadian oscillator. Phys Rev E Stat Nonlin Soft Matter Phys 73: 041923.

Yin, H. S., H. G. Qi, J. W. Xu, W. N. N. Hung and X. Y. Song, 2014 Generalized Framework for Similarity Measure of Time Series. Mathematical Problems in Engineering.

Yoshii, T., T. Todo, C. Wülbeck, R. Stanewsky and C. Helfrich-Förster, 2008 Cryptochrome is present in the compound eyes and a subset of Drosophila’s clock neurons. The Journal of comparative neurology 508: 952–966.

Yoshii, T., S. Vanin, R. Costa and C. Helfrich-Förster, 2009 Synergic entrainment of Drosophila’s circadian clock by light and temperature., pp. 452–464 in J Biol Rhythms.

Young, M. W., and S. A. Kay, 2001 Time zones: a comparative genetics of circadian clocks. Nature reviews. Genetics 2: 702–715.

Yu, W., H. Zheng, J. H. Houl, B. Dauwalder and P. E. Hardin, 2006 PER-dependent rhythms in CLK phosphorylation and E-box binding regulate circadian transcription. Genes & development 20: 723–733.

Yu, W. W., J. H. J. Houl and P. E. P. Hardin, 2011 NEMO Kinase Contributes to Core Period Determination by Slowing the Pace of the Drosophila Circadian Oscillator. Current biology: CB 21: 6–6.

Zak, D. E., F. J. Doyle, D. G. Vlachos and J. S. Schwaber, 2001 Stochastic kinetic analysis of transcriptional feedback models for circadian rhythms. Proceedings of the 40th Ieee Conference on Decision and Control, Vols 1-5: 849–854.

Zeeberg, B. R., J. Riss, D. W. Kane, K. J. Bussey, E. Uchio et al., 2004 Mistaken identifiers: gene name errors can be introduced inadvertently when using Excel in bioinformatics. BMC Bioinformatics 5: 80.

Zeigler, B. P., 2012 Guide to modeling and simulation of systems of systems. Springer, New York.

Zeigler, B. P., and P. E. Hammonds, 2007 Modeling & simulation-based data engineering: introducing pragmatics into ontologies for net-centric information exchange. Academic, Burlington, MA.

Zeigler, B. P., H. Praehofer and T. G. Kim, 2000 Theory of modeling and simulation: integrating discrete event and continuous complex dynamic systems. Academic Press, San Diego.

Zeng, H., P. E. Hardin and M. Rosbash, 1994 Constitutive overexpression of the Drosophila period protein inhibits period mRNA cycling., pp. 3590–3598 in EMBO J.

Zerr, D. M., J. C. Hall, M. Rosbash and K. K. Siwicki, 1990 Circadian fluctuations of period protein immunoreactivity in the CNS and the visual system of Drosophila., pp. 2749–2762 in J Neurosci.

Zhang, R., N. F. Lahens, H. I. Ballance, M. E. Hughes and J. B. Hogenesch, 2014a A circadian gene expression atlas in mammals: implications for biology and medicine. Proc Natl Acad Sci U S A 111: 16219–16224.

Zhang, Z., W. Zhu and J. Luo, 2014b Bringing biocuration to China. Genomics Proteomics Bioinformatics 12: 153–155.

Zheng, X., K. Koh, M. Sowcik, C. J. Smith, D. Chen et al., 2009 An isoform-specific mutant reveals a role of PDP1 epsilon in the circadian oscillator., pp. 10920–10927 in J Neurosci.

Zheng, X., M. Sowcik, D. Chen and A. Sehgal, 2014 Casein kinase 1 promotes synchrony of the circadian clock network., pp. 2682–2694 in Mol Cell Biol.

Zielinski, T., A. M. Moore, E. Troup, K. J. Halliday and A. J. Millar, 2014 Strengths and Limitations of Period Estimation Methods for Circadian Data. PLOS ONE 9.

Ziemann, M., Y. Eren and A. El-Osta, 2016 Gene name errors are widespread in the scientific literature. Genome Biol 17: 177.

Zinn, K., D. Dimaio and T. Maniatis, 1983 Identification of 2 Distinct Regulatory Regions Adjacent to the Human Beta-Interferon Gene. Cell 34: 865–879.

Zordan, M. A., and F. Sandrelli, 2015 Circadian clock dysfunction and psychiatric disease: could fruit flies have a say? Frontiers in Neurology 6.

Zweig, K. A., 2016 Section 11.8 Data responsibility and Data hygiene, pp. 350–361 in Network analysis literacy: a practical approach to the analysis of Networks, edited by K. A. Zweig. Springer Berlin Heidelberg, New York, NY.

Zwiebel, L. J., P. E. Hardin, J. C. Hall and M. Rosbash, 1991a Circadian oscillations in protein and mRNA levels of the period gene of Drosophila melanogaster., pp. 533–537 in Biochem Soc Trans.

Zwiebel, L. J., P. E. Hardin, X. Liu, J. C. Hall and M. Rosbash, 1991b A Posttranscriptional Mechanism Contributes to Circadian Cycling of a Per-Beta-Galactosidase Fusion Protein. Proceedings of the National Academy of Sciences of the United States of America 88: 3882–3886.

